# Bovine in vitro blastocysts with distinct morphokinetic patterns show transcriptomic differences at genome activation

**DOI:** 10.64898/2026.06.28.733532

**Authors:** Alline de Paula Reis, Mounya Belghiti, Ludivine Laffont, Sylvie Ruffini, Catherine Archilla, Nadège Le Brusq, Audrey Teste, Brigitte Marquant-LeGuienne, Eugénie Canon, Luc Jouneau, Yan Jaszczyszyn, Andrew A. Ponter, Mélanie Bernard Caciarella, Jean Unrug, Eve-Marie Stamler, Véronique Duranthon, Alain Trubuil

## Abstract

Bovine embryo in vitro production (IVP) is characterised by low efficiency and variable outcomes. Monitoring early embryonic development by videomicroscopy revealed substantial morphokinetic heterogeneity in the first four embryonic cycles (EC, conventionally referred to as the 2-, 4-, 8- and 16-cell stages). Morphokinetic analysis offers a promising approach to characterize divergent developmental trajectories and provide a better understanding of underlying molecular mechanisms. We developed a Random Forest classification system (Bovine Embryo Analyser based on Morphokinetics: BEAM) to predict embryo phenotype. It is based on morphokinetic variables collected from the 1st to the 4th EC and predicts four blastocyst categories (EHB: Early Hatching Blastocyst, HB: Hatching Blastocyst, SSB: Subtle Developmental Shift Blastocyst, ADB: Arrhythmic Development Blastocyst). Classification performance on an independent dataset was good (F1 score = 0.59; Accuracy = 0.78), indicating that the BEAM can be useful for embryo development studies. The BEAM was further applied to embryos submitted to 4.3 days of culture and having completed the 4th EC (16-32 cells). RNA sequencing was performed on sixteen samples (4 x 8 pooled embryos/category). The ADB category was significantly enriched in transcripts involved in the regulation of transcriptional activity compared to the EHB category. In addition, in the ADB category, 22.7% (n = 185/816) of the upregulated genes were of maternal origin while only 6.2% were of embryonic origin (n = 54/816). In conclusion, despite being at a comparable developmental stage and transcriptionally competent, the ADB category showed delayed maternal transcript degradation suggesting delayed transition to embryonic transcriptional autonomy.

**Illustrated Abstract:** 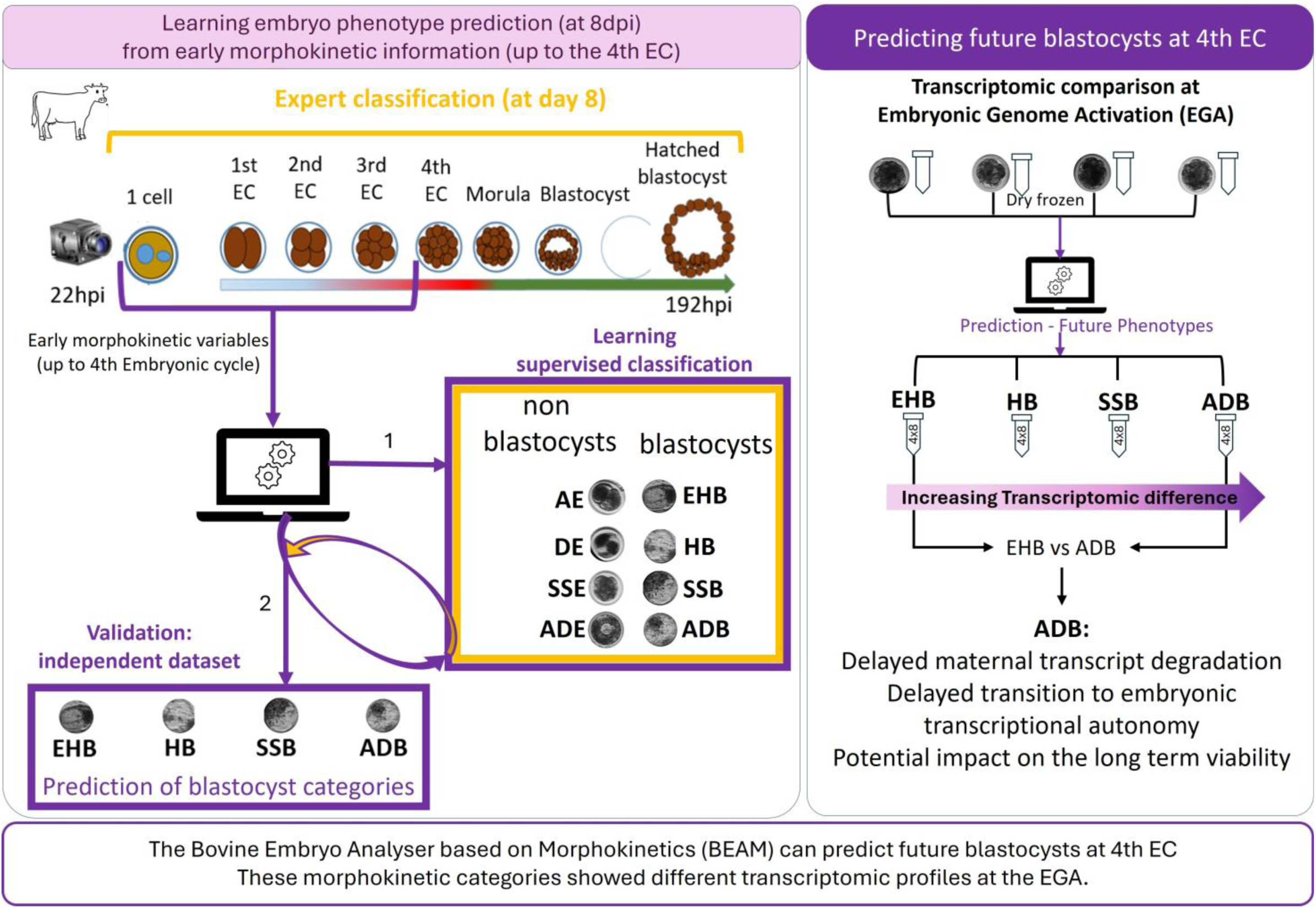

**Summary Sentence:** In vitro produced bovine embryo morphokinetic patterns allow prediction of four blastocyst categories (up to the 4^th^ embryonic cycle) and are associated with distinct transcriptomic profiles at embryonic genome activation in competent embryos.

## Introduction

Bovine embryo in vitro production (IVP) has developed significantly and, since 2018, its field use (embryo production and transfer to a female recipient) has exceeded that of in vivo derived embryos [1]. Nonetheless, the overall efficiency of this technique remains low and variable, with higher pregnancy and neonatal losses compared to pregnancies from in vivo developed embryos [2]. Consequently, enhancing the overall performance of bovine IVP procedures will necessitate improvements in the production process and the selection of viable embryos for transfer.

In this context, morphokinetics is a promising tool to improve our understanding of the mechanisms involved in early embryonic development and could contribute to improving the production processes and decision-making for the selection of embryos prior to transfer. In human medicine, the availability of large dataset cohorts of embryos combining morphokinetic data and clinical outcomes has enabled the development of deep learning approaches to predict implantation potential with moderate discriminatory power [3]. However, currently the usefulness of morphokinetics to predict the possibility to obtain a birth has been questioned [4, 5, 6].

In cattle, only Sugimura et al. [7] assessed the ability of morphokinetic variables to predict the establishment of pregnancy after a small number of embryo transfers (n = 52). The best results were obtained by associating morphokinetic variables and oxygen consumption during in vitro culture. However, the limited availability of large bovine datasets combining morphokinetics and pregnancy outcomes restricts the direct transposition of the approaches used in human medicine. Other studies in cattle aimed at distinguishing blastocysts and non-blastocysts either relied on supervised deep learning directly from images [8] or conventional statistical methods [9], [10]. Also, Yaacobi-Artzi et al. [10] showed that morphokinetics is not correlated with the gold standard morphological classification of the International Embryo Technology Society (IETS) grading blastocysts into “Excellent or good” (Code 1), “Fair” (Code 2), “Poor” (Code 3) and “Dead or degenerating” (Code 4) based on the static observation of the Inner Cell Mass and Trophectoderm, usually at day seven of development [11], [12].

We observed that blastocysts, even those with identical IETS grades, can develop through a wide range of morphokinetic patterns, in particular during the first four embryonic cycles (ECs) - conventionally referred to as the 2-, 4-, 8- and 16-cell stages. Furthermore, blastocyst hatching times can vary by up to 24 hours. This suggests that it may be possible to predictively classify blastocysts into different morphokinetic phenotypes according to their developmental patterns during the very first ECs, independently of their IETS morphological grade. Such an innovative approach would allow us to assess the biological information of future blastocysts of different morphokinetic phenotypes prior to blastocyst formation. It may reveal early biological heterogeneity among embryos competent to form blastocysts and, in the future, improve understanding of the sources of performance variation of IVP embryos. We therefore hypothesise that an automated classification system could be used to predictively classify embryos into a small number of blastocyst phenotypes (n=4) based on the numerous morphokinetic variables measured up to the 4^th^ EC. Secondly, we hypothesise that this predictive classification, obtained around the completion of major Embryo Genome Activation (EGA), could be used to study intermediary stages (16-32 cells, morula) to explore biological differences between competent embryos, including at EGA.

The objectives of this study were to develop an automated morphokinetic classification system based on early development morphokinetic variables, enabling the predictive classification of embryos as early as the 4^th^ EC; and to compare the transcriptomic profiles of embryos predicted to belong to one of the four proposed blastocyst morphokinetic categories, immediately following the presumed major wave of embryonic genome activation (EGA) (i.e. at the end of the 4^th^ EC), to evaluate the extent to which morphokinetics can reveal molecular and functional differences among embryos at a similar developmental stage.

## Materials and methods

### Ethical statement

The study was conducted using ovaries collected post-mortem in commercial slaughterhouses and frozen-thawed semen collected from bulls at breeding stations whose commercial activities are duly regulated by the relevant ministry in France. No ethical committee experimental authorisation was required.

### 1. Development of a semi-automated morphokinetic classification system enabling the prospective classification of bovine embryos

#### Bovine embryo in vitro production and image acquisition

Cumulus oocyte complex (COC) collection and in vitro maturation (IVM) were performed as described by [13]. Briefly, COCs were aspirated from follicles 2 to 8 mm in diameter from bovine ovaries collected at a slaughterhouse and transported in flasks containing Euroflush media for 2 hours in an insulated box at 34°C.

The harvested COCs were rinsed in an embryo collection medium (Euroflush; IMV Technologies) and in vitro matured in groups of 50-60, placed in 500 µL of TCM 199 (Sigma) supplemented with 10% foetal calf serum (FCS, Invitrogen), 10 μg/ml FSH (Folltropin-V; Reprobiol) and LH (Lutropin-V; Reprobiol), 1 μg/ml oestradiol (Sigma), 1 μg/ml EGF (Epidermal Growth Factor; Sigma) and 50 μg/ml gentamicin for 22 hours at 38.5°C in 5% CO₂ in humidified air.

Frozen semen from four Holstein bulls was used. Following thawing for 30 seconds at 37°C, motile spermatozoa were selected using a BoviPure® / Bovidilute® discontinuous gradient (40/80) according to the manufacturer’s instructions (Nidacon International AB, Gothenburg, Sweden).

Groups of 50 COCs were placed in 500 µl fertilisation medium containing 10 µg/ml heparin, 20 µM penicillamine, 10 µM hypotaurine and 1 µM epinephrine [14] with 1×10^6^ spermatozoa/ml and were incubated at 38.5°C, 5% CO_2_ in humidified air for 22 hours.

The moment of gamete contact was considered as the initiation of embryonic development. At 22h ± 2h after gamete contact, the COCs were denuded and the presumed zygotes were placed in semi-individual culture until 8 days post gamete contact (_dpi_) in 16 microwell dishes covered with a single 120 µl drop of synthetic oviductal fluid medium (SOF; Minitub/Inventio) supplemented with 1% heifer serum collected 3 days after oestrus (SVJ3, homemade) covered with 3 mL of mineral oil (Origio, ART-4008-5) and equilibrated overnight in an atmosphere of 5% CO_2_, 5% O_2_ and 90% N_2_ in humidified air at 38.5°C.

The incubator was equipped with a video microscope (PrimoVision®). Images were acquired at 15-minute intervals throughout the culture and compiled into time-lapse sequences using the dedicated PrimoVision® analysis software.

#### Morphokinetic classification and supervised learning

##### Expert annotation and Dataset

The morphokinetic variables (Table 1) were annotated by an expert, following the recommendations of the annotation guidelines developed in our laboratory by Le Brusq [15] (an extract of the guidelines is translated and supplied in Supplementary Material 1). Within the 8 days of culture, the expert assigned a label to each embryo corresponding to a morphokinetic category, which grouped together similar morphokinetic phenotypes:

- **Degenerated embryo (DE):** embryo showing no cellular activity and no intact cell membrane at 8_dpi_.
- **Arrested embryo (AE):** an embryo that shows signs of life characterised by the presence of cytoplasmic particle movement and at least one cell with an intact membrane. However, at 8_dpi_, the embryo no longer exhibits signs of cleavage.
- **Arrhythmic development embryo (ADE):** embryo exhibiting at least one cell presenting abnormal cleavage: direct cleavage (1 blastomere resulting in 3 or more blastomeres) or reverse cleavage (fusion of blastomeres) or their combination during at least one of the first three EC, associated with marked developmental delay (completion of the fourth EC beyond 4.3_dpi_), with or without cell degeneration during the first three EC.
- **Arrhythmic development blastocyst (ADB):** embryo exhibiting at least one cell presenting abnormal cleavage: direct cleavage (1 blastomere resulting in 3 or more blastomeres) or reverse cleavage (fusion of blastomeres) or their combination during at least one of the first three EC, without developmental delay (completion of the fourth EC up to 4.3_dpi_) and without cell degeneration during the first three EC.
- **Subtle developmental shift embryo (SSE):** embryo exhibiting morphologically normal cleavage (1 blastomere resulting in 2 blastomeres) during the first three EC, associated with a slight disturbance in development in at least one EC (i.e. slightly lengthened EC or lengthened cytokinesis, or combination of both) and developmental delay, defined as completion of the fourth EC beyond 4.3_dpi_, with or without cellular degeneration during the first three EC.
- **Subtle developmental shift blastocyst (SSB):** embryo with morphologically normal cleavage (1 blastomere resulting in 2 blastomeres) during the first three EC, associated with a slight disturbance in development in at least one EC (i.e. slightly lengthened EC or lengthened cytokinesis, or combination of both), without developmental delay, defined as completion of the fourth EC up to the fourth EC up to 4.3_dpi_) and without cellular degeneration during the first three EC.
- **Hatching blastocyst (HB):** embryo with morphologically normal cleavage (1 blastomere resulting in 2 blastomeres) during the first three EC, without detectable developmental disturbance or delay (completion of the fourth EC up to 4.3_dpi_), without cellular degeneration during the first three EC and Lag Phase (defined as the prolonged interphase commonly observed after the completion of the third EC) and reaching the hatching stage between 7.5 and 8_dpi_.
- **Early hatching blastocyst (EHB):** embryo showing morphologically normal cleavage (1 blastomere resulting in 2 blastomeres) during the first three EC, with no detectable developmental disturbance or delay (completion of the fourth EC up to 4.3_dpi_), without cellular degeneration during the first three EC and Lag Phase and reaching the hatching stage rapidly (before 7.4_dpi_).

**Table 1:** Morphological and kinetic variables described by LeBrusq (2018). Only variables observed between the presumed zygote stage and the end of the Lag Phase were used for the learning process.

| Morphokinetic variables |  |  | Developmental stage (embryonic cycle - EC) |  |  |  |  |  |  |
| --- | --- | --- | --- | --- | --- | --- | --- | --- | --- |
| Type | Number | Name | Zygote | 1st EC | 2nd EC | 3rd EC | Lag Phase | 4th EC | Blastocyst |
| Morphologic variables | 1 | Second polar body extrusion observed (Yes/No) | X |  |  |  |  |  |  |
|  | 2 | Presence of vacuole-like structures | X | X |  |  |  |  |  |
|  | 3 | Initiation of the cleavages of an embryonic cycle (Yes/No) |  | X | X | X |  | X |  |
|  | 4 | Cleavage pattern (normal (1-2 cells); abnormal: (1-3, 1-4; reverse cleavage) |  | X | O | O |  |  |  |
|  | 5 | Total cell number at the end of the EC |  | X | X | X |  |  |  |
|  | 6 | Number of degenerated cells during the EC |  | X | X | X | X |  |  |
|  | 7 | Plasma membrane dynamics during cell cleavage |  | X | X |  |  |  |  |
|  | 8 | Cellular asymetry |  | X |  |  |  |  |  |
|  | 9 | Cleavage axis orientation (visual assessment) |  |  | X |  |  |  |  |
|  | 10 | Presence of fragmentation (Yes/No) |  | X |  |  |  |  |  |
|  | 11 | Reaching the Hatching stage (Yes/No) |  |  |  |  |  |  | O |
|  | 12 | Signs of life (particle movements and membrane integrity) | O | O | O | O | O | O | O |
| Kinetic variables | 13 | Timing of second polar body extrusion | O |  |  |  |  |  |  |
| | 14 | Initiation ($t_{ie}$) and completion ($t_{fe}$) of cytokinesis of the embryonic cycle | | X | X | X | | O | |
|  | 15 | Duration of the cytokinesis of each cell |  | X |  |  |  |  |  |
|  | 16 | Duration of the cytokinesis of the embryonic cycle |  | X | X | X |  | O |  |
|  | 17 | Synchronisation of cytokinesis within an embryonic cycle |  |  | X | X |  | X |  |
| | 18 | Initiation ($t_{iec}$) and completion ($t_{fec}$) of the embryonic cycle | | X | X | X | | X | |
|  | 19 | Duration of an embryonic cycle (EC) |  | X | X | X |  | O |  |
| | 20 | Initiation ($t_{il}$) and completion ($t_{fl}$) of the Lag Phase | | | | | X | | |
|  | 21 | Duration of the Lag Phase |  |  |  |  | X |  |  |
| | 22 | Initiation ($t_{ih}$) of hatching | | | | | | | O |

Note: X: variable used for the machine learning process; O: additional variable used by the expert to perform expert annotation.

Figure 1 provides an illustrative representation of this expert classification.

Following expert annotation, the dataset was compiled (Table 2). It was initially unbalanced and was artificially balanced through random duplication of individuals belonging to less represented classes, as commonly performed in medical research dealing with unbalanced data [16]. In addition, to reduce the number of morphokinetic categories and thereby improve the learning performance of the model, classes exhibiting similar early morphokinetic characteristics with divergent phenotypes but easily discernible by the expert (SSB and SSE categories; ADB and ADE categories) were merged as shown in Table 2. The distinction between the transferable (SSB or ADB) and non-transferable (SSE or ADE) embryos can be easily performed by an expert following the last step of the expert classification tree (Figure 1).

**Figure 1:**
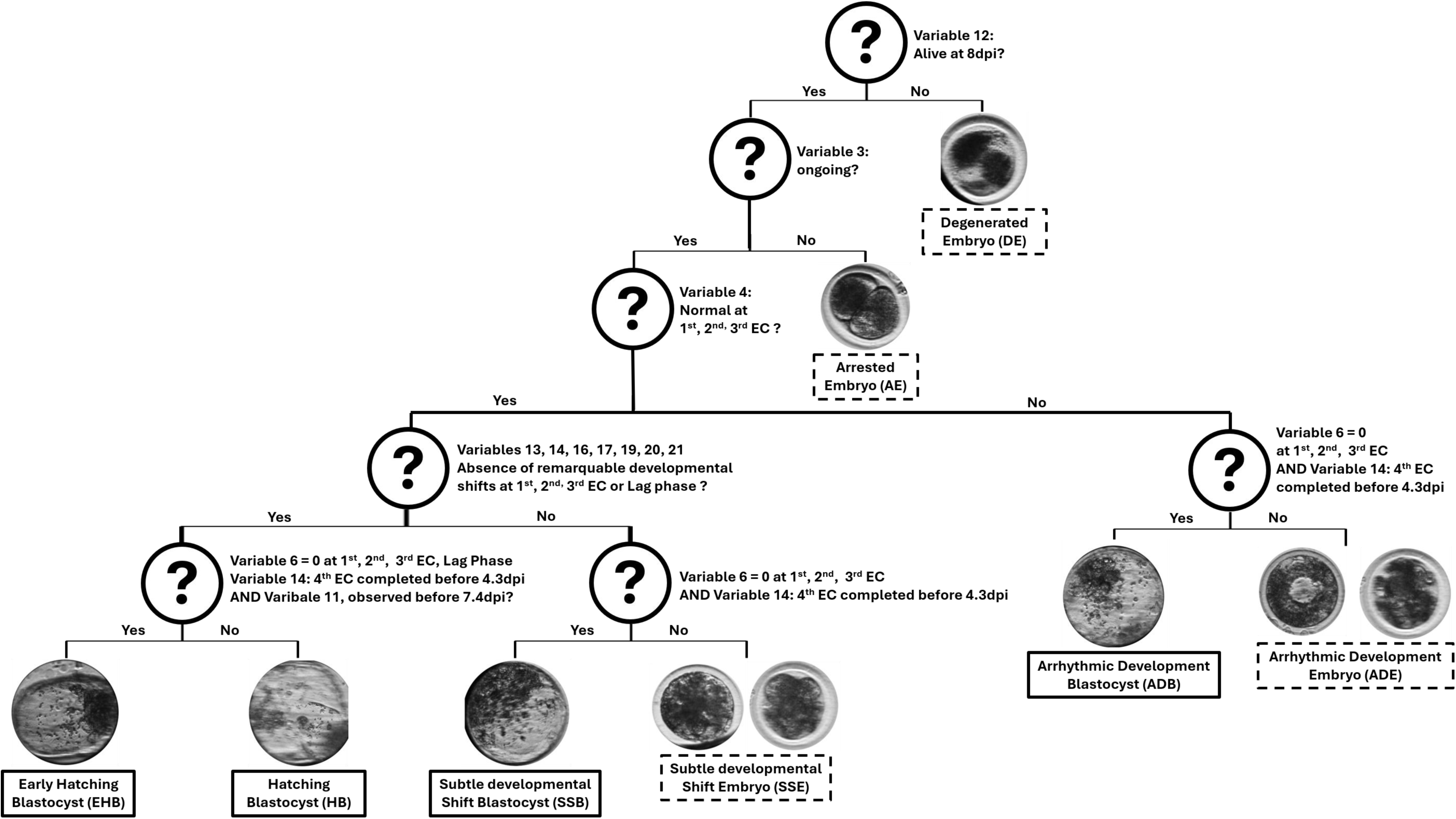
Visual expert classification of bovine IVP embryos into morphokinetic categories: Degenerated embryo (DE); Arrested embryo (AE); Arrhythmic Development Embryo (ADE); Arrhythmic Development Blastocyst (ADB); Subtle developmental Shift Embryo (SSE); Subtle developmental Shift Blastocyst (SSB); Hatching blastocyst (HB); Early hatching blastocyst (EHB). Transferable embryos are surrounded by a solid outline. Non-transferable embryos are surrounded by a dotted outline. The list of variables is supplied in Table 1.

**Table 2:** Bovine IVP embryo cohorts included in the learning process (Learning Dataset) and the independent test (Independent Test Dataset). The model was trained to predict six labels: EHB: Early hatching blastocyst, HB: Hatching blastocyst, SSE + SSB: resulting from grouping Subtle development shift embryo (SSE) and Subtle development shift blastocyst (SSB), ADE + ADB: resulting from grouping Arrhythmic Development Embryo (ADE) and Arrhythmic Development Blastocyst (ADB), Arrested Embryo (AE) and Degenerated Embryo (DE).

| Morphokinetic category | Learning embryo dataset |  |  |  |  |  | Independent Test Dataset n |
| --- | --- | --- | --- | --- | --- | --- | --- |
|  | Original n | Randomly Duplicated n | Learning morphokinetic category after merging | Total n | Train n | Test n |  |
| Early Hatching Blastocyst (EHB) | 11 | 37 | EHB | 48 | 38 | 10 | 19 |
| Hatching Blastocyst (HB) | 28 | 33 | HB | 61 | 51 | 10 | 15 |
| Subtle developmental Shift Blastocyst (SSB) | 29 | 0 | SSE + SSB | 31 | 20 | 11 | 10 |
| Subtle developmental Shift Embryo (SSE) | 2 | 0 |  |  |  |  | – |
| Arrhythmic Development Blastocyst (ADB) | 3 | 0 | ADE + ADB | 38 | 27 | 11 | 10 |
| Arrhythmic Development Embryo (ADE) | 35 | 0 |  |  |  |  | – |
| Arrested Embryo (AE) | 17 | 29 | AE | 46 | 35 | 11 | – |
| Degenerated Embryo (DE) | 46 | 0 | DE | 46 | 35 | 11 | – |
| Total | 171 | 99 | Total | 270 | 206 | 64 | 54 |

#### Learning process

Training was performed using the Random Forest algorithm [17] in the randomForest package of R [18] with a random resampling strategy with replacement (bootstrap) [19] to limit the risk of overfitting [16]. The Random Forest model consisted of 500 trees, and at each split, a random subset of predictors was considered to determine the optimal node. Final predictions were obtained by a majority vote across all trees. A 10-fold cross validation was performed using the caret package of R [20].

The performance of the model was assessed via the Sensitivity (True Positive Rate (TPR) or Recall), Specificity (or True Negative Rate (TNR)), Positive Predictive Value (PPV or Precision), Negative Predictive Value (NPV), F1-score (an indicator that combines Precision and Sensitivity into a single metric) and Accuracy (the proportion of correct predictions out of all predictions) [21].

The importance of the variables was calculated using the Mean Decrease in Impurity (MDI) and Mean Decrease in Accuracy (MDA) measures from Random Forest [17] and the predictive variables were automatically selected with the Variable Selection Using Random Forests – V-SURF – R VSURF package, as described by [22].

#### Test: prediction under field conditions

After learning and selection of the predictive variables, a test was performed to assess how well the model could discriminate between the future types of blastocysts before the phenotype observation on an independent dataset.

A new, independent, dataset (Table 2) was produced under the conditions described previously, except for:

- the COCs were collected from slaughterhouse ovaries of Prim’Holstein, Montbeliard or Charolais cows, without distinction, with a transport duration of 3 hours instead of 2 hours;
- a new batch of media, including ingredients from different production batches;
- only the predictive variables defined by the learning process were used by the classification system.

The previously described performance indicators were used to assess the prediction performance on the field simulation sample.

### 2. Comparison of transcriptomic profiles among four morphokinetic competent categories (EHB, HB, SSB, ADB) immediately after embryonic genome activation (EGA)

#### Sample production

A total of six embryo production sessions and time-lapse imaging were performed as described but embryo culture was stopped at 4.3_dpi_. Only embryos that were considered by the classification system as belonging to the EHB, HB, SSB or ADB categories and had completed the 4^th^ EC before 4.3 (Variable 21, Table 1) were included in the study, since these embryos will go on to form a blastocyst (Figure 1).

#### RNA extraction, library production and sequencing

On completion of embryo culture, the embryos were rinsed three times in PBS and placed individually in 0.5 mL RNase-free Eppendorf tubes with a minimum amount of medium (<2 µL), immediately freeze-dried and stored at −80°C. Sixteen samples comprising eight embryos were prepared and named as follows: (EHB category: EHB-1, EHB-2, EHB-3, EHB-4), (HB category: HB-1, HB-2, HB-3, HB-4), (SSB category: SSB-1, SSB-2, SSB-3, SSB-4), (ADB category: ADB-1, ADB-2, ADB-3 and ADB-4). To eliminate session and/or bull effects, the different sessions and bulls were equally represented within the samples analysed.

Total RNA (RNAtot) from the 16 samples was extracted using the ArcturusTM PicoPure kit and purified using RNase-Free DNase Set (Qiagen), according to the supplier’s instructions. The quality of RNAtot was assessed by reading the RIN (RNA Integrity Number) [23] and the amount was estimated with the RNA 6000 Pico Kit (Agilent). All samples had more than 500 pg of RNAtot and a RIN > 8.

Due to the low amounts of material, cDNA was prepared starting from 0.5 ng total RNA using the SMART seq v4 ultra-low input RNA kit from Takarabio (www.takarabio.com, #634888). RNA-Seq libraries were prepared from 0.15 ng cDNA using the Nextera XT Illumina library preparation kit. All libraries were subjected to sequencing (paired-end, 50–34 bp) using a NextSeq 500 instrument at the I2BC High-throughput Sequencing Platform. Each sample yielded over 75 million paired-end reads. Demultiplexing and adapter removal were performed using bcl2fastq 2-2.18.12 and Cutadapt 1.15. The quality of the raw RNA-Seq data was assessed using FastQC v0.11.5. Only reads longer than 10 bp were used for subsequent analysis.

#### Data processing and analysis of results

Fastq files were mapped to the transcriptome reference genome (Ensembl 95, Bos taurus ARS-UCD1.2) using Tophat (v2.1.1) [24]. Tables of counts were generated with featureCounts (v1.6.0) [25]. Analysis of differential expression was made using the DESeq2 package (v1.28.1) [26] and raw p-values were corrected for multiple testing using Benjamini-Hochberg procedure [27]. Genes were considered as differentially expressed (DEG) if adjusted P-value <0.05 and |log₂FC| > 1. The results were illustrated in Venn diagrams.

A descriptive analysis was performed using Hierarchical clustering based on correlation [28] and Principal Component Analysis (PCA) [29].

Enrichment analysis of biological functions and mammalian phenotypes was performed on the Enrichr platform [30, 31]. The platform was accessed in September 2025. The KEGG Mouse, Gene Ontology – Biological Processes, Molecular Functions, Cellular Components, and MGI mammalian phenotypes libraries were explored.

Only *pathways* with a q-value (False Discovery Rate, FDR) below 0.05 were retained. Differentially expressed gene profiles were determined using the GSE52415 dataset [32], including oocyte (germinal vesicle and metaphase II) and embryonic (4-, 8-, 16-cell and blastocyst) stages. FASTQ files were processed as described above, and genomic annotations were retrieved from Ensembl (release 92) for *Bos taurus* (ARS-UCD1.2).

Transcripts were classified as maternal when median expression in GV or MII-oocytes was >5 reads and decreased after further stages, embryonic when median MII-oocyte was ≤5 reads but >5 reads in at least one embryonic stage or mixed otherwise. A transcript with a median of zero across all stages was considered “not represented”.

## Results

### The automated classification system based on morphokinetics distinguishes between four categories of blastocysts

An expert can reliably determine the blastocyst phenotype only once it is expressed, i.e., from 7 to 8 days of culture for blastocysts. The automated classification system developed in this study is called BEAM (Bovine Embryo Analyser based on Morphokinetics) (Figure 2). It takes advantage of divergent morphokinetic trajectories from the earliest stages that ultimately lead to either transferable (blastocysts) or non-transferable embryos with a double purpose: to structure these diverse trajectories into a limited number of morphokinetic categories, thereby reducing subjectivity through standardised annotation and classification, and 2) to carry out the classification of four categories of blastocysts predictively (by the end of 4^th^ EC, i.e.: before the phenotype is observed). This predictive property shifts the paradigm from observing a phenotype once it is established to being able to predict it in advance and provide a unique analytical framework to better characterise the biological mechanisms underlying this diversity.

**Figure 2:**
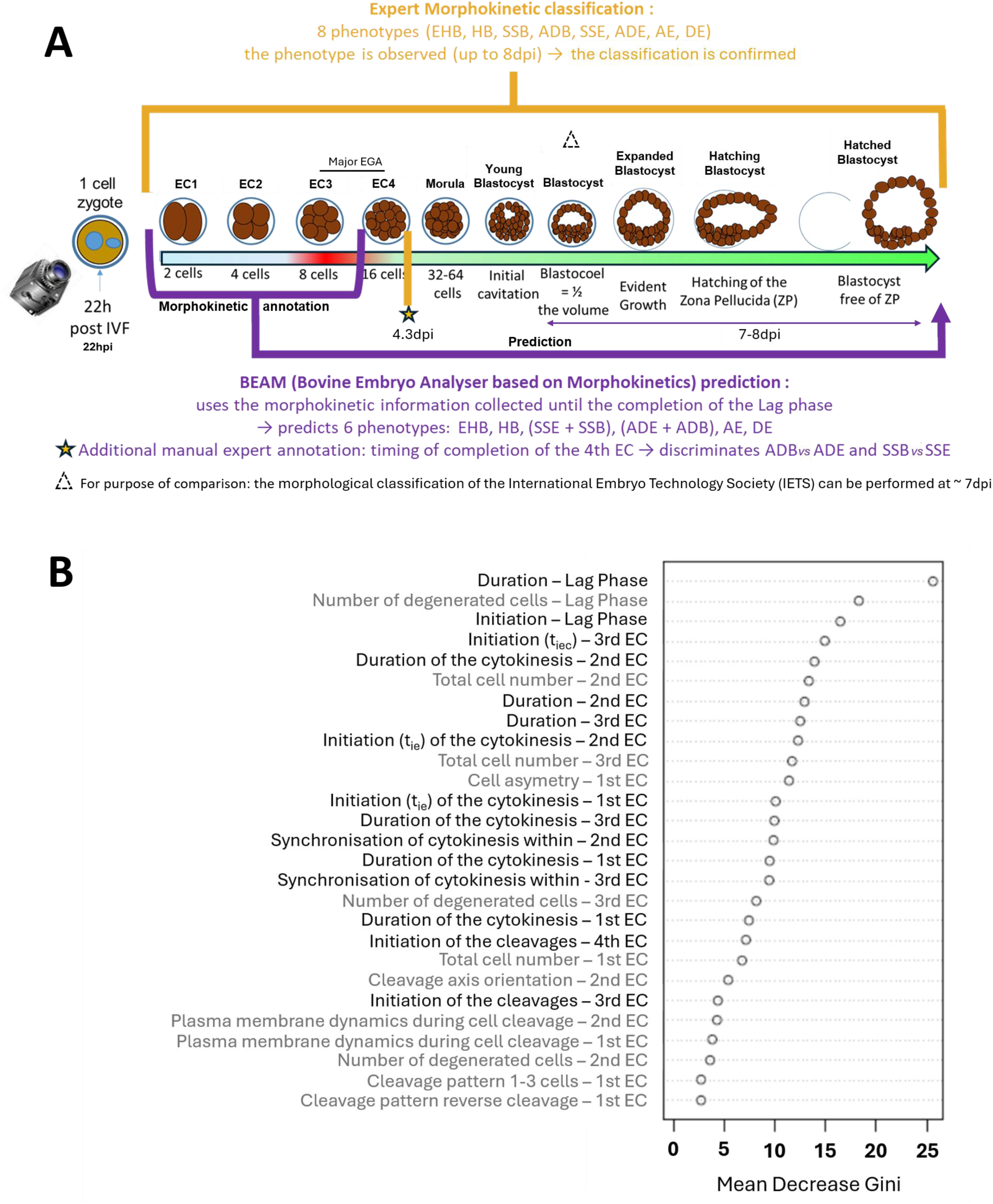
Principle of the Bovine Embryo Analyser based on Morphokinetics (BEAM) learned from expert-labelled embryos (A) and the predictive kinetic (black) and morphological (grey) variables used to classify embryos into the morphokinetic categories (B). Note: EHB: Early Hatching Blastocysts; HB: Hatching Blastocysts; SSE + SSB: Subtle Developmental Shift Embryos + Subtle Developmental Shift Blastocysts; ADE + ADB: Arrhythmic Development Embryos + Arrhythmic Development Blastocysts; AE: Arrested Embryos; DE: Degenerated Embryos.

The learning dataset comprised 270 embryos, including 171 original embryos and 99 from the data augmentation (Table 2), for which fifty-four morphokinetic variables were annotated from the 1^st^ to the 4^th^ EC, in accordance with the recommendations set out in our morphokinetic annotation guide [15].

The validation of this stochastic classifier over 10 runs showed high performance (F1-score: 0.80), albeit with some variability shown by a standard deviation of 0.07 (Table 3). A set of 15 kinetic and 12 morphologic predictive variables was identified over the first four EC (including the Lag Phase) (Figure 2).

**Table 3:** Performance of the BEAM morphokinetic classification system obtained during the learning process (Top) and in predicting the assignment of embryos at the completion of the 4th EC to one of the four blastocyst categories: EHB, HB, SSB, ADB obtained after the independent field test (Bottom).

| Learning dataset | Performance |  |  |  |  |  |  |  |
| --- | --- | --- | --- | --- | --- | --- | --- | --- |
|  | MK category | N | Sensitivity (Recall) | Specificity | PPV (Precision) | NPV | F1-score | Accuracy |
| | EHB | 10 | 0.96 ± 0.06 | 0.97 ± 0.02 | 0.88 ± 0.07 | 0.99 ± 0.01 | 0.92 ± 0.05 | 0.97 ± 0.02 |
| | HB | 10 | 0.93 ± 0.07 | 0.96 ± 0.02 | 0.85 ± 0.09 | 0.99 ± 0.1 | 0.88 ± 0.05 | 0.95 ± 0.02 |
| | SSE + SSB | 11 | 0.64 ± 0.11 | 0.97 ± 0.03 | 0.86 ± 0.15 | 0.93 ± 0.02 | 0.72 ± 0.09 | 0.89 ± 0.03 |
| | ADE + ADB | 11 | 0.86 ± 0.12 | 0.98 ± 0.02 | 0.89 ± 0.09 | 0.97 ± 0.02 | 0.87 ± 0.07 | 0.96 ± 0.03 |
| | AE | 11 | 0.81 ± 0.13 | 0.94 ± 0.03 | 0.73 ± 0.09 | 0.96 ± 0.03 | 0.76 ± 0.06 | 0.92 ± 0.03 |
| | DE | 11 | 0.67 ± 0.15 | 0.94 ± 0.03 | 0.72 ± 0.11 | 0.93 ± 0.03 | 0.69 ± 0.01 | 0.89 ± 0.04 |
|  | Global | 64 | **0.80 ± 0.10** | **0.96 ± 0.02** | **0.82 ± 0.10** | **0.96 ± 0.02** | **0.80 ± 0.07** | **0.93 ± 0.01** |
| Independent test dataset | Performance |  |  |  |  |  |  |  |
|  | MK category | N | Sensitivity (Recall) | Specificity | PPV (Precision) | NPV | F1-score | Accuracy |
|  | EHB | 19 | 0.37 | 0.94 | 0.78 | 0.73 | 0.50 | 0.74 |
|  | HB | 15 | 0.60 | 0.72 | 0.45 | 0.83 | 0.51 | 0.69 |
|  | SSB | 10 | 0.50 | 0.84 | 0.42 | 0.88 | 0.45 | 0.78 |
|  | ADB | 10 | 1.00 | 1.00 | 1.00 | 1.00 | 1.00 | 1.00 |
|  | Global | 54 | **0.87** | **0.57** | **0.66** | **0.83** | **0.59** | **0.78** |

Note: MK category: Morphokinetic category. EHB: Early hatching blastocysts; HB: Hatching blastocysts; SSE + SSB: Subtle developmental Shift Embryo + Subtle developmental Shift Blastocyst; ADE + ADB: Arrhythmic Development Embryo + Arrhythmic Development Blastocyst; AE: Arrested embryo and DE: Degenerated embryo. The performance of the learning process was calculated over 10 runs.

Finally, despite a decline in performance, the test under field conditions confirmed the ability of BEAM to predict whether an embryo belongs to one of the four morphokinetic categories of blastocysts (EHB, HB, SSB or ADB) with good overall results (F-score: 0.59 and Accuracy: 0.78) (Table 3). The high specificity indicates that incorrect classifications are rare, while the moderate sensitivity indicates that some true positives may be under-detected, especially in the EHB category.

### The automated morphokinetic classification system highlighted gene expression differences between the four morphokinetic categories immediately post-EGA

To investigate whether these morphokinetic categories reflect intrinsic characteristics of embryos around the EGA, the transcriptomic profile of 16 BEAM classified samples (4 samples per category (EHB, HB, SSB and ADB), each containing 8 embryos at the 16–32 cell stage), was analysed. The data have been deposited in the European Nucleotide Archive (ENA), https://www.ebi.ac.uk/ena/browser/home, under accession number PRJEB111128. The correlation clustering and PCA analysis showed clearly that the ADB category differed from the EHB and HB categories (Figure 3). The SSB category showed a more variable profile that could be close to both the ADB category (SSB-3 and SSB-4 samples) and to the EHB and HB categories (SSB-1 and SSB-2 samples).

**Figure 3:**
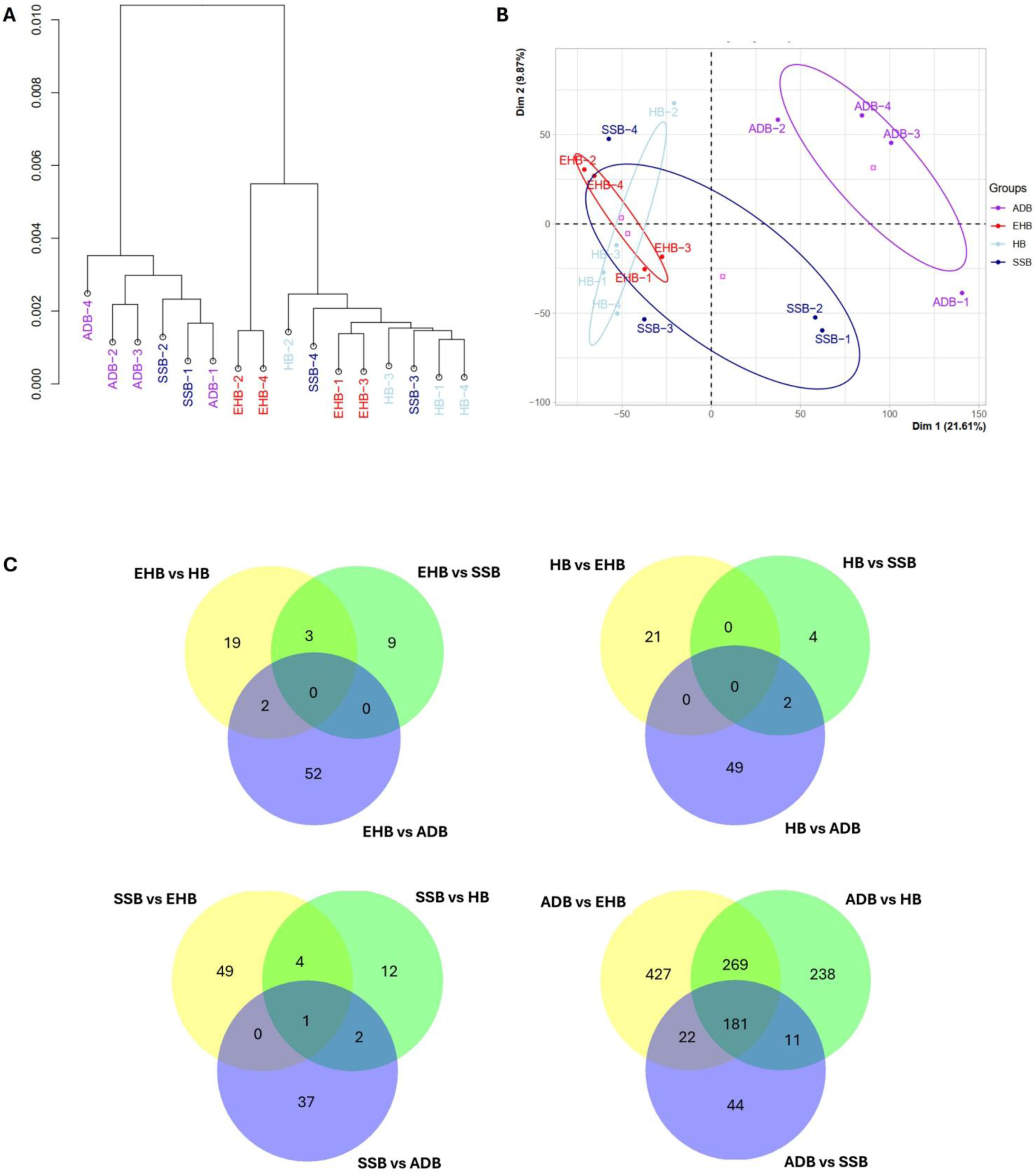
(EHB-1, EHB-2, EHB-3, EHB-4, HB-1, HB-2, HB-3, HB-4, SSB-1, SSB-2, SSB-3, SSB-4, ADB-1, ADB-2, ADCorrelation clustering (A) and Principal Component Analysis (B) showing the similarity and spatial distribution in multivariate space of the samples B-3, ADB-4), and a Venn diagram (C) representing the number of upregulated genes in each morphokinetic category (EHB, HB, SSB, ADB) compared to the others. EHB: Early hatching blastocysts, HB: Hatching blastocysts, SSB: Subtle developmental Shift Blastocyst and ADB: Arrhythmic Development Blastocyst.

The ADB embryos were characterised molecularly by a significant upregulation of 899, 699, and 258 genes compared with the EHB, HB, and SSB categories, respectively (Figure 3). Among these genes, 181 were significantly upregulated in ADB compared to all the other categories (Supplementary material 2, 3, 4, 5, 6, 7, 8). However, no genes upregulated in the EHB or HB categories were shared across all comparisons. The ADB and EHB categories showed the greatest transcriptomic difference, while HB and SSB showed the greatest similarity (only 25 differentially expressed genes, of which 19 were downregulated and 6 were upregulated in HB compared to SSB).

### The ADB category overexpressed numerous transcripts associated with transcriptional pathways

Only the comparison showing the greatest difference in gene expression (EHB vs ADB) was retained for subsequent analyses. A total of 54 genes were significantly downregulated, and 899 genes were upregulated in the ADB category compared to the EHB category.

Among the 899 upregulated and 54 downregulated transcripts in ADB compared to EHB, 816 and 40, respectively were present in the reference dataset [32] and a profile could not be established for 23 transcripts due to insufficient data (Figure 4). A high proportion of maternal and mixed profile genes were upregulated while the embryonic and mixed profiles represented the majority of the downregulated transcripts in ADB.

**Figure 4:**
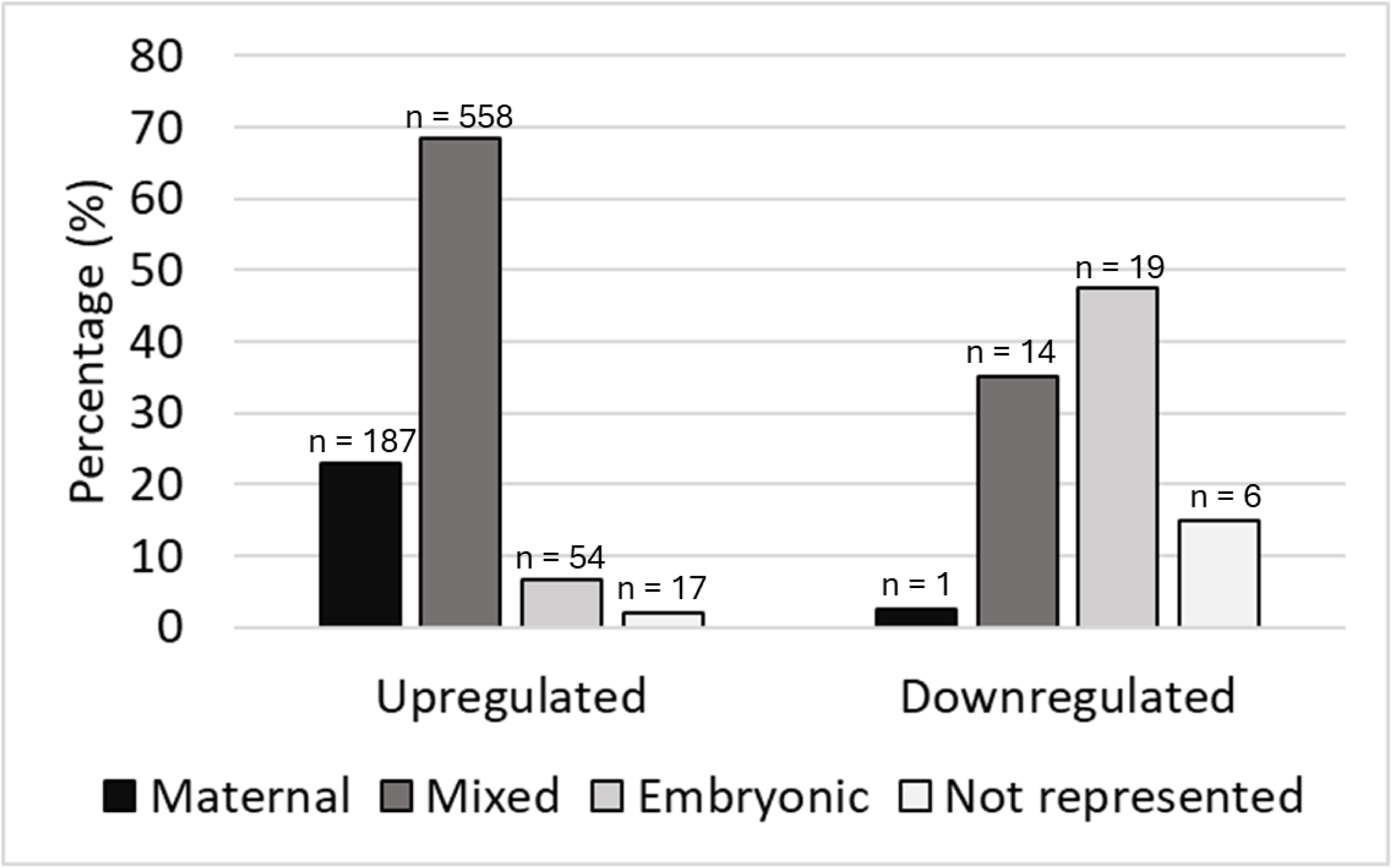
Percentage of transcripts of maternal, embryonic, mixed profiles upregulated or downregulated in category ADB compared to EHB. The dataset GSE52415 [32] was the reference to perform the transcript profile determination. A transcript with a median of zero across all stages in the reference dataset was considered “not represented”.

No functional pathways were significantly enriched among the genes downregulated in the ADB category, suggesting that this gene set (n=54) was too small or functionally dispersed. In contrast, a significant enrichment of 39 functional pathways from the *Biological Processes* and *Molecular Functions* (Gene Ontology, GO) and *Mammalian Phenotypes* (Mouse Genome Informatics, MGI) libraries was observed among the upregulated genes in ADB (Table 4). The list of overlapping genes per pathway is supplied in Supplementary Material 9.

**Table 4:**
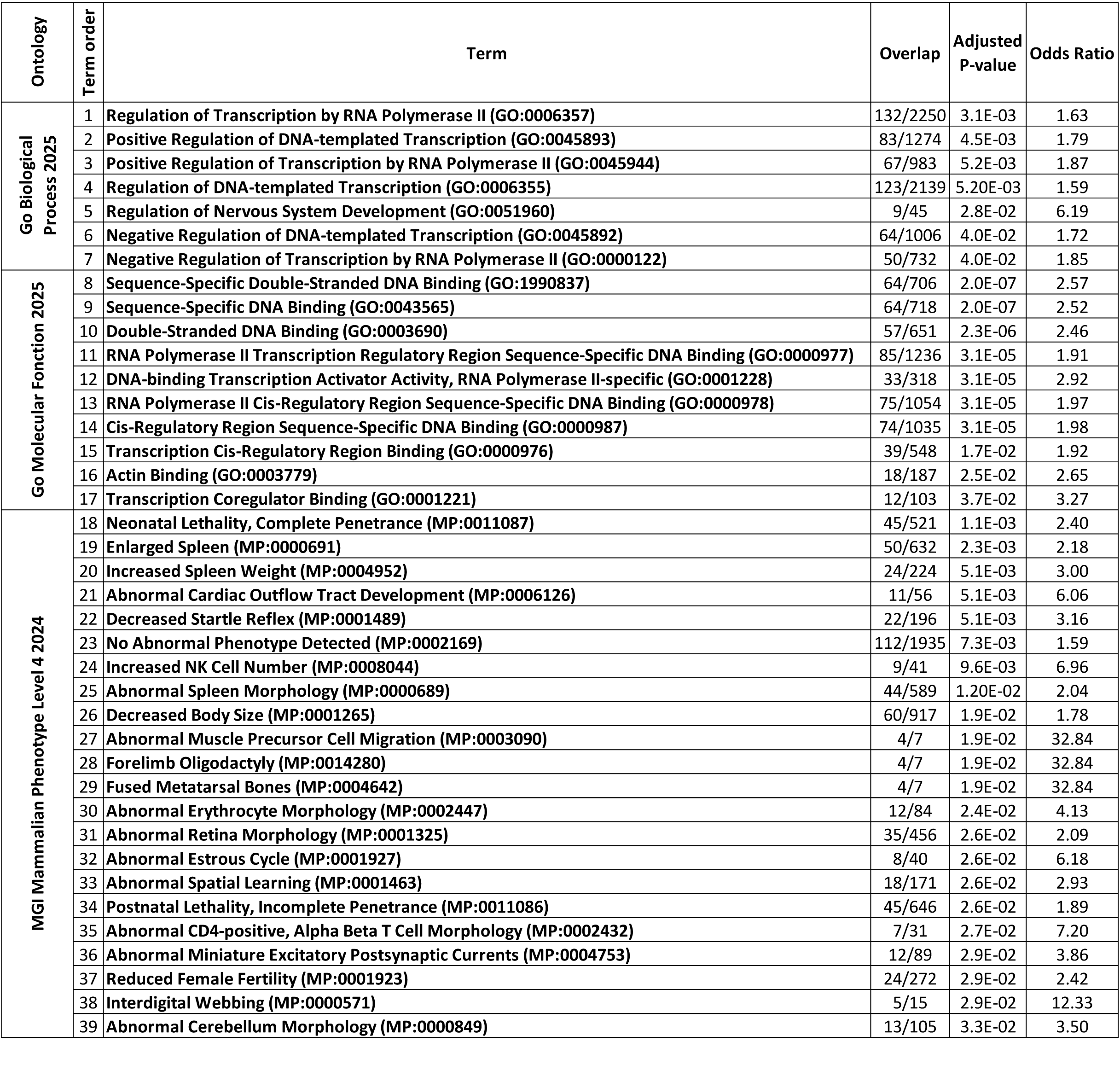
Functional pathway enrichment of genes overexpressed in the ADB category compared to the EHB category at the completion of the 4th embryonic cycle.

The enriched functional pathways in the Gene Ontology libraries are mainly related to transcription and its positive or negative regulation (n = 15 pathways / 17). Only two of these functional pathways are related to processes other than transcription: Regulation of Nervous System Development (Term order 5: related to the modulation of the formation, organisation and growth of the nervous system via transcriptional regulation, signalling and cell migration) and Actin binding (Term order 16: involved in cell motility and intracellular transport).

Twenty-one of the 22 enriched functional pathways of the MGI Mammalian phenotype library are linked to developmental abnormalities, suggesting a higher probability of developmental abnormalities in the ADB category compared to the EHB category. Among these functional pathways, established in the mouse, three seemed particularly relevant to cattle: Neonatal lethality, complete penetrance (term 18), Decreased body size (term 26) and Postnatal lethality, incomplete penetrance (term 34), because of high pregnancy loss and mortality in the perinatal period of IVP individuals.

## Discussion

The objectives of this study were twofold. Firstly, to develop a multiclass classification system capable of predicting the different morphokinetic categories of embryos (Early Hatching Blastocyst (EHB), Hatching Blastocyst (HB), Subtle Developmental Shift Embryo + Subtle Developmental Shift Blastocyst (SSE + SSB), Arrhythmic Development Embryo + Arrhythmic Development Blastocyst (ADE + ADB), Degenerated Embryos (DE) and Arrested Embryos (AE)). A particular focus was placed on the early identification of four blastocyst morphokinetic categories (EHB, HB, SSB and ADB prior to phenotype observation). Secondly, to evaluate whether morphokinetic trajectories leading to the EHB, HB, SSB and ADB categories were associated with molecular differences immediately following EGA.

Although expert classification of these morphokinetic categories is possible, it relies on observing the embryonic phenotype, in particular, blastocyst categories and hatching which can occur as late as 8_dpi_. In the field this is impractical because embryo transfer is done before hatching. The BEAM classifier (Bovine Embryo Analyser Based on Morphokinetics) circumvents this timing constraint by anticipating embryonic categories at an earlier stage (as early as the 4^th^ EC) based on a generalisable automatic learning process using the Random Forest method.

Other authors have used morphokinetic analysis to predict the developmental potential of an embryo. In general, these studies are based on a binary classification approach, such as blastocyst versus non-blastocyst [8, 9, 10], or pregnancy versus non-pregnancy [7]. Both deep learning [8] and more conventional statistics [7, 9, 10] have been employed. Deep learning has the advantage of being less dependent on human annotations, but it requires substantially larger datasets to achieve stable, satisfactory performance. In contrast, conventional statistical approaches rely on strong parametric assumptions and limited interaction structures.

We selected the Random Forest algorithm [17] due to its ability to accommodate complex, nonlinear relationships and high-order interactions among multiple variables, while remaining robust to noise, multicollinearity, and overfitting. Furthermore, it provides reliable estimates of feature importance, enabling the evaluation of each variable’s contribution across all interactions [17, 33, 34]. This methodological framework was therefore particularly well suited to our research question.

Binary classification is easier to perform than multiclass classification and requires fewer variables. Indeed, in [7, 9, 10], good accuracy was achieved with just two to five variables, whereas complex combinations of 27 variables were necessary for the BEAM multiclass prediction.

Six of these variables were common with at least one of the variables in the cited publications: the timing of the 1^st^ EC cytokinesis, cell asymmetry, synchronisation of the cleavages, cleavage type and synchronisation within a cycle, and the number of cells at the onset of the lag Phase. This suggests that these variables are relevant for embryo morphokinetic studies, regardless of the study’s objective.

However, it should be noted that each study used a different annotation system, which prevents direct comparisons. BEAM was trained using the standard annotation system developed in our laboratory [15] – translation supplied in Supplementary Data. All of the work presented in this article was performed using the same annotation system. The standardisation of the annotation is fundamental to improve inter-study comparisons and make further improvements in this area.

One final methodological point: BEAM is not intended to predict the IETS morphological classification [11, 12]. As previously demonstrated [10], we also found no relationship between morphokinetics and the IETS morphological classification.

The learning process was followed by a field test on an independent dataset, which confirmed BEAM’s ability to automatically predict whether an embryo belongs to one of the four morphokinetic categories of blastocysts before the phenotype is observed. However, the performance was lower than in the training phase. Several factors may have contributed to this difference in performance. These include the intrinsic variability of the classification system (with a standard deviation ranging from 0.05 to 0.09, depending on the category), the small size of the training dataset, and differences in the production of the second cohort of embryos: the change of donor breed used and the use of a new batch of culture media. Nevertheless, the F1 score and accuracy – performance metrics ranging from 0 (lowest performance) to 1 (highest performance) – showed good performance (0.59 and 0.78, respectively) for the model on the independent dataset and supported the continued use of this classification system for morphokinetic analysis and, in particular, for discriminating between blastocysts.

The second objective of our work was to determine whether morphokinetic trajectories leading to the EHB, HB, SSB and ADB categories predicted by BEAM were associated with molecular differences immediately following EGA, a pivotal event in embryonic development marking the transition to the embryo taking control of its developmental programme [35, 36].

Studies in Drosophila, zebrafish, mouse, human, sheep, goat, rabbit and cattle [37, 38, 39, 40, 41] have demonstrated that the maternal-to-embryonic transition (MET) is a two-way process. Maternal mRNA clearance is required to remove repressive factors and enable EGA. In turn, embryonic transcription is necessary for the degradation of subsets of maternal transcripts.

Studies [32, 41] in cattle have shown that the gradual degradation of maternal transcripts during early embryonic development (1^st^ to 3^rd^ EC) allows the full embryonic transcriptional programme to be established and the major wave of embryonic genome activation to be completed. It is characterised by a marked increase in transcriptional activity at the 8-16 cell stage (or 4^th^ EC) [32, 41, 42, 43]. Therefore, embryos that are unable to activate their genome at this stage will stop developing [44].

In this context, the exclusive selection of embryos that had completed the 4^th^ EC (16-cell stage completed), at 4.3_dpi_ in our study, made it possible to compare competent embryos at a strictly equivalent stage of development, presumably immediately after the major wave of EGA.

Given the highly comparable developmental stage between the different embryonic categories in our study, we anticipated a broadly similar proportion of maternal, embryonic and mixed profile transcripts within the differentially expressed genes between categories but with differences in encoded molecular functions.

Our comparison showed a large number of genes differentially expressed between the two extreme morphokinetic categories EHB and ADB (Figure 3), while the two central categories HB and SSB showed a fairly similar transcriptomic signature.

The comparison of the EHB and ADB categories revealed an upregulation of numerous maternal transcripts (22.6%) in ADB embryos which is consistent with a delay in their degradation. However, the presence of upregulated embryonic transcripts demonstrated that ADB embryos started to transcribe and have started EGA despite being slightly different from EHB. These two processes could explain the upregulation of numerous mixed transcripts (n = 558) in this category given that mixed transcripts can have both maternal and embryonic origins.

Similarly, the high number of enriched functional pathways related to transcription in the ADB category (n = 15) suggests differences in the dynamics or regulation of EGA between ADB and EHB categories. Indeed, these functional pathways are related to the key process of EGA [32] and concern different levels of transcription regulation [45, 46].

In parallel, the enrichment of the Actin binding pathway in the ADB category suggests an involvement of mechanisms related to the cytoskeleton, cell adhesion, or cell-cell signalling [47]. The cell-cell signalling pathway was enriched at the transcriptionally active 8-cell stage (3^rd^ EC) in cattle [41, 43]. These results confirm that different early morphokinetic patterns can capture variations in EGA dynamics among competent embryos.

Finally, the analysis of the mammalian phenotype library (MGI), revealed a significant enrichment of 22 pathways in the ADB category, 21 of which are linked to developmental abnormalities. These enriched functional pathways are mainly related to neonatal or perinatal mortality and developmental abnormalities such as small size, abnormal morphology (spleen, cerebellum, retina, bones) or abnormal functions (cardiac outflow, cell migration, spatial learning, excitatory postsynaptic currents). However, these phenotypes cannot be verified in our study because they can only be observed in the foetus or newborns. Indeed, although the frequency of neonatal or perinatal mortality varies between studies (from 7.8% to 22.2%), numerous data indicate its higher prevalence in IVP compared to IVD calves. Various causes, such as variations in birth size, malformations, calving difficulties, and alterations in biochemical profiles were reported [48, 49, 50, 51]. Our results encourage further studies in this area because they suggest that a number of abnormalities observed in the IVP newborn calf could be determined as early as the EGA stage.

In conclusion, our study showed that an automated classification system based on morphokinetics is able to identify four categories of competent embryos as early as the end of the 4^th^ EC. These categories show relevant gene expression differences immediately after the EGA. Particularly, embryos showing an arrhythmic development (category ADB) present a different pattern of MET including both a delay in the degradation of maternal transcripts and modified EGA. These transcriptomic differences could have long term effects, potentially causing higher neonatal mortality and abnormalities. Although this association requires functional validation, it reinforces the biological relevance of the proposed morphokinetic classification system.

Overall, the ability to predictively assign embryos to blastocyst categories prior to phenotypic observation opens up new avenues for embryo research and provides a powerful tool to refine embryo selection strategies and improve in vitro production (IVP) procedures.

## Supporting information

Supplementary material 1- Morphokinetic notation guidelines

Supplementary material 2: DEG_Comparison_EHB_vs_HB

Supplementary material 3: DEG_Comparison_EHB_vs_SSB

Supplementary material 4: DEG_Comparison_EHB_vs_ADB

Supplementary material 5: DEG_Comparison_HB_vs_SSB

Supplementary material 6: DEG_Comparison_HB_vs_ADB

Supplementary material 7: DEG_Comparison_SSB_vs_ADB

Supplementary material 8: DEG_Comparison_ADB_vs_ALL

Supplementary material 9: Gene overlap list par pathway

## Acknowledgements

The authors thank Patrick Bouthemy and Yasmine Hachani for the careful review of the manuscript and for the constructive comments provided prior to submission.

They acknowledge the sequencing and bioinformatics expertise of the I2BC High-throughput sequencing facility (https://www.i2bc.paris-saclay.fr/sequencing/ng-sequencing/; Université Paris-Saclay, Gif-sur-Yvette, France), supported by France Génomique (funded by the French National Program “Investissement d’Avenir” ANR-10-INBS-09)

## Use of AI

The authors declared that Generative AI was used in the creation of this manuscript. The authors used ChatGPT and DeepL to assist in correcting sentence structure and rephrasing. prior to final review, including additional editions, by a native English speaker. The authors take full responsibility for the content of this publication.

## Authors’ contributions

Co-senior authors: V.D., A.T.

Project design and coordination: A.P.R., V.D., A.T.

Production of embryos and videos: S.R., B.M-L, E.C., A.P.R., L.L.

Creation of the standardised annotation guide: N.L, A.P.R.

Expert morphokinetic annotation: A.P.R.

Morphokinetic data processing: M.B., M.B.C, J.U., E-M. S.

Predictive model development: M.B., A.P.R., A.T.

RNAm extraction: C.A.

RNA-seq library construction and sequencing: Y.J

Transcriptomic analysis: C.A., L.J., A.T., A.P.R., V.D.

Writing: A.P.R., M.B., N.L, A.T., A.P., V.D., A.T.

Proofreading: A.P.R., V.D., A.T., A.P.

## Declaration of Conflict of Interest

The authors declare that they have no financial or non-financial competing interests that could have influenced the work reported in this manuscript.

## Data availability

Data deposition: The data reported in this paper have been deposited in the European Nucleotide Archive (ENA), https://www.ebi.ac.uk/ena/browser/home, under accession number PRJEB111128.

## Notes

https://www.ebi.ac.uk/ena/browser/home

### Competing Interest Statement

The authors have declared no competing interest.

