## Supplementary material 1- Morphokinetic notation guidelines for "Bovine in vitro blastocysts with distinct morphokinetic patterns show transcriptomic differences at genome activation": Sup_Mat_1_Morphokinetics_notation_guidelines_vf.pdf

### Guidelines to Standardise the Notation of Morphokinetic parameters

#### Preamble:

This supplementary data is an English translation of part of Dr LeBrusq's work [15], developed to standardise the notation of morphokinetic parameters in our laboratory. Dr LeBrusq is a co-author of this article and has given her consent for the publication of this section of the notation guide.

Important note: The aim of this guide is not to explain the physiology of developmental events. Its aim is to describe the morphokinetic characteristics and the precise method for annotating the morphokinetic parameters observed in the videos in order to obtain standardised data.

#### Introduction:

Embryonic development is the result of a succession of embryonic cycles. From a morphokinetic perspective, an embryonic cycle (EC) refers to the sequence of events that leads all the cells of the embryo to divide once. It includes a phase of absence of division (corresponding to interphase), followed by a phase of cell division in which all the cells present at this stage commence division in a more or less synchronised manner (it corresponds to the cytokinesis) and this depends on the embryo. Morphokinetic parameters can be observed as the embryo progresses through the ECs. Certain parameters may be detected across several ECs and may have varying levels of importance depending on whether they arise during the initial embryonic cycles or during subsequent stages of development. Consequently, the notation of a given observed parameter must always be linked to the EC in which it is observed.

Final consideration: developmental events have both a morphological component (what is seen) and a kinetic component (when and for how long), since the progression from the stage before the event to the stage after the event is dynamic and can last from a few minutes to several hours. Both the morphological and kinetic components of these events can vary and do so relatively independently of one another. Our notation system therefore proposes to systematically separate morphological information (morphological variables) from kinetic information (kinetic variables), to improve the accuracy of the information obtained.

#### Morphological variables

##### Second polar body extrusion observed (Yes/No)

At the start of embryo culture (around 22h post-IVF), most embryos have already expelled their second polar body. However, if expulsion occurs later, it can be observed by the biologist since video recording started at 22h post-IVF. The expulsion of the second polar body follows a pattern like cytokinesis, involving membrane movements as seen during cytokinesis (for details, see Introduction of the variables describing the cytokinesis. This movement is followed by the expulsion of a small cell, (containing the chromosomes) surrounded by its cytoplasmic membrane. As the video progresses, it becomes apparent that this structure does not undergo subsequent division.

- Observation window:
    - between the start of the video (around 22h post-IVF) and the beginning of the first EC cytokinesis.
- Notation:
    - 0 – if the event of expulsion is not observed
    - 1 – if the event of expulsion is observed

If the polar body is already present at the beginning of the video, the notation is 0.

##### Presence of vacuole-like structures

A vacuole-like structure is any rounded cytoplasmic structure with a content that is distinctly more transparent than the embryo’s cytoplasm (whitish appearance) and has a defined outline, suggesting a vacuolar structure. Without additional staining, it is not possible to determine the true nature of these structures (i.e. a genuine vacuole or a lipid droplet), hence the term 'vacuole-like structure'. By playing the video forward, one can verify that the vacuole-like structure(s) do(es) not divide and is(are) stable in size. They can move in the cytoplasm and may be expelled along with cellular fragments following cell cytokinesis and disappear from the cytoplasm as development progresses. They are most frequently observed at the initial phases of development (from the zygote through to the second EC) or from the morula stage onwards.

- Observation window:
  - from the start of the video (around 22h post-IVF) until the onset of cytokinesis in the $1^{st}$ EC
- Notation:
  - 0 – if no vacuole-like structures are observed
  - 1 – if one or more vacuole-like structures are observed

If morula stage is studied, the same notation system can be used.

**Initiation of cytokinesis of an embryonic cycle (Yes/No)**

It is important to indicate if the initiation of cytokinesis in an embryonic cycle is observed (Yes or No). This is important because it informs us when there is no data whether this corresponds to an absence of notation or developmental arrest. In some cases, cytokinesis of two different ECs can be observed simultaneously. Consequently, noting this parameter requires individual monitoring of each cell in order to assign with certainty the EC corresponding to a cytokinesis event.

- Time of observation:
  - throughout the entire video, up to the start of cytokinesis in the $4^{th}$ EC.
- Notation:
  - For each EC ($1^{st}$, $2^{nd}$, $3^{rd}$ and $4^{th}$ ECs separately)
  - No – if none of the cells have initiated cytokinesis
  - Yes – if at least one of the cells begins cytokinesis

**Cytokinesis pattern (normal (1–2 cells); abnormal: (1–3; 1–4; reverse cytokinesis))**

Cytokinesis of a cell is considered normal when it produces two daughter cells. Any cytokinesis resulting in a different number of daughter cells or a reverse cytokinesis (fusion of the membranes of at least two daughter cells) is considered abnormal. Abnormal cytokinesis is most frequently observed during the 1$^{st}$ EC. Its incidence in subsequent embryonic cycles and at Lag phase is more sporadic. In these subsequent cycles, it can be identified by counting cells at the end of the cycle to prevent the notation process from becoming excessively burdensome.

Several types of cell cytokinesis are illustrated in **Figure 1S.**

- Observation window:
    - during the cytokinesis of the 1$^{st}$ EC.
- Notation:
    - Direct 1-3 (Yes/No);
    - Direct 1-4 (Yes/No);
    - Reverse cytokinesis (Yes/No).
    - Cytokinesis is normal if 'No' is noted for each of the three previous options

**Total cell number at the end of the EC**

At the end of an EC, an embryo theoretically has doubled its cell number. However, this number may differ due to abnormal cytokinesis, cellular degeneration or cell arrest.

- Observation window:
    - at the end of cytokinesis in a cycle (1$^{st}$, 2$^{nd}$ and 3$^{rd}$ EC).
- Notation:
    - The total number of cells is recorded for each EC separately
    - The total cell number includes living cells and degenerated cells during the EC.
    - The degenerated cells will be indicated in a dedicated column. Consequently, the combination of the number of cells and the number of degenerated cells provides

information on the total number of cells at the end of an EC expected to follow development.

**Number of degenerated cells during an EC**

Cell degeneration is sometimes observed during early embryo development, with varying severity depending on the stage at which it occurs. During the 1st, 2nd, 3rd EC and Lag Phase the onset of cell degeneration is marked by a halt in the movement of particles within the cytoplasm, associated with a gradual swelling of the cell until the membrane ruptures. The entire process can last from a few minutes to several hours. Thus, a cell may begin a process of degeneration during cycle n, whilst morphological confirmation of this can be observable during cycles n+1.

- Observation window:
    - throughout the 1st, 2nd, 3rd EC and Lag Phase.
- Notation:
    - Only cells which degenerated (via cytoplasmic membrane rupture) during a completed EC are noted for the cycle in question.

**Plasma membrane dynamics during cell cytokinesis**

During cytokinesis, the membrane is not static; it is actively remodelled: it is pulled inwards to form a cytokinesis furrow (driven by a contractile actomyosin ring beneath the membrane) and to form daughter cells.

In most cases, membrane movement is coordinated and is observed both immediately prior to and shortly after cytokinesis.

- Observation window:
    - during the cytokinesis of the 1st and 2nd ECs.
- Notation:
    - Membrane behaviour can be classified into four categories:

- Coordinated membrane movements: the cell outlines are smooth and rounded, and the cells separate harmoniously;
- Uncoordinated membrane movements: the cell outlines are smooth, but cell separation occurs more vigorously, as reflected by an increase in the number and/or speed of membrane movements;
- Membrane waves: The embryo's outline is irregular: all or part of the cell membrane exhibits wave-like movements;
- Absence of observable membrane movement: cytokinesis occurs within a time lapse shorter than the interval between two consecutive video frames (15 min). This may indicate either the absence of membrane movement during cytokinesis, or very rapid progression.

**Cellular asymmetry**

During cytokinesis, the resulting cells may not be of an identical size. To estimate asymmetry accurately, it is necessary to measure the volume of the daughter cells. However, 2D imaging and the dark appearance of the embryo prevent a reliable estimation of volume. Surface area is also difficult to measure, as this requires manually tracing the outline of each cell. We opted to calculate the difference between the products of the axes of each cell present. When more than two cells are present, only the largest and the smallest are considered. The degree of asymmetry between two daughter cells can be calculated.

- Observation window:
    - immediately after completion of cytokinesis of the 1st EC.
- Notation:
    - Follow the procedure below:
        1) Go to the first post-cytokinesis image where the two daughter cells are independent (enclosed by their own membranes).

2) Measure the two perpendicular axes of each daughter cell: the longest axis ($A_1$) and the widest axis ($A_2$). The length of each axis corresponds to the distance between two opposite membranes. The ends of the ruler must be placed on the internal face of the membrane (**Figure 2S**).
3) Multiply the two axes for each cell.
4) Calculate the asymmetry between the daughter cells using the formula:

$$Cellular\ asymmetry = 1 - \frac{(A1xA2)min}{(A1xA2)max}$$

**Cytokinesis axis orientation (visual assessment)**

This parameter is dynamic and its characterisation is limited by a 2D observation. Nevertheless, the position of the cytokinesis axis of the 1st EC allows three classes of cytokinesis axis orientations to be defined during the 2nd EC. The three main configurations are: (i) parallel orientation compared to the axis of the first cytokinesis, (ii) perpendicular orientation compared to this axis, and (iii) intermediate orientations between these two extremes. Embryos with a cell count different from four at the time of analysis are not eligible for notation of this parameter.

- Observation window:
    - the position of the cytokinesis axis at the end of the 1st EC is the reference. A second observation is performed during the cytokinesis of the 2nd EC. The formation of the furrow must be observed.
- Notation:
    - Four categories (Figure 3S):
        - Cytokinesis with perpendicular axes: the furrows of the 2nd EC are predominantly perpendicular to the cytokinesis axis of the 1st EC.
        - Cytokinesis with predominantly parallel axes: the furrows of the 2nd EC are predominantly parallel to the cytokinesis axis of the 1st EC.

- Mixed or other division axes: the furrows of the second EC show a combination of orientations (one axis can be parallel while the other can be perpendicular to the cytokinesis axis of the 1$^{st}$ EC
- Ineligible cases: annotated with '?' for embryos with a cell count which is different from four cells at the time of observation or when the embryologist is unable to define the orientation.

**Presence of fragmentation (Yes/No)**

Cell fragmentation is observable around the cytokinesis process. This generally occurs during or immediately after cytokinesis where some cells expel part of their cellular mass. It is often observed around furrow. These fragments vary in number and size. Furthermore, whilst it is easy to measure their size in some embryos, in others it can be more difficult because the fragments are partly hidden. We therefore recommend only noting the presence or absence of fragmentation to obtain a reliable parameter.

- Observation window:
    - during cytokinesis of the 1$^{st}$ EC.
- Notation:
    1) Position the video just before cytokinesis begins and play forward frame by frame until one hour after the end of cytokinesis.
    2) Observe the expulsion and/or the presence of new fragments (not previously present) in the perivitelline space.
    3) Notation:
        a. 0 if no new fragments are observed
        b. 1 if at least one new fragment is observed

**Reaching the hatching stage (Yes/No)**

Hatching is a characteristic seen at the expanded blastocyst stage. An embryo is hatching when the zona pellucida ruptures, or when partial or complete emergence of the embryo is observed through the opening.

- Observation window:
  - hatching is assessed by viewing the video from the onset of the blastocyst stage
- Notation:
  - Yes: if a hatching groove or the partial or complete emergence of the embryo from the zona pellucida is observed
  - No: if neither a hatching groove nor the partial or complete emergence of the embryo from the zona pellucida is observed before the end of the movie

**Signs of life (particle movements and membrane integrity)**

The signs of life in a cell are the presence of movement of intracellular particles and the presence of an intact membrane. When all the cells in an embryo have undergone degeneration, no particle movement is observed and the cell outlines are irregular and/or lack the smooth, well-defined edge resulting from the rupture of the cytoplasmic membrane. This variable allows us to verify whether the embryo is still alive or not.

- Observation window:
  - throughout the video, particularly in embryos no longer showing cytokinesis activity
- Notation:
  - when the embryo shows no signs of life, the stage of development at the time of detection is noted and the embryo is classified as degenerate (DE).
  - If there are signs of life without cell division, it is considered arrested (AE).

### Kinetic variables

The kinetic variables provide information about the timing and duration of events. The most appropriate temporal resolution for notation is in minutes. We therefore recommend noting all the kinetic variables in minutes.

**Timing of second polar body expulsion**

As explained previously (Variable “Second polar body extrusion observed”), at the time of embryo culture (>22h post-IVF), most embryos have already expelled their second polar body. When this expulsion occurs later, it can be observed in the video. It is possible to note the time of expulsion.

- Observation window:
  - between the start of the video and the start of cytokinesis of the 1st EC.
- Notation:
  - the time when polar body expulsion is completed.

**Variables related to the cytokinesis**

From a morphokinetic point of view, cytokinesis is characterised, in the first instance, by the formation of the cytokinesis furrow, which gradually deepens until the newly formed daughter cells are distinct and surrounded by their own membranes. This process may take varying lengths of time and may occur with varying degrees of synchronisation within an EC. In the moments preceding, during and immediately following cytokinesis, some specific movement of cytoplasmic particles as well as slight membrane movement, which does not result in the formation of a cytokinesis furrow, may be observed. These movements are not noted due to their limited objective visibility. Thus, only changes in the membrane resulting in the formation of the cytokinesis furrow through to the separation of the newly formed daughter cells are notated.

**Duration of cytokinesis of each cell**

- Observation window:
  - 1st, 2nd and 3rd C: from the start up to the end of cytokinesis for each cell in an embryonic cycle.
- Notation:
  - the observer must queue the video just before the start of cell division. The film must then be advanced frame by frame until the end of cell division. The cells of an embryo often overlap. Therefore, to determine the end of cytokinesis, it is necessary to observe the membrane outlines of the newly formed daughter cells through transparency.
  - The following information must be noted:
    - $Ti_{cn}$ = Start of cytokinesis of cell n, i.e. time T in the image corresponding to the onset of the cytokinesis furrow in the dividing cell.
    - $Tf_{cn}$ = End of cytokinesis of the cell n, i.e. time T in the image corresponding to the first appearance of newly formed distinct daughter cells, surrounded by their own membranes.
    - The duration of cytokinesis of the cell n is calculated as follows:

$$Duration\ of\ cytokinesis\ of\ the\ cell\ n = Tf_{cn} - Ti_{cn}$$

**Initiation ($t_{ie}$) and completion ($t_{fe}$) of cytokinesis of an embryonic cycle**

This parameter is intended to identify the start and end of the cytokinesis within an embryonic cycle. Cytokinesis occurring between these two points are not considered. This is particularly useful as development progresses and the number of cells increases, making detailed cell-by-cell notation more difficult.

- Observation window:

- from the start of cytokinesis of the first cell to enter division during a given EC to the end of cytokinesis of the last cell of that same EC to complete its cytokinesis. This parameter is annotated for the 1st, 2nd and 3rd EC. The expert uses this parameter at the 4th EC to distinguish between ADE/ADB and SSE/SSB.

- Notation:
  - The initiation of cytokinesis in the embryonic cycle ($t_{ie}$) corresponds to the onset of cytokinesis with the first cell to divide ($Ti_{c1}$), whereas its completion ($t_{fe}$) corresponds to the end of cytokinesis in the last dividing cell (l) within the embryonic cycle ($Tf_{cl}$) (see "Duration of cytokinesis of each cell").

**Duration of the cytokinesis of the embryonic cycle**

Once the variable "Initiation ($t_{ie}$) and completion ($t_{fe}$) of cytokinesis of the embryonic cycle" have been noted, the duration of cytokinesis of the embryonic cycle is calculated:

$$Duration\ of\ cytokinesis\ of\ the\ embryonic\ cycle\ n = Tf_{cl} - Ti_{c1}$$

This parameter is calculated for the 1st, 2nd, and 3rd EC.

**Synchronisation of cytokinesis within an embryonic cycle**

The synchronisation of cell cytokinesis within a EC corresponds to the interval between the initiation of cytokinesis in the first ($Ti_{c1}$) , and the last cells ($Ti_{cl}$) cleaved within an embryonic cycle.

- Observation window:
  - This parameter is noted for the 2nd and 3rd ECs. The observation period is defined as the interval between the onset of cytokinesis in the first cell to enter division during a given EC and the onset of cytokinesis in the last cell of that EC to complete division.
- Notation:

- the initiation of the cytokinesis of the first cell ($Ti_{c1}$) and the initiation of the cytokinesis of the last cell ($Ti_{cl}$) of an embryonic cycle are initially noted (see Variable "Duration of cytokinesis of each cell"). The duration is then calculated:

$$Synchronisation\ of\ cytokinesis\ within\ an\ embryonic\ cycle\ n = Ti_{cl} - Ti_{c1}$$

**Initiation ($Ti_{EC}$) and completion ($Tf_{EC}$) of an embryonic cycle**

The initiation and completion of an embryonic cycle allow its precocity to be assessed based on both the moment it starts and its completion.

Specificities:

- **The case of the first EC ($EC_1$):** the 1st EC is a special case. Its start should be counted from the moment of fertilisation. However, the exact time of fertilisation is unknown in In Vitro Fertilisation. By convention, the start of the 1st EC is the beginning of the co-incubation of the oocyte with the spermatozoa ($T_0 = 0$). The end of cytokinesis of the 1st cell coincides with the end of the 1st EC.
- **The case of the other ECs ($EC_2$, $EC_3$, $EC_4$):**
- Notation:
  - the initial time T of embryonic cycle n is counted from the first minute following the end of the last cytokinetic division of the preceding embryonic cycle. $Ti_{ECn} = Tf_{cl(ECn-1)} + 1$ (see Variable "Initiation ($t_{ie}$) and completion ($t_{fe}$) of cytokinesis of the embryonic cycle");
  - the final time T of embryonic cycle n, denoted $Tf_{ECn}$ , corresponds to the end of cytokinesis of the embryonic cycle ($Tf_{cl(ECn)}$)

**Duration of an embryonic cycle**

The duration of an embryonic cycle corresponds to the time required to complete interphase and cytokinesis for all cells present which are capable of division. Morphokinetics enables the determination of the duration of each embryonic cycle with an accuracy of ± X minutes (where X = the interval between two successive scheduled image captures – in our case: 15 minutes).

**The case of the first EC ($EC_1$):** As seen in Variable "**Initiation ($Ti_{EC}$) and completion ($Tf_{EC}$) of an embryonic cycle"** the start of EC1 is the moment the gametes meet ($T_0$= 0). Thus, the duration of the first EC is calculated as follows:

- $Duration\ of\ the\ 1st\ EC = Tf_{cl(EC1)} - 0$

**The case of the other ECs ($EC_2$, $EC_3$, $EC_4$):** for the other ECs, the duration includes the interval between the end of the previous embryonic cycle ($EC_{n-1}$) and the end of the $EC_n$, and is calculated as follows:

$$Duration\ of\ the\ \mathrm{EC}_n = Tf_{cl(ECn)} - (Tf_{cl(ECn-1)} + 1)$$

**Figure 4S** illustrates the set of kinetic parameters involving cytokinesis and the embryonic cycle.

**Initiation ($T_{il}$) and completion ($T_{fl}$) of the Lag phase.**

The Lag Phase is a period during which no cell division is visible for a duration exceeding the average length of an embryonic cycle, here, eleven hours.

- Observation window:
    - in normally developing embryos: after the 3rd EC, cytokinesis of a few cells at the 4th EC can be seen before the Lag Phase;
    - in abnormally developing embryos (not blastocysts): it may be observed at any stage of development, most commonly during the 2nd EC or during the 3rd EC. In some abnormally developing embryos, it is not observed.
- Notation:
    - $Ti_l$ = the initial time T of the Lag Phase. According to our method, the start of the Lag phase corresponds to the time T representing three images (45 minutes) after the final time of the last observed cytokinesis. This ensures that cytoplasmic and membrane movements or cell rearrangement associated with the end of a cycle's cytokinesis are not included in the Lag Phase.

- $Tf_l$ = the final time T of the Lag Phase, corresponding to the last image before the onset of spatial reorganisation of the cells immediately prior to the start of the next cytokinesis or, where applicable, the last image before the start of the next cytokinesis.

**Figure 5S** illustrates some examples of Lag Phase configurations.

**Duration of the Lag Phase.**

The total duration of the Lag Phase is calculated as follows:

$$Duration\ of\ the\ Lag\ Phase = Tf_l - Ti_l$$

**Initiation ($t_{ih}$) of hatching**

This parameter is recorded by focusing on the blastocyst. The film is then advanced whilst observing the entire surface and outline of the embryo. The initiation of hatching is recorded as soon as an open slit is observed on the surface of the zona pellucida or the embryo emerges partially or completely from its zona pellucida. The first of these phenomena to be observed indicates the initiation of hatching.

- Observation window:
    - from the start of blastocyst expansion, defined in our methodology as an intra-zona pellucida diameter >135 µm and the narrowing of the zona pellucida of at least 2 µm
- Notation:
    - $Ti_h$ corresponds to the first instance at which one of the signs of hatching is observed (an open slit on the surface of the zona pellucida or the partial or complete emergence of the embryo from its zona pellucida)

Note: some blastocysts can expel cells. It is not considered as a hatching process.

**Figures Legend**

**Figure 1S:** Different types of cell division observed in the in vitro-produced bovine embryo.

**Figure 2S:** Cell asymmetry notation at the end of the 1$^{st}$ EC.

**Figure 3S:** Cytokinesis axis orientation at the 2$^{nd}$ EC.

**Figure 4S:** Kinetic parameters related to cytokinesis and embryonic cycle (EC) duration.

**Figure 5S:** Examples of Phase Lag configurations in the bovine embryo.

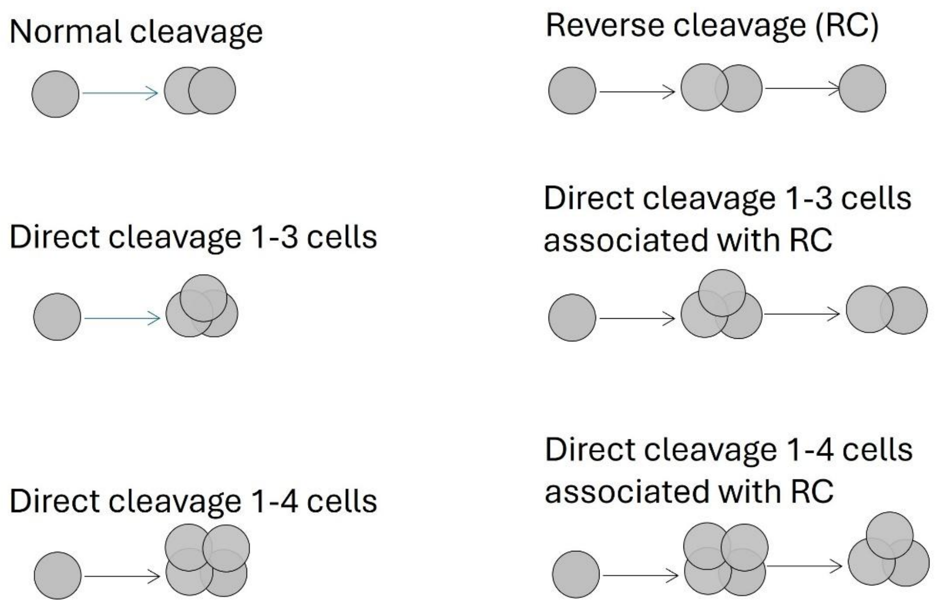


**Figure 4S:** Different types of cell division observed in the in vitro-produced bovine embryo.

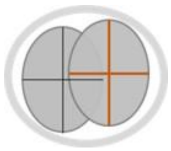

**Figure 5S:** Cell asymmetry notation at the end of the 1$^{st}$ EC.

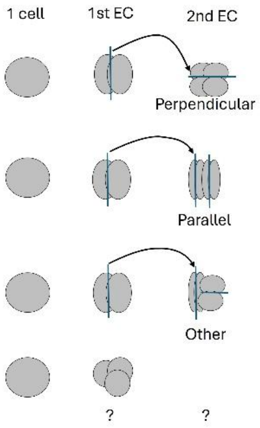


**Figure 6S:** Cytokinesis axis orientation at the 2$^{nd}$ EC.

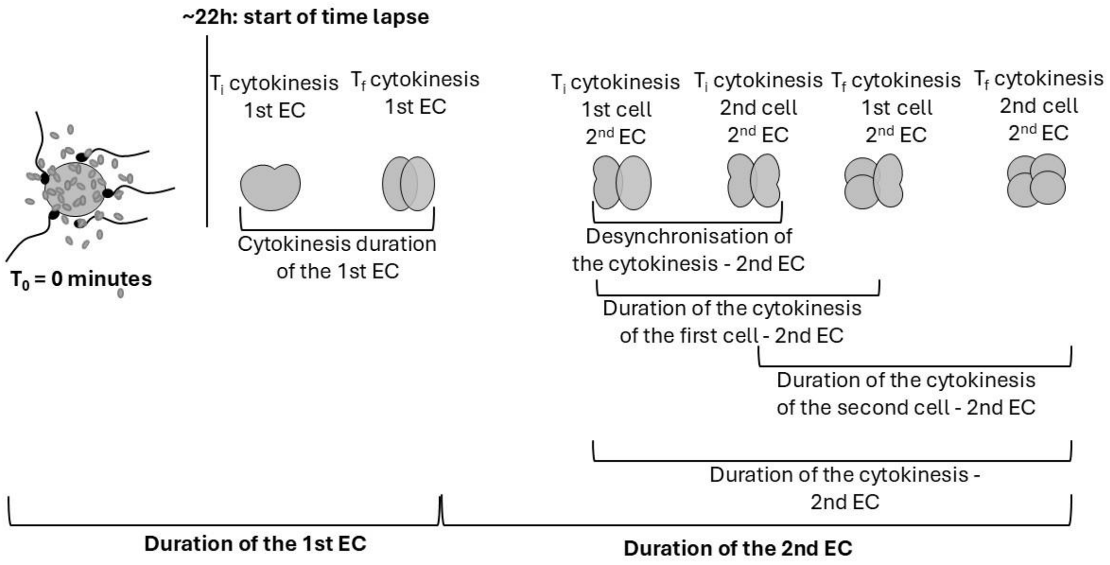


**Figure 4S:** Kinetic parameters related to cytokinesis and embryonic cycle (EC) duration.

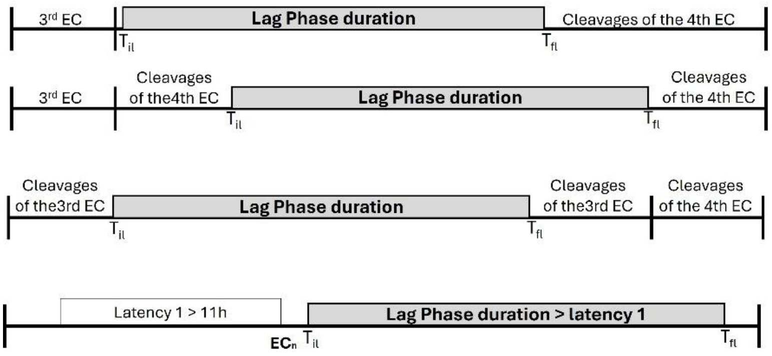


**Figure 5S:** Examples of Phase Lag configurations in the bovine embryo.
