## Supplementary material 2: DEG_Comparison_EHB_vs_HB for "Bovine in vitro blastocysts with distinct morphokinetic patterns show transcriptomic differences at genome activation"

| DEG_comparison_EHB_vs_HB | | | | | | | | | | | |
|---|---|---|---|---|---|---|---|---|---|---|---|
| **Geneid** | **Gene name** | **log2FC** | **padj** | **EHB-1** | **EHB-2** | **EHB-3** | **EHB-4** | **HB-1** | **HB-2** | **HB-3** | **HB-4** |
| **ENSBTAG00000049405** | | -19.70 | 4.90E-05 | 0.00 | 0.00 | 0.00 | 0.00 | 0.00 | 93.24 | 0.00 | 0.00 |
| **ENSBTAG00000021408** | FMO1 | -19.52 | 4.20E-08 | 0.00 | 0.00 | 0.00 | 0.00 | 0.00 | 77.83 | 0.00 | 0.00 |
| **ENSBTAG00000045714** | DSPP | -15.56 | 3.82E-02 | 0.00 | 0.00 | 0.00 | 0.00 | 4.97 | 0.00 | 0.00 | 0.00 |
| **ENSBTAG00000006350** | SLC22A7 | -7.51 | 6.85E-04 | 0.00 | 0.00 | 0.00 | 0.00 | 73.53 | 29.19 | 17.04 | 24.97 |
| **ENSBTAG00000016365** | ABCG5 | -7.03 | 1.80E-04 | 0.00 | 1.02 | 0.00 | 0.00 | 5.96 | 100.53 | 52.74 | 11.24 |
| **ENSBTAG00000001764** | NCAN | -7.02 | 6.95E-03 | 0.00 | 0.00 | 0.00 | 0.00 | 24.84 | 17.03 | 30.02 | 31.22 |
| **ENSBTAG00000013984** | KL | -6.99 | 1.82E-02 | 0.00 | 0.00 | 0.00 | 0.00 | 35.77 | 7.30 | 52.74 | 4.99 |
| **ENSBTAG00000027490** | OR5A1 | -6.71 | 1.00E-02 | 0.00 | 0.00 | 0.00 | 0.00 | 8.94 | 51.89 | 21.91 | 0.00 |
| **ENSBTAG00000007090** | MYH2 | -6.46 | 4.60E-03 | 0.00 | 1.02 | 0.00 | 0.00 | 8.94 | 57.56 | 13.79 | 34.96 |
| **ENSBTAG00000013319** | SMCO1 | -6.42 | 2.21E-02 | 0.00 | 0.00 | 0.00 | 0.00 | 19.87 | 1.62 | 24.34 | 22.48 |
| **ENSBTAG00000048611** | | -6.38 | 1.14E-02 | 0.00 | 0.00 | 1.23 | 0.00 | 12.92 | 0.00 | 31.64 | 64.93 |
| **ENSBTAG00000046298** | SYCE3 | -5.85 | 4.94E-02 | 0.00 | 0.00 | 0.00 | 0.00 | 0.00 | 9.73 | 11.36 | 24.97 |
| **ENSBTAG00000001243** | AP3M2 | -5.33 | 7.70E-05 | 3.02 | 1.02 | 0.00 | 0.00 | 50.68 | 64.86 | 31.64 | 22.48 |
| **ENSBTAG00000044012** | TMEM196 | -4.82 | 4.95E-03 | 3.02 | 0.00 | 3.69 | 0.00 | 0.00 | 129.72 | 28.40 | 27.47 |
| **ENSBTAG00000015525** | SLC39A12 | -3.72 | 4.83E-03 | 7.05 | 1.02 | 7.38 | 0.00 | 29.81 | 71.35 | 42.19 | 58.69 |
| **ENSBTAG00000040202** | | -2.65 | 4.62E-02 | 13.10 | 2.03 | 23.38 | 5.38 | 47.70 | 132.96 | 47.06 | 43.70 |
| **ENSBTAG00000014300** | SLC6A5 | -2.58 | 4.60E-03 | 18.14 | 9.14 | 20.92 | 10.76 | 83.47 | 163.77 | 52.74 | 49.95 |
| **ENSBTAG00000053488** | | -2.34 | 3.01E-02 | 19.14 | 6.09 | 4.92 | 16.15 | 81.48 | 22.70 | 44.63 | 88.66 |
| **ENSBTAG00000043996** | SAMD12 | -2.34 | 3.75E-02 | 33.25 | 20.31 | 41.84 | 7.53 | 24.84 | 293.49 | 89.25 | 112.38 |
| **ENSBTAG00000004974** | ZDHHC23 | -1.58 | 7.92E-03 | 53.40 | 42.64 | 43.07 | 88.26 | 237.48 | 130.53 | 139.56 | 172.32 |
| **ENSBTAG00000044044** | CLOCK | -1.03 | 4.66E-02 | 380.86 | 228.44 | 388.90 | 589.84 | 614.08 | 789.66 | 1053.17 | 792.92 |
| **ENSBTAG00000018527** | HDGFL3 | 1.35 | 4.62E-02 | 415.12 | 866.03 | 367.98 | 593.07 | 242.45 | 209.17 | 223.13 | 207.28 |
| **ENSBTAG00000010368** | TPST2 | 1.70 | 8.59E-03 | 308.32 | 698.51 | 311.37 | 821.26 | 152.03 | 208.36 | 184.18 | 114.88 |
| **ENSBTAG00000010487** | SMIM3 | 1.75 | 3.82E-02 | 169.27 | 503.57 | 376.60 | 273.39 | 74.52 | 144.31 | 100.61 | 73.67 |
| **ENSBTAG00000001116** | P4HA2 | 1.80 | 4.62E-02 | 144.08 | 791.91 | 172.30 | 491.89 | 157.00 | 141.88 | 99.80 | 61.19 |
| **ENSBTAG00000000899** | TMEM17 | 1.94 | 4.62E-02 | 117.89 | 250.77 | 119.38 | 192.67 | 15.90 | 43.78 | 46.25 | 72.42 |
| **ENSBTAG00000013456** | TMEM207 | 1.96 | 4.66E-02 | 73.55 | 152.29 | 115.69 | 148.54 | 25.83 | 43.78 | 28.40 | 27.47 |
| **ENSBTAG00000003752** | SLC25A24 | 2.00 | 1.24E-02 | 262.98 | 587.84 | 92.30 | 437.00 | 137.12 | 58.37 | 68.16 | 82.41 |
| **ENSBTAG00000051910** | FGF10 | 3.58 | 3.82E-02 | 58.44 | 63.96 | 19.69 | 27.99 | 0.00 | 8.11 | 0.00 | 6.24 |
| **ENSBTAG00000002526** | BDH2 | 3.65 | 1.42E-02 | 18.14 | 366.51 | 50.46 | 254.02 | 14.90 | 5.68 | 30.02 | 3.75 |
| **ENSBTAG00000011274** | ENO2 | 4.23 | 2.27E-02 | 35.26 | 15.23 | 23.38 | 186.21 | 3.97 | 0.81 | 6.49 | 2.50 |

| | | | | | | | | | | | |
|---|---|---|---|---|---|---|---|---|---|---|---|
| **ENSBTAG00000025401** | DOK3 | 5.43 | 2.34E-02 | 25.19 | 20.31 | 30.77 | 7.53 | 0.00 | 0.00 | 0.00 | 2.50 |
| **ENSBTAG00000006039** | ARHGDIB | 6.20 | 1.82E-02 | 24.18 | 15.23 | 0.00 | 11.84 | 0.00 | 0.00 | 0.00 | 0.00 |
| **ENSBTAG00000013339** | NEBL | 7.03 | 1.80E-04 | 13.10 | 25.38 | 9.85 | 43.05 | 0.00 | 0.00 | 0.00 | 0.00 |
| **ENSBTAG00000046155** | RGN | 7.08 | 8.59E-03 | 15.11 | 61.93 | 16.00 | 62.43 | 0.00 | 0.81 | 0.00 | 0.00 |
| **ENSBTAG00000033352** | DMRT1 | 7.62 | 1.81E-02 | 79.60 | 0.00 | 20.92 | 36.60 | 0.00 | 0.00 | 0.00 | 0.00 |
| **ENSBTAG00000003892** | CMAH | 7.68 | 3.96E-02 | 11.08 | 91.37 | 1.23 | 39.83 | 0.00 | 0.00 | 0.00 | 0.00 |
| **ENSBTAG00000016026** | PCOLCE2 | 9.72 | 3.38E-03 | 46.35 | 238.59 | 6.15 | 683.48 | 0.00 | 0.00 | 0.00 | 1.25 |
| **ENSBTAG00000053807** | | 20.57 | 1.55E-03 | 1.01 | 27.41 | 14.77 | 0.00 | 0.00 | 0.00 | 0.00 | 0.00 |
| **ENSBTAG00000007881** | IFIT1 | 21.16 | 8.67E-04 | 0.00 | 0.00 | 0.00 | 71.04 | 0.00 | 0.00 | 0.00 | 0.00 |
| **ENSBTAG00000006432** | KCNE4 | 21.30 | 7.84E-04 | 0.00 | 73.10 | 3.69 | 0.00 | 0.00 | 0.00 | 0.00 | 0.00 |
| **ENSBTAG00000019428** | CCR1 | 21.48 | 1.02E-05 | 0.00 | 0.00 | 0.00 | 91.49 | 0.00 | 0.00 | 0.00 | 0.00 |
| **ENSBTAG00000016444** | RETREG1 | 21.49 | 6.85E-04 | 0.00 | 0.00 | 0.00 | 85.03 | 0.00 | 0.00 | 0.00 | 0.00 |
| **ENSBTAG00000007596** | GEM | 24.73 | 3.76E-09 | 0.00 | 581.75 | 0.00 | 349.81 | 0.00 | 0.00 | 0.00 | 0.00 |
| **ENSBTAG00000004547** | OLR1 | 25.16 | 1.15E-05 | 0.00 | 966.54 | 0.00 | 445.61 | 0.00 | 0.00 | 0.00 | 0.00 |
