## Supplementary material 3: DEG_Comparison_EHB_vs_SSB for "Bovine in vitro blastocysts with distinct morphokinetic patterns show transcriptomic differences at genome activation"

| DEG_comparison_EHB_vs_SSB | | | | | | | | | | | |
|---|---|---|---|---|---|---|---|---|---|---|---|
| **Geneid** | **Gene name** | **log2FC** | **padj** | **EHB-1** | **EHB-2** | **EHB-3** | **EHB-4** | **SSB-1** | **SSB-2** | **SSB-3** | **SSB-4** |
| **ENSBTAG00000007927** | SLAMF1 | -20.45 | 1.46E-03 | 0.00 | 0.00 | 0.00 | 0.00 | 68.79 | 0.00 | 0.00 | 0.00 |
| **ENSBTAG00000000707** | WISP1 | -20.43 | 1.46E-03 | 0.00 | 0.00 | 0.00 | 0.00 | 0.00 | 134.58 | 0.00 | 0.00 |
| **ENSBTAG00000022759** | PAG11 | -19.62 | 1.67E-03 | 0.00 | 0.00 | 0.00 | 0.00 | 0.00 | 74.32 | 0.00 | 0.00 |
| **ENSBTAG00000045714** | DSPP | -16.03 | 1.71E-02 | 0.00 | 0.00 | 0.00 | 0.00 | 6.79 | 0.00 | 0.00 | 0.00 |
| **ENSBTAG00000017294** | ORM1 | -15.56 | 4.84E-02 | 0.00 | 0.00 | 0.00 | 0.00 | 4.25 | 0.00 | 0.00 | 0.00 |
| **ENSBTAG00000049405** | | -14.40 | 2.02E-02 | 0.00 | 0.00 | 0.00 | 0.00 | 0.00 | 0.00 | 2.41 | 0.00 |
| **ENSBTAG00000021408** | FMO1 | -13.26 | 7.57E-03 | 0.00 | 0.00 | 0.00 | 0.00 | 0.00 | 1.00 | 0.00 | 0.00 |
| **ENSBTAG00000027490** | OR5A1 | -7.94 | 4.39E-04 | 0.00 | 0.00 | 0.00 | 0.00 | 26.33 | 8.03 | 99.84 | 60.47 |
| **ENSBTAG00000016365** | ABCG5 | -7.20 | 1.05E-04 | 0.00 | 1.02 | 0.00 | 0.00 | 38.22 | 80.35 | 40.90 | 33.08 |
| **ENSBTAG00000049599** | FAM163A | -6.86 | 1.95E-02 | 0.00 | 0.00 | 0.00 | 0.00 | 49.26 | 37.16 | 0.00 | 5.70 |
| **ENSBTAG00000054666** | | -6.82 | 1.56E-02 | 0.00 | 0.00 | 0.00 | 0.00 | 0.00 | 32.14 | 18.04 | 39.93 |
| **ENSBTAG00000031573** | NMRK2 | -6.71 | 2.46E-02 | 0.00 | 0.00 | 0.00 | 0.00 | 8.49 | 46.20 | 0.00 | 28.52 |
| **ENSBTAG00000002199** | CORIN | -6.45 | 6.38E-03 | 0.00 | 0.00 | 0.00 | 0.00 | 19.53 | 22.10 | 9.62 | 18.25 |
| **ENSBTAG00000013319** | SMCO1 | -6.32 | 1.87E-02 | 0.00 | 0.00 | 0.00 | 0.00 | 10.19 | 9.04 | 15.64 | 28.52 |
| **ENSBTAG00000006350** | SLC22A7 | -6.29 | 1.19E-02 | 0.00 | 0.00 | 0.00 | 0.00 | 0.00 | 29.13 | 21.65 | 11.41 |
| **ENSBTAG00000001764** | NCAN | -6.18 | 2.46E-02 | 0.00 | 0.00 | 0.00 | 0.00 | 11.89 | 43.19 | 0.00 | 2.28 |
| **ENSBTAG00000048611** | | -5.91 | 2.03E-02 | 0.00 | 0.00 | 1.23 | 0.00 | 28.88 | 14.06 | 8.42 | 27.38 |
| **ENSBTAG00000007090** | MYH2 | -5.77 | 1.58E-02 | 0.00 | 1.02 | 0.00 | 0.00 | 11.89 | 20.09 | 34.88 | 4.56 |
| **ENSBTAG00000001067** | GRM2 | -5.59 | 3.61E-02 | 0.00 | 0.00 | 0.00 | 1.08 | 33.97 | 19.08 | 9.62 | 0.00 |
| **ENSBTAG00000001243** | AP3M2 | -5.24 | 1.17E-04 | 3.02 | 1.02 | 0.00 | 0.00 | 48.41 | 25.11 | 9.62 | 76.44 |
| **ENSBTAG00000044012** | TMEM196 | -5.06 | 1.67E-03 | 3.02 | 0.00 | 3.69 | 0.00 | 58.60 | 68.30 | 66.16 | 26.24 |
| **ENSBTAG00000015525** | SLC39A12 | -3.97 | 1.30E-03 | 7.05 | 1.02 | 7.38 | 0.00 | 74.74 | 134.58 | 9.62 | 19.39 |
| **ENSBTAG00000006937** | ABCA10 | -3.92 | 4.65E-02 | 3.02 | 5.08 | 2.46 | 1.08 | 5.10 | 29.13 | 18.04 | 125.49 |
| **ENSBTAG00000018650** | HEPACAM | -3.37 | 2.43E-02 | 13.10 | 8.12 | 3.69 | 0.00 | 57.75 | 16.07 | 143.14 | 42.21 |
| **ENSBTAG00000006064** | C7H19orf57 | -3.13 | 4.27E-02 | 10.08 | 3.05 | 4.92 | 0.00 | 48.41 | 41.18 | 38.49 | 30.80 |
| **ENSBTAG00000011733** | GIPC2 | -2.78 | 1.70E-05 | 51.39 | 15.23 | 32.00 | 19.37 | 225.92 | 215.94 | 187.65 | 181.40 |
| **ENSBTAG00000043996** | SAMD12 | -2.75 | 2.89E-03 | 33.25 | 20.31 | 41.84 | 7.53 | 163.92 | 227.99 | 81.79 | 215.62 |
| **ENSBTAG00000047658** | PHF7 | -2.46 | 1.01E-02 | 2.02 | 43.66 | 13.54 | 18.30 | 60.30 | 119.52 | 137.13 | 110.66 |
| **ENSBTAG00000014300** | SLC6A5 | -2.09 | 4.27E-02 | 18.14 | 9.14 | 20.92 | 10.76 | 123.15 | 57.25 | 16.84 | 51.34 |
| **ENSBTAG00000003103** | EIF4E1B | -2.05 | 1.31E-02 | 443.33 | 133.00 | 380.29 | 105.48 | 1204.33 | 1484.43 | 716.90 | 990.26 |
| **ENSBTAG00000038891** | | -1.96 | 1.19E-02 | 30.23 | 86.30 | 265.83 | 73.19 | 295.56 | 506.19 | 566.54 | 405.00 |

| | | | | | | | | | | | |
|---|---|---|---|---|---|---|---|---|---|---|---|
| **ENSBTAG00000009800** | MSI1 | -1.96 | 2.92E-02 | 22.17 | 14.21 | 45.54 | 23.68 | 110.41 | 89.39 | 64.95 | 142.61 |
| **ENSBTAG00000013219** | PEX5L | -1.94 | 1.91E-02 | 83.63 | 47.72 | 115.69 | 16.15 | 223.37 | 336.46 | 191.25 | 257.83 |
| **ENSBTAG00000000015** | FOXRED2 | -1.75 | 2.08E-04 | 125.95 | 300.52 | 126.76 | 172.22 | 540.16 | 904.92 | 433.03 | 559.02 |
| **ENSBTAG00000000623** | CPEB1 | -1.69 | 1.64E-02 | 2218.67 | 1317.82 | 3151.84 | 923.51 | 6756.30 | 6351.51 | 4809.02 | 6629.53 |
| **ENSBTAG00000026972** | MYF5 | -1.60 | 2.26E-02 | 324.44 | 256.86 | 358.14 | 99.02 | 536.77 | 1062.60 | 761.41 | 788.33 |
| **ENSBTAG00000005729** | FBXL4 | -1.60 | 1.19E-02 | 153.15 | 60.92 | 203.07 | 121.63 | 485.81 | 553.40 | 340.41 | 246.43 |
| **ENSBTAG00000039522** | CYCT | -1.51 | 4.72E-02 | 467.51 | 391.89 | 146.45 | 354.12 | 431.45 | 1355.87 | 682.02 | 1395.27 |
| **ENSBTAG00000002202** | CRAMP1 | -1.46 | 4.27E-02 | 147.11 | 93.40 | 136.61 | 55.97 | 276.88 | 585.54 | 170.81 | 157.44 |
| **ENSBTAG00000019791** | GRHL3 | -1.42 | 3.85E-02 | 75.57 | 67.01 | 25.84 | 52.74 | 191.95 | 79.34 | 150.36 | 173.41 |
| **ENSBTAG00000012682** | UNC13A | -1.38 | 2.89E-03 | 168.26 | 113.71 | 147.68 | 130.24 | 425.51 | 464.01 | 194.86 | 374.20 |
| **ENSBTAG00000004974** | ZDHHC23 | -1.36 | 3.36E-02 | 53.40 | 42.64 | 43.07 | 88.26 | 214.88 | 131.57 | 133.52 | 102.68 |
| **ENSBTAG00000007494** | SMARCA2 | -1.22 | 4.27E-02 | 324.44 | 110.66 | 422.13 | 246.48 | 510.44 | 839.64 | 597.82 | 616.06 |
| **ENSBTAG00000037710** | | -1.18 | 4.27E-02 | 1432.76 | 651.80 | 931.64 | 539.25 | 1997.59 | 2604.28 | 1477.11 | 2001.07 |
| **ENSBTAG00000018307** | SEMA3F | -1.15 | 4.27E-02 | 172.29 | 118.79 | 163.68 | 137.77 | 387.29 | 484.10 | 200.88 | 243.00 |
| **ENSBTAG00000048876** | FAM117B | -1.15 | 4.31E-02 | 297.23 | 241.63 | 439.36 | 207.74 | 889.23 | 877.80 | 469.11 | 385.61 |
| **ENSBTAG00000002370** | ZNF792 | -1.14 | 9.58E-04 | 138.04 | 234.53 | 273.22 | 250.79 | 558.85 | 503.18 | 401.75 | 514.53 |
| **ENSBTAG00000025441** | HSPA1A | -1.14 | 9.86E-03 | 2591.47 | 1773.68 | 2914.31 | 2651.06 | 5752.41 | 6104.44 | 6079.23 | 3909.72 |
| **ENSBTAG00000001294** | PPP1R15A | -1.10 | 4.27E-02 | 445.35 | 337.07 | 493.51 | 257.25 | 917.26 | 1158.02 | 525.65 | 674.25 |
| **ENSBTAG00000020878** | DMTF1 | -1.07 | 1.14E-03 | 202.52 | 172.60 | 180.91 | 179.75 | 338.03 | 290.26 | 560.53 | 359.37 |
| **ENSBTAG00000046701** | FRAT2 | -1.03 | 4.91E-02 | 570.28 | 237.57 | 433.21 | 463.91 | 849.31 | 1067.62 | 983.94 | 581.84 |
| **ENSBTAG00000003889** | PER1 | -1.03 | 2.46E-02 | 382.88 | 231.48 | 374.13 | 354.12 | 980.96 | 715.10 | 611.05 | 430.10 |
| **ENSBTAG00000011041** | ZFYVE1 | -1.01 | 1.58E-02 | 577.34 | 510.68 | 689.20 | 478.98 | 1403.07 | 1433.21 | 819.14 | 883.02 |
| **ENSBTAG00000014387** | PRKAB2 | -1.00 | 2.09E-02 | 1554.68 | 1292.44 | 1304.55 | 1122.64 | 3216.35 | 3855.70 | 1835.56 | 1651.96 |
| **ENSBTAG00000019453** | PTGES | 1.03 | 1.19E-02 | 1859.97 | 3071.19 | 2547.56 | 2399.19 | 1586.52 | 797.45 | 1046.48 | 1398.69 |
| **ENSBTAG00000003470** | TTYH1 | 1.22 | 4.76E-02 | 102.77 | 144.17 | 98.46 | 88.26 | 58.60 | 40.17 | 42.10 | 44.49 |
| **ENSBTAG00000011424** | TPM2 | 2.23 | 4.77E-02 | 712.35 | 4782.94 | 873.80 | 3384.06 | 619.15 | 641.78 | 532.86 | 281.79 |
| **ENSBTAG00000011990** | ALOX15 | 3.88 | 1.27E-02 | 23.17 | 35.53 | 19.69 | 55.97 | 2.55 | 3.01 | 1.20 | 2.28 |
| **ENSBTAG00000055051** | RNF125 | 5.16 | 3.67E-05 | 87.66 | 90.36 | 67.69 | 32.29 | 0.00 | 0.00 | 8.42 | 0.00 |
| **ENSBTAG00000052709** | | 6.53 | 3.52E-02 | 10.08 | 25.38 | 4.92 | 76.42 | 0.00 | 1.00 | 0.00 | 0.00 |
| **ENSBTAG00000005017** | ZNF169 | 7.28 | 4.27E-02 | 88.67 | 0.00 | 1.23 | 29.06 | 0.00 | 0.00 | 0.00 | 0.00 |
| **ENSBTAG00000003892** | CMAH | 7.55 | 2.95E-02 | 11.08 | 91.37 | 1.23 | 39.83 | 0.00 | 0.00 | 0.00 | 0.00 |
| **ENSBTAG00000007881** | IFIT1 | 21.01 | 9.58E-04 | 0.00 | 0.00 | 0.00 | 71.04 | 0.00 | 0.00 | 0.00 | 0.00 |
| **ENSBTAG00000007196** | TAGLN | 25.28 | 3.45E-06 | 0.00 | 877.19 | 0.00 | 656.58 | 0.00 | 0.00 | 0.00 | 0.00 |

| | | | | | | | | | | | |
|---|---|---|---|---|---|---|---|---|---|---|---|
| **ENSBTAG00000004547** | OLR1 | 25.30 | 1.24E-05 | 0.00 | 966.54 | 0.00 | 445.61 | 0.00 | 0.00 | 0.00 | 0.00 |
| **ENSBTAG00000018563** | SFRP2 | 26.42 | 4.05E-14 | 0.00 | 2347.30 | 0.00 | 1176.45 | 0.00 | 0.00 | 0.00 | 0.00 |
