## Supplementary material 4: DEG_Comparison_EHB_vs_ADB for "Bovine in vitro blastocysts with distinct morphokinetic patterns show transcriptomic differences at genome activation"

| DEG comparison EHB vs ADB | | | | | | | | | | | |
|---|---|---|---|---|---|---|---|---|---|---|---|
| **Geneid** | **Gene name** | **log2FC** | **padj** | **EHB-1** | **EHB-2** | **EHB-3** | **EHB-4** | **ADB-1** | **ADB-2** | **ADB-3** | **ADB-4** |
| **ENSBTAG00000017294** | ORM1 | -21.90 | 1.87E-05 | 0.00 | 0.00 | 0.00 | 0.00 | 0.69 | 0.00 | 0.00 | 346.81 |
| **ENSBTAG00000000707** | WISP1 | -19.53 | 2.00E-04 | 0.00 | 0.00 | 0.00 | 0.00 | 0.00 | 0.00 | 0.00 | 71.91 |
| **ENSBTAG00000045714** | DSPP | -19.16 | 6.37E-05 | 0.00 | 0.00 | 0.00 | 0.00 | 0.00 | 0.00 | 0.00 | 60.08 |
| **ENSBTAG00000021408** | FMO1 | -18.36 | 2.29E-08 | 0.00 | 0.00 | 0.00 | 0.00 | 0.00 | 10.40 | 20.91 | 3.64 |
| **ENSBTAG00000022759** | PAG11 | -16.06 | 2.69E-03 | 0.00 | 0.00 | 0.00 | 0.00 | 6.25 | 0.00 | 0.00 | 0.00 |
| **ENSBTAG00000049405** | | -14.27 | 2.04E-03 | 0.00 | 0.00 | 0.00 | 0.00 | 2.08 | 0.00 | 0.00 | 0.00 |
| **ENSBTAG00000049599** | FAM163A | -8.82 | 1.96E-05 | 0.00 | 0.00 | 0.00 | 0.00 | 47.21 | 64.74 | 93.23 | 152.93 |
| **ENSBTAG00000052530** | | -8.71 | 4.85E-02 | 0.00 | 0.00 | 0.00 | 0.00 | 0.00 | 2.31 | 162.06 | 167.49 |
| **ENSBTAG00000038700** | FAM124B | -8.00 | 1.49E-03 | 0.00 | 0.00 | 0.00 | 0.00 | 20.14 | 78.61 | 35.72 | 69.18 |
| **ENSBTAG00000013047** | GRM7 | -7.89 | 1.56E-04 | 0.00 | 0.00 | 0.00 | 0.00 | 28.47 | 65.89 | 27.88 | 65.54 |
| **ENSBTAG00000027490** | OR5A1 | -7.75 | 3.44E-05 | 0.00 | 0.00 | 0.00 | 0.00 | 6.94 | 82.08 | 72.32 | 10.01 |
| **ENSBTAG00000017043** | GPR1 | -7.67 | 3.81E-03 | 0.00 | 0.00 | 0.00 | 0.00 | 6.94 | 58.96 | 53.15 | 42.78 |
| **ENSBTAG00000047203** | TSPAN19 | -7.63 | 2.81E-04 | 0.00 | 1.02 | 0.00 | 0.00 | 41.66 | 16.18 | 56.63 | 144.73 |
| **ENSBTAG00000046298** | SYCE3 | -7.53 | 5.73E-05 | 0.00 | 0.00 | 0.00 | 0.00 | 13.19 | 27.74 | 46.18 | 59.17 |
| **ENSBTAG00000052951** | | -7.52 | 2.55E-03 | 0.00 | 0.00 | 0.00 | 0.00 | 18.05 | 33.52 | 44.44 | 50.07 |
| **ENSBTAG00000000970** | ASIC1 | -7.48 | 6.17E-04 | 0.00 | 0.00 | 0.00 | 0.00 | 42.35 | 32.37 | 24.40 | 42.78 |
| **ENSBTAG00000002199** | CORIN | -7.41 | 2.30E-05 | 0.00 | 0.00 | 0.00 | 0.00 | 13.19 | 11.56 | 26.14 | 83.75 |
| **ENSBTAG00000021639** | ARC | -7.40 | 1.55E-03 | 0.00 | 0.00 | 0.00 | 0.00 | 4.86 | 12.72 | 88.87 | 27.31 |
| **ENSBTAG00000046630** | | -7.30 | 9.36E-04 | 4.03 | 0.00 | 8.61 | 7.53 | 261.07 | 4.62 | 518.41 | 2394.93 |
| **ENSBTAG00000040461** | | -7.26 | 8.32E-03 | 0.00 | 0.00 | 0.00 | 0.00 | 15.97 | 5.78 | 21.78 | 78.28 |
| **ENSBTAG00000033748** | IL18RAP | -7.18 | 5.95E-03 | 0.00 | 0.00 | 0.00 | 0.00 | 6.94 | 24.28 | 57.50 | 26.40 |
| **ENSBTAG00000051262** | | -7.18 | 2.28E-03 | 0.00 | 0.00 | 0.00 | 0.00 | 14.58 | 46.24 | 27.88 | 26.40 |
| **ENSBTAG00000000470** | IBSP | -7.11 | 4.05E-02 | 0.00 | 0.00 | 0.00 | 0.00 | 29.86 | 0.00 | 28.75 | 50.98 |
| **ENSBTAG00000001764** | NCAN | -7.10 | 2.80E-04 | 0.00 | 0.00 | 0.00 | 0.00 | 5.55 | 35.84 | 60.99 | 6.37 |
| **ENSBTAG00000006141** | WDR93 | -7.09 | 3.81E-02 | 0.00 | 0.00 | 0.00 | 0.00 | 11.11 | 0.00 | 55.76 | 40.96 |
| **ENSBTAG00000044012** | TMEM196 | -7.02 | 7.82E-09 | 3.02 | 0.00 | 3.69 | 0.00 | 107.62 | 252.01 | 359.84 | 135.63 |
| **ENSBTAG00000001231** | CDHR1 | -6.99 | 1.35E-02 | 0.00 | 0.00 | 0.00 | 0.00 | 14.58 | 20.81 | 0.87 | 64.63 |
| **ENSBTAG00000015315** | | -6.95 | 1.08E-02 | 0.00 | 0.00 | 0.00 | 0.00 | 18.05 | 21.96 | 45.31 | 12.74 |
| **ENSBTAG00000001243** | AP3M2 | -6.93 | 1.82E-10 | 3.02 | 1.02 | 0.00 | 0.00 | 132.62 | 85.55 | 201.27 | 97.40 |
| **ENSBTAG00000000606** | SYT10 | -6.88 | 2.11E-02 | 0.00 | 0.00 | 0.00 | 0.00 | 25.00 | 24.28 | 21.78 | 22.76 |
| **ENSBTAG00000006350** | SLC22A7 | -6.88 | 1.72E-04 | 0.00 | 0.00 | 0.00 | 0.00 | 3.47 | 34.68 | 35.72 | 20.03 |

| | | | | | | | | | | | |
|---|---|---|---|---|---|---|---|---|---|---|---|
| **ENSBTAG00000009837** | ANKS4B | -6.85 | 2.40E-02 | 0.00 | 0.00 | 0.00 | 0.00 | 46.52 | 27.74 | 17.43 | 0.00 |
| **ENSBTAG00000005259** | UCP3 | -6.74 | 1.70E-04 | 0.00 | 0.00 | 2.46 | 0.00 | 27.77 | 17.34 | 46.18 | 142.00 |
| **ENSBTAG00000002963** | SAA4 | -6.72 | 6.02E-03 | 0.00 | 0.00 | 0.00 | 0.00 | 37.49 | 13.87 | 18.30 | 13.65 |
| **ENSBTAG00000012121** | NOXA1 | -6.70 | 7.41E-03 | 0.00 | 0.00 | 0.00 | 0.00 | 0.00 | 16.18 | 36.59 | 30.04 |
| **ENSBTAG00000018817** | POU4F3 | -6.67 | 5.41E-03 | 0.00 | 0.00 | 0.00 | 0.00 | 43.05 | 23.12 | 4.36 | 10.01 |
| **ENSBTAG00000016365** | ABCG5 | -6.62 | 3.17E-05 | 0.00 | 1.02 | 0.00 | 0.00 | 15.28 | 53.18 | 23.52 | 37.32 |
| **ENSBTAG00000054666** | | -6.62 | 1.87E-03 | 0.00 | 0.00 | 0.00 | 0.00 | 8.33 | 10.40 | 42.69 | 16.38 |
| **ENSBTAG00000017664** | HGF | -6.61 | 4.77E-02 | 0.00 | 0.00 | 0.00 | 0.00 | 7.64 | 64.74 | 0.00 | 5.46 |
| **ENSBTAG00000013984** | KL | -6.58 | 2.40E-03 | 0.00 | 0.00 | 0.00 | 0.00 | 37.49 | 9.25 | 20.91 | 8.19 |
| **ENSBTAG00000006451** | GAP43 | -6.57 | 6.88E-03 | 0.00 | 0.00 | 0.00 | 0.00 | 9.72 | 17.34 | 12.20 | 36.41 |
| **ENSBTAG00000019146** | OSBP2 | -6.57 | 4.66E-03 | 0.00 | 0.00 | 0.00 | 0.00 | 2.78 | 5.78 | 13.07 | 53.71 |
| **ENSBTAG00000019942** | PABPC5 | -6.56 | 4.05E-02 | 0.00 | 0.00 | 0.00 | 0.00 | 22.91 | 21.96 | 0.00 | 30.04 |
| **ENSBTAG00000001067** | GRM2 | -6.55 | 3.87E-04 | 0.00 | 0.00 | 0.00 | 1.08 | 58.32 | 4.62 | 31.37 | 28.22 |
| **ENSBTAG00000018157** | IFT172 | -6.53 | 3.23E-02 | 0.00 | 0.00 | 0.00 | 0.00 | 27.77 | 0.00 | 41.82 | 3.64 |
| **ENSBTAG00000015936** | BRINP1 | -6.52 | 8.98E-04 | 1.01 | 0.00 | 1.23 | 0.00 | 29.16 | 3.47 | 41.82 | 122.89 |
| **ENSBTAG00000046343** | CCNJL | -6.43 | 2.45E-03 | 0.00 | 0.00 | 0.00 | 0.00 | 25.69 | 27.74 | 4.36 | 10.92 |
| **ENSBTAG00000013319** | SMCO1 | -6.43 | 1.28E-03 | 0.00 | 0.00 | 0.00 | 0.00 | 6.25 | 0.00 | 33.11 | 29.13 |
| **ENSBTAG00000017136** | SSTR2 | -6.41 | 1.47E-02 | 0.00 | 0.00 | 0.00 | 0.00 | 17.36 | 21.96 | 15.68 | 12.74 |
| **ENSBTAG00000037786** | NALCN | -6.40 | 4.69E-03 | 0.00 | 0.00 | 1.23 | 0.00 | 40.97 | 3.47 | 51.41 | 14.56 |
| **ENSBTAG00000017318** | TMEM178A | -6.38 | 1.73E-06 | 0.00 | 0.00 | 8.61 | 0.00 | 98.60 | 209.24 | 259.64 | 115.60 |
| **ENSBTAG00000004580** | MS4A1 | -6.36 | 3.99E-03 | 0.00 | 1.02 | 0.00 | 0.00 | 20.14 | 5.78 | 43.56 | 38.23 |
| **ENSBTAG00000030892** | | -6.35 | 1.29E-03 | 0.00 | 0.00 | 1.23 | 0.00 | 14.58 | 15.03 | 26.14 | 50.98 |
| **ENSBTAG00000010145** | SLC12A1 | -6.34 | 1.63E-02 | 0.00 | 0.00 | 0.00 | 0.00 | 21.52 | 21.96 | 5.23 | 15.47 |
| **ENSBTAG00000004674** | HAAO | -6.31 | 3.04E-02 | 0.00 | 0.00 | 0.00 | 0.00 | 11.11 | 15.03 | 9.58 | 27.31 |
| **ENSBTAG00000053503** | | -6.27 | 6.53E-03 | 0.00 | 1.02 | 0.00 | 0.00 | 6.25 | 8.09 | 28.75 | 58.26 |
| **ENSBTAG00000021833** | GJA8 | -6.27 | 2.86E-02 | 0.00 | 0.00 | 0.00 | 0.00 | 9.03 | 0.00 | 33.98 | 18.21 |
| **ENSBTAG00000040460** | KRT23 | -6.27 | 2.19E-02 | 0.00 | 0.00 | 0.00 | 0.00 | 24.30 | 10.40 | 0.00 | 26.40 |
| **ENSBTAG00000049009** | | -6.27 | 2.12E-02 | 0.00 | 0.00 | 0.00 | 0.00 | 20.14 | 1.16 | 27.88 | 11.83 |
| **ENSBTAG00000000066** | LRRN4 | -6.24 | 8.21E-03 | 0.00 | 0.00 | 0.00 | 0.00 | 27.08 | 9.25 | 3.49 | 20.03 |
| **ENSBTAG00000010089** | CALCA | -6.16 | 1.22E-02 | 0.00 | 0.00 | 0.00 | 1.08 | 38.19 | 38.15 | 17.43 | 0.00 |
| **ENSBTAG00000040394** | PRSS27 | -6.12 | 7.98E-03 | 0.00 | 0.00 | 0.00 | 0.00 | 5.55 | 26.59 | 9.58 | 13.65 |
| **ENSBTAG00000049077** | | -6.07 | 3.68E-02 | 0.00 | 0.00 | 0.00 | 0.00 | 2.78 | 26.59 | 9.58 | 14.56 |
| **ENSBTAG00000050189** | | -6.05 | 2.92E-02 | 0.00 | 0.00 | 0.00 | 0.00 | 6.94 | 16.18 | 14.81 | 14.56 |

| | | | | | | | | | | | |
|---|---|---|---|---|---|---|---|---|---|---|---|
| **ENSBTAG00000004955** | SYTL5 | -6.04 | 8.23E-06 | 5.04 | 0.00 | 0.00 | 0.00 | 75.68 | 169.94 | 48.79 | 50.07 |
| **ENSBTAG00000000437** | FFAR4 | -6.01 | 2.46E-02 | 0.00 | 0.00 | 0.00 | 0.00 | 11.80 | 0.00 | 14.81 | 24.58 |
| **ENSBTAG00000016012** | WNT3 | -6.01 | 3.11E-02 | 0.00 | 1.02 | 0.00 | 0.00 | 43.74 | 12.72 | 13.94 | 13.65 |
| **ENSBTAG00000020580** | TCN1 | -5.96 | 1.17E-02 | 0.00 | 0.00 | 0.00 | 1.08 | 20.14 | 5.78 | 44.44 | 10.92 |
| **ENSBTAG00000015483** | CCR8 | -5.94 | 1.38E-02 | 4.03 | 0.00 | 0.00 | 0.00 | 6.25 | 114.45 | 89.74 | 41.87 |
| **ENSBTAG00000047321** | SPRR4 | -5.93 | 4.40E-03 | 0.00 | 0.00 | 1.23 | 0.00 | 13.19 | 21.96 | 37.47 | 7.28 |
| **ENSBTAG00000003959** | ARHGAP24 | -5.92 | 4.65E-02 | 0.00 | 0.00 | 0.00 | 1.08 | 47.21 | 21.96 | 1.74 | 8.19 |
| **ENSBTAG00000049550** | | -5.91 | 2.12E-03 | 0.00 | 1.02 | 0.00 | 0.00 | 20.14 | 13.87 | 19.17 | 25.49 |
| **ENSBTAG00000015525** | SLC39A12 | -5.87 | 1.88E-10 | 7.05 | 1.02 | 7.38 | 0.00 | 115.26 | 369.93 | 268.36 | 140.18 |
| **ENSBTAG00000039340** | SCN4B | -5.84 | 3.19E-02 | 0.00 | 0.00 | 0.00 | 0.00 | 1.39 | 26.59 | 12.20 | 5.46 |
| **ENSBTAG00000001952** | PROM2 | -5.75 | 2.40E-02 | 0.00 | 1.02 | 0.00 | 0.00 | 4.17 | 26.59 | 28.75 | 10.92 |
| **ENSBTAG00000046755** | | -5.75 | 1.94E-03 | 1.01 | 0.00 | 0.00 | 1.08 | 10.42 | 30.06 | 45.31 | 30.04 |
| **ENSBTAG00000012706** | RTP1 | -5.71 | 2.13E-02 | 0.00 | 0.00 | 1.23 | 0.00 | 11.11 | 10.40 | 24.40 | 22.76 |
| **ENSBTAG00000017507** | SV2C | -5.69 | 1.38E-02 | 0.00 | 0.00 | 0.00 | 0.00 | 4.17 | 8.09 | 9.58 | 19.12 |
| **ENSBTAG00000007090** | MYH2 | -5.60 | 2.02E-03 | 0.00 | 1.02 | 0.00 | 0.00 | 12.50 | 3.47 | 40.08 | 7.28 |
| **ENSBTAG00000006998** | RASGRP4 | -5.44 | 2.79E-03 | 0.00 | 0.00 | 2.46 | 0.00 | 16.66 | 12.72 | 36.59 | 29.13 |
| **ENSBTAG00000048611** | | -5.40 | 5.36E-03 | 0.00 | 0.00 | 1.23 | 0.00 | 22.91 | 11.56 | 8.71 | 11.83 |
| **ENSBTAG00000053833** | | -5.39 | 2.05E-02 | 0.00 | 0.00 | 1.23 | 0.00 | 0.00 | 13.87 | 19.17 | 21.85 |
| **ENSBTAG00000008017** | UNC79 | -5.36 | 1.04E-03 | 2.02 | 0.00 | 2.46 | 0.00 | 1.39 | 39.31 | 102.81 | 36.41 |
| **ENSBTAG00000052032** | | -5.36 | 1.49E-02 | 0.00 | 0.00 | 2.46 | 0.00 | 6.94 | 36.99 | 17.43 | 29.13 |
| **ENSBTAG00000013614** | TMEM38A | -5.32 | 2.02E-02 | 2.02 | 0.00 | 0.00 | 0.00 | 23.61 | 26.59 | 11.33 | 23.67 |
| **ENSBTAG00000031573** | NMRK2 | -5.30 | 2.32E-02 | 0.00 | 0.00 | 0.00 | 0.00 | 2.78 | 0.00 | 15.68 | 12.74 |
| **ENSBTAG00000020755** | SELP | -5.28 | 2.27E-02 | 0.00 | 0.00 | 3.69 | 0.00 | 26.38 | 16.18 | 47.05 | 46.42 |
| **ENSBTAG00000019172** | IGSF21 | -5.23 | 3.21E-02 | 0.00 | 0.00 | 0.00 | 0.00 | 4.86 | 8.09 | 6.97 | 10.01 |
| **ENSBTAG00000024394** | SEMA3D | -5.22 | 3.76E-03 | 0.00 | 1.02 | 0.00 | 5.38 | 27.77 | 126.01 | 86.26 | 0.00 |
| **ENSBTAG00000016662** | CPS1 | -5.16 | 5.37E-03 | 0.00 | 0.00 | 0.00 | 9.69 | 72.21 | 98.26 | 162.93 | 13.65 |
| **ENSBTAG00000053317** | YPEL4 | -5.13 | 1.30E-06 | 3.02 | 1.02 | 6.15 | 1.08 | 29.86 | 157.22 | 97.58 | 103.77 |
| **ENSBTAG00000053102** | | -5.13 | 5.45E-05 | 9.07 | 2.03 | 12.31 | 3.23 | 180.53 | 21.96 | 252.67 | 469.70 |
| **ENSBTAG00000019975** | IL7R | -5.06 | 2.34E-03 | 0.00 | 0.00 | 2.46 | 1.08 | 27.77 | 20.81 | 33.98 | 29.13 |
| **ENSBTAG00000001324** | SLCO2A1 | -5.05 | 3.67E-02 | 0.00 | 0.00 | 1.23 | 0.00 | 23.61 | 3.47 | 6.97 | 9.10 |
| **ENSBTAG00000016359** | SLCO4C1 | -4.90 | 2.09E-04 | 11.08 | 0.00 | 12.31 | 1.08 | 91.65 | 242.77 | 145.50 | 247.59 |
| **ENSBTAG00000049108** | TRAT1 | -4.89 | 6.42E-04 | 23.17 | 6.09 | 0.00 | 0.00 | 95.12 | 298.26 | 209.98 | 268.53 |
| **ENSBTAG00000035013** | | -4.86 | 2.60E-03 | 1.01 | 0.00 | 3.69 | 2.15 | 34.72 | 57.80 | 54.02 | 48.24 |

| | | | | | | | | | | | |
|---|---|---|---|---|---|---|---|---|---|---|---|
| **ENSBTAG00000039049** | RAI2 | -4.86 | 2.50E-02 | 3.02 | 0.00 | 0.00 | 0.00 | 27.08 | 9.25 | 18.30 | 35.50 |
| **ENSBTAG00000005938** | BARHL2 | -4.85 | 3.10E-03 | 9.07 | 0.00 | 3.69 | 0.00 | 49.30 | 58.96 | 176.00 | 85.57 |
| **ENSBTAG00000040202** | | -4.85 | 5.40E-10 | 13.10 | 2.03 | 23.38 | 5.38 | 205.52 | 223.11 | 249.19 | 576.20 |
| **ENSBTAG00000024058** | EGR4 | -4.85 | 1.51E-02 | 4.03 | 0.00 | 0.00 | 0.00 | 31.25 | 20.81 | 25.27 | 41.87 |
| **ENSBTAG00000046635** | | -4.84 | 1.82E-02 | 0.00 | 0.00 | 24.61 | 0.00 | 40.27 | 10.40 | 613.38 | 35.50 |
| **ENSBTAG00000032908** | TMEM253 | -4.78 | 3.76E-02 | 1.01 | 0.00 | 0.00 | 0.00 | 9.03 | 16.18 | 0.87 | 10.01 |
| **ENSBTAG00000012629** | ZNF362 | -4.74 | 4.79E-07 | 5.04 | 7.11 | 8.61 | 6.46 | 132.62 | 105.20 | 253.54 | 233.03 |
| **ENSBTAG00000003992** | LHX4 | -4.73 | 2.35E-02 | 6.05 | 0.00 | 0.00 | 0.00 | 22.22 | 55.49 | 44.44 | 40.96 |
| **ENSBTAG00000020843** | PCSK1 | -4.72 | 2.21E-02 | 5.04 | 0.00 | 0.00 | 4.31 | 39.58 | 47.40 | 136.79 | 24.58 |
| **ENSBTAG00000015157** | GRAP2 | -4.69 | 1.68E-03 | 0.00 | 0.00 | 12.31 | 0.00 | 77.77 | 83.23 | 57.50 | 89.21 |
| **ENSBTAG00000051714** | | -4.59 | 4.55E-03 | 16.12 | 0.00 | 2.46 | 0.00 | 54.16 | 194.21 | 101.94 | 100.13 |
| **ENSBTAG00000025400** | PARP4 | -4.57 | 1.52E-02 | 1.01 | 0.00 | 6.15 | 3.23 | 37.49 | 90.17 | 66.22 | 50.07 |
| **ENSBTAG00000047818** | FAM19A2 | -4.56 | 4.56E-04 | 1.01 | 1.02 | 29.54 | 0.00 | 113.18 | 212.71 | 304.95 | 100.13 |
| **ENSBTAG00000011733** | GIPC2 | -4.50 | 3.80E-18 | 51.39 | 15.23 | 32.00 | 19.37 | 510.34 | 1175.68 | 649.98 | 328.61 |
| **ENSBTAG00000012827** | | -4.44 | 3.95E-02 | 0.00 | 0.00 | 2.46 | 3.23 | 33.33 | 33.52 | 14.81 | 40.05 |
| **ENSBTAG00000007506** | CACNG7 | -4.43 | 2.11E-03 | 4.03 | 5.08 | 0.00 | 2.15 | 28.47 | 76.30 | 49.66 | 92.85 |
| **ENSBTAG00000043996** | SAMD12 | -4.41 | 4.41E-11 | 33.25 | 20.31 | 41.84 | 7.53 | 229.13 | 633.50 | 598.57 | 728.22 |
| **ENSBTAG00000046430** | ZNF804B | -4.39 | 4.81E-02 | 2.02 | 0.00 | 12.31 | 0.00 | 118.73 | 72.83 | 32.24 | 72.82 |
| **ENSBTAG00000034069** | MAB21L1 | -4.38 | 7.34E-03 | 0.00 | 0.00 | 12.31 | 0.00 | 40.27 | 99.42 | 33.98 | 74.64 |
| **ENSBTAG00000050454** | | -4.28 | 6.19E-03 | 0.00 | 5.08 | 0.00 | 0.00 | 11.11 | 26.59 | 42.69 | 20.94 |
| **ENSBTAG00000016920** | EOMES | -4.27 | 4.66E-03 | 1.01 | 2.03 | 33.23 | 2.15 | 348.56 | 3.47 | 145.50 | 234.85 |
| **ENSBTAG00000018133** | SEMA3A | -4.18 | 3.69E-06 | 3.02 | 7.11 | 3.69 | 5.38 | 42.35 | 102.89 | 54.89 | 149.28 |
| **ENSBTAG00000021082** | TMEM125 | -4.18 | 2.33E-03 | 4.03 | 0.00 | 4.92 | 0.00 | 32.63 | 32.37 | 41.82 | 52.80 |
| **ENSBTAG00000003276** | PRKCH | -4.17 | 3.44E-05 | 36.27 | 0.00 | 4.92 | 11.84 | 212.47 | 181.50 | 309.31 | 258.52 |
| **ENSBTAG00000012394** | CCDC85A | -4.15 | 1.28E-03 | 10.08 | 1.02 | 2.46 | 0.00 | 28.47 | 49.71 | 48.79 | 115.60 |
| **ENSBTAG00000014443** | ODF4 | -4.14 | 2.52E-02 | 11.08 | 0.00 | 44.31 | 0.00 | 78.46 | 404.61 | 231.76 | 257.61 |
| **ENSBTAG00000004007** | MCF2 | -4.12 | 1.70E-04 | 6.05 | 6.09 | 2.46 | 0.00 | 65.96 | 50.87 | 81.03 | 58.26 |
| **ENSBTAG00000009197** | | -4.10 | 1.55E-03 | 680.11 | 133.00 | 671.97 | 189.44 | 9017.31 | 393.05 | 7050.42 | 12285.06 |
| **ENSBTAG00000002674** | GNGT1 | -4.08 | 2.21E-02 | 4.03 | 0.00 | 0.00 | 9.69 | 56.24 | 24.28 | 4.36 | 148.37 |
| **ENSBTAG00000027017** | SIX3 | -4.08 | 2.25E-06 | 9.07 | 23.35 | 50.46 | 11.84 | 349.25 | 279.76 | 426.93 | 539.79 |
| **ENSBTAG00000026344** | MAFA | -4.08 | 7.50E-03 | 12.09 | 0.00 | 2.46 | 2.15 | 24.30 | 0.00 | 99.33 | 160.21 |
| **ENSBTAG00000021807** | TAC3 | -4.08 | 3.86E-02 | 7.05 | 0.00 | 0.00 | 0.00 | 29.86 | 23.12 | 42.69 | 25.49 |
| **ENSBTAG00000048560** | TMEM233 | -4.01 | 3.10E-07 | 9.07 | 5.08 | 19.69 | 16.15 | 141.64 | 278.60 | 156.83 | 225.75 |

| | | | | | | | | | | | |
|---|---|---|---|---|---|---|---|---|---|---|---|
| **ENSBTAG00000032517** | BCL7A | -4.01 | 3.75E-04 | 4.03 | 11.17 | 3.69 | 0.00 | 68.04 | 72.83 | 78.42 | 87.39 |
| **ENSBTAG00000021885** | RSPO4 | -4.01 | 3.29E-03 | 40.30 | 0.00 | 0.00 | 0.00 | 156.92 | 112.13 | 214.34 | 169.31 |
| **ENSBTAG00000051889** | TDRD15 | -3.97 | 5.88E-03 | 7.05 | 17.26 | 0.00 | 0.00 | 69.43 | 141.04 | 44.44 | 130.17 |
| **ENSBTAG00000000189** | LHFPL3 | -3.96 | 2.31E-04 | 44.33 | 25.38 | 168.61 | 1.08 | 691.56 | 1088.98 | 1043.80 | 895.71 |
| **ENSBTAG00000044125** | | -3.94 | 4.97E-09 | 34.26 | 69.04 | 56.61 | 26.91 | 373.55 | 1132.91 | 573.30 | 783.75 |
| **ENSBTAG00000037550** | | -3.92 | 1.51E-02 | 1.01 | 4.06 | 2.46 | 0.00 | 0.69 | 36.99 | 11.33 | 65.54 |
| **ENSBTAG00000012043** | PADI3 | -3.91 | 1.02E-02 | 12.09 | 0.00 | 0.00 | 1.08 | 20.14 | 19.65 | 76.67 | 83.75 |
| **ENSBTAG00000003137** | PLPPR1 | -3.90 | 1.03E-02 | 3.02 | 0.00 | 7.38 | 0.00 | 15.97 | 39.31 | 64.47 | 32.77 |
| **ENSBTAG00000003103** | EIF4E1B | -3.89 | 1.57E-12 | 443.33 | 133.00 | 380.29 | 105.48 | 2664.85 | 4269.22 | 3690.75 | 5072.96 |
| **ENSBTAG00000026972** | MYF5 | -3.87 | 3.80E-18 | 324.44 | 256.86 | 358.14 | 99.02 | 1991.35 | 4783.65 | 4497.56 | 3931.47 |
| **ENSBTAG00000012150** | | -3.84 | 1.77E-02 | 9.07 | 0.00 | 1.23 | 0.00 | 31.25 | 47.40 | 40.08 | 30.95 |
| **ENSBTAG00000053893** | | -3.84 | 1.47E-08 | 498.75 | 181.73 | 674.43 | 132.39 | 5685.20 | 4277.31 | 6288.92 | 5036.55 |
| **ENSBTAG00000015449** | PPEF2 | -3.84 | 1.38E-03 | 6.05 | 0.00 | 9.85 | 3.23 | 25.00 | 100.57 | 54.89 | 90.12 |
| **ENSBTAG00000019421** | DACT1 | -3.83 | 2.97E-02 | 13.10 | 0.00 | 0.00 | 4.31 | 88.87 | 56.65 | 46.18 | 57.35 |
| **ENSBTAG00000009223** | SFN | -3.83 | 1.55E-03 | 15.11 | 0.00 | 17.23 | 11.84 | 90.26 | 18.50 | 262.26 | 253.97 |
| **ENSBTAG00000026309** | ZHX2 | -3.82 | 1.65E-02 | 2.02 | 0.00 | 12.31 | 0.00 | 11.80 | 71.67 | 70.57 | 43.69 |
| **ENSBTAG00000009219** | CADM2 | -3.82 | 2.81E-02 | 0.00 | 1.02 | 6.15 | 0.00 | 7.64 | 4.62 | 32.24 | 52.80 |
| **ENSBTAG00000045782** | BMP15 | -3.80 | 1.11E-06 | 1271.55 | 177.67 | 939.03 | 171.14 | 8818.04 | 9617.02 | 9364.55 | 7907.55 |
| **ENSBTAG00000053475** | | -3.78 | 4.36E-02 | 428.22 | 3.05 | 367.98 | 27.99 | 3594.56 | 277.45 | 1800.94 | 5672.83 |
| **ENSBTAG00000050197** | C10orf99 | -3.77 | 1.60E-02 | 0.00 | 0.00 | 13.54 | 0.00 | 36.80 | 21.96 | 93.23 | 27.31 |
| **ENSBTAG00000030317** | PPP1R26 | -3.76 | 2.58E-02 | 0.00 | 5.08 | 1.23 | 3.23 | 59.02 | 26.59 | 11.33 | 32.77 |
| **ENSBTAG00000013760** | COL4A6 | -3.76 | 1.79E-02 | 1.01 | 0.00 | 0.00 | 7.53 | 34.02 | 28.90 | 10.46 | 42.78 |
| **ENSBTAG00000036087** | ARMC2 | -3.72 | 1.36E-11 | 152.14 | 54.82 | 152.61 | 38.75 | 841.53 | 1205.74 | 1526.49 | 1676.72 |
| **ENSBTAG00000013920** | | -3.72 | 8.70E-05 | 13.10 | 9.14 | 52.92 | 0.00 | 127.76 | 375.71 | 238.73 | 239.40 |
| **ENSBTAG00000006722** | FGF16 | -3.71 | 2.59E-06 | 19.14 | 58.89 | 43.07 | 16.15 | 351.33 | 508.65 | 399.05 | 532.51 |
| **ENSBTAG00000020984** | RAPGEF4 | -3.68 | 1.46E-02 | 3.02 | 0.00 | 23.38 | 0.00 | 78.46 | 137.57 | 35.72 | 82.83 |
| **ENSBTAG00000022360** | SOX5 | -3.67 | 2.86E-04 | 0.00 | 0.00 | 13.54 | 6.46 | 33.33 | 94.79 | 67.96 | 51.89 |
| **ENSBTAG00000009478** | GDF9 | -3.66 | 1.75E-08 | 237.79 | 142.14 | 204.30 | 184.06 | 1496.98 | 3499.30 | 1931.63 | 2772.69 |
| **ENSBTAG00000047600** | ROS1 | -3.65 | 1.72E-04 | 12.09 | 6.09 | 56.61 | 5.38 | 119.43 | 439.29 | 176.00 | 263.98 |
| **ENSBTAG00000000418** | TOX3 | -3.65 | 1.53E-05 | 0.00 | 27.41 | 38.15 | 26.91 | 204.13 | 381.49 | 312.79 | 256.70 |
| **ENSBTAG00000004958** | PTPRN2 | -3.65 | 1.31E-02 | 1.01 | 0.00 | 1.23 | 4.31 | 11.11 | 20.81 | 28.75 | 20.94 |
| **ENSBTAG00000054147** | | -3.64 | 1.06E-03 | 39.30 | 14.21 | 82.46 | 15.07 | 47.21 | 45.09 | 247.44 | 1541.09 |
| **ENSBTAG00000046744** | PALM3 | -3.62 | 3.47E-06 | 23.17 | 0.00 | 20.92 | 3.23 | 120.12 | 172.25 | 94.10 | 191.16 |

| | | | | | | | | | | | |
|---|---|---|---|---|---|---|---|---|---|---|---|
| **ENSBTAG00000014300** | SLC6A5 | -3.61 | 2.29E-08 | 18.14 | 9.14 | 20.92 | 10.76 | 172.19 | 231.21 | 169.90 | 145.64 |
| **ENSBTAG00000012139** | SIX1 | -3.58 | 4.09E-04 | 73.55 | 4.06 | 60.30 | 0.00 | 331.89 | 284.38 | 473.98 | 558.00 |
| **ENSBTAG00000052496** | FGF14 | -3.55 | 3.71E-02 | 9.07 | 1.02 | 9.85 | 1.08 | 74.29 | 31.21 | 54.02 | 85.57 |
| **ENSBTAG00000051816** | | -3.55 | 6.26E-03 | 6.05 | 1.02 | 6.15 | 0.00 | 13.89 | 53.18 | 32.24 | 54.62 |
| **ENSBTAG00000010745** | THRA | -3.55 | 4.74E-02 | 7.05 | 0.00 | 0.00 | 6.46 | 32.63 | 27.74 | 67.09 | 31.86 |
| **ENSBTAG00000051802** | | -3.55 | 2.60E-02 | 0.00 | 0.00 | 4.92 | 10.76 | 41.66 | 38.15 | 53.15 | 49.15 |
| **ENSBTAG00000034366** | RGS2 | -3.53 | 9.44E-06 | 1367.27 | 395.96 | 2313.73 | 287.39 | 7097.48 | 17061.85 | 14700.28 | 11699.75 |
| **ENSBTAG00000006064** | C7H19orf57 | -3.53 | 9.71E-04 | 10.08 | 3.05 | 4.92 | 0.00 | 88.18 | 38.15 | 52.28 | 30.04 |
| **ENSBTAG00000013027** | CCK | -3.51 | 1.74E-02 | 8.06 | 0.00 | 0.00 | 8.61 | 36.11 | 35.84 | 42.69 | 76.46 |
| **ENSBTAG00000001322** | SAXO1 | -3.50 | 8.02E-09 | 793.96 | 375.65 | 776.58 | 328.29 | 4121.56 | 8569.65 | 5602.35 | 7514.31 |
| **ENSBTAG00000007216** | DPF1 | -3.49 | 1.70E-05 | 18.14 | 2.03 | 34.46 | 22.60 | 222.88 | 194.21 | 263.13 | 182.05 |
| **ENSBTAG00000001326** | SLC44A5 | -3.48 | 2.16E-05 | 24.18 | 12.18 | 16.00 | 0.00 | 71.52 | 310.97 | 132.43 | 70.09 |
| **ENSBTAG00000015251** | | -3.45 | 1.08E-04 | 154.16 | 53.81 | 86.15 | 52.74 | 712.39 | 1039.27 | 803.32 | 1247.08 |
| **ENSBTAG00000053445** | | -3.45 | 7.81E-03 | 0.00 | 5.08 | 2.46 | 9.69 | 25.69 | 54.33 | 54.02 | 54.62 |
| **ENSBTAG00000005104** | MGAT5B | -3.44 | 7.82E-08 | 11.08 | 16.24 | 61.54 | 9.69 | 181.92 | 343.34 | 260.51 | 275.81 |
| **ENSBTAG00000019892** | HAS2 | -3.44 | 1.90E-02 | 115.87 | 44.67 | 81.23 | 3.23 | 556.86 | 683.21 | 784.15 | 626.27 |
| **ENSBTAG00000034346** | BTG4 | -3.42 | 1.00E-04 | 3545.64 | 1071.11 | 7651.30 | 1505.82 | 16773.01 | 58927.49 | 30122.86 | 42014.61 |
| **ENSBTAG00000011266** | ZBTB16 | -3.41 | 1.27E-04 | 27.20 | 47.72 | 83.69 | 31.21 | 401.32 | 537.55 | 534.97 | 546.16 |
| **ENSBTAG00000000830** | GPR137B | -3.40 | 7.59E-08 | 124.94 | 36.55 | 179.68 | 58.12 | 775.57 | 1102.85 | 1150.09 | 1195.19 |
| **ENSBTAG00000014523** | DIRAS2 | -3.40 | 3.50E-03 | 184.39 | 2.03 | 98.46 | 26.91 | 767.93 | 1094.76 | 722.29 | 712.74 |
| **ENSBTAG00000002997** | ADAMTSL1 | -3.40 | 8.24E-04 | 23.17 | 11.17 | 36.92 | 21.53 | 272.18 | 156.06 | 278.81 | 271.26 |
| **ENSBTAG00000054056** | | -3.39 | 7.96E-05 | 9.07 | 13.20 | 13.54 | 0.00 | 49.30 | 90.17 | 134.18 | 101.95 |
| **ENSBTAG00000048635** | | -3.39 | 3.44E-03 | 2.02 | 1.02 | 6.15 | 3.23 | 13.19 | 26.59 | 42.69 | 45.51 |
| **ENSBTAG00000019580** | TCL1A | -3.39 | 9.55E-06 | 25.19 | 12.18 | 35.69 | 11.84 | 121.51 | 210.40 | 158.57 | 395.97 |
| **ENSBTAG00000022381** | TET3 | -3.38 | 2.28E-02 | 82.62 | 3.05 | 49.23 | 15.07 | 654.76 | 245.08 | 366.81 | 298.57 |
| **ENSBTAG00000012206** | SNX33 | -3.35 | 1.80E-02 | 5.04 | 8.12 | 3.69 | 0.00 | 68.74 | 67.05 | 6.10 | 30.95 |
| **ENSBTAG00000053831** | | -3.35 | 4.37E-06 | 53.40 | 37.56 | 70.15 | 5.38 | 120.12 | 426.58 | 676.12 | 473.34 |
| **ENSBTAG00000017420** | ZMAT4 | -3.34 | 1.78E-02 | 31.23 | 0.00 | 28.31 | 0.00 | 100.68 | 211.55 | 168.16 | 121.98 |
| **ENSBTAG00000006689** | TECTB | -3.34 | 2.38E-03 | 89.67 | 0.00 | 169.84 | 79.65 | 592.96 | 1062.39 | 921.82 | 853.84 |
| **ENSBTAG00000000623** | | -3.34 | 8.11E-13 | 2218.67 | 1317.82 | 3151.84 | 923.51 | 11675.22 | 22879.00 | 22287.42 | 20046.96 |
| **ENSBTAG00000016799** | ERN1 | -3.33 | 4.41E-03 | 11.08 | 22.34 | 9.85 | 1.08 | 163.86 | 33.52 | 95.84 | 154.75 |
| **ENSBTAG00000000074** | NFIA | -3.33 | 2.60E-03 | 27.20 | 38.58 | 6.15 | 0.00 | 136.09 | 163.00 | 309.31 | 119.25 |
| **ENSBTAG00000030838** | | -3.33 | 4.52E-02 | 6.05 | 6.09 | 1.23 | 4.31 | 9.03 | 10.40 | 38.34 | 121.07 |

| | | | | | | | | | | | |
|---|---|---|---|---|---|---|---|---|---|---|---|
| **ENSBTAG00000000886** | | -3.33 | 3.90E-02 | 12.09 | 0.00 | 4.92 | 1.08 | 24.30 | 105.20 | 17.43 | 35.50 |
| **ENSBTAG00000013525** | MPIG6B | -3.33 | 5.64E-04 | 465.50 | 64.98 | 363.06 | 27.99 | 2030.23 | 2483.15 | 2227.87 | 2513.27 |
| **ENSBTAG00000050101** | NRXN3 | -3.32 | 4.06E-04 | 26.20 | 9.14 | 29.54 | 8.61 | 120.81 | 161.84 | 244.83 | 205.72 |
| **ENSBTAG00000019946** | USP44 | -3.31 | 1.02E-06 | 94.71 | 12.18 | 135.38 | 48.44 | 340.22 | 1273.95 | 557.62 | 710.01 |
| **ENSBTAG00000037508** | EBF1 | -3.30 | 4.05E-07 | 65.49 | 19.29 | 7.38 | 77.50 | 300.65 | 396.52 | 476.59 | 496.10 |
| **ENSBTAG00000006188** | USH2A | -3.29 | 3.30E-10 | 56.42 | 15.23 | 118.15 | 62.43 | 379.80 | 664.72 | 598.57 | 821.98 |
| **ENSBTAG00000004769** | NEIL2 | -3.29 | 1.94E-04 | 5.04 | 8.12 | 60.30 | 34.44 | 61.80 | 197.68 | 362.45 | 426.92 |
| **ENSBTAG00000050418** | | -3.28 | 5.58E-03 | 0.00 | 12.18 | 1.23 | 0.00 | 15.28 | 24.28 | 25.27 | 67.36 |
| **ENSBTAG00000013219** | PEX5L | -3.28 | 3.07E-09 | 83.63 | 47.72 | 115.69 | 16.15 | 527.00 | 616.16 | 860.83 | 544.34 |
| **ENSBTAG00000030440** | | -3.26 | 2.03E-02 | 2.02 | 4.06 | 1.23 | 3.23 | 20.14 | 12.72 | 58.38 | 10.01 |
| **ENSBTAG00000018650** | HEPACAM | -3.25 | 3.44E-03 | 13.10 | 8.12 | 3.69 | 0.00 | 72.21 | 87.86 | 40.08 | 39.14 |
| **ENSBTAG00000047658** | PHF7 | -3.24 | 1.41E-06 | 2.02 | 43.66 | 13.54 | 18.30 | 190.94 | 124.85 | 197.78 | 222.11 |
| **ENSBTAG00000037964** | HIST3H2A | -3.23 | 2.57E-04 | 30.23 | 39.60 | 25.84 | 90.41 | 52.08 | 139.88 | 1004.59 | 548.90 |
| **ENSBTAG00000016377** | IPP | -3.22 | 3.71E-04 | 1.01 | 28.43 | 93.53 | 43.05 | 213.85 | 534.09 | 461.78 | 334.98 |
| **ENSBTAG00000005230** | SHOX2 | -3.22 | 1.80E-05 | 21.16 | 38.58 | 48.00 | 0.00 | 229.13 | 208.09 | 277.07 | 284.92 |
| **ENSBTAG00000019622** | TPBGL | -3.21 | 5.59E-03 | 9.07 | 0.00 | 3.69 | 0.00 | 32.63 | 21.96 | 25.27 | 38.23 |
| **ENSBTAG00000006823** | CMYA5 | -3.20 | 1.70E-02 | 114.86 | 4.06 | 132.92 | 1.08 | 410.35 | 851.99 | 399.92 | 668.14 |
| **ENSBTAG00000013995** | MAGEB3 | -3.19 | 4.31E-02 | 0.00 | 1.02 | 28.31 | 1.08 | 53.46 | 55.49 | 40.95 | 122.89 |
| **ENSBTAG00000026384** | ZAR1L | -3.18 | 8.45E-06 | 1244.35 | 510.68 | 1970.36 | 360.58 | 4869.36 | 11211.18 | 10930.24 | 10103.13 |
| **ENSBTAG00000009800** | MSI1 | -3.17 | 6.89E-08 | 22.17 | 14.21 | 45.54 | 23.68 | 211.08 | 224.27 | 299.72 | 209.36 |
| **ENSBTAG00000048053** | SLBP2 | -3.16 | 2.14E-06 | 577.34 | 149.24 | 694.12 | 266.94 | 2447.53 | 4520.08 | 4087.19 | 3996.10 |
| **ENSBTAG00000024061** | NOTO | -3.15 | 1.95E-02 | 53.40 | 0.00 | 22.15 | 0.00 | 241.63 | 167.62 | 159.44 | 101.95 |
| **ENSBTAG00000008091** | SELENBP1 | -3.15 | 1.93E-11 | 99.75 | 64.98 | 65.23 | 27.99 | 412.43 | 582.64 | 633.42 | 656.31 |
| **ENSBTAG00000007306** | RAB3C | -3.14 | 7.23E-10 | 63.48 | 24.37 | 48.00 | 52.74 | 231.91 | 544.49 | 321.50 | 568.01 |
| **ENSBTAG00000000502** | DAZL | -3.12 | 1.59E-03 | 84.64 | 27.41 | 40.61 | 0.00 | 227.05 | 347.97 | 293.62 | 462.42 |
| **ENSBTAG00000002733** | DBX1 | -3.10 | 1.06E-09 | 86.65 | 33.50 | 114.46 | 181.90 | 798.48 | 765.29 | 1004.59 | 1007.67 |
| **ENSBTAG00000030425** | ID3 | -3.10 | 3.87E-05 | 1133.52 | 619.31 | 1676.22 | 339.05 | 3149.50 | 11844.69 | 7628.95 | 9749.03 |
| **ENSBTAG00000007778** | TRPM3 | -3.10 | 2.84E-05 | 34.26 | 13.20 | 77.53 | 33.37 | 169.42 | 419.64 | 345.90 | 420.55 |
| **ENSBTAG00000011439** | EPB42 | -3.10 | 1.69E-02 | 1.01 | 9.14 | 3.69 | 6.46 | 42.35 | 60.11 | 20.04 | 51.89 |
| **ENSBTAG00000007403** | SLC7A3 | -3.09 | 1.11E-06 | 156.17 | 19.29 | 150.15 | 49.51 | 583.24 | 1164.12 | 681.34 | 756.44 |
| **ENSBTAG00000048279** | BCAR4 | -3.09 | 1.82E-10 | 141.06 | 89.34 | 198.14 | 82.88 | 601.99 | 1396.48 | 1165.78 | 1174.25 |
| **ENSBTAG00000015907** | IRX4 | -3.08 | 1.86E-02 | 26.20 | 0.00 | 0.00 | 1.08 | 33.33 | 48.55 | 65.35 | 86.48 |
| **ENSBTAG00000014225** | SLC5A9 | -3.08 | 1.41E-02 | 8.06 | 0.00 | 56.61 | 41.98 | 100.68 | 216.18 | 302.34 | 281.27 |

| | | | | | | | | | | | |
|---|---|---|---|---|---|---|---|---|---|---|---|
| **ENSBTAG00000025760** | | -3.07 | 3.27E-05 | 67.51 | 3.05 | 108.30 | 99.02 | 399.94 | 623.10 | 616.87 | 685.44 |
| **ENSBTAG00000050574** | | -3.06 | 2.11E-02 | 3.02 | 2.03 | 3.69 | 2.15 | 14.58 | 4.62 | 56.63 | 14.56 |
| **ENSBTAG00000019145** | MOS | -3.05 | 6.21E-04 | 362.72 | 70.05 | 845.49 | 187.29 | 1948.30 | 4537.42 | 2320.23 | 3308.85 |
| **ENSBTAG00000002929** | IRF4 | -3.04 | 9.96E-08 | 12.09 | 12.18 | 16.00 | 11.84 | 123.59 | 65.89 | 152.47 | 85.57 |
| **ENSBTAG00000043972** | SLC24A2 | -3.03 | 4.05E-02 | 24.18 | 0.00 | 12.31 | 0.00 | 36.80 | 101.73 | 115.01 | 45.51 |
| **ENSBTAG00000006844** | LEF1 | -3.01 | 6.09E-08 | 178.34 | 67.01 | 338.44 | 137.77 | 1042.19 | 910.95 | 2053.61 | 1816.00 |
| **ENSBTAG00000046162** | ZNF385C | -3.01 | 4.78E-02 | 7.05 | 0.00 | 13.54 | 0.00 | 40.27 | 31.21 | 44.44 | 48.24 |
| **ENSBTAG00000003827** | RIMS2 | -3.01 | 9.16E-07 | 125.95 | 39.60 | 183.38 | 207.74 | 731.83 | 1396.48 | 1247.68 | 1104.16 |
| **ENSBTAG00000031246** | CCNI2 | -3.01 | 3.39E-09 | 113.86 | 42.64 | 68.92 | 47.36 | 377.72 | 649.69 | 639.52 | 529.78 |
| **ENSBTAG00000020426** | TFAP2B | -3.00 | 1.06E-05 | 53.40 | 19.29 | 172.30 | 43.05 | 495.75 | 731.77 | 476.59 | 594.41 |
| **ENSBTAG00000001094** | MTUS2 | -3.00 | 9.55E-03 | 15.11 | 3.05 | 48.00 | 9.69 | 70.13 | 146.82 | 92.36 | 294.02 |
| **ENSBTAG00000004023** | KIAA1324L | -2.99 | 5.89E-05 | 16.12 | 11.17 | 32.00 | 29.06 | 220.80 | 144.50 | 238.73 | 96.49 |
| **ENSBTAG00000053626** | | -2.99 | 1.07E-02 | 8.06 | 48.73 | 17.23 | 5.38 | 156.92 | 149.13 | 187.33 | 138.36 |
| **ENSBTAG00000021779** | MGST2 | -2.98 | 1.17E-05 | 107.81 | 128.94 | 103.38 | 83.96 | 658.23 | 1063.55 | 911.36 | 724.58 |
| **ENSBTAG00000002908** | GNG4 | -2.97 | 5.37E-07 | 32.24 | 40.61 | 83.69 | 34.44 | 182.61 | 361.84 | 400.79 | 544.34 |
| **ENSBTAG00000002673** | FIGNL2 | -2.96 | 3.46E-02 | 3.02 | 0.00 | 3.69 | 1.08 | 18.75 | 19.65 | 10.46 | 10.92 |
| **ENSBTAG00000020620** | RGS2 | -2.96 | 2.04E-03 | 141.06 | 23.35 | 236.30 | 34.44 | 484.64 | 1343.31 | 776.31 | 779.19 |
| **ENSBTAG00000014020** | RTKN2 | -2.96 | 1.75E-02 | 3.02 | 28.43 | 32.00 | 6.46 | 127.06 | 143.35 | 107.17 | 163.85 |
| **ENSBTAG00000023377** | SH3BP5 | -2.96 | 2.93E-06 | 90.68 | 26.40 | 22.15 | 75.34 | 358.97 | 306.35 | 518.41 | 484.27 |
| **ENSBTAG00000020214** | LRRC8B | -2.94 | 4.15E-03 | 61.46 | 54.82 | 29.54 | 69.96 | 389.52 | 439.29 | 298.85 | 534.33 |
| **ENSBTAG00000004407** | KCNK2 | -2.94 | 4.01E-03 | 7.05 | 8.12 | 2.46 | 3.23 | 72.21 | 19.65 | 41.82 | 27.31 |
| **ENSBTAG00000051503** | EFCAB8 | -2.92 | 1.36E-02 | 0.00 | 6.09 | 8.61 | 2.15 | 9.03 | 33.52 | 20.91 | 62.81 |
| **ENSBTAG00000023752** | | -2.92 | 1.01E-03 | 30.23 | 10.15 | 121.84 | 7.53 | 137.48 | 463.57 | 261.38 | 416.00 |
| **ENSBTAG00000006727** | | -2.91 | 1.39E-02 | 5.04 | 1.02 | 18.46 | 1.08 | 21.52 | 76.30 | 27.88 | 63.72 |
| **ENSBTAG00000015592** | GPR84 | -2.90 | 5.33E-03 | 54.41 | 3.05 | 20.92 | 1.08 | 147.89 | 157.22 | 150.73 | 139.27 |
| **ENSBTAG00000017679** | PAK5 | -2.90 | 1.18E-05 | 21.16 | 107.62 | 105.84 | 66.73 | 585.32 | 502.87 | 385.11 | 774.64 |
| **ENSBTAG00000018838** | IQCA1L | -2.90 | 6.58E-03 | 6.05 | 2.03 | 3.69 | 3.23 | 39.58 | 16.18 | 36.59 | 19.12 |
| **ENSBTAG00000009832** | NOBOX | -2.90 | 7.75E-04 | 15.11 | 15.23 | 73.84 | 1.08 | 233.99 | 145.66 | 208.24 | 190.25 |
| **ENSBTAG00000013541** | LMO3 | -2.89 | 8.77E-03 | 20.15 | 20.31 | 68.92 | 0.00 | 72.21 | 366.46 | 124.59 | 247.59 |
| **ENSBTAG00000001729** | DUSP10 | -2.89 | 4.70E-07 | 336.53 | 67.01 | 219.07 | 117.32 | 1251.88 | 1419.61 | 1614.49 | 1180.63 |
| **ENSBTAG00000010694** | BICC1 | -2.89 | 4.12E-05 | 84.64 | 54.82 | 82.46 | 5.38 | 458.95 | 305.19 | 611.64 | 302.21 |
| **ENSBTAG00000046199** | | -2.88 | 7.26E-05 | 57.43 | 19.29 | 139.07 | 48.44 | 506.17 | 483.22 | 482.69 | 466.97 |
| **ENSBTAG00000052473** | | -2.88 | 2.40E-07 | 429.22 | 272.09 | 979.64 | 304.61 | 2681.52 | 5034.51 | 2980.66 | 3898.70 |

| | | | | | | | | | | | |
|---|---|---|---|---|---|---|---|---|---|---|---|
| **ENSBTAG00000048673** | | -2.88 | 8.21E-03 | 8.06 | 7.11 | 17.23 | 1.08 | 35.41 | 77.45 | 37.47 | 93.76 |
| **ENSBTAG00000016462** | TCF4 | -2.87 | 1.19E-03 | 26.20 | 12.18 | 223.99 | 39.83 | 343.00 | 499.41 | 862.57 | 493.37 |
| **ENSBTAG00000001021** | CYP1A1 | -2.87 | 3.75E-03 | 1031.75 | 198.99 | 156.30 | 58.12 | 3716.07 | 2273.91 | 1401.89 | 3139.54 |
| **ENSBTAG00000020643** | BARX2 | -2.85 | 4.29E-03 | 9.07 | 0.00 | 8.61 | 17.22 | 45.83 | 93.64 | 31.37 | 80.10 |
| **ENSBTAG00000053278** | | -2.84 | 2.54E-05 | 784.90 | 268.03 | 1335.32 | 240.03 | 3188.38 | 5628.71 | 5698.19 | 4287.39 |
| **ENSBTAG00000051340** | | -2.84 | 2.62E-03 | 52.39 | 2.03 | 27.08 | 5.38 | 93.74 | 135.26 | 195.17 | 197.53 |
| **ENSBTAG00000026307** | ZNF629 | -2.84 | 8.32E-03 | 11.08 | 11.17 | 7.38 | 4.31 | 65.27 | 30.06 | 73.19 | 74.64 |
| **ENSBTAG00000048544** | | -2.84 | 1.09E-07 | 52.39 | 85.28 | 72.61 | 18.30 | 293.70 | 397.67 | 446.10 | 496.10 |
| **ENSBTAG00000005729** | FBXL4 | -2.82 | 3.62E-11 | 153.15 | 60.92 | 203.07 | 121.63 | 1010.95 | 890.14 | 892.19 | 1017.69 |
| **ENSBTAG00000015952** | ADAP1 | -2.82 | 2.48E-13 | 86.65 | 55.84 | 72.61 | 87.18 | 563.80 | 463.57 | 568.08 | 544.34 |
| **ENSBTAG00000004305** | RGS16 | -2.81 | 8.74E-04 | 990.44 | 170.57 | 1126.10 | 139.93 | 4198.63 | 5306.18 | 4454.87 | 3098.57 |
| **ENSBTAG00000016879** | KIAA1147 | -2.80 | 1.52E-03 | 75.57 | 2.03 | 9.85 | 7.53 | 177.06 | 143.35 | 209.98 | 133.81 |
| **ENSBTAG00000054434** | | -2.79 | 6.22E-06 | 82.62 | 93.40 | 64.00 | 24.76 | 261.07 | 647.38 | 451.32 | 478.80 |
| **ENSBTAG00000017370** | ARFGEF3 | -2.79 | 4.18E-05 | 32.24 | 30.46 | 27.08 | 24.76 | 194.41 | 227.74 | 201.27 | 167.49 |
| **ENSBTAG00000048977** | | -2.79 | 2.36E-03 | 28.21 | 4.06 | 18.46 | 5.38 | 38.19 | 112.13 | 119.37 | 117.43 |
| **ENSBTAG00000049459** | | -2.78 | 1.46E-03 | 6.05 | 5.08 | 16.00 | 11.84 | 40.27 | 86.70 | 81.03 | 57.35 |
| **ENSBTAG00000009704** | CECR2 | -2.76 | 1.70E-07 | 245.85 | 96.45 | 345.83 | 298.15 | 1669.87 | 1784.91 | 1706.84 | 1520.16 |
| **ENSBTAG00000005301** | KLHL32 | -2.75 | 5.34E-03 | 41.31 | 0.00 | 22.15 | 12.92 | 95.82 | 137.57 | 129.82 | 151.11 |
| **ENSBTAG00000019675** | ATXN1 | -2.75 | 7.67E-06 | 48.36 | 18.27 | 78.77 | 23.68 | 222.88 | 379.18 | 224.79 | 307.67 |
| **ENSBTAG00000012590** | ACP7 | -2.75 | 3.55E-03 | 9.07 | 28.43 | 0.00 | 9.69 | 52.08 | 33.52 | 107.17 | 126.53 |
| **ENSBTAG00000023976** | ARHGAP25 | -2.74 | 7.77E-05 | 79.60 | 45.69 | 188.30 | 27.99 | 363.83 | 679.75 | 503.60 | 730.95 |
| **ENSBTAG00000013274** | | -2.73 | 9.66E-08 | 1850.90 | 1027.45 | 2814.62 | 741.61 | 6068.47 | 14260.79 | 10654.92 | 11768.93 |
| **ENSBTAG00000017205** | WISP3 | -2.73 | 6.42E-04 | 15.11 | 45.69 | 59.07 | 6.46 | 203.44 | 204.62 | 189.07 | 237.58 |
| **ENSBTAG00000007732** | ARPP21 | -2.73 | 3.47E-02 | 38.29 | 15.23 | 81.23 | 0.00 | 140.26 | 319.06 | 305.82 | 125.62 |
| **ENSBTAG00000037710** | | -2.73 | 2.68E-14 | 1432.76 | 651.80 | 931.64 | 539.25 | 3646.64 | 6054.13 | 5509.12 | 8335.38 |
| **ENSBTAG00000017354** | MPP7 | -2.72 | 7.08E-07 | 294.21 | 96.45 | 424.59 | 137.77 | 1105.38 | 2036.93 | 1690.29 | 1448.25 |
| **ENSBTAG00000018753** | TMEM163 | -2.72 | 2.00E-06 | 201.51 | 74.11 | 326.14 | 91.49 | 700.58 | 1327.12 | 1201.50 | 1319.90 |
| **ENSBTAG00000031052** | CDK5R2 | -2.71 | 5.00E-04 | 44.33 | 39.60 | 78.77 | 3.23 | 228.44 | 216.18 | 311.92 | 328.61 |
| **ENSBTAG00000004886** | ZAR1 | -2.71 | 1.19E-06 | 134.01 | 125.89 | 171.07 | 77.50 | 540.89 | 833.50 | 843.40 | 1099.61 |
| **ENSBTAG00000010856** | SLC8A2 | -2.69 | 1.42E-02 | 48.36 | 16.24 | 3.69 | 0.00 | 107.62 | 151.44 | 87.13 | 97.40 |
| **ENSBTAG00000021971** | SNCAIP | -2.69 | 1.67E-02 | 20.15 | 4.06 | 57.84 | 4.31 | 162.47 | 128.32 | 111.52 | 152.02 |
| **ENSBTAG00000039720** | TMEM225B | -2.68 | 1.40E-11 | 259.95 | 190.87 | 262.14 | 298.15 | 1037.33 | 2206.86 | 1697.26 | 1561.12 |
| **ENSBTAG00000034925** | KHDC3L | -2.68 | 2.37E-05 | 1883.15 | 745.21 | 3016.46 | 726.54 | 4633.98 | 10497.91 | 10792.58 | 15008.60 |

| | | | | | | | | | | | |
|---|---|---|---|---|---|---|---|---|---|---|---|
| **ENSBTAG00000008100** | GOLGA7B | -2.68 | 2.54E-05 | 100.76 | 75.13 | 129.22 | 59.20 | 343.70 | 635.82 | 500.12 | 858.39 |
| **ENSBTAG00000022819** | ONECUT2 | -2.68 | 3.09E-02 | 14.11 | 7.11 | 41.84 | 0.00 | 82.63 | 63.58 | 119.37 | 135.63 |
| **ENSBTAG00000010002** | IRF2 | -2.67 | 3.49E-02 | 7.05 | 4.06 | 3.69 | 6.46 | 9.72 | 4.62 | 33.98 | 87.39 |
| **ENSBTAG00000002289** | NLRP14 | -2.67 | 1.29E-07 | 412.10 | 196.96 | 198.14 | 124.86 | 1383.81 | 2100.51 | 1307.80 | 1131.47 |
| **ENSBTAG00000033319** | CD200 | -2.66 | 6.84E-07 | 111.84 | 22.34 | 135.38 | 48.44 | 232.60 | 631.19 | 528.87 | 616.26 |
| **ENSBTAG00000005215** | IL10RA | -2.66 | 4.74E-02 | 4.03 | 2.03 | 4.92 | 7.53 | 27.77 | 3.47 | 77.54 | 7.28 |
| **ENSBTAG00000000172** | PFKFB1 | -2.65 | 1.91E-02 | 44.33 | 8.12 | 11.08 | 5.38 | 35.41 | 124.85 | 82.77 | 191.16 |
| **ENSBTAG00000033726** | GRIP1 | -2.64 | 2.18E-02 | 22.17 | 2.03 | 23.38 | 0.00 | 46.52 | 129.48 | 46.18 | 74.64 |
| **ENSBTAG00000017644** | KIF17 | -2.64 | 1.42E-02 | 4.03 | 5.08 | 19.69 | 0.00 | 57.63 | 32.37 | 53.15 | 33.68 |
| **ENSBTAG00000000245** | NHSL1 | -2.64 | 1.81E-04 | 512.85 | 178.69 | 431.98 | 307.84 | 1671.96 | 2256.57 | 2907.47 | 2079.07 |
| **ENSBTAG00000019009** | | -2.62 | 1.08E-03 | 134.01 | 23.35 | 107.07 | 10.76 | 401.32 | 362.99 | 549.78 | 381.40 |
| **ENSBTAG00000011350** | CCNQ | -2.62 | 9.39E-11 | 1091.20 | 673.12 | 1281.16 | 733.00 | 2039.95 | 7767.37 | 6128.60 | 7340.45 |
| **ENSBTAG00000007553** | KCNA5 | -2.62 | 6.46E-03 | 12.09 | 0.00 | 73.84 | 24.76 | 334.67 | 101.73 | 135.92 | 103.77 |
| **ENSBTAG00000046918** | NPM2 | -2.61 | 1.68E-08 | 397.99 | 255.85 | 318.75 | 116.25 | 1276.88 | 2177.96 | 1346.13 | 1862.42 |
| **ENSBTAG00000007898** | CYBRD1 | -2.61 | 2.69E-07 | 86.65 | 70.05 | 52.92 | 11.84 | 342.31 | 375.71 | 302.34 | 332.25 |
| **ENSBTAG00000000476** | THPO | -2.60 | 6.59E-03 | 46.35 | 20.31 | 24.61 | 16.15 | 26.38 | 61.27 | 415.60 | 148.37 |
| **ENSBTAG00000001812** | H1FOO | -2.59 | 1.63E-03 | 541.06 | 210.16 | 1361.16 | 149.61 | 2125.35 | 4488.87 | 4000.06 | 2979.33 |
| **ENSBTAG00000006618** | HLF | -2.58 | 3.30E-10 | 216.63 | 78.18 | 231.37 | 206.66 | 747.10 | 1166.43 | 1326.09 | 1154.23 |
| **ENSBTAG00000053802** | | -2.58 | 2.86E-02 | 15.11 | 0.00 | 83.69 | 4.31 | 132.62 | 195.37 | 128.95 | 157.48 |
| **ENSBTAG00000019024** | JAZF1 | -2.57 | 1.46E-05 | 111.84 | 56.86 | 52.92 | 13.99 | 337.45 | 375.71 | 385.98 | 304.03 |
| **ENSBTAG00000010875** | MSX1 | -2.57 | 2.26E-14 | 785.90 | 466.01 | 1309.47 | 588.77 | 4063.24 | 5272.65 | 5053.44 | 4325.62 |
| **ENSBTAG00000015444** | LETM2 | -2.56 | 5.94E-03 | 17.13 | 24.37 | 60.30 | 0.00 | 163.17 | 137.57 | 172.51 | 122.89 |
| **ENSBTAG00000015737** | KPNA7 | -2.55 | 3.07E-03 | 925.96 | 257.88 | 1087.94 | 193.74 | 3187.69 | 4038.01 | 4175.19 | 3060.34 |
| **ENSBTAG00000006961** | | -2.55 | 1.45E-04 | 33.25 | 40.61 | 27.08 | 19.37 | 257.60 | 184.96 | 185.58 | 75.55 |
| **ENSBTAG00000043950** | LCA5 | -2.54 | 5.35E-03 | 56.42 | 0.00 | 35.69 | 19.37 | 126.37 | 134.10 | 284.91 | 104.68 |
| **ENSBTAG00000027348** | OOSP2 | -2.54 | 4.42E-04 | 2470.56 | 439.61 | 1660.22 | 452.07 | 4768.68 | 11807.69 | 6643.53 | 5980.50 |
| **ENSBTAG00000004710** | HNF1B | -2.54 | 3.17E-04 | 64.48 | 41.63 | 45.54 | 11.84 | 148.59 | 317.91 | 152.47 | 331.34 |
| **ENSBTAG00000025898** | TBC1D8 | -2.53 | 6.23E-07 | 211.59 | 106.60 | 61.54 | 93.64 | 547.83 | 598.82 | 1049.90 | 535.24 |
| **ENSBTAG00000008078** | TRIM59 | -2.52 | 1.42E-05 | 211.59 | 141.12 | 161.22 | 212.04 | 384.66 | 1780.29 | 750.17 | 1260.73 |
| **ENSBTAG00000021553** | PTPN14 | -2.52 | 1.17E-03 | 57.43 | 65.99 | 97.23 | 17.22 | 281.90 | 405.77 | 391.21 | 286.74 |
| **ENSBTAG00000007053** | ZFHX2 | -2.52 | 1.67E-02 | 13.10 | 3.05 | 14.77 | 7.53 | 94.43 | 41.62 | 54.89 | 28.22 |
| **ENSBTAG00000022580** | INKA2 | -2.50 | 2.59E-06 | 57.43 | 58.89 | 100.92 | 29.06 | 358.28 | 476.28 | 250.06 | 309.49 |
| **ENSBTAG00000018446** | GCA | -2.50 | 3.94E-07 | 325.44 | 135.03 | 211.68 | 89.34 | 892.91 | 1218.46 | 1137.90 | 1058.65 |

| | | | | | | | | | | | |
|---|---|---|---|---|---|---|---|---|---|---|---|
| **ENSBTAG00000050187** | | -2.49 | 2.38E-05 | 71.54 | 31.47 | 43.07 | 67.81 | 142.34 | 428.89 | 210.85 | 422.37 |
| **ENSBTAG00000020433** | NLRP1 | -2.49 | 3.68E-02 | 13.10 | 19.29 | 43.07 | 13.99 | 113.18 | 128.32 | 122.85 | 136.54 |
| **ENSBTAG00000008916** | CYFIP2 | -2.49 | 4.95E-04 | 143.07 | 58.89 | 124.30 | 97.95 | 585.32 | 575.70 | 491.40 | 727.31 |
| **ENSBTAG00000012436** | HES7 | -2.48 | 3.76E-03 | 40.30 | 1.02 | 39.38 | 11.84 | 73.60 | 110.98 | 127.21 | 202.08 |
| **ENSBTAG00000013279** | FOXP3 | -2.47 | 3.05E-02 | 8.06 | 7.11 | 3.69 | 0.00 | 31.94 | 12.72 | 40.95 | 19.12 |
| **ENSBTAG00000008756** | ELF3 | -2.46 | 9.19E-03 | 25.19 | 19.29 | 71.38 | 44.13 | 183.30 | 197.68 | 212.59 | 284.01 |
| **ENSBTAG00000020406** | GPC3 | -2.45 | 2.28E-03 | 100.76 | 4.06 | 116.92 | 43.05 | 206.22 | 524.84 | 464.39 | 254.88 |
| **ENSBTAG00000015273** | CAND2 | -2.45 | 7.70E-05 | 56.42 | 30.46 | 93.53 | 40.90 | 605.46 | 248.55 | 142.89 | 211.18 |
| **ENSBTAG00000006095** | PRDM13 | -2.45 | 7.03E-06 | 94.71 | 30.46 | 39.38 | 54.89 | 180.53 | 387.27 | 372.91 | 262.16 |
| **ENSBTAG00000021487** | CIART | -2.45 | 6.51E-09 | 990.44 | 339.10 | 452.90 | 613.52 | 2614.86 | 3714.33 | 3723.86 | 3053.97 |
| **ENSBTAG00000054800** | HES6 | -2.44 | 4.72E-03 | 65.49 | 41.63 | 28.31 | 58.12 | 149.98 | 48.55 | 400.79 | 455.14 |
| **ENSBTAG00000006005** | MPND | -2.44 | 4.85E-06 | 58.44 | 86.30 | 194.45 | 64.58 | 332.59 | 665.87 | 594.21 | 598.96 |
| **ENSBTAG00000021872** | RASGRP1 | -2.44 | 2.45E-02 | 5.04 | 12.18 | 66.46 | 21.53 | 152.06 | 116.76 | 162.93 | 136.54 |
| **ENSBTAG00000014179** | SLC12A5 | -2.44 | 2.69E-04 | 138.04 | 77.16 | 223.99 | 44.13 | 440.21 | 880.90 | 602.06 | 692.72 |
| **ENSBTAG00000014560** | HLX | -2.44 | 2.75E-02 | 11.08 | 0.00 | 14.77 | 20.45 | 77.07 | 67.05 | 41.82 | 63.72 |
| **ENSBTAG00000002253** | FKBP6 | -2.43 | 8.98E-05 | 54.41 | 161.43 | 94.76 | 45.21 | 363.83 | 442.76 | 574.18 | 542.52 |
| **ENSBTAG00000022006** | TRIM65 | -2.43 | 8.65E-04 | 123.93 | 153.31 | 136.61 | 17.22 | 381.19 | 691.31 | 940.11 | 312.22 |
| **ENSBTAG00000003936** | PNKD | -2.42 | 1.64E-02 | 6.05 | 19.29 | 4.92 | 19.37 | 118.73 | 6.94 | 58.38 | 81.01 |
| **ENSBTAG00000038322** | | -2.42 | 3.80E-02 | 30.23 | 0.00 | 6.15 | 5.38 | 72.21 | 46.24 | 29.62 | 75.55 |
| **ENSBTAG00000014580** | FCRLB | -2.41 | 2.51E-02 | 5.04 | 17.26 | 0.00 | 10.76 | 47.21 | 15.03 | 51.41 | 63.72 |
| **ENSBTAG00000025329** | IRF2BPL | -2.41 | 4.02E-02 | 4.03 | 13.20 | 14.77 | 0.00 | 19.44 | 32.37 | 38.34 | 79.19 |
| **ENSBTAG00000021709** | SPI1 | -2.41 | 3.49E-02 | 7.05 | 0.00 | 12.31 | 0.00 | 16.66 | 34.68 | 22.65 | 27.31 |
| **ENSBTAG00000001080** | SPAG17 | -2.40 | 6.59E-07 | 199.50 | 186.81 | 339.67 | 111.94 | 769.32 | 1595.32 | 792.87 | 1275.29 |
| **ENSBTAG00000010785** | FAS | -2.40 | 1.34E-02 | 9.07 | 5.08 | 12.31 | 6.46 | 25.00 | 90.17 | 15.68 | 42.78 |
| **ENSBTAG00000017063** | EPB41L4B | -2.39 | 5.46E-10 | 228.72 | 177.67 | 150.15 | 69.96 | 763.77 | 1120.19 | 777.18 | 624.45 |
| **ENSBTAG00000002594** | ZNF436 | -2.38 | 7.82E-09 | 87.66 | 139.09 | 66.46 | 75.34 | 328.42 | 458.94 | 372.91 | 767.36 |
| **ENSBTAG00000054813** | | -2.38 | 2.85E-07 | 152.14 | 58.89 | 180.91 | 71.04 | 411.05 | 698.24 | 700.51 | 601.69 |
| **ENSBTAG00000013881** | GJA4 | -2.37 | 7.28E-05 | 56.42 | 49.75 | 62.77 | 34.44 | 331.89 | 317.91 | 209.11 | 194.80 |
| **ENSBTAG00000014699** | FAM81A | -2.37 | 7.59E-08 | 485.65 | 409.15 | 820.88 | 176.52 | 1845.54 | 2858.86 | 2628.66 | 2472.30 |
| **ENSBTAG00000010351** | SNX25 | -2.37 | 6.05E-05 | 151.14 | 116.76 | 175.99 | 64.58 | 581.85 | 649.69 | 778.06 | 609.88 |
| **ENSBTAG00000017024** | PPARGC1A | -2.35 | 1.76E-02 | 61.46 | 119.80 | 127.99 | 1.08 | 299.26 | 440.45 | 488.79 | 352.28 |
| **ENSBTAG00000017251** | SLC26A8 | -2.35 | 4.85E-02 | 7.05 | 4.06 | 14.77 | 0.00 | 18.75 | 52.02 | 24.40 | 35.50 |
| **ENSBTAG00000023398** | UBASH3A | -2.35 | 1.58E-02 | 16.12 | 3.05 | 9.85 | 3.23 | 56.94 | 28.90 | 13.07 | 64.63 |

| | | | | | | | | | | | |
|---|---|---|---|---|---|---|---|---|---|---|---|
| **ENSBTAG00000023172** | ITSN2 | -2.34 | 1.44E-05 | 346.60 | 103.56 | 195.68 | 69.96 | 488.81 | 1238.11 | 993.26 | 917.56 |
| **ENSBTAG00000004725** | NLRP9 | -2.34 | 1.59E-05 | 162.22 | 63.96 | 98.46 | 69.96 | 402.02 | 606.92 | 512.31 | 483.36 |
| **ENSBTAG00000010597** | GGCT | -2.34 | 1.76E-05 | 471.54 | 377.68 | 531.66 | 206.66 | 958.87 | 2762.91 | 2027.48 | 2301.17 |
| **ENSBTAG00000018600** | VGLL4 | -2.34 | 1.69E-10 | 221.67 | 211.18 | 280.60 | 223.88 | 1000.53 | 1308.63 | 1413.22 | 1008.58 |
| **ENSBTAG00000039015** | TMEM145 | -2.33 | 1.65E-02 | 101.76 | 11.17 | 35.69 | 13.99 | 180.53 | 235.83 | 314.53 | 89.21 |
| **ENSBTAG00000001002** | TCF7 | -2.33 | 1.76E-05 | 68.51 | 20.31 | 52.92 | 40.90 | 169.42 | 216.18 | 286.65 | 246.68 |
| **ENSBTAG00000004613** | CSGALNACT1 | -2.33 | 9.03E-03 | 21.16 | 26.40 | 9.85 | 25.83 | 65.27 | 141.04 | 99.33 | 113.78 |
| **ENSBTAG00000037804** | IKZF2 | -2.33 | 1.49E-03 | 27.20 | 20.31 | 28.31 | 23.68 | 73.60 | 99.42 | 203.88 | 121.98 |
| **ENSBTAG00000032933** | TTC23L | -2.33 | 3.67E-02 | 4.03 | 10.15 | 30.77 | 23.68 | 42.35 | 163.00 | 54.02 | 83.75 |
| **ENSBTAG00000016711** | PPIF | -2.32 | 1.74E-02 | 540.06 | 113.71 | 430.75 | 175.45 | 944.29 | 1569.89 | 1779.16 | 2018.08 |
| **ENSBTAG00000046122** | NRARP | -2.32 | 4.96E-02 | 30.23 | 31.47 | 11.08 | 0.00 | 11.11 | 70.52 | 114.14 | 168.40 |
| **ENSBTAG00000030542** | KRT25 | -2.31 | 9.78E-03 | 55.42 | 68.02 | 39.38 | 106.56 | 105.54 | 717.89 | 494.89 | 22.76 |
| **ENSBTAG00000024021** | NRXN1 | -2.31 | 6.73E-03 | 37.28 | 35.53 | 120.61 | 67.81 | 301.34 | 412.70 | 281.42 | 297.66 |
| **ENSBTAG00000020445** | ZNF398 | -2.31 | 8.26E-07 | 178.34 | 70.05 | 239.99 | 121.63 | 683.22 | 705.18 | 961.03 | 661.77 |
| **ENSBTAG00000044155** | BSPRY | -2.30 | 6.57E-05 | 23.17 | 13.20 | 8.61 | 61.35 | 112.48 | 121.38 | 148.99 | 143.82 |
| **ENSBTAG00000020491** | SOX30 | -2.30 | 3.17E-03 | 21.16 | 12.18 | 30.77 | 13.99 | 87.49 | 113.29 | 72.32 | 110.14 |
| **ENSBTAG00000001879** | PER2 | -2.29 | 2.43E-06 | 53.40 | 34.52 | 39.38 | 24.76 | 186.08 | 232.36 | 167.29 | 159.30 |
| **ENSBTAG00000048876** | FAM117B | -2.29 | 9.67E-11 | 297.23 | 241.63 | 439.36 | 207.74 | 1337.29 | 1397.64 | 1541.30 | 1517.43 |
| **ENSBTAG00000017767** | GPR173 | -2.28 | 1.28E-03 | 96.73 | 38.58 | 84.92 | 93.64 | 377.02 | 320.22 | 436.51 | 395.06 |
| **ENSBTAG00000053612** | | -2.27 | 2.03E-03 | 142.07 | 25.38 | 109.53 | 44.13 | 335.36 | 84.39 | 449.58 | 679.06 |
| **ENSBTAG00000032686** | HOMEZ | -2.25 | 2.89E-03 | 27.20 | 47.72 | 66.46 | 47.36 | 135.40 | 208.09 | 261.38 | 293.11 |
| **ENSBTAG00000007490** | SULF2 | -2.25 | 3.77E-05 | 33.25 | 21.32 | 35.69 | 15.07 | 61.10 | 154.91 | 153.35 | 131.08 |
| **ENSBTAG00000054976** | MCTP1 | -2.25 | 4.24E-02 | 7.05 | 31.47 | 50.46 | 1.08 | 76.38 | 53.18 | 141.15 | 154.75 |
| **ENSBTAG00000003889** | PER1 | -2.24 | 2.26E-14 | 382.88 | 231.48 | 374.13 | 354.12 | 1752.50 | 1346.78 | 1804.43 | 1443.69 |
| **ENSBTAG00000005314** | MFN2 | -2.24 | 3.40E-03 | 17.13 | 45.69 | 19.69 | 50.59 | 250.65 | 45.09 | 177.74 | 154.75 |
| **ENSBTAG00000052808** | | -2.23 | 1.85E-02 | 17.13 | 30.46 | 9.85 | 4.31 | 63.88 | 25.43 | 112.40 | 89.21 |
| **ENSBTAG00000052435** | | -2.23 | 4.21E-02 | 2.02 | 10.15 | 6.15 | 6.46 | 22.22 | 55.49 | 27.01 | 11.83 |
| **ENSBTAG00000018249** | KCNN3 | -2.23 | 8.07E-05 | 48.36 | 41.63 | 39.38 | 101.18 | 310.37 | 203.46 | 325.86 | 238.49 |
| **ENSBTAG00000016691** | LACC1 | -2.22 | 6.20E-04 | 58.44 | 45.69 | 49.23 | 41.98 | 154.14 | 215.02 | 349.38 | 190.25 |
| **ENSBTAG00000013537** | FER1L6 | -2.21 | 1.57E-02 | 21.16 | 28.43 | 20.92 | 9.69 | 79.15 | 127.16 | 135.05 | 30.95 |
| **ENSBTAG00000007767** | TBX15 | -2.21 | 6.65E-03 | 29.22 | 13.20 | 13.54 | 41.98 | 67.35 | 84.39 | 174.26 | 127.44 |
| **ENSBTAG00000051941** | | -2.20 | 3.85E-02 | 8.06 | 11.17 | 11.08 | 34.44 | 75.68 | 67.05 | 113.27 | 41.87 |
| **ENSBTAG00000018438** | RRAGD | -2.20 | 1.50E-02 | 119.90 | 82.24 | 162.45 | 121.63 | 343.00 | 499.41 | 758.02 | 634.46 |

| | | | | | | | | | | | |
|---|---|---|---|---|---|---|---|---|---|---|---|
| **ENSBTAG00000017717** | RALYL | -2.19 | 2.04E-02 | 15.11 | 6.09 | 13.54 | 10.76 | 40.97 | 62.43 | 57.50 | 46.42 |
| **ENSBTAG00000022991** | NBEA | -2.19 | 2.17E-02 | 7.05 | 15.23 | 73.84 | 32.29 | 105.54 | 105.20 | 176.00 | 194.80 |
| **ENSBTAG00000044018** | | -2.18 | 8.24E-04 | 50.38 | 48.73 | 35.69 | 33.37 | 194.41 | 161.84 | 330.22 | 78.28 |
| **ENSBTAG00000004588** | KCNN4 | -2.17 | 1.53E-05 | 34.26 | 45.69 | 68.92 | 23.68 | 188.16 | 154.91 | 193.42 | 239.40 |
| **ENSBTAG00000012053** | ACCSL | -2.17 | 3.30E-03 | 1702.79 | 305.60 | 1043.64 | 278.78 | 4680.50 | 3036.89 | 4655.26 | 2632.51 |
| **ENSBTAG00000007719** | MFSD6 | -2.17 | 2.08E-02 | 25.19 | 16.24 | 27.08 | 34.44 | 151.36 | 78.61 | 115.88 | 115.60 |
| **ENSBTAG00000019674** | CLIP3 | -2.16 | 3.29E-02 | 102.77 | 26.40 | 67.69 | 7.53 | 329.81 | 75.14 | 235.25 | 274.90 |
| **ENSBTAG00000010947** | PHYHIPL | -2.16 | 4.17E-07 | 923.94 | 190.87 | 552.59 | 294.92 | 1471.99 | 3028.80 | 2221.77 | 2049.94 |
| **ENSBTAG00000003102** | GCC1 | -2.16 | 1.81E-02 | 57.43 | 62.95 | 111.99 | 13.99 | 307.59 | 278.60 | 276.20 | 236.67 |
| **ENSBTAG00000033967** | | -2.16 | 4.23E-03 | 19.14 | 24.37 | 61.54 | 11.84 | 66.66 | 135.26 | 90.61 | 226.66 |
| **ENSBTAG00000014915** | ETV5 | -2.15 | 7.80E-03 | 93.70 | 26.40 | 225.22 | 46.28 | 389.52 | 230.05 | 457.42 | 662.68 |
| **ENSBTAG00000051066** | ZNF71 | -2.15 | 4.16E-02 | 28.21 | 22.34 | 22.15 | 0.00 | 46.52 | 104.04 | 108.04 | 64.63 |
| **ENSBTAG00000038891** | | -2.15 | 1.82E-04 | 30.23 | 86.30 | 265.83 | 73.19 | 720.72 | 382.65 | 590.73 | 319.51 |
| **ENSBTAG00000003171** | SHANK2 | -2.15 | 2.90E-04 | 49.37 | 57.87 | 57.84 | 13.99 | 271.48 | 166.47 | 213.46 | 140.18 |
| **ENSBTAG00000038815** | | -2.14 | 4.38E-04 | 41.31 | 72.08 | 22.15 | 40.90 | 133.31 | 246.23 | 240.47 | 161.12 |
| **ENSBTAG00000005877** | FAM83A | -2.14 | 1.20E-02 | 4.03 | 5.08 | 8.61 | 11.84 | 24.30 | 15.03 | 47.92 | 41.87 |
| **ENSBTAG00000000357** | PRAG1 | -2.14 | 2.50E-03 | 55.42 | 89.34 | 30.77 | 46.28 | 302.73 | 234.67 | 224.79 | 215.73 |
| **ENSBTAG00000038945** | PADI6 | -2.13 | 4.74E-06 | 112.85 | 74.11 | 153.84 | 96.87 | 607.54 | 505.19 | 460.91 | 341.35 |
| **ENSBTAG00000014467** | | -2.13 | 4.99E-02 | 5.04 | 6.09 | 18.46 | 5.38 | 7.64 | 42.77 | 48.79 | 52.80 |
| **ENSBTAG00000012922** | PHF2 | -2.12 | 6.84E-03 | 10.08 | 20.31 | 20.92 | 24.76 | 126.37 | 31.21 | 40.95 | 130.17 |
| **ENSBTAG00000012855** | LPL | -2.11 | 4.87E-02 | 21.16 | 0.00 | 45.54 | 90.41 | 127.06 | 287.85 | 147.25 | 114.69 |
| **ENSBTAG00000020762** | HUNK | -2.11 | 3.89E-02 | 3.02 | 14.21 | 12.31 | 18.30 | 66.66 | 39.31 | 36.59 | 62.81 |
| **ENSBTAG00000018845** | DND1 | -2.11 | 8.30E-07 | 80.61 | 55.84 | 89.84 | 107.64 | 333.28 | 387.27 | 441.74 | 273.99 |
| **ENSBTAG00000005429** | | -2.10 | 2.49E-06 | 341.57 | 305.60 | 748.27 | 158.22 | 888.05 | 2001.09 | 2087.59 | 1668.53 |
| **ENSBTAG00000006472** | GPR143 | -2.09 | 9.83E-03 | 77.58 | 67.01 | 34.46 | 69.96 | 104.15 | 58.96 | 347.64 | 549.81 |
| **ENSBTAG00000014412** | ELAVL3 | -2.09 | 5.29E-04 | 178.34 | 41.63 | 82.46 | 45.21 | 240.93 | 190.75 | 552.39 | 494.28 |
| **ENSBTAG00000004194** | RALGDS | -2.09 | 9.03E-03 | 16.12 | 38.58 | 59.07 | 94.72 | 227.74 | 188.43 | 229.15 | 239.40 |
| **ENSBTAG00000016381** | REM2 | -2.09 | 1.64E-02 | 60.45 | 7.11 | 50.46 | 22.60 | 95.12 | 129.48 | 116.75 | 254.88 |
| **ENSBTAG00000027868** | DLG2 | -2.08 | 1.16E-05 | 79.60 | 24.37 | 67.69 | 54.89 | 181.92 | 211.55 | 264.00 | 301.30 |
| **ENSBTAG00000054829** | | -2.08 | 1.08E-02 | 16.12 | 6.09 | 22.15 | 21.53 | 35.41 | 115.60 | 87.13 | 40.05 |
| **ENSBTAG00000004780** | CDKL1 | -2.07 | 7.33E-05 | 64.48 | 98.48 | 64.00 | 58.12 | 149.28 | 383.80 | 319.76 | 347.72 |
| **ENSBTAG00000004115** | MYLIP | -2.07 | 9.37E-11 | 590.44 | 519.82 | 849.19 | 480.05 | 1658.07 | 2705.11 | 3499.07 | 2381.28 |
| **ENSBTAG00000008232** | DAB2IP | -2.07 | 1.09E-02 | 28.21 | 11.17 | 132.92 | 39.83 | 236.07 | 122.54 | 249.19 | 278.54 |

| | | | | | | | | | | | |
|---|---|---|---|---|---|---|---|---|---|---|---|
| **ENSBTAG00000030721** | UNCX | -2.07 | 7.65E-08 | 346.60 | 106.60 | 364.29 | 268.01 | 1047.75 | 1055.46 | 1096.07 | 1350.85 |
| **ENSBTAG00000009739** | CSMD1 | -2.07 | 2.83E-02 | 3.02 | 4.06 | 9.85 | 10.76 | 28.47 | 41.62 | 29.62 | 15.47 |
| **ENSBTAG00000020887** | CACNA1E | -2.06 | 2.11E-02 | 79.60 | 11.17 | 39.38 | 62.43 | 99.98 | 330.62 | 176.00 | 197.53 |
| **ENSBTAG00000017116** | RASD2 | -2.06 | 4.19E-03 | 46.35 | 70.05 | 60.30 | 49.51 | 268.71 | 150.28 | 234.38 | 288.56 |
| **ENSBTAG00000033015** | FAHD1 | -2.06 | 2.06E-03 | 30.23 | 32.49 | 83.69 | 26.91 | 58.32 | 201.15 | 136.79 | 323.15 |
| **ENSBTAG00000012653** | CAMK2B | -2.05 | 1.64E-03 | 11.08 | 19.29 | 6.15 | 9.69 | 52.77 | 33.52 | 54.02 | 52.80 |
| **ENSBTAG00000007746** | AHR | -2.05 | 1.30E-03 | 174.31 | 22.34 | 87.38 | 61.35 | 347.17 | 401.14 | 345.03 | 334.07 |
| **ENSBTAG00000054352** | | -2.04 | 8.84E-03 | 45.34 | 4.06 | 41.84 | 20.45 | 100.68 | 89.01 | 128.08 | 142.00 |
| **ENSBTAG00000020548** | AZIN2 | -2.04 | 1.63E-02 | 73.55 | 10.15 | 48.00 | 63.50 | 193.02 | 268.20 | 210.85 | 131.08 |
| **ENSBTAG00000018179** | PCDH8 | -2.04 | 3.49E-04 | 347.61 | 176.66 | 278.14 | 296.00 | 930.41 | 520.21 | 1894.17 | 1160.60 |
| **ENSBTAG00000019456** | SPRED2 | -2.04 | 6.79E-06 | 237.79 | 164.47 | 379.06 | 85.03 | 735.30 | 756.04 | 1086.49 | 969.44 |
| **ENSBTAG00000004705** | ILDR1 | -2.04 | 2.63E-04 | 42.32 | 119.80 | 100.92 | 46.28 | 486.73 | 190.75 | 340.67 | 247.59 |
| **ENSBTAG00000009950** | PAX3 | -2.03 | 3.05E-07 | 322.42 | 424.38 | 327.37 | 190.51 | 984.56 | 1145.63 | 1334.81 | 1717.69 |
| **ENSBTAG00000047582** | CSNK1E | -2.03 | 3.13E-13 | 557.19 | 258.89 | 364.29 | 259.40 | 1631.68 | 1399.95 | 1589.22 | 1268.01 |
| **ENSBTAG00000004974** | ZDHHC23 | -2.03 | 1.40E-06 | 53.40 | 42.64 | 43.07 | 88.26 | 318.00 | 168.78 | 261.38 | 180.23 |
| **ENSBTAG00000045526** | INSM1 | -2.03 | 1.13E-03 | 60.45 | 18.27 | 34.46 | 33.37 | 104.84 | 120.23 | 199.52 | 173.86 |
| **ENSBTAG00000013802** | DAB1 | -2.02 | 4.29E-02 | 172.29 | 20.31 | 27.08 | 41.98 | 361.75 | 176.87 | 223.92 | 301.30 |
| **ENSBTAG00000008004** | NCF2 | -2.02 | 6.67E-03 | 37.28 | 11.17 | 23.38 | 17.22 | 34.02 | 145.66 | 58.38 | 124.71 |
| **ENSBTAG00000049186** | | -2.02 | 1.42E-02 | 126.95 | 6.09 | 72.61 | 138.85 | 546.44 | 91.33 | 439.13 | 320.42 |
| **ENSBTAG00000037547** | OLIG1 | -2.02 | 1.02E-02 | 75.57 | 39.60 | 93.53 | 23.68 | 222.88 | 146.82 | 352.87 | 217.56 |
| **ENSBTAG00000009749** | USP2 | -2.01 | 1.79E-16 | 648.87 | 687.34 | 887.34 | 740.53 | 3114.78 | 2897.01 | 3127.91 | 2805.46 |
| **ENSBTAG00000008535** | SOCS7 | -2.01 | 1.81E-04 | 63.48 | 71.07 | 172.30 | 94.72 | 305.51 | 387.27 | 528.87 | 390.51 |
| **ENSBTAG00000040398** | KIAA1211 | -2.01 | 8.21E-05 | 64.48 | 71.07 | 142.76 | 97.95 | 420.77 | 441.60 | 259.64 | 387.78 |
| **ENSBTAG00000003512** | MYH7B | -2.00 | 1.86E-03 | 42.32 | 9.14 | 52.92 | 65.66 | 186.78 | 161.84 | 234.38 | 94.67 |
| **ENSBTAG00000006965** | NLRP8 | -1.99 | 1.11E-06 | 1438.81 | 579.72 | 1745.14 | 891.22 | 6637.83 | 3525.89 | 4045.37 | 4334.72 |
| **ENSBTAG00000046626** | CNTD2 | -1.98 | 1.70E-02 | 21.16 | 42.64 | 48.00 | 12.92 | 143.03 | 79.77 | 108.04 | 161.12 |
| **ENSBTAG00000052570** | | -1.98 | 1.81E-03 | 58.44 | 144.17 | 27.08 | 93.64 | 354.80 | 373.40 | 304.95 | 243.95 |
| **ENSBTAG00000012012** | CYB5A | -1.97 | 3.43E-06 | 361.72 | 309.66 | 377.83 | 180.83 | 638.79 | 1369.90 | 1270.33 | 1556.57 |
| **ENSBTAG00000039766** | FBRSL1 | -1.97 | 3.25E-08 | 358.69 | 268.03 | 380.29 | 456.37 | 2422.53 | 876.27 | 1229.38 | 1212.48 |
| **ENSBTAG00000012509** | DYRK1B | -1.97 | 1.26E-06 | 200.51 | 106.60 | 214.14 | 116.25 | 490.89 | 675.12 | 701.38 | 628.09 |
| **ENSBTAG00000017983** | PRELID3A | -1.96 | 9.17E-03 | 45.34 | 42.64 | 22.15 | 32.29 | 158.31 | 104.04 | 175.13 | 119.25 |
| **ENSBTAG00000019791** | GRHL3 | -1.96 | 1.67E-05 | 75.57 | 67.01 | 25.84 | 52.74 | 233.99 | 217.33 | 148.99 | 265.80 |
| **ENSBTAG00000009493** | BCL3 | -1.96 | 1.44E-02 | 119.90 | 88.33 | 72.61 | 34.44 | 265.93 | 396.52 | 406.89 | 160.21 |

| | | | | | | | | | | | |
|---|---|---|---|---|---|---|---|---|---|---|---|
| **ENSBTAG00000010192** | LYSMD4 | -1.96 | 1.53E-05 | 116.88 | 105.59 | 241.22 | 149.61 | 508.25 | 759.51 | 636.04 | 483.36 |
| **ENSBTAG00000032455** | HIST2H2BF | -1.96 | 3.17E-20 | 308.32 | 255.85 | 235.06 | 326.14 | 809.59 | 1151.41 | 1008.07 | 1420.03 |
| **ENSBTAG00000035995** | TNFAIP2 | -1.96 | 3.84E-03 | 21.16 | 18.27 | 52.92 | 32.29 | 213.85 | 36.99 | 103.68 | 127.44 |
| **ENSBTAG00000007841** | WTIP | -1.96 | 4.62E-04 | 178.34 | 135.03 | 103.38 | 41.98 | 203.44 | 358.37 | 617.74 | 608.06 |
| **ENSBTAG00000006209** | EXT1 | -1.96 | 5.78E-06 | 170.28 | 101.53 | 287.99 | 136.70 | 508.95 | 721.36 | 568.95 | 901.17 |
| **ENSBTAG00000009489** | CACNA2D2 | -1.96 | 1.62E-03 | 46.35 | 71.07 | 64.00 | 37.67 | 297.17 | 241.61 | 208.24 | 102.86 |
| **ENSBTAG00000004131** | RAP2C | -1.95 | 1.37E-02 | 143.07 | 192.90 | 48.00 | 27.99 | 229.13 | 538.71 | 345.03 | 486.09 |
| **ENSBTAG00000016455** | CACHD1 | -1.95 | 3.97E-03 | 140.05 | 128.94 | 379.06 | 119.48 | 672.11 | 775.70 | 787.64 | 730.95 |
| **ENSBTAG00000005581** | CCL25 | -1.94 | 4.71E-03 | 69.52 | 82.24 | 105.84 | 11.84 | 203.44 | 196.53 | 247.44 | 387.78 |
| **ENSBTAG00000021183** | MAP3K21 | -1.94 | 1.42E-03 | 210.58 | 36.55 | 162.45 | 130.24 | 434.65 | 359.53 | 670.89 | 600.78 |
| **ENSBTAG00000001509** | ELK3 | -1.93 | 1.41E-02 | 10.08 | 32.49 | 97.23 | 52.74 | 128.45 | 250.86 | 204.75 | 149.28 |
| **ENSBTAG00000003051** | FER | -1.93 | 2.05E-09 | 1438.81 | 964.51 | 1321.78 | 629.67 | 2389.90 | 5514.27 | 4755.46 | 3911.45 |
| **ENSBTAG00000016841** | ATP10B | -1.92 | 9.59E-03 | 17.13 | 21.32 | 9.85 | 34.44 | 58.32 | 100.57 | 83.64 | 72.82 |
| **ENSBTAG00000002037** | ATXN7L1 | -1.92 | 2.16E-03 | 94.71 | 79.19 | 54.15 | 31.21 | 243.02 | 277.45 | 230.02 | 234.85 |
| **ENSBTAG00000020348** | ASTL | -1.92 | 8.86E-03 | 34.26 | 14.21 | 57.84 | 37.67 | 162.47 | 80.92 | 146.38 | 153.84 |
| **ENSBTAG00000009918** | GSE1 | -1.92 | 1.17E-02 | 126.95 | 75.13 | 49.23 | 48.44 | 398.55 | 249.70 | 273.58 | 213.91 |
| **ENSBTAG00000038600** | LRRC15 | -1.92 | 3.82E-02 | 11.08 | 9.14 | 38.15 | 76.42 | 149.98 | 89.01 | 90.61 | 179.32 |
| **ENSBTAG00000038235** | NOD1 | -1.92 | 1.85E-02 | 113.86 | 23.35 | 50.46 | 9.69 | 187.47 | 161.84 | 170.77 | 224.84 |
| **ENSBTAG00000006054** | ACBD4 | -1.91 | 3.80E-03 | 23.17 | 21.32 | 41.84 | 33.37 | 128.45 | 99.42 | 144.63 | 76.46 |
| **ENSBTAG00000012370** | | -1.91 | 1.69E-02 | 163.23 | 67.01 | 76.30 | 90.41 | 385.36 | 397.67 | 549.78 | 161.12 |
| **ENSBTAG00000027134** | DYNC1I1 | -1.91 | 3.63E-03 | 95.72 | 15.23 | 105.84 | 85.03 | 161.78 | 378.02 | 330.22 | 263.98 |
| **ENSBTAG00000017824** | IRF8 | -1.91 | 5.61E-06 | 349.63 | 334.02 | 748.27 | 377.80 | 1421.99 | 1935.19 | 1938.60 | 1494.67 |
| **ENSBTAG00000017733** | CA2 | -1.91 | 3.72E-02 | 193.45 | 100.51 | 46.77 | 49.51 | 224.96 | 187.28 | 211.72 | 841.09 |
| **ENSBTAG00000016836** | PDK1 | -1.90 | 6.85E-10 | 233.76 | 282.25 | 568.59 | 283.08 | 973.46 | 1443.88 | 1296.47 | 1400.00 |
| **ENSBTAG00000003669** | BNC2 | -1.90 | 2.74E-07 | 282.12 | 143.15 | 172.30 | 116.25 | 717.25 | 455.48 | 877.38 | 620.81 |
| **ENSBTAG00000024426** | PPP1R9A | -1.90 | 2.25E-03 | 18.14 | 11.17 | 14.77 | 17.22 | 41.66 | 69.36 | 66.22 | 51.89 |
| **ENSBTAG00000005888** | MDGA1 | -1.89 | 2.16E-02 | 83.63 | 27.41 | 96.00 | 10.76 | 186.78 | 197.68 | 137.66 | 286.74 |
| **ENSBTAG00000050206** | | -1.89 | 1.71E-02 | 24.18 | 36.55 | 24.61 | 9.69 | 43.74 | 100.57 | 115.01 | 94.67 |
| **ENSBTAG00000031194** | PHLDA2 | -1.89 | 2.32E-02 | 2554.19 | 975.67 | 2382.65 | 1525.19 | 2105.91 | 3587.16 | 16200.63 | 5739.28 |
| **ENSBTAG00000046768** | IGFBP1 | -1.89 | 4.99E-02 | 3.02 | 48.73 | 18.46 | 17.22 | 59.02 | 71.67 | 93.23 | 101.04 |
| **ENSBTAG00000011234** | FOXO3 | -1.89 | 6.12E-05 | 136.02 | 150.26 | 107.07 | 104.41 | 449.23 | 361.84 | 538.45 | 495.19 |
| **ENSBTAG00000012681** | CORO2B | -1.89 | 2.75E-03 | 26.20 | 50.76 | 19.69 | 31.21 | 143.73 | 87.86 | 191.68 | 50.98 |
| **ENSBTAG00000020458** | STX6 | -1.89 | 3.72E-03 | 54.41 | 152.29 | 93.53 | 45.21 | 337.45 | 304.04 | 337.19 | 297.66 |

| | | | | | | | | | | | |
|---|---|---|---|---|---|---|---|---|---|---|---|
| **ENSBTAG00000008940** | NPTX1 | -1.89 | 1.18E-05 | 212.60 | 209.15 | 159.99 | 237.87 | 1040.11 | 297.10 | 900.91 | 788.30 |
| **ENSBTAG00000015979** | SEZ6 | -1.88 | 3.01E-04 | 140.05 | 34.52 | 180.91 | 72.12 | 361.05 | 409.23 | 415.60 | 388.69 |
| **ENSBTAG00000002224** | UHRF1 | -1.88 | 1.03E-09 | 1634.28 | 907.65 | 1358.70 | 837.40 | 4277.79 | 3968.65 | 5680.76 | 3545.52 |
| **ENSBTAG00000053238** | | -1.88 | 3.05E-02 | 45.34 | 37.56 | 66.46 | 6.46 | 154.84 | 120.23 | 142.02 | 155.66 |
| **ENSBTAG00000003880** | EMILIN2 | -1.88 | 1.25E-02 | 90.68 | 36.55 | 38.15 | 34.44 | 79.85 | 260.11 | 276.20 | 119.25 |
| **ENSBTAG00000002081** | BMPR1B | -1.87 | 3.28E-03 | 66.50 | 37.56 | 132.92 | 121.63 | 155.53 | 320.22 | 426.06 | 405.98 |
| **ENSBTAG00000015498** | ELOVL4 | -1.87 | 2.36E-03 | 287.16 | 108.63 | 369.21 | 59.20 | 573.52 | 806.91 | 796.35 | 829.26 |
| **ENSBTAG00000009587** | NRG3 | -1.86 | 1.81E-02 | 19.14 | 20.31 | 23.38 | 23.68 | 31.25 | 78.61 | 92.36 | 112.87 |
| **ENSBTAG00000004300** | LRRC8E | -1.86 | 1.36E-03 | 59.45 | 74.11 | 46.77 | 109.79 | 277.73 | 215.02 | 357.23 | 205.72 |
| **ENSBTAG00000004593** | TOP2B | -1.86 | 3.97E-08 | 1205.05 | 691.40 | 1175.32 | 961.18 | 2676.66 | 4657.65 | 3088.70 | 4232.77 |
| **ENSBTAG00000039522** | CYCT | -1.86 | 3.11E-04 | 467.51 | 391.89 | 146.45 | 354.12 | 845.00 | 545.65 | 1854.09 | 1696.75 |
| **ENSBTAG00000027182** | NR3C2 | -1.86 | 1.93E-11 | 479.60 | 469.05 | 414.75 | 386.41 | 1919.83 | 1152.56 | 1541.30 | 1737.71 |
| **ENSBTAG00000008353** | CDKN1A | -1.86 | 1.06E-03 | 155.17 | 77.16 | 120.61 | 121.63 | 254.13 | 247.39 | 690.06 | 526.14 |
| **ENSBTAG00000007494** | SMARCA2 | -1.85 | 2.25E-06 | 324.44 | 110.66 | 422.13 | 246.48 | 740.16 | 941.01 | 1092.59 | 1197.92 |
| **ENSBTAG00000011044** | TACC3 | -1.84 | 1.08E-23 | 5569.84 | 4202.20 | 5416.34 | 3752.17 | 18095.02 | 14713.95 | 19635.23 | 15537.47 |
| **ENSBTAG00000048628** | HHAT | -1.84 | 1.94E-02 | 47.36 | 54.82 | 36.92 | 32.29 | 97.90 | 223.11 | 152.47 | 141.09 |
| **ENSBTAG00000021339** | SCN8A | -1.84 | 1.11E-04 | 175.32 | 164.47 | 232.60 | 291.69 | 785.99 | 1063.55 | 659.56 | 583.49 |
| **ENSBTAG00000005946** | USP13 | -1.84 | 3.26E-03 | 130.98 | 141.12 | 137.84 | 68.89 | 352.72 | 388.43 | 569.82 | 402.34 |
| **ENSBTAG00000000495** | HAVCR2 | -1.84 | 4.93E-02 | 26.20 | 29.44 | 20.92 | 11.84 | 58.32 | 6.94 | 110.65 | 140.18 |
| **ENSBTAG00000010241** | UNC5D | -1.84 | 9.69E-04 | 25.19 | 32.49 | 17.23 | 32.29 | 80.54 | 105.20 | 72.32 | 126.53 |
| **ENSBTAG00000026842** | ZRANB3 | -1.84 | 2.28E-02 | 151.14 | 58.89 | 64.00 | 119.48 | 269.40 | 441.60 | 334.57 | 360.47 |
| **ENSBTAG00000046348** | ELAVL2 | -1.84 | 8.26E-05 | 483.63 | 296.46 | 454.13 | 470.37 | 879.03 | 1891.27 | 1844.51 | 1468.27 |
| **ENSBTAG00000037389** | TRIM44 | -1.83 | 1.03E-02 | 34.26 | 32.49 | 16.00 | 51.66 | 96.51 | 99.42 | 111.52 | 171.13 |
| **ENSBTAG00000017268** | PROCA1 | -1.82 | 1.80E-06 | 422.17 | 320.83 | 500.90 | 196.97 | 1027.61 | 1227.70 | 1586.61 | 1255.27 |
| **ENSBTAG00000008936** | ABCC2 | -1.82 | 4.96E-02 | 15.11 | 58.89 | 46.77 | 12.92 | 90.26 | 115.60 | 132.43 | 133.81 |
| **ENSBTAG00000002888** | TMTC2 | -1.82 | 1.56E-04 | 121.92 | 67.01 | 273.22 | 333.67 | 649.90 | 802.29 | 731.88 | 621.72 |
| **ENSBTAG00000018161** | TBX18 | -1.82 | 4.20E-04 | 71.54 | 35.53 | 72.61 | 62.43 | 188.16 | 151.44 | 304.08 | 207.54 |
| **ENSBTAG00000004462** | NPAS3 | -1.82 | 4.28E-02 | 7.05 | 7.11 | 12.31 | 8.61 | 24.30 | 42.77 | 26.14 | 30.04 |
| **ENSBTAG00000026085** | ZP4 | -1.81 | 1.01E-05 | 288.16 | 360.42 | 412.29 | 290.62 | 1199.81 | 1174.53 | 865.18 | 1512.88 |
| **ENSBTAG00000016834** | LHFPL5 | -1.81 | 2.77E-02 | 34.26 | 27.41 | 24.61 | 21.53 | 124.29 | 141.04 | 33.11 | 81.01 |
| **ENSBTAG00000017285** | FGF19 | -1.81 | 1.10E-03 | 39.30 | 136.05 | 124.30 | 48.44 | 427.02 | 272.82 | 326.73 | 193.89 |
| **ENSBTAG00000048478** | | -1.81 | 4.29E-02 | 120.91 | 17.26 | 125.53 | 39.83 | 159.70 | 36.99 | 642.14 | 222.11 |
| **ENSBTAG00000002545** | | -1.81 | 4.85E-03 | 247.86 | 145.18 | 201.84 | 103.33 | 538.11 | 199.99 | 418.22 | 1286.22 |

| | | | | | | | | | | | |
|---|---|---|---|---|---|---|---|---|---|---|---|
| **ENSBTAG00000013588** | ZNF532 | -1.81 | 9.01E-04 | 127.96 | 64.98 | 178.45 | 107.64 | 325.64 | 473.97 | 528.87 | 344.99 |
| **ENSBTAG00000015981** | ETV1 | -1.80 | 1.34E-02 | 29.22 | 50.76 | 82.46 | 48.44 | 106.93 | 176.87 | 230.89 | 219.38 |
| **ENSBTAG00000022917** | FANK1 | -1.80 | 7.41E-03 | 156.17 | 153.31 | 203.07 | 11.84 | 388.13 | 569.92 | 360.71 | 508.84 |
| **ENSBTAG00000021844** | COCH | -1.80 | 8.33E-03 | 25.19 | 45.69 | 33.23 | 44.13 | 171.50 | 60.11 | 117.62 | 166.58 |
| **ENSBTAG00000004313** | HOXD13 | -1.80 | 4.99E-02 | 31.23 | 68.02 | 28.31 | 32.29 | 193.72 | 165.31 | 55.76 | 142.00 |
| **ENSBTAG00000002103** | PLCG2 | -1.80 | 5.85E-03 | 41.31 | 95.44 | 124.30 | 161.45 | 670.73 | 342.18 | 136.79 | 317.69 |
| **ENSBTAG00000047594** | CCNO | -1.80 | 7.60E-04 | 67.51 | 33.50 | 55.38 | 43.05 | 203.44 | 131.79 | 189.94 | 167.49 |
| **ENSBTAG00000001414** | KCTD12 | -1.80 | 7.63E-03 | 73.55 | 50.76 | 23.38 | 50.59 | 247.18 | 105.20 | 179.48 | 158.39 |
| **ENSBTAG00000010581** | MAGI1 | -1.80 | 1.59E-03 | 162.22 | 77.16 | 227.68 | 86.11 | 270.79 | 507.50 | 474.85 | 667.23 |
| **ENSBTAG00000013247** | NLRP5 | -1.80 | 5.57E-03 | 78.59 | 43.66 | 86.15 | 40.90 | 193.02 | 293.63 | 192.55 | 185.70 |
| **ENSBTAG00000002361** | TMCC2 | -1.79 | 1.02E-03 | 78.59 | 64.98 | 68.92 | 45.21 | 237.46 | 121.38 | 228.28 | 303.12 |
| **ENSBTAG00000016259** | KIAA1958 | -1.79 | 1.00E-04 | 102.77 | 98.48 | 51.69 | 39.83 | 218.02 | 334.09 | 194.30 | 268.53 |
| **ENSBTAG00000011639** | STK11 | -1.79 | 1.36E-05 | 236.78 | 170.57 | 344.60 | 311.07 | 1133.85 | 583.80 | 1013.30 | 933.94 |
| **ENSBTAG00000018394** | SDR42E1 | -1.78 | 6.59E-07 | 172.29 | 198.99 | 162.45 | 150.69 | 422.85 | 751.42 | 488.79 | 692.72 |
| **ENSBTAG00000044111** | EPHA6 | -1.78 | 4.24E-02 | 19.14 | 16.24 | 41.84 | 12.92 | 39.58 | 110.98 | 79.29 | 79.19 |
| **ENSBTAG00000006194** | FOSL1 | -1.77 | 1.90E-02 | 25.19 | 71.07 | 56.61 | 52.74 | 147.20 | 55.49 | 287.52 | 212.09 |
| **ENSBTAG00000026637** | | -1.77 | 4.29E-03 | 27.20 | 26.40 | 27.08 | 11.84 | 48.60 | 94.79 | 79.29 | 92.85 |
| **ENSBTAG00000008105** | RBM38 | -1.77 | 1.33E-07 | 829.23 | 413.21 | 508.28 | 483.28 | 1521.98 | 1349.09 | 2788.11 | 1943.44 |
| **ENSBTAG00000008814** | ADGRA2 | -1.76 | 4.91E-02 | 6.05 | 15.23 | 8.61 | 3.23 | 22.22 | 30.06 | 30.49 | 30.04 |
| **ENSBTAG00000021505** | USP36 | -1.76 | 5.49E-10 | 1141.58 | 699.52 | 999.33 | 1177.53 | 4377.08 | 2335.18 | 4000.06 | 2869.18 |
| **ENSBTAG00000044195** | SDK2 | -1.75 | 8.70E-05 | 53.40 | 60.92 | 54.15 | 31.21 | 158.31 | 187.28 | 223.05 | 105.59 |
| **ENSBTAG00000003719** | TDRD7 | -1.75 | 1.75E-02 | 28.21 | 89.34 | 52.92 | 77.50 | 130.53 | 197.68 | 162.06 | 344.99 |
| **ENSBTAG00000002202** | CRAMP1 | -1.75 | 3.96E-04 | 147.11 | 93.40 | 136.61 | 55.97 | 428.40 | 252.01 | 417.34 | 355.01 |
| **ENSBTAG00000025071** | TENM2 | -1.75 | 1.70E-02 | 44.33 | 40.61 | 25.84 | 9.69 | 106.93 | 68.21 | 92.36 | 137.45 |
| **ENSBTAG00000006667** | EPB41 | -1.74 | 4.68E-07 | 352.65 | 240.62 | 511.97 | 386.41 | 1347.01 | 1297.07 | 1381.85 | 969.44 |
| **ENSBTAG00000004036** | GJC1 | -1.74 | 1.57E-09 | 292.20 | 485.30 | 600.58 | 391.79 | 1326.18 | 1262.39 | 1633.66 | 1665.80 |
| **ENSBTAG00000048486** | IQANK1 | -1.73 | 3.99E-02 | 16.12 | 13.20 | 25.84 | 6.46 | 23.61 | 57.80 | 56.63 | 66.45 |
| **ENSBTAG00000006438** | ACTL8 | -1.73 | 4.23E-03 | 75.57 | 86.30 | 43.07 | 64.58 | 231.91 | 158.38 | 245.70 | 260.34 |
| **ENSBTAG00000017976** | ABL1 | -1.73 | 3.26E-06 | 417.13 | 234.53 | 332.29 | 222.81 | 1009.56 | 941.01 | 1228.51 | 821.07 |
| **ENSBTAG00000007584** | INPP1 | -1.73 | 3.87E-04 | 177.33 | 79.19 | 127.99 | 151.77 | 351.33 | 576.86 | 312.79 | 534.33 |
| **ENSBTAG00000006870** | RASGEF1A | -1.73 | 2.96E-05 | 164.23 | 102.54 | 185.84 | 220.65 | 357.58 | 410.39 | 656.95 | 801.04 |
| **ENSBTAG00000034140** | MEGF11 | -1.73 | 4.69E-02 | 26.20 | 4.06 | 34.46 | 25.83 | 35.41 | 91.33 | 55.76 | 116.52 |
| **ENSBTAG00000012106** | ADCK1 | -1.72 | 3.44E-02 | 40.30 | 56.86 | 111.99 | 40.90 | 164.56 | 338.72 | 181.23 | 140.18 |

| | | | | | | | | | | | |
|---|---|---|---|---|---|---|---|---|---|---|---|
| **ENSBTAG00000019517** | ELN | -1.72 | 9.07E-05 | 115.87 | 86.30 | 182.14 | 146.38 | 305.51 | 450.85 | 382.49 | 608.97 |
| **ENSBTAG00000026977** | PTPRQ | -1.72 | 6.45E-05 | 253.91 | 246.71 | 290.45 | 156.07 | 633.23 | 1095.92 | 689.18 | 700.00 |
| **ENSBTAG00000044126** | SNTB1 | -1.72 | 4.35E-03 | 75.57 | 15.23 | 92.30 | 32.29 | 153.45 | 130.63 | 193.42 | 227.57 |
| **ENSBTAG00000000923** | HOXC13 | -1.71 | 3.92E-02 | 501.77 | 318.79 | 210.45 | 254.02 | 848.48 | 399.99 | 1805.30 | 1165.15 |
| **ENSBTAG00000049111** | | -1.71 | 6.67E-06 | 116.88 | 53.81 | 91.07 | 83.96 | 174.97 | 331.78 | 320.63 | 304.94 |
| **ENSBTAG00000032021** | RALB | -1.71 | 1.57E-12 | 2200.53 | 1914.80 | 2590.64 | 1734.01 | 5516.48 | 9249.40 | 6122.50 | 6678.68 |
| **ENSBTAG00000053150** | | -1.71 | 3.21E-03 | 32.24 | 48.73 | 30.77 | 20.45 | 49.30 | 132.94 | 112.40 | 138.36 |
| **ENSBTAG00000018084** | CASTOR1 | -1.71 | 1.74E-08 | 697.24 | 342.15 | 575.97 | 278.78 | 1112.32 | 1491.28 | 2000.47 | 1572.95 |
| **ENSBTAG00000047902** | ULBP21 | -1.70 | 1.96E-02 | 15.11 | 60.92 | 107.07 | 67.81 | 153.45 | 89.01 | 222.18 | 350.46 |
| **ENSBTAG00000014605** | ETV6 | -1.70 | 6.09E-03 | 112.85 | 187.82 | 238.76 | 95.80 | 175.67 | 624.26 | 580.27 | 685.44 |
| **ENSBTAG00000004862** | TUB | -1.70 | 1.10E-03 | 72.54 | 62.95 | 48.00 | 54.89 | 156.23 | 190.75 | 311.05 | 115.60 |
| **ENSBTAG00000002471** | CHL1 | -1.69 | 5.34E-03 | 272.04 | 167.52 | 204.30 | 170.06 | 305.51 | 1127.13 | 522.77 | 672.69 |
| **ENSBTAG00000002417** | RAB30 | -1.69 | 1.68E-04 | 249.88 | 414.23 | 236.30 | 289.54 | 783.21 | 1295.91 | 882.61 | 876.59 |
| **ENSBTAG00000012389** | | -1.69 | 1.95E-03 | 609.58 | 268.03 | 433.21 | 200.20 | 1852.48 | 543.33 | 1132.67 | 1330.82 |
| **ENSBTAG00000044105** | FOXO1 | -1.68 | 4.21E-08 | 636.78 | 510.68 | 695.35 | 540.33 | 2005.24 | 1871.61 | 1802.68 | 1979.85 |
| **ENSBTAG00000055035** | | -1.68 | 3.52E-02 | 341.57 | 43.66 | 131.69 | 277.70 | 1051.92 | 226.58 | 872.15 | 402.34 |
| **ENSBTAG00000001874** | DOCK3 | -1.68 | 3.49E-02 | 4.03 | 54.82 | 32.00 | 25.83 | 136.78 | 79.77 | 92.36 | 64.63 |
| **ENSBTAG00000013726** | RNPEP | -1.68 | 4.40E-04 | 136.02 | 182.75 | 255.99 | 206.66 | 347.86 | 569.92 | 888.71 | 692.72 |
| **ENSBTAG00000023765** | C18H16orf46 | -1.68 | 3.76E-03 | 56.42 | 97.47 | 86.15 | 55.97 | 303.42 | 154.91 | 215.21 | 271.26 |
| **ENSBTAG00000046509** | TENT5C | -1.68 | 2.06E-08 | 476.58 | 277.17 | 438.13 | 355.20 | 1184.53 | 1019.62 | 1157.06 | 1577.50 |
| **ENSBTAG00000053488** | | -1.67 | 3.92E-02 | 19.14 | 6.09 | 4.92 | 16.15 | 24.30 | 27.74 | 41.82 | 54.62 |
| **ENSBTAG00000005412** | NEDD4L | -1.67 | 1.07E-04 | 296.23 | 193.92 | 168.61 | 168.99 | 751.27 | 482.06 | 845.14 | 558.00 |
| **ENSBTAG00000023730** | TUBB3 | -1.67 | 1.50E-06 | 432.25 | 296.46 | 503.36 | 529.57 | 1412.27 | 891.30 | 1976.94 | 1321.72 |
| **ENSBTAG00000004791** | VPS26C | -1.67 | 1.45E-03 | 282.12 | 272.09 | 104.61 | 115.17 | 492.98 | 567.61 | 816.39 | 588.04 |
| **ENSBTAG00000007578** | SHTN1 | -1.67 | 1.95E-02 | 96.73 | 145.18 | 216.60 | 110.86 | 261.07 | 610.38 | 472.24 | 463.33 |
| **ENSBTAG00000001514** | ASB11 | -1.66 | 1.73E-06 | 221.67 | 288.34 | 376.60 | 367.04 | 599.90 | 1394.17 | 1043.80 | 929.39 |
| **ENSBTAG00000017263** | MXI1 | -1.66 | 3.03E-07 | 578.34 | 451.80 | 439.36 | 395.02 | 946.38 | 1486.66 | 1643.24 | 1821.46 |
| **ENSBTAG00000010371** | CHAC1 | -1.66 | 4.79E-03 | 208.57 | 162.44 | 317.52 | 156.07 | 481.17 | 574.55 | 839.92 | 763.72 |
| **ENSBTAG00000018773** | RND1 | -1.65 | 3.60E-03 | 766.76 | 120.82 | 1055.95 | 351.97 | 1384.50 | 1371.05 | 2088.47 | 2369.44 |
| **ENSBTAG00000008005** | DRC7 | -1.65 | 7.11E-03 | 47.36 | 48.73 | 139.07 | 76.42 | 236.07 | 199.99 | 221.31 | 318.60 |
| **ENSBTAG00000024204** | | -1.65 | 3.04E-05 | 94.71 | 51.78 | 108.30 | 69.96 | 236.77 | 198.84 | 319.76 | 262.16 |
| **ENSBTAG00000015535** | NEK8 | -1.65 | 4.57E-02 | 45.34 | 31.47 | 17.23 | 13.99 | 61.10 | 93.64 | 100.20 | 85.57 |
| **ENSBTAG00000011209** | SYT11 | -1.65 | 1.60E-02 | 425.19 | 206.10 | 332.29 | 97.95 | 818.62 | 267.04 | 811.16 | 1428.22 |

| | | | | | | | | | | | |
|---|---|---|---|---|---|---|---|---|---|---|---|
| **ENSBTAG00000015313** | CEACAM19 | -1.65 | 9.59E-03 | 18.14 | 30.46 | 28.31 | 24.76 | 72.91 | 38.15 | 108.04 | 98.31 |
| **ENSBTAG00000019138** | KIF3C | -1.64 | 1.55E-03 | 298.24 | 248.74 | 518.13 | 345.51 | 1278.96 | 1046.21 | 957.54 | 1107.80 |
| **ENSBTAG00000004658** | WEE2 | -1.64 | 8.32E-03 | 235.77 | 51.78 | 445.52 | 113.02 | 740.16 | 525.99 | 633.42 | 730.04 |
| **ENSBTAG00000011540** | SPG21 | -1.63 | 2.46E-07 | 1458.96 | 944.20 | 1158.09 | 795.43 | 1907.33 | 5277.28 | 3396.26 | 2948.38 |
| **ENSBTAG00000011000** | HDAC10 | -1.63 | 3.04E-04 | 97.73 | 86.30 | 62.77 | 85.03 | 264.54 | 347.97 | 198.65 | 220.29 |
| **ENSBTAG00000018909** | CREB5 | -1.63 | 1.06E-02 | 41.31 | 118.79 | 108.30 | 154.99 | 194.41 | 112.13 | 548.91 | 452.41 |
| **ENSBTAG00000021747** | | -1.63 | 1.02E-08 | 409.07 | 249.76 | 409.83 | 487.59 | 793.62 | 1310.94 | 1427.16 | 1276.20 |
| **ENSBTAG00000000260** | ZNRF2 | -1.63 | 8.77E-09 | 698.25 | 650.79 | 772.88 | 838.48 | 1375.47 | 2891.23 | 2855.19 | 2024.45 |
| **ENSBTAG00000020796** | UBE2D1 | -1.62 | 2.22E-02 | 174.31 | 253.82 | 183.38 | 166.83 | 209.69 | 879.74 | 566.33 | 730.04 |
| **ENSBTAG00000005275** | PKIG | -1.62 | 6.59E-03 | 115.87 | 45.69 | 116.92 | 77.50 | 119.43 | 361.84 | 295.36 | 314.04 |
| **ENSBTAG00000018577** | SLC7A1 | -1.61 | 4.60E-02 | 34.26 | 45.69 | 40.61 | 16.15 | 127.76 | 113.29 | 99.33 | 76.46 |
| **ENSBTAG00000023929** | FOSL2 | -1.61 | 3.33E-03 | 104.79 | 47.72 | 77.53 | 58.12 | 213.85 | 264.73 | 186.45 | 213.00 |
| **ENSBTAG00000050550** | FAM110B | -1.60 | 1.02E-03 | 127.96 | 110.66 | 212.91 | 94.72 | 311.06 | 431.20 | 413.86 | 495.19 |
| **ENSBTAG00000033089** | HACD1 | -1.60 | 3.14E-02 | 52.39 | 82.24 | 46.77 | 32.29 | 110.40 | 206.93 | 105.43 | 224.84 |
| **ENSBTAG00000020520** | RASD1 | -1.60 | 4.60E-02 | 142.07 | 51.78 | 72.61 | 35.52 | 163.17 | 158.38 | 417.34 | 173.86 |
| **ENSBTAG00000014543** | PLXNA4 | -1.59 | 5.52E-03 | 67.51 | 112.69 | 61.54 | 96.87 | 443.68 | 249.70 | 208.24 | 118.34 |
| **ENSBTAG00000003865** | CASR | -1.59 | 4.12E-05 | 215.62 | 181.73 | 285.52 | 107.64 | 765.16 | 373.40 | 692.67 | 546.16 |
| **ENSBTAG00000012262** | FBXO10 | -1.59 | 3.48E-03 | 66.50 | 120.82 | 48.00 | 52.74 | 374.25 | 102.89 | 190.81 | 198.44 |
| **ENSBTAG00000014267** | ZCCHC14 | -1.59 | 2.13E-07 | 467.51 | 394.94 | 520.59 | 396.10 | 1037.33 | 1195.34 | 1836.66 | 1280.76 |
| **ENSBTAG00000017731** | ZFAT | -1.59 | 3.11E-03 | 62.47 | 29.44 | 35.69 | 32.29 | 147.89 | 113.29 | 121.11 | 98.31 |
| **ENSBTAG00000012382** | KCTD9 | -1.59 | 3.39E-09 | 449.38 | 679.22 | 559.97 | 597.38 | 1131.07 | 2053.11 | 1712.94 | 1968.01 |
| **ENSBTAG00000005212** | C1H3orf70 | -1.59 | 2.52E-02 | 18.14 | 22.34 | 45.54 | 41.98 | 117.34 | 117.92 | 77.54 | 70.09 |
| **ENSBTAG00000016658** | ABHD4 | -1.58 | 4.56E-04 | 217.63 | 173.61 | 313.83 | 216.35 | 739.47 | 517.90 | 664.79 | 838.36 |
| **ENSBTAG00000017527** | CRYBG1 | -1.58 | 3.44E-03 | 39.30 | 111.68 | 131.69 | 141.00 | 430.49 | 310.97 | 310.18 | 211.18 |
| **ENSBTAG00000048850** | | -1.58 | 6.17E-03 | 35.26 | 95.44 | 48.00 | 36.60 | 100.68 | 204.62 | 196.04 | 142.00 |
| **ENSBTAG00000024957** | SNCA | -1.57 | 1.13E-04 | 282.12 | 178.69 | 322.44 | 204.51 | 495.75 | 808.07 | 677.86 | 957.61 |
| **ENSBTAG00000003836** | ADAM19 | -1.57 | 1.26E-06 | 670.03 | 607.13 | 659.66 | 578.00 | 1243.55 | 1930.57 | 2171.24 | 2130.95 |
| **ENSBTAG00000033890** | PRAME | -1.57 | 4.52E-04 | 1937.56 | 1083.29 | 951.34 | 683.48 | 4813.81 | 1284.35 | 2991.99 | 4742.53 |
| **ENSBTAG00000001294** | PPP1R15A | -1.57 | 9.55E-06 | 445.35 | 337.07 | 493.51 | 257.25 | 1562.94 | 758.36 | 1301.70 | 923.93 |
| **ENSBTAG00000019497** | ILDR2 | -1.57 | 8.21E-03 | 91.69 | 61.93 | 59.07 | 65.66 | 112.48 | 223.11 | 156.83 | 334.07 |
| **ENSBTAG00000030483** | KLK7 | -1.57 | 3.06E-05 | 199.50 | 282.25 | 291.68 | 322.91 | 1152.59 | 908.64 | 585.50 | 599.87 |
| **ENSBTAG00000018272** | RERE | -1.56 | 2.19E-09 | 286.15 | 208.13 | 173.53 | 163.61 | 679.75 | 530.62 | 740.59 | 509.75 |
| **ENSBTAG00000014612** | DOCK2 | -1.56 | 4.12E-02 | 37.28 | 11.17 | 81.23 | 29.06 | 124.29 | 117.92 | 135.92 | 88.30 |

| | | | | | | | | | | | |
|---|---|---|---|---|---|---|---|---|---|---|---|
| **ENSBTAG00000043961** | MPDZ | -1.56 | 7.54E-03 | 52.39 | 37.56 | 73.84 | 58.12 | 217.33 | 147.97 | 88.87 | 197.53 |
| **ENSBTAG00000015355** | PHKA2 | -1.56 | 1.20E-03 | 353.66 | 229.45 | 598.12 | 218.50 | 1344.92 | 670.50 | 1364.43 | 735.50 |
| **ENSBTAG00000012010** | PANX1 | -1.56 | 1.49E-02 | 128.97 | 108.63 | 61.54 | 79.65 | 356.89 | 298.26 | 250.93 | 209.36 |
| **ENSBTAG00000014619** | WASF3 | -1.55 | 8.70E-05 | 215.62 | 103.56 | 375.37 | 350.89 | 654.76 | 752.58 | 730.14 | 914.83 |
| **ENSBTAG00000008661** | C15H11orf52 | -1.55 | 2.35E-02 | 20.15 | 115.74 | 48.00 | 30.14 | 119.43 | 141.04 | 217.82 | 147.46 |
| **ENSBTAG00000040001** | MARVELD2 | -1.55 | 8.34E-03 | 471.54 | 155.34 | 198.14 | 52.74 | 665.87 | 647.38 | 506.22 | 745.51 |
| **ENSBTAG00000004383** | FNBP1L | -1.54 | 7.98E-05 | 464.49 | 603.07 | 292.91 | 473.60 | 638.09 | 1781.44 | 1443.72 | 1484.66 |
| **ENSBTAG00000019707** | GATA2 | -1.54 | 9.05E-10 | 321.41 | 283.26 | 436.90 | 368.11 | 838.75 | 969.91 | 1177.10 | 1115.09 |
| **ENSBTAG00000013994** | BSX | -1.54 | 2.69E-02 | 17.13 | 37.56 | 38.15 | 29.06 | 47.21 | 119.07 | 100.20 | 88.30 |
| **ENSBTAG00000017839** | TIAM1 | -1.54 | 9.04E-03 | 137.03 | 125.89 | 83.69 | 77.50 | 585.32 | 293.63 | 131.56 | 223.02 |
| **ENSBTAG00000030898** | MS4A13 | -1.54 | 6.15E-03 | 134.01 | 262.95 | 215.37 | 79.65 | 249.27 | 324.84 | 733.62 | 702.73 |
| **ENSBTAG00000000436** | TNFAIP3 | -1.53 | 4.76E-02 | 204.54 | 59.90 | 67.69 | 54.89 | 293.01 | 124.85 | 175.13 | 526.14 |
| **ENSBTAG00000000656** | NFATC1 | -1.53 | 2.39E-03 | 104.79 | 184.78 | 119.38 | 55.97 | 483.95 | 211.55 | 351.13 | 293.11 |
| **ENSBTAG00000023666** | | -1.53 | 1.14E-03 | 446.35 | 248.74 | 308.91 | 145.31 | 665.17 | 950.26 | 653.46 | 1043.17 |
| **ENSBTAG00000007139** | WSB2 | -1.52 | 1.07E-04 | 602.53 | 419.31 | 696.58 | 428.39 | 1478.24 | 1907.45 | 1258.13 | 1530.17 |
| **ENSBTAG00000051378** | | -1.52 | 2.66E-04 | 74.56 | 143.15 | 82.46 | 142.08 | 256.21 | 506.34 | 280.55 | 230.30 |
| **ENSBTAG00000007071** | RAI14 | -1.52 | 2.06E-03 | 418.14 | 328.95 | 339.67 | 226.03 | 1181.76 | 863.55 | 762.37 | 953.06 |
| **ENSBTAG00000019164** | RHOBTB1 | -1.52 | 4.65E-02 | 56.42 | 13.20 | 34.46 | 38.75 | 47.91 | 95.95 | 144.63 | 121.07 |
| **ENSBTAG00000023283** | JAML | -1.52 | 4.52E-04 | 119.90 | 100.51 | 75.07 | 151.77 | 257.60 | 292.48 | 325.86 | 405.07 |
| **ENSBTAG00000021654** | GAS2 | -1.51 | 8.77E-03 | 116.88 | 79.19 | 96.00 | 136.70 | 166.64 | 161.84 | 279.68 | 616.26 |
| **ENSBTAG00000001597** | PITPNM2 | -1.51 | 2.03E-02 | 31.23 | 52.79 | 18.46 | 37.67 | 115.95 | 82.08 | 97.58 | 104.68 |
| **ENSBTAG00000024482** | | -1.51 | 3.63E-04 | 118.89 | 175.64 | 109.53 | 67.81 | 214.55 | 315.60 | 298.85 | 517.04 |
| **ENSBTAG00000008842** | JPH1 | -1.51 | 4.50E-03 | 325.44 | 110.66 | 83.69 | 130.24 | 724.88 | 399.99 | 356.35 | 371.39 |
| **ENSBTAG00000020797** | CABLES2 | -1.50 | 7.46E-04 | 141.06 | 101.53 | 145.22 | 139.93 | 422.85 | 368.77 | 455.68 | 249.42 |
| **ENSBTAG00000055205** | | -1.50 | 9.54E-05 | 96.73 | 50.76 | 83.69 | 97.95 | 174.97 | 189.59 | 301.46 | 266.71 |
| **ENSBTAG00000011041** | ZFYVE1 | -1.50 | 1.97E-07 | 577.34 | 510.68 | 689.20 | 478.98 | 1664.32 | 1268.17 | 1615.36 | 1833.29 |
| **ENSBTAG00000017486** | ADARB1 | -1.50 | 1.58E-02 | 52.39 | 36.55 | 27.08 | 34.44 | 115.26 | 63.58 | 110.65 | 135.63 |
| **ENSBTAG00000000015** | FOXRED2 | -1.50 | 2.57E-04 | 125.95 | 300.52 | 126.76 | 172.22 | 647.12 | 332.94 | 439.13 | 628.09 |
| **ENSBTAG00000053699** | | -1.49 | 1.06E-03 | 114.86 | 142.14 | 201.84 | 146.38 | 375.63 | 566.45 | 284.91 | 477.89 |
| **ENSBTAG00000017039** | ARHGEF33 | -1.49 | 1.32E-02 | 66.50 | 22.34 | 38.15 | 87.18 | 210.38 | 91.33 | 149.86 | 151.11 |
| **ENSBTAG00000031737** | TMEM102 | -1.49 | 8.98E-03 | 40.30 | 98.48 | 62.77 | 36.60 | 166.64 | 168.78 | 182.10 | 152.93 |
| **ENSBTAG00000004259** | HPCAL1 | -1.49 | 1.32E-02 | 75.57 | 103.56 | 57.84 | 133.47 | 208.99 | 416.17 | 150.73 | 263.98 |
| **ENSBTAG00000010677** | LIMCH1 | -1.48 | 3.67E-02 | 79.60 | 81.22 | 51.69 | 51.66 | 172.19 | 135.26 | 241.35 | 186.61 |

| | | | | | | | | | | | |
|---|---|---|---|---|---|---|---|---|---|---|---|
| **ENSBTAG00000016108** | BTBD2 | -1.47 | 9.72E-03 | 75.57 | 114.73 | 174.76 | 262.63 | 450.62 | 507.50 | 483.56 | 291.29 |
| **ENSBTAG00000007439** | TENM4 | -1.46 | 1.41E-03 | 72.54 | 44.67 | 43.07 | 105.48 | 193.72 | 153.75 | 212.59 | 173.86 |
| **ENSBTAG00000016091** | KLHL25 | -1.46 | 1.15E-03 | 230.73 | 225.39 | 99.69 | 163.61 | 504.09 | 286.70 | 697.03 | 496.10 |
| **ENSBTAG00000052673** | | -1.46 | 5.27E-04 | 89.67 | 117.77 | 111.99 | 284.16 | 340.22 | 490.16 | 383.36 | 449.68 |
| **ENSBTAG00000020533** | FBXO34 | -1.46 | 6.59E-07 | 560.21 | 437.58 | 440.59 | 342.28 | 967.21 | 1567.58 | 1069.94 | 1301.69 |
| **ENSBTAG00000016910** | EEF1AKMT3 | -1.46 | 3.09E-02 | 29.22 | 34.52 | 132.92 | 96.87 | 204.83 | 297.10 | 155.09 | 149.28 |
| **ENSBTAG00000021527** | IGF1R | -1.46 | 2.27E-04 | 204.54 | 146.20 | 171.07 | 68.89 | 498.53 | 393.05 | 362.45 | 371.39 |
| **ENSBTAG00000004561** | PAX6 | -1.46 | 9.18E-04 | 60.45 | 36.55 | 38.15 | 52.74 | 113.18 | 76.30 | 155.09 | 172.04 |
| **ENSBTAG00000019213** | CNNM4 | -1.46 | 8.27E-03 | 297.23 | 362.45 | 379.06 | 396.10 | 1409.50 | 624.26 | 1081.26 | 825.62 |
| **ENSBTAG00000000895** | TAF9B | -1.46 | 3.67E-02 | 46.35 | 25.38 | 109.53 | 107.64 | 112.48 | 262.42 | 200.40 | 217.56 |
| **ENSBTAG00000047473** | | -1.46 | 4.38E-02 | 269.02 | 283.26 | 279.37 | 117.32 | 1001.92 | 483.22 | 499.25 | 618.99 |
| **ENSBTAG00000045904** | PIM3 | -1.46 | 1.44E-05 | 558.19 | 513.73 | 781.50 | 498.35 | 2105.91 | 973.38 | 1828.82 | 1540.18 |
| **ENSBTAG00000007129** | MRVI1 | -1.46 | 4.89E-03 | 72.54 | 51.78 | 60.30 | 53.82 | 110.40 | 276.29 | 135.05 | 133.81 |
| **ENSBTAG00000016546** | PARP12 | -1.45 | 2.88E-05 | 882.63 | 675.15 | 716.27 | 807.27 | 1831.65 | 2069.29 | 2564.19 | 1978.03 |
| **ENSBTAG00000010582** | BRD3 | -1.45 | 2.69E-08 | 957.19 | 438.60 | 996.87 | 753.45 | 2280.89 | 1910.92 | 2216.54 | 2192.85 |
| **ENSBTAG00000016164** | LMBRD1 | -1.45 | 3.35E-04 | 1325.96 | 1040.65 | 1540.84 | 1098.96 | 2519.04 | 4920.07 | 2863.04 | 3390.77 |
| **ENSBTAG00000013278** | MAGIX | -1.45 | 5.89E-05 | 96.73 | 124.88 | 156.30 | 177.60 | 322.17 | 410.39 | 496.63 | 284.92 |
| **ENSBTAG00000004694** | RPGRIP1 | -1.45 | 4.64E-03 | 468.52 | 195.95 | 297.83 | 315.37 | 626.98 | 1221.92 | 670.89 | 961.25 |
| **ENSBTAG00000010542** | SPIRE1 | -1.45 | 3.00E-05 | 401.01 | 365.50 | 557.51 | 284.16 | 1294.93 | 897.08 | 1120.47 | 1063.20 |
| **ENSBTAG00000004078** | KCNH2 | -1.44 | 5.39E-03 | 68.51 | 65.99 | 77.53 | 78.57 | 120.81 | 215.02 | 214.34 | 240.31 |
| **ENSBTAG00000034393** | FAM149A | -1.44 | 1.56E-02 | 58.44 | 169.55 | 158.76 | 198.05 | 419.38 | 376.87 | 420.83 | 368.66 |
| **ENSBTAG00000011892** | SRGAP1 | -1.44 | 4.47E-03 | 592.45 | 246.71 | 422.13 | 447.76 | 816.54 | 947.94 | 1286.01 | 1582.06 |
| **ENSBTAG00000011101** | EML6 | -1.44 | 4.39E-02 | 39.30 | 10.15 | 22.15 | 64.58 | 57.63 | 113.29 | 71.45 | 127.44 |
| **ENSBTAG00000004442** | AGPAT1 | -1.44 | 5.74E-03 | 96.73 | 131.99 | 98.46 | 226.03 | 290.23 | 240.45 | 582.02 | 384.14 |
| **ENSBTAG00000007101** | F3 | -1.43 | 2.80E-06 | 968.27 | 690.38 | 1025.18 | 717.93 | 1461.57 | 2410.32 | 3069.53 | 2232.90 |
| **ENSBTAG00000016293** | TRIM77 | -1.43 | 3.43E-06 | 885.65 | 389.86 | 999.33 | 645.81 | 1162.31 | 2166.40 | 2436.98 | 2100.00 |
| **ENSBTAG00000004498** | ESR2 | -1.43 | 5.69E-03 | 70.53 | 63.96 | 71.38 | 29.06 | 138.17 | 175.72 | 165.54 | 152.02 |
| **ENSBTAG00000020311** | USP20 | -1.42 | 1.55E-03 | 68.51 | 104.57 | 83.69 | 39.83 | 220.10 | 129.48 | 203.88 | 241.22 |
| **ENSBTAG00000014501** | FERMT2 | -1.42 | 5.98E-04 | 1631.25 | 1183.80 | 1590.07 | 911.67 | 2478.08 | 3678.49 | 4663.98 | 3439.93 |
| **ENSBTAG00000051215** | AGBL2 | -1.42 | 4.12E-05 | 215.62 | 149.24 | 205.53 | 243.26 | 427.02 | 383.80 | 670.02 | 699.09 |
| **ENSBTAG00000007519** | ADAR | -1.42 | 1.01E-05 | 418.14 | 278.18 | 342.14 | 302.46 | 1115.10 | 667.03 | 880.87 | 923.93 |
| **ENSBTAG00000014011** | TMOD2 | -1.41 | 2.63E-06 | 250.88 | 119.80 | 317.52 | 205.58 | 603.38 | 579.17 | 647.36 | 547.07 |
| **ENSBTAG00000017053** | ABCB10 | -1.41 | 4.23E-03 | 262.98 | 348.24 | 588.28 | 266.94 | 702.67 | 1265.85 | 837.30 | 1095.97 |

| | | | | | | | | | | | |
|---|---|---|---|---|---|---|---|---|---|---|---|
| **ENSBTAG00000012278** | MTMR12 | -1.41 | 3.77E-04 | 370.79 | 253.82 | 242.45 | 190.51 | 860.97 | 360.68 | 858.21 | 732.77 |
| **ENSBTAG00000000854** | SEC16B | -1.41 | 2.67E-04 | 212.60 | 167.52 | 287.99 | 365.96 | 915.83 | 567.61 | 460.91 | 794.67 |
| **ENSBTAG00000020780** | SBNO2 | -1.40 | 1.35E-02 | 77.58 | 34.52 | 129.22 | 101.18 | 345.78 | 123.70 | 233.50 | 198.44 |
| **ENSBTAG00000044097** | KLF7 | -1.40 | 9.41E-03 | 30.23 | 35.53 | 54.15 | 62.43 | 126.37 | 97.11 | 96.71 | 159.30 |
| **ENSBTAG00000010452** | PODXL | -1.40 | 3.86E-03 | 113.86 | 121.83 | 115.69 | 58.12 | 360.36 | 228.89 | 232.63 | 256.70 |
| **ENSBTAG00000043957** | TDRP | -1.40 | 1.39E-02 | 66.50 | 70.05 | 98.46 | 51.66 | 152.06 | 213.87 | 203.01 | 185.70 |
| **ENSBTAG00000033008** | MYOZ1 | -1.40 | 2.72E-03 | 48.36 | 36.55 | 19.69 | 47.36 | 118.73 | 105.20 | 103.68 | 73.73 |
| **ENSBTAG00000020193** | DCLRE1A | -1.39 | 9.35E-03 | 430.23 | 193.92 | 155.07 | 175.45 | 453.40 | 722.52 | 679.60 | 654.49 |
| **ENSBTAG00000015177** | PRSS23 | -1.39 | 8.55E-05 | 422.17 | 793.94 | 804.88 | 597.38 | 1540.03 | 1873.93 | 2196.50 | 1258.00 |
| **ENSBTAG00000013249** | SALL2 | -1.38 | 3.46E-02 | 32.24 | 119.80 | 66.46 | 96.87 | 366.61 | 213.87 | 124.59 | 117.43 |
| **ENSBTAG00000020999** | DDB2 | -1.38 | 1.90E-02 | 66.50 | 43.66 | 142.76 | 60.28 | 286.76 | 215.02 | 189.07 | 119.25 |
| **ENSBTAG00000009565** | RASA1 | -1.37 | 2.50E-05 | 450.38 | 680.23 | 937.80 | 811.57 | 1159.54 | 2072.76 | 1829.69 | 2363.98 |
| **ENSBTAG00000014906** | VCAN | -1.37 | 2.86E-02 | 47.36 | 141.12 | 141.53 | 105.48 | 145.12 | 341.03 | 313.66 | 323.15 |
| **ENSBTAG00000010360** | LRIG1 | -1.37 | 2.58E-03 | 128.97 | 114.73 | 132.92 | 132.39 | 428.40 | 151.44 | 382.49 | 347.72 |
| **ENSBTAG00000021458** | DLX6 | -1.36 | 5.51E-03 | 291.19 | 114.73 | 204.30 | 176.52 | 426.32 | 497.09 | 572.43 | 525.23 |
| **ENSBTAG00000015982** | KCTD13 | -1.36 | 3.71E-09 | 540.06 | 552.31 | 927.95 | 663.03 | 1471.99 | 1702.83 | 1732.98 | 1960.73 |
| **ENSBTAG00000011910** | STK33 | -1.35 | 3.72E-02 | 48.36 | 64.98 | 59.07 | 83.96 | 115.26 | 191.90 | 189.94 | 156.57 |
| **ENSBTAG00000053114** | | -1.35 | 9.39E-03 | 360.71 | 82.24 | 283.06 | 129.16 | 335.36 | 421.95 | 616.87 | 798.31 |
| **ENSBTAG00000002144** | ADRB2 | -1.35 | 5.29E-04 | 1344.10 | 754.35 | 617.81 | 863.24 | 1355.34 | 2070.45 | 3971.31 | 1699.48 |
| **ENSBTAG00000012500** | RARA | -1.34 | 5.29E-03 | 48.36 | 44.67 | 43.07 | 104.41 | 182.61 | 176.87 | 135.92 | 115.60 |
| **ENSBTAG00000005475** | TCAF2 | -1.34 | 3.73E-02 | 57.43 | 146.20 | 105.84 | 100.10 | 219.41 | 286.70 | 180.36 | 353.19 |
| **ENSBTAG00000009290** | FAM161A | -1.34 | 2.44E-06 | 983.39 | 661.96 | 1142.10 | 1061.28 | 1601.13 | 2656.56 | 2522.36 | 2964.76 |
| **ENSBTAG00000023179** | TRIB1 | -1.34 | 3.88E-04 | 947.12 | 1058.93 | 958.72 | 494.05 | 1997.60 | 2038.08 | 2470.96 | 2249.29 |
| **ENSBTAG00000005244** | RASL11A | -1.34 | 1.44E-05 | 434.26 | 266.00 | 601.82 | 327.21 | 770.71 | 1201.12 | 1299.95 | 850.20 |
| **ENSBTAG00000054030** | | -1.34 | 5.99E-03 | 84.64 | 91.37 | 123.07 | 127.01 | 111.79 | 406.92 | 192.55 | 366.84 |
| **ENSBTAG00000002363** | SESN2 | -1.34 | 4.96E-02 | 339.55 | 141.12 | 210.45 | 103.33 | 901.24 | 253.17 | 447.84 | 402.34 |
| **ENSBTAG00000031793** | NUDT2 | -1.33 | 4.14E-06 | 244.84 | 244.68 | 305.22 | 135.62 | 479.09 | 628.88 | 555.01 | 681.80 |
| **ENSBTAG00000008898** | GBX2 | -1.33 | 5.18E-03 | 111.84 | 40.61 | 121.84 | 88.26 | 177.75 | 130.63 | 303.21 | 300.39 |
| **ENSBTAG00000009355** | SNAP91 | -1.33 | 7.34E-03 | 89.67 | 67.01 | 103.38 | 109.79 | 204.13 | 262.42 | 266.61 | 197.53 |
| **ENSBTAG00000015462** | BNC1 | -1.33 | 3.96E-02 | 127.96 | 105.59 | 62.77 | 87.18 | 286.76 | 186.12 | 273.58 | 217.56 |
| **ENSBTAG00000019187** | ZNF462 | -1.33 | 1.05E-02 | 198.49 | 161.43 | 129.22 | 90.41 | 316.62 | 343.34 | 313.66 | 481.53 |
| **ENSBTAG00000013175** | KIAA0355 | -1.33 | 7.19E-05 | 439.30 | 386.82 | 334.75 | 336.90 | 806.82 | 879.74 | 969.74 | 1102.34 |
| **ENSBTAG00000009076** | ADD2 | -1.33 | 4.99E-02 | 52.39 | 54.82 | 71.38 | 104.41 | 217.33 | 201.15 | 184.71 | 105.59 |

| | | | | | | | | | | | |
|---|---|---|---|---|---|---|---|---|---|---|---|
| **ENSBTAG00000004018** | DGKA | -1.33 | 8.46E-03 | 329.48 | 787.85 | 409.83 | 340.13 | 1457.41 | 862.40 | 1323.48 | 1034.98 |
| **ENSBTAG00000019234** | BMP6 | -1.32 | 2.26E-02 | 61.46 | 21.32 | 68.92 | 69.96 | 108.32 | 152.60 | 151.60 | 142.00 |
| **ENSBTAG00000033621** | SMCO4 | -1.32 | 3.98E-13 | 651.90 | 702.57 | 815.96 | 595.22 | 1456.71 | 1706.30 | 2174.72 | 1584.79 |
| **ENSBTAG00000001693** | RPAP1 | -1.32 | 2.06E-06 | 514.87 | 309.66 | 370.44 | 438.08 | 1293.54 | 714.43 | 1025.50 | 1047.73 |
| **ENSBTAG00000008895** | BPGM | -1.32 | 3.38E-06 | 1087.17 | 793.94 | 1053.48 | 575.85 | 1441.44 | 2626.50 | 2582.48 | 2103.64 |
| **ENSBTAG00000000987** | OTOG | -1.31 | 2.49E-02 | 66.50 | 46.70 | 105.84 | 103.33 | 165.25 | 78.61 | 276.20 | 279.45 |
| **ENSBTAG00000026676** | SDC3 | -1.31 | 1.05E-02 | 188.42 | 75.13 | 139.07 | 95.80 | 377.02 | 299.41 | 380.75 | 179.32 |
| **ENSBTAG00000012032** | PDE4A | -1.31 | 2.93E-03 | 86.65 | 180.72 | 114.46 | 171.14 | 376.33 | 307.50 | 386.85 | 301.30 |
| **ENSBTAG00000025634** | FMN1 | -1.31 | 1.26E-02 | 121.92 | 139.09 | 142.76 | 100.10 | 233.30 | 194.21 | 368.55 | 451.50 |
| **ENSBTAG00000006747** | LTBP3 | -1.31 | 3.71E-03 | 128.97 | 37.56 | 102.15 | 85.03 | 171.50 | 158.38 | 285.78 | 259.43 |
| **ENSBTAG00000016600** | CCDC142 | -1.31 | 2.64E-02 | 18.14 | 31.47 | 44.31 | 49.51 | 140.95 | 62.43 | 96.71 | 52.80 |
| **ENSBTAG00000038532** | | -1.31 | 3.78E-02 | 55.42 | 166.50 | 36.92 | 45.21 | 151.36 | 228.89 | 210.85 | 162.94 |
| **ENSBTAG00000017281** | OPLAH | -1.31 | 5.80E-04 | 120.91 | 142.14 | 130.45 | 116.25 | 372.86 | 337.56 | 321.50 | 227.57 |
| **ENSBTAG00000002988** | MACO1 | -1.31 | 8.72E-05 | 419.15 | 486.31 | 300.29 | 278.78 | 634.62 | 1069.33 | 1017.66 | 950.33 |
| **ENSBTAG00000018404** | PRKG1 | -1.30 | 1.32E-02 | 49.37 | 48.73 | 81.23 | 91.49 | 81.24 | 218.49 | 203.88 | 165.67 |
| **ENSBTAG00000000181** | SUSD3 | -1.30 | 8.71E-05 | 123.93 | 169.55 | 194.45 | 104.41 | 280.51 | 478.60 | 433.90 | 269.44 |
| **ENSBTAG00000001068** | ZCWPW1 | -1.30 | 4.82E-03 | 256.93 | 167.52 | 327.37 | 135.62 | 497.84 | 658.94 | 661.30 | 371.39 |
| **ENSBTAG00000018902** | ZC2HC1A | -1.30 | 8.33E-09 | 4820.21 | 3980.87 | 4789.91 | 3413.12 | 7380.07 | 13468.91 | 12121.29 | 8897.93 |
| **ENSBTAG00000016175** | SPNS3 | -1.30 | 1.10E-02 | 25.19 | 43.66 | 32.00 | 38.75 | 60.41 | 80.92 | 71.45 | 131.08 |
| **ENSBTAG00000008844** | EXTL3 | -1.30 | 3.87E-02 | 143.07 | 67.01 | 156.30 | 151.77 | 381.19 | 199.99 | 296.24 | 394.15 |
| **ENSBTAG00000046701** | FRAT2 | -1.29 | 2.59E-04 | 570.28 | 237.57 | 433.21 | 463.91 | 1245.63 | 624.26 | 1392.31 | 913.92 |
| **ENSBTAG00000044070** | SNX30 | -1.29 | 6.43E-03 | 185.39 | 197.98 | 196.91 | 117.32 | 517.97 | 306.35 | 407.76 | 474.25 |
| **ENSBTAG00000009744** | FAM193A | -1.29 | 6.72E-03 | 218.64 | 213.21 | 248.60 | 76.42 | 506.86 | 369.93 | 478.33 | 488.82 |
| **ENSBTAG00000033563** | ZNF529 | -1.28 | 2.20E-04 | 357.69 | 203.05 | 275.68 | 269.09 | 367.30 | 1048.52 | 619.48 | 661.77 |
| **ENSBTAG00000019406** | IGF2BP3 | -1.28 | 2.81E-05 | 2290.21 | 1959.47 | 2041.74 | 1003.16 | 3150.89 | 4940.87 | 5848.05 | 3825.88 |
| **ENSBTAG00000009194** | OSBPL10 | -1.28 | 4.79E-03 | 108.82 | 114.73 | 147.68 | 191.59 | 298.56 | 261.26 | 456.55 | 347.72 |
| **ENSBTAG00000021210** | HES2 | -1.28 | 5.75E-03 | 162.22 | 172.60 | 275.68 | 274.47 | 681.14 | 334.09 | 410.37 | 714.57 |
| **ENSBTAG00000018297** | PLEKHH1 | -1.28 | 4.90E-02 | 54.41 | 78.18 | 119.38 | 38.75 | 286.07 | 124.85 | 137.66 | 152.93 |
| **ENSBTAG00000009653** | | -1.27 | 5.37E-03 | 312.35 | 358.39 | 280.60 | 249.71 | 642.26 | 465.88 | 959.28 | 832.90 |
| **ENSBTAG00000048576** | | -1.27 | 1.71E-02 | 125.95 | 90.36 | 49.23 | 53.82 | 201.36 | 199.99 | 202.14 | 169.31 |
| **ENSBTAG00000002385** | SLC35E4 | -1.27 | 1.94E-02 | 120.91 | 161.43 | 100.92 | 167.91 | 536.72 | 225.43 | 338.06 | 227.57 |
| **ENSBTAG00000008991** | EIF4ENIF1 | -1.27 | 1.43E-06 | 787.92 | 524.89 | 814.73 | 696.40 | 2087.86 | 1521.34 | 1579.64 | 1609.36 |
| **ENSBTAG00000014791** | CTH | -1.27 | 1.30E-02 | 69.52 | 113.71 | 86.15 | 146.38 | 164.56 | 245.08 | 359.84 | 232.12 |

| | | | | | | | | | | | |
|---|---|---|---|---|---|---|---|---|---|---|---|
| **ENSBTAG00000012074** | MYB | -1.26 | 2.90E-02 | 529.98 | 112.69 | 323.68 | 91.49 | 742.94 | 586.11 | 610.77 | 601.69 |
| **ENSBTAG00000054533** | GNG12 | -1.26 | 8.07E-05 | 641.82 | 337.07 | 414.75 | 635.05 | 906.11 | 1152.56 | 1351.36 | 1465.54 |
| **ENSBTAG00000025046** | ALKBH5 | -1.26 | 2.23E-12 | 3931.54 | 3971.74 | 5221.89 | 3935.15 | 11280.84 | 9139.58 | 10416.19 | 10089.48 |
| **ENSBTAG00000008111** | ESYT3 | -1.26 | 8.27E-03 | 87.66 | 144.17 | 148.92 | 82.88 | 418.68 | 257.79 | 244.83 | 187.52 |
| **ENSBTAG00000001060** | CXCR4 | -1.26 | 7.37E-06 | 2406.08 | 1295.48 | 1735.30 | 1511.20 | 3539.02 | 3693.52 | 4298.04 | 5104.82 |
| **ENSBTAG00000016275** | AMDHD1 | -1.25 | 1.70E-03 | 460.46 | 209.15 | 324.91 | 286.31 | 518.67 | 865.87 | 880.00 | 788.30 |
| **ENSBTAG00000052516** | CDA | -1.25 | 1.45E-03 | 79.60 | 159.40 | 153.84 | 122.70 | 377.02 | 285.54 | 280.55 | 281.27 |
| **ENSBTAG00000018598** | HSPB6 | -1.25 | 4.81E-03 | 144.08 | 129.95 | 145.22 | 161.45 | 406.88 | 290.16 | 360.71 | 322.24 |
| **ENSBTAG00000008330** | RNF19B | -1.25 | 1.18E-03 | 626.71 | 379.71 | 795.04 | 565.09 | 1719.17 | 860.09 | 1587.48 | 1456.44 |
| **ENSBTAG00000013048** | NIPAL3 | -1.25 | 3.14E-02 | 113.86 | 98.48 | 109.53 | 22.60 | 211.08 | 255.48 | 166.42 | 185.70 |
| **ENSBTAG00000011761** | LRP6 | -1.25 | 2.14E-06 | 658.95 | 406.11 | 337.21 | 406.86 | 993.59 | 1073.95 | 1105.66 | 1127.83 |
| **ENSBTAG00000020854** | BCL6B | -1.25 | 1.25E-02 | 170.28 | 102.54 | 135.38 | 92.57 | 199.97 | 418.48 | 376.39 | 195.71 |
| **ENSBTAG00000003994** | IGFBP3 | -1.25 | 5.36E-06 | 317.38 | 396.97 | 508.28 | 362.73 | 1124.13 | 1119.04 | 587.24 | 925.75 |
| **ENSBTAG00000011011** | SSH2 | -1.24 | 2.40E-03 | 309.32 | 176.66 | 264.60 | 284.16 | 572.83 | 628.88 | 764.12 | 485.18 |
| **ENSBTAG00000000816** | PRDM1 | -1.24 | 4.31E-02 | 190.43 | 255.85 | 35.69 | 119.48 | 236.77 | 320.22 | 602.93 | 266.71 |
| **ENSBTAG00000019553** | | -1.24 | 4.80E-02 | 271.04 | 173.61 | 172.30 | 99.02 | 212.47 | 597.67 | 325.86 | 558.00 |
| **ENSBTAG00000021752** | DNAJB4 | -1.24 | 1.99E-03 | 114.86 | 292.40 | 148.92 | 160.38 | 485.34 | 336.40 | 327.60 | 544.34 |
| **ENSBTAG00000032137** | PNPLA6 | -1.24 | 2.88E-03 | 75.57 | 81.22 | 157.53 | 83.96 | 198.58 | 240.45 | 264.87 | 234.85 |
| **ENSBTAG00000019581** | PLEKHF2 | -1.24 | 1.39E-04 | 1449.89 | 840.64 | 1145.79 | 991.32 | 2061.48 | 3165.21 | 2854.32 | 2372.17 |
| **ENSBTAG00000011772** | PPP1R12B | -1.24 | 4.07E-02 | 49.37 | 84.27 | 70.15 | 47.36 | 220.80 | 146.82 | 108.91 | 115.60 |
| **ENSBTAG00000005252** | NUDT11 | -1.24 | 8.97E-03 | 214.61 | 141.12 | 238.76 | 252.94 | 419.38 | 489.00 | 651.72 | 437.84 |
| **ENSBTAG00000010693** | LMO7 | -1.24 | 3.71E-04 | 359.70 | 202.04 | 354.44 | 259.40 | 633.93 | 898.24 | 709.22 | 529.78 |
| **ENSBTAG00000002689** | NME7 | -1.24 | 4.90E-03 | 83.63 | 228.44 | 114.46 | 158.22 | 231.91 | 412.70 | 331.09 | 404.16 |
| **ENSBTAG00000024509** | TMEM52 | -1.23 | 3.19E-02 | 33.25 | 16.24 | 23.38 | 32.29 | 72.21 | 80.92 | 50.53 | 43.69 |
| **ENSBTAG00000010626** | SH3GLB2 | -1.23 | 5.65E-03 | 121.92 | 223.36 | 232.60 | 307.84 | 649.20 | 393.05 | 528.87 | 505.20 |
| **ENSBTAG00000010450** | WIZ | -1.23 | 1.12E-03 | 233.76 | 147.21 | 114.46 | 188.36 | 571.44 | 290.16 | 392.95 | 346.81 |
| **ENSBTAG00000021420** | EPHA7 | -1.23 | 2.02E-02 | 80.61 | 161.43 | 142.76 | 76.42 | 233.30 | 228.89 | 281.42 | 335.89 |
| **ENSBTAG00000011504** | ZP2 | -1.23 | 3.33E-04 | 576.33 | 525.91 | 343.37 | 414.40 | 978.32 | 1109.79 | 828.59 | 1435.50 |
| **ENSBTAG00000034368** | PRSS33 | -1.22 | 4.82E-02 | 100.76 | 89.34 | 70.15 | 118.40 | 106.93 | 110.98 | 426.06 | 241.22 |
| **ENSBTAG00000020878** | DMTF1 | -1.22 | 2.25E-06 | 202.52 | 172.60 | 180.91 | 179.75 | 436.04 | 375.71 | 463.52 | 439.66 |
| **ENSBTAG00000005824** | SPOP | -1.22 | 1.42E-04 | 677.09 | 434.54 | 751.96 | 597.38 | 1010.26 | 1908.61 | 1281.66 | 1531.99 |
| **ENSBTAG00000020644** | GPC4 | -1.22 | 2.26E-02 | 358.69 | 337.07 | 281.83 | 226.03 | 858.89 | 825.41 | 622.97 | 494.28 |
| **ENSBTAG00000020346** | CCDC92 | -1.22 | 2.07E-02 | 97.73 | 60.92 | 94.76 | 62.43 | 197.19 | 169.94 | 196.04 | 170.22 |

| | | | | | | | | | | | |
|---|---|---|---|---|---|---|---|---|---|---|---|
| **ENSBTAG00000038084** | NAA80 | -1.22 | 1.21E-02 | 110.83 | 74.11 | 91.07 | 41.98 | 174.28 | 132.94 | 313.66 | 117.43 |
| **ENSBTAG00000015887** | FOXJ3 | -1.22 | 6.37E-05 | 632.75 | 620.33 | 527.97 | 572.62 | 1185.92 | 1454.29 | 1623.20 | 1205.20 |
| **ENSBTAG00000016297** | CCNYL1 | -1.22 | 1.19E-05 | 632.75 | 306.61 | 404.90 | 373.49 | 729.05 | 1069.33 | 1307.80 | 885.70 |
| **ENSBTAG00000019168** | PHOX2A | -1.21 | 9.33E-04 | 164.23 | 169.55 | 204.30 | 121.63 | 236.77 | 483.22 | 508.83 | 300.39 |
| **ENSBTAG00000017602** | TMEM45B | -1.21 | 3.12E-02 | 244.84 | 167.52 | 259.68 | 133.47 | 212.47 | 831.19 | 656.95 | 162.03 |
| **ENSBTAG00000015913** | MFHAS1 | -1.21 | 1.19E-02 | 248.87 | 258.89 | 217.83 | 559.70 | 845.00 | 591.89 | 651.72 | 879.32 |
| **ENSBTAG00000009126** | YBX2 | -1.21 | 5.18E-05 | 462.47 | 382.76 | 559.97 | 430.54 | 1039.42 | 845.06 | 1322.61 | 1027.70 |
| **ENSBTAG00000015904** | RORA | -1.21 | 3.26E-02 | 99.75 | 103.56 | 43.07 | 100.10 | 185.39 | 253.17 | 220.43 | 142.00 |
| **ENSBTAG00000005376** | FBXO30 | -1.20 | 4.79E-04 | 877.59 | 438.60 | 829.50 | 579.08 | 1826.10 | 1211.52 | 1508.19 | 1696.75 |
| **ENSBTAG00000014191** | QSOX1 | -1.20 | 4.66E-02 | 76.58 | 56.86 | 102.15 | 120.55 | 180.53 | 149.13 | 175.13 | 310.40 |
| **ENSBTAG00000030599** | SMOC1 | -1.20 | 6.57E-05 | 193.45 | 146.20 | 274.45 | 200.20 | 420.07 | 443.92 | 514.93 | 484.27 |
| **ENSBTAG00000047717** | FAM222B | -1.19 | 7.29E-06 | 2487.69 | 1716.82 | 2758.01 | 2005.25 | 6689.21 | 3900.45 | 5428.09 | 4494.02 |
| **ENSBTAG00000001120** | CORO6 | -1.19 | 1.18E-05 | 606.56 | 315.75 | 502.13 | 475.75 | 1062.33 | 981.47 | 1174.49 | 1123.28 |
| **ENSBTAG00000019479** | | -1.19 | 8.27E-03 | 116.88 | 130.97 | 49.23 | 116.25 | 181.92 | 189.59 | 297.11 | 277.63 |
| **ENSBTAG00000004899** | ABLIM1 | -1.19 | 1.77E-05 | 910.84 | 688.35 | 633.81 | 712.55 | 1166.48 | 2102.82 | 1665.02 | 1770.48 |
| **ENSBTAG00000007103** | ITGAL | -1.19 | 5.03E-04 | 171.29 | 123.86 | 164.91 | 148.54 | 243.71 | 343.34 | 419.96 | 377.76 |
| **ENSBTAG00000000638** | CDT1 | -1.18 | 4.47E-03 | 321.41 | 174.63 | 503.36 | 450.99 | 1047.05 | 715.58 | 752.79 | 760.99 |
| **ENSBTAG00000006573** | KLHL18 | -1.18 | 1.24E-02 | 436.28 | 478.19 | 633.81 | 496.20 | 1757.36 | 1077.42 | 1014.17 | 771.00 |
| **ENSBTAG00000007634** | HOOK3 | -1.18 | 1.17E-04 | 1096.24 | 953.34 | 898.42 | 562.93 | 2045.51 | 2199.93 | 1706.84 | 1984.40 |
| **ENSBTAG00000000561** | OCLN | -1.18 | 2.55E-08 | 5144.65 | 3837.72 | 3290.91 | 3047.16 | 6852.38 | 11264.36 | 8380.87 | 8127.84 |
| **ENSBTAG00000054342** | | -1.18 | 1.59E-02 | 54.41 | 32.49 | 35.69 | 39.83 | 63.88 | 82.08 | 113.27 | 108.32 |
| **ENSBTAG00000000949** | DNAJC9 | -1.17 | 3.57E-04 | 763.74 | 482.25 | 716.27 | 591.99 | 1250.49 | 1462.38 | 1474.21 | 1579.33 |
| **ENSBTAG00000016247** | TACR3 | -1.17 | 1.99E-02 | 288.16 | 91.37 | 201.84 | 315.37 | 374.25 | 731.77 | 561.98 | 351.37 |
| **ENSBTAG00000048999** | TRIM6 | -1.17 | 6.14E-03 | 1076.08 | 980.75 | 1531.00 | 653.35 | 2804.41 | 2358.30 | 2285.37 | 2087.26 |
| **ENSBTAG00000033747** | SIL1 | -1.17 | 1.51E-02 | 67.51 | 193.92 | 166.15 | 91.49 | 286.07 | 309.82 | 341.54 | 228.48 |
| **ENSBTAG00000008147** | MICAL1 | -1.17 | 1.48E-02 | 1023.69 | 634.54 | 1054.71 | 474.67 | 3100.89 | 1095.92 | 1954.29 | 1008.58 |
| **ENSBTAG00000021021** | C21H15orf39 | -1.17 | 2.06E-03 | 766.76 | 686.32 | 1021.49 | 675.95 | 3040.49 | 1246.20 | 1429.78 | 1344.47 |
| **ENSBTAG00000011905** | RRAS2 | -1.16 | 1.78E-07 | 4417.18 | 4517.95 | 5420.03 | 4242.99 | 9926.19 | 12739.46 | 9263.48 | 9663.47 |
| **ENSBTAG00000021538** | MYO1E | -1.15 | 9.75E-03 | 182.37 | 78.18 | 185.84 | 244.33 | 439.51 | 487.84 | 256.16 | 353.19 |
| **ENSBTAG00000014126** | | -1.15 | 7.00E-03 | 124.94 | 249.76 | 205.53 | 105.48 | 502.70 | 278.60 | 351.13 | 390.51 |
| **ENSBTAG00000009443** | GCNT3 | -1.15 | 2.36E-02 | 350.63 | 224.37 | 194.45 | 195.90 | 556.16 | 452.01 | 575.92 | 560.73 |
| **ENSBTAG00000020764** | CNN2 | -1.15 | 3.78E-03 | 144.08 | 198.99 | 237.53 | 252.94 | 304.12 | 450.85 | 538.45 | 556.18 |
| **ENSBTAG00000004976** | CDCA7L | -1.15 | 1.53E-02 | 258.95 | 327.93 | 182.14 | 433.77 | 458.26 | 760.67 | 729.26 | 722.76 |

| | | | | | | | | | | | |
|---|---|---|---|---|---|---|---|---|---|---|---|
| **ENSBTAG00000049752** | | -1.15 | 4.48E-04 | 9525.56 | 6092.64 | 7280.86 | 6579.75 | 21294.51 | 6776.65 | 17935.36 | 19349.69 |
| **ENSBTAG00000053030** | GPR27 | -1.14 | 3.28E-02 | 55.42 | 25.38 | 113.22 | 78.57 | 154.14 | 159.53 | 156.83 | 128.35 |
| **ENSBTAG00000020861** | CHAMP1 | -1.14 | 4.11E-06 | 1042.83 | 1051.82 | 1323.01 | 702.86 | 1935.11 | 2308.59 | 2756.74 | 2066.32 |
| **ENSBTAG00000002315** | RNF34 | -1.14 | 7.66E-08 | 3391.48 | 2599.09 | 3108.76 | 2686.58 | 4221.55 | 6864.51 | 7549.67 | 7294.03 |
| **ENSBTAG00000047975** | DOK5 | -1.14 | 3.99E-02 | 138.04 | 112.69 | 66.46 | 71.04 | 74.29 | 211.55 | 352.00 | 217.56 |
| **ENSBTAG00000009512** | EPS8L1 | -1.14 | 4.79E-03 | 254.92 | 174.63 | 191.99 | 75.34 | 471.45 | 267.04 | 421.70 | 369.57 |
| **ENSBTAG00000004266** | DDX25 | -1.13 | 1.32E-02 | 801.02 | 431.49 | 319.98 | 418.70 | 946.38 | 562.99 | 1922.05 | 885.70 |
| **ENSBTAG00000047502** | FKBP5 | -1.13 | 2.03E-03 | 561.22 | 446.72 | 555.05 | 745.91 | 1176.90 | 1676.24 | 1016.79 | 1183.36 |
| **ENSBTAG00000021535** | CROT | -1.13 | 3.68E-03 | 1029.74 | 433.52 | 884.88 | 725.46 | 868.61 | 2246.17 | 1448.94 | 2148.25 |
| **ENSBTAG00000026829** | TMEM185B | -1.13 | 1.00E-04 | 325.44 | 492.41 | 294.14 | 515.57 | 666.56 | 961.82 | 856.47 | 1071.39 |
| **ENSBTAG00000005650** | SKAP2 | -1.12 | 3.80E-02 | 180.35 | 151.28 | 191.99 | 90.41 | 347.86 | 338.72 | 216.95 | 431.47 |
| **ENSBTAG00000001481** | IGSF9 | -1.12 | 4.10E-02 | 94.71 | 151.28 | 71.38 | 39.83 | 185.39 | 158.38 | 239.60 | 193.89 |
| **ENSBTAG00000002630** | MRTFA | -1.12 | 3.63E-02 | 50.38 | 76.15 | 38.15 | 106.56 | 168.72 | 141.04 | 174.26 | 105.59 |
| **ENSBTAG00000010681** | NR1H3 | -1.12 | 4.38E-02 | 145.09 | 154.32 | 238.76 | 44.13 | 327.73 | 199.99 | 325.86 | 406.89 |
| **ENSBTAG00000006695** | VCPIP1 | -1.11 | 2.49E-04 | 486.66 | 368.54 | 550.13 | 370.27 | 1372.00 | 720.21 | 1011.56 | 737.32 |
| **ENSBTAG00000018910** | WDR6 | -1.11 | 3.97E-03 | 2066.52 | 1315.79 | 1823.91 | 1966.50 | 6380.23 | 2034.61 | 4023.58 | 3080.37 |
| **ENSBTAG00000055062** | SMAD5 | -1.11 | 1.20E-03 | 466.50 | 237.57 | 319.98 | 292.77 | 420.07 | 869.33 | 981.06 | 580.75 |
| **ENSBTAG00000007666** | IGF2BP2 | -1.11 | 3.72E-02 | 146.10 | 101.53 | 134.15 | 192.67 | 494.37 | 213.87 | 342.41 | 188.43 |
| **ENSBTAG00000017442** | CDO1 | -1.11 | 3.81E-03 | 682.12 | 1182.79 | 601.82 | 729.77 | 892.91 | 1990.68 | 2003.95 | 2018.08 |
| **ENSBTAG00000011857** | PPM1H | -1.11 | 4.57E-02 | 105.79 | 106.60 | 143.99 | 115.17 | 335.36 | 322.53 | 225.66 | 131.99 |
| **ENSBTAG00000018050** | CDR2 | -1.10 | 1.68E-02 | 613.61 | 555.35 | 676.89 | 416.55 | 2171.18 | 598.82 | 1202.37 | 887.52 |
| **ENSBTAG00000015835** | PIWIL2 | -1.10 | 2.36E-02 | 420.16 | 270.06 | 342.14 | 188.36 | 665.87 | 428.89 | 833.82 | 690.90 |
| **ENSBTAG00000012751** | BTBD6 | -1.10 | 7.07E-03 | 406.05 | 227.42 | 540.28 | 215.27 | 994.98 | 872.80 | 676.12 | 431.47 |
| **ENSBTAG00000033727** | RBPMS | -1.10 | 1.31E-03 | 1504.30 | 2415.33 | 2284.19 | 1766.30 | 3100.89 | 6068.00 | 4102.00 | 3781.28 |
| **ENSBTAG00000018900** | CARMIL2 | -1.10 | 4.62E-04 | 223.68 | 225.39 | 235.06 | 214.19 | 532.55 | 442.76 | 512.31 | 430.56 |
| **ENSBTAG00000007186** | ARHGAP39 | -1.09 | 1.45E-02 | 56.42 | 56.86 | 65.23 | 61.35 | 106.23 | 216.18 | 90.61 | 101.04 |
| **ENSBTAG00000005788** | VANGL1 | -1.09 | 3.27E-02 | 135.01 | 63.96 | 132.92 | 90.41 | 231.21 | 297.10 | 252.67 | 118.34 |
| **ENSBTAG00000003474** | RAB15 | -1.09 | 2.99E-02 | 405.04 | 378.70 | 290.45 | 438.08 | 1176.90 | 431.20 | 887.84 | 722.76 |
| **ENSBTAG00000015166** | PNLDC1 | -1.09 | 9.44E-06 | 405.04 | 537.08 | 589.51 | 550.02 | 960.26 | 1092.45 | 1180.59 | 1196.10 |
| **ENSBTAG00000046684** | FOXN3 | -1.09 | 1.39E-04 | 421.16 | 335.04 | 525.51 | 318.60 | 868.61 | 773.38 | 859.08 | 899.35 |
| **ENSBTAG00000013901** | PTDSS1 | -1.09 | 1.35E-02 | 934.02 | 596.98 | 903.34 | 518.80 | 2421.14 | 1268.17 | 1281.66 | 1297.14 |
| **ENSBTAG00000003390** | SF3A1 | -1.09 | 4.05E-03 | 2325.47 | 2257.96 | 2685.40 | 2294.79 | 9115.90 | 2512.05 | 4710.16 | 3957.87 |
| **ENSBTAG00000011150** | PFN2 | -1.08 | 4.35E-03 | 507.81 | 609.16 | 701.50 | 635.05 | 1188.00 | 1530.58 | 1143.12 | 1324.45 |

| | | | | | | | | | | | |
|---|---|---|---|---|---|---|---|---|---|---|---|
| **ENSBTAG00000007007** | WDR20 | -1.08 | 9.52E-13 | 1839.82 | 1769.62 | 1945.75 | 1838.41 | 3616.09 | 4224.14 | 3953.01 | 3820.42 |
| **ENSBTAG00000020283** | DUSP5 | -1.07 | 1.68E-05 | 1011.60 | 898.51 | 876.26 | 685.64 | 1649.74 | 2300.50 | 1677.22 | 1685.83 |
| **ENSBTAG00000021570** | PLEKHG4 | -1.07 | 1.57E-02 | 916.89 | 802.06 | 1235.63 | 1301.31 | 4315.98 | 1125.97 | 1512.55 | 1991.68 |
| **ENSBTAG00000010324** | GTPBP1 | -1.07 | 2.39E-04 | 505.80 | 513.73 | 495.97 | 425.16 | 1349.09 | 938.70 | 940.99 | 847.47 |
| **ENSBTAG00000011904** | HCFC1 | -1.06 | 3.48E-04 | 1950.65 | 1207.16 | 2070.05 | 1758.76 | 5344.28 | 2714.36 | 3882.44 | 2654.36 |
| **ENSBTAG00000014855** | MICAL2 | -1.06 | 2.10E-02 | 221.67 | 238.59 | 403.67 | 163.61 | 713.77 | 357.21 | 494.89 | 576.20 |
| **ENSBTAG00000019107** | GAS7 | -1.06 | 4.47E-03 | 454.41 | 166.50 | 457.82 | 278.78 | 597.13 | 605.76 | 927.92 | 698.18 |
| **ENSBTAG00000017245** | COPG2 | -1.06 | 1.03E-02 | 355.67 | 527.94 | 633.81 | 276.62 | 636.70 | 1267.01 | 837.30 | 999.48 |
| **ENSBTAG00000018307** | SEMA3F | -1.06 | 1.06E-02 | 172.29 | 118.79 | 163.68 | 137.77 | 357.58 | 173.40 | 385.98 | 314.04 |
| **ENSBTAG00000004675** | ZNF18 | -1.05 | 4.48E-02 | 247.86 | 150.26 | 465.21 | 132.39 | 587.41 | 525.99 | 433.90 | 517.04 |
| **ENSBTAG00000004190** | ARHGAP29 | -1.05 | 2.67E-05 | 627.72 | 551.29 | 728.58 | 851.40 | 1276.18 | 1708.61 | 1316.51 | 1411.83 |
| **ENSBTAG00000037832** | ASPHD2 | -1.05 | 4.81E-02 | 80.61 | 162.44 | 116.92 | 156.07 | 392.99 | 351.43 | 161.19 | 161.12 |
| **ENSBTAG00000014137** | CEP164 | -1.05 | 9.30E-04 | 540.06 | 561.44 | 651.04 | 550.02 | 747.10 | 1528.27 | 1274.69 | 1205.20 |
| **ENSBTAG00000007147** | CUEDC1 | -1.04 | 3.20E-02 | 302.27 | 360.42 | 180.91 | 164.68 | 583.93 | 394.21 | 569.82 | 532.51 |
| **ENSBTAG00000053034** | H4 | -1.04 | 3.68E-02 | 116.88 | 48.73 | 129.22 | 45.21 | 171.50 | 194.21 | 170.77 | 163.85 |
| **ENSBTAG00000009760** | | -1.04 | 4.14E-03 | 563.23 | 346.21 | 808.57 | 571.54 | 1353.95 | 1218.46 | 1251.16 | 893.89 |
| **ENSBTAG00000047756** | CNTROB | -1.04 | 3.04E-02 | 234.76 | 196.96 | 164.91 | 121.63 | 359.66 | 485.53 | 310.18 | 324.06 |
| **ENSBTAG00000025441** | HSPA1A | -1.04 | 2.19E-03 | 2591.47 | 1773.68 | 2914.31 | 2651.06 | 5365.11 | 2967.53 | 4510.63 | 7526.15 |
| **ENSBTAG00000020014** | CEP104 | -1.04 | 3.04E-02 | 272.04 | 222.34 | 157.53 | 340.13 | 840.84 | 438.14 | 430.41 | 323.15 |
| **ENSBTAG00000019250** | BTK | -1.03 | 1.69E-02 | 548.12 | 495.45 | 484.90 | 254.02 | 1056.08 | 1101.70 | 602.93 | 884.79 |
| **ENSBTAG00000021128** | ADD1 | -1.03 | 2.48E-13 | 1007.57 | 958.41 | 983.33 | 978.41 | 1960.10 | 1883.17 | 2242.68 | 1938.88 |
| **ENSBTAG00000007123** | ENSA | -1.03 | 1.27E-07 | 2397.01 | 2253.90 | 2449.10 | 1792.13 | 3675.11 | 5555.88 | 4156.89 | 4769.84 |
| **ENSBTAG00000005234** | LYPD1 | -1.03 | 1.39E-04 | 879.61 | 709.67 | 910.72 | 440.23 | 1187.31 | 1677.40 | 1505.58 | 1616.65 |
| **ENSBTAG00000040093** | FOXI3 | -1.03 | 1.70E-02 | 741.57 | 570.58 | 1046.10 | 870.77 | 2966.89 | 1032.34 | 1378.37 | 1191.55 |
| **ENSBTAG00000021250** | RALBP1 | -1.02 | 1.83E-10 | 2431.26 | 2273.19 | 2383.88 | 2280.79 | 4047.96 | 4885.38 | 4656.14 | 5464.37 |
| **ENSBTAG00000009874** | ZXDC | -1.02 | 3.22E-02 | 150.13 | 92.39 | 126.76 | 157.15 | 214.55 | 202.31 | 303.21 | 349.55 |
| **ENSBTAG00000015817** | ELK1 | -1.02 | 3.77E-02 | 441.32 | 316.76 | 371.67 | 428.39 | 1487.26 | 326.00 | 706.61 | 642.65 |
| **ENSBTAG00000046218** | KLF11 | -1.02 | 8.86E-03 | 577.34 | 456.87 | 302.75 | 490.82 | 843.62 | 1057.77 | 809.42 | 1003.12 |
| **ENSBTAG00000021151** | MYH10 | -1.02 | 4.43E-02 | 449.38 | 341.13 | 369.21 | 440.23 | 793.62 | 1034.65 | 546.29 | 874.77 |
| **ENSBTAG00000043981** | SBF2 | -1.02 | 4.00E-02 | 239.80 | 181.73 | 241.22 | 134.54 | 273.57 | 499.41 | 419.09 | 426.92 |
| **ENSBTAG00000008436** | CDC25B | -1.02 | 7.70E-05 | 1634.28 | 747.24 | 1532.23 | 1125.87 | 2706.51 | 2373.33 | 2621.69 | 2514.18 |
| **ENSBTAG00000007112** | HSPB11 | -1.02 | 4.90E-03 | 151.14 | 294.43 | 145.22 | 232.49 | 254.82 | 551.43 | 467.01 | 399.61 |
| **ENSBTAG00000006609** | ERRFI1 | -1.02 | 5.89E-05 | 1475.08 | 2182.83 | 2586.94 | 1984.80 | 5242.91 | 3379.08 | 3779.63 | 4239.15 |

| | | | | | | | | | | | |
|---|---|---|---|---|---|---|---|---|---|---|---|
| **ENSBTAG00000004991** | ELP1 | -1.02 | 4.74E-02 | 372.80 | 414.23 | 374.13 | 122.70 | 1046.36 | 525.99 | 444.35 | 577.11 |
| **ENSBTAG00000018583** | | -1.01 | 4.53E-02 | 119.90 | 54.82 | 49.23 | 105.48 | 161.09 | 195.37 | 161.19 | 149.28 |
| **ENSBTAG00000009314** | FYB2 | -1.01 | 1.89E-02 | 142.07 | 120.82 | 98.46 | 73.19 | 195.11 | 253.17 | 165.54 | 263.98 |
| **ENSBTAG00000020495** | SH3YL1 | -1.01 | 1.90E-02 | 110.83 | 162.44 | 223.99 | 129.16 | 188.16 | 293.63 | 349.38 | 428.74 |
| **ENSBTAG00000002116** | ZCCHC3 | -1.01 | 1.26E-04 | 1788.44 | 1203.10 | 2435.57 | 1509.05 | 4375.00 | 3205.67 | 3521.73 | 2839.14 |
| **ENSBTAG00000018691** | RHOU | -1.01 | 4.69E-02 | 204.54 | 221.33 | 343.37 | 71.04 | 513.81 | 520.21 | 329.34 | 323.15 |
| **ENSBTAG00000007118** | SMURF1 | -1.01 | 5.44E-04 | 283.13 | 219.30 | 478.74 | 365.96 | 867.92 | 618.48 | 609.90 | 603.51 |
| **ENSBTAG00000019044** | BAIAP2 | -1.00 | 1.35E-02 | 975.33 | 737.09 | 1053.48 | 769.59 | 1874.01 | 1285.51 | 2174.72 | 1755.01 |
| **ENSBTAG00000013645** | RFFL | -1.00 | 3.82E-02 | 189.42 | 80.21 | 119.38 | 47.36 | 169.42 | 202.31 | 258.77 | 244.86 |
| **ENSBTAG00000000919** | HEY2 | -1.00 | 1.99E-02 | 184.39 | 238.59 | 241.22 | 190.51 | 436.74 | 301.72 | 466.14 | 506.11 |
| **ENSBTAG00000009252** | KLRA1 | 1.00 | 3.11E-02 | 498.75 | 278.18 | 347.06 | 381.03 | 193.02 | 99.42 | 196.91 | 260.34 |
| **ENSBTAG00000015147** | S100A10 | 1.01 | 7.54E-03 | 10109.95 | 20398.81 | 6871.03 | 14403.76 | 3688.99 | 8547.69 | 6917.99 | 6569.45 |
| **ENSBTAG00000032775** | | 1.02 | 4.23E-02 | 69.52 | 163.46 | 83.69 | 85.03 | 34.02 | 45.09 | 49.66 | 69.18 |
| **ENSBTAG00000011334** | NADK2 | 1.04 | 2.59E-03 | 514.87 | 530.99 | 292.91 | 525.26 | 209.69 | 273.98 | 227.40 | 199.35 |
| **ENSBTAG00000003027** | EMX2 | 1.04 | 1.57E-04 | 356.68 | 535.05 | 431.98 | 487.59 | 191.64 | 208.09 | 184.71 | 295.84 |
| **ENSBTAG00000002166** | CRISP2 | 1.05 | 1.30E-02 | 1128.48 | 1002.07 | 1423.93 | 1541.34 | 663.78 | 778.01 | 710.10 | 309.49 |
| **ENSBTAG00000051295** | FXYD4 | 1.07 | 7.56E-03 | 1650.40 | 4116.92 | 2218.96 | 3113.89 | 976.93 | 2062.36 | 1205.86 | 1058.65 |
| **ENSBTAG00000001595** | | 1.07 | 2.11E-02 | 1769.29 | 1478.23 | 1517.46 | 2036.46 | 410.35 | 910.95 | 1144.87 | 766.45 |
| **ENSBTAG00000020105** | | 1.07 | 5.98E-04 | 1128.48 | 1206.14 | 990.72 | 1622.06 | 475.62 | 596.51 | 514.93 | 763.72 |
| **ENSBTAG00000005085** | TRIM63 | 1.07 | 1.25E-04 | 1448.88 | 1806.17 | 1499.00 | 1969.73 | 586.71 | 1277.41 | 600.31 | 732.77 |
| **ENSBTAG00000021292** | ANKFN1 | 1.09 | 2.74E-02 | 743.59 | 631.50 | 486.13 | 696.40 | 178.44 | 485.53 | 227.40 | 312.22 |
| **ENSBTAG00000033308** | | 1.10 | 3.26E-03 | 100.76 | 180.72 | 166.15 | 175.45 | 65.96 | 87.86 | 64.47 | 73.73 |
| **ENSBTAG00000050728** | | 1.11 | 4.52E-02 | 100.76 | 124.88 | 109.53 | 89.34 | 68.74 | 53.18 | 37.47 | 37.32 |
| **ENSBTAG00000002041** | SASH3 | 1.11 | 2.78E-02 | 238.79 | 134.02 | 94.76 | 107.64 | 73.60 | 53.18 | 66.22 | 73.73 |
| **ENSBTAG00000020676** | MMP9 | 1.11 | 9.02E-06 | 453.41 | 483.27 | 492.28 | 420.85 | 223.58 | 262.42 | 199.52 | 173.86 |
| **ENSBTAG00000021098** | | 1.11 | 1.36E-02 | 147.11 | 100.51 | 147.68 | 146.38 | 65.27 | 48.55 | 55.76 | 80.10 |
| **ENSBTAG00000022564** | AKR1C4 | 1.15 | 2.10E-02 | 608.57 | 320.83 | 262.14 | 438.08 | 150.67 | 270.51 | 103.68 | 212.09 |
| **ENSBTAG00000032812** | | 1.17 | 1.09E-02 | 119.90 | 98.48 | 110.76 | 83.96 | 31.25 | 63.58 | 46.18 | 43.69 |
| **ENSBTAG00000020550** | | 1.18 | 2.30E-03 | 1905.31 | 718.81 | 734.73 | 1445.54 | 426.32 | 571.08 | 610.77 | 518.86 |
| **ENSBTAG00000032057** | | 1.18 | 1.91E-03 | 4968.32 | 3167.64 | 4047.79 | 4272.05 | 1090.80 | 2737.48 | 1599.68 | 1837.84 |
| **ENSBTAG00000044490** | RF01241 | 1.20 | 7.46E-03 | 355.67 | 290.37 | 205.53 | 306.76 | 92.35 | 110.98 | 131.56 | 170.22 |
| **ENSBTAG00000054661** | bta-mir-2887-2 | 1.20 | 1.69E-02 | 709.33 | 891.41 | 904.57 | 1045.14 | 270.10 | 331.78 | 416.47 | 526.14 |
| **ENSBTAG00000010368** | TPST2 | 1.21 | 1.95E-02 | 308.32 | 698.51 | 311.37 | 821.26 | 135.40 | 248.55 | 152.47 | 390.51 |

| | | | | | | | | | | | |
|---|---|---|---|---|---|---|---|---|---|---|---|
| **ENSBTAG00000015828** | FKBP11 | 1.23 | 1.70E-04 | 3769.32 | 4441.81 | 3779.50 | 8066.19 | 1491.43 | 3031.11 | 1873.26 | 2152.80 |
| **ENSBTAG00000038706** | MT1E | 1.24 | 1.49E-02 | 788.93 | 307.63 | 500.90 | 757.75 | 136.09 | 255.48 | 358.10 | 247.59 |
| **ENSBTAG00000050377** | | 1.25 | 1.62E-02 | 123.93 | 368.54 | 175.99 | 503.73 | 88.87 | 158.38 | 131.56 | 113.78 |
| **ENSBTAG00000042256** | RF00090 | 1.28 | 2.88E-03 | 206.55 | 194.93 | 247.37 | 153.92 | 105.54 | 54.33 | 56.63 | 112.87 |
| **ENSBTAG00000043258** | RF00425 | 1.30 | 2.08E-02 | 219.65 | 157.37 | 158.76 | 128.09 | 49.99 | 49.71 | 64.47 | 105.59 |
| **ENSBTAG00000004950** | BRB | 1.34 | 2.75E-02 | 3892.24 | 9140.48 | 5711.71 | 7689.47 | 2728.04 | 4424.13 | 1477.70 | 1799.61 |
| **ENSBTAG00000033802** | | 1.34 | 1.91E-02 | 62.47 | 85.28 | 64.00 | 82.88 | 21.52 | 42.77 | 21.78 | 30.95 |
| **ENSBTAG00000015307** | FBN2 | 1.36 | 4.19E-03 | 209.57 | 255.85 | 551.36 | 207.74 | 127.76 | 144.50 | 101.07 | 103.77 |
| **ENSBTAG00000001043** | LUZP2 | 1.44 | 7.99E-04 | 189.42 | 454.84 | 559.97 | 698.55 | 103.46 | 159.53 | 179.48 | 259.43 |
| **ENSBTAG00000021118** | CYP26A1 | 1.45 | 8.71E-05 | 1445.86 | 1148.27 | 1479.31 | 1048.37 | 366.61 | 409.23 | 481.82 | 610.79 |
| **ENSBTAG00000006397** | | 1.46 | 2.89E-02 | 613.61 | 140.11 | 127.99 | 171.14 | 86.10 | 119.07 | 78.42 | 100.13 |
| **ENSBTAG00000004175** | HPD | 1.55 | 1.19E-02 | 76.58 | 96.45 | 93.53 | 137.77 | 25.00 | 24.28 | 59.25 | 29.13 |
| **ENSBTAG00000004347** | ADGRF5 | 1.62 | 1.05E-02 | 200.51 | 304.58 | 205.53 | 201.28 | 109.70 | 31.21 | 128.08 | 27.31 |
| **ENSBTAG00000043222** | RF00586 | 1.68 | 1.45E-06 | 228.72 | 235.54 | 226.45 | 178.67 | 79.85 | 40.46 | 67.96 | 81.01 |
| **ENSBTAG00000044441** | RF00553 | 1.68 | 2.40E-03 | 173.30 | 138.08 | 91.07 | 144.23 | 54.16 | 26.59 | 30.49 | 58.26 |
| **ENSBTAG00000000898** | F2RL2 | 1.77 | 2.72E-02 | 59.45 | 228.44 | 129.22 | 308.91 | 49.99 | 35.84 | 92.36 | 34.59 |
| **ENSBTAG00000004817** | | 2.00 | 7.20E-04 | 369.78 | 292.40 | 131.69 | 102.25 | 56.24 | 67.05 | 48.79 | 51.89 |
| **ENSBTAG00000055264** | | 2.01 | 1.76E-02 | 49.37 | 61.93 | 141.53 | 257.25 | 37.49 | 52.02 | 27.88 | 9.10 |
| **ENSBTAG00000005330** | KRTDAP | 2.02 | 1.54E-04 | 586.41 | 1600.07 | 493.51 | 1424.02 | 306.20 | 397.67 | 107.17 | 198.44 |
| **ENSBTAG00000050159** | | 2.18 | 3.88E-02 | 25.19 | 12.18 | 8.61 | 17.22 | 4.17 | 3.47 | 1.74 | 4.55 |
| **ENSBTAG00000017071** | C1QTNF3 | 2.29 | 3.87E-02 | 42.32 | 912.73 | 43.07 | 287.39 | 19.44 | 31.21 | 22.65 | 189.34 |
| **ENSBTAG00000021522** | PLA2G10 | 2.59 | 1.87E-02 | 32.24 | 40.61 | 41.84 | 52.74 | 3.47 | 0.00 | 10.46 | 13.65 |
| **ENSBTAG00000000620** | MCMDC2 | 3.13 | 3.40E-02 | 7.05 | 41.63 | 45.54 | 4.31 | 9.72 | 0.00 | 0.00 | 0.91 |
| **ENSBTAG00000021358** | BLNK | 3.41 | 6.12E-05 | 64.48 | 192.90 | 206.76 | 228.19 | 3.47 | 4.62 | 47.05 | 10.01 |
| **ENSBTAG00000050086** | | 4.31 | 1.93E-05 | 62.47 | 75.13 | 75.07 | 55.97 | 2.08 | 10.40 | 1.74 | 0.00 |
| **ENSBTAG00000054664** | | 5.35 | 2.40E-02 | 82.62 | 145.18 | 1.23 | 0.00 | 2.78 | 0.00 | 0.00 | 2.73 |
| **ENSBTAG00000038496** | CR2 | 6.21 | 4.31E-02 | 41.31 | 7.11 | 0.00 | 0.00 | 0.00 | 0.00 | 0.00 | 0.00 |
| **ENSBTAG00000045739** | OR5R1 | 6.38 | 1.42E-02 | 1.01 | 0.00 | 7.38 | 46.28 | 0.00 | 0.00 | 0.00 | 0.00 |
| **ENSBTAG00000012887** | FCER1A | 7.01 | 2.15E-02 | 0.00 | 42.64 | 41.84 | 0.00 | 0.00 | 0.00 | 0.00 | 0.00 |
| **ENSBTAG00000025952** | | 7.39 | 2.75E-03 | 79.60 | 0.00 | 0.00 | 102.25 | 0.00 | 1.16 | 0.00 | 0.00 |
| **ENSBTAG00000006432** | KCNE4 | 21.46 | 3.04E-05 | 0.00 | 73.10 | 3.69 | 0.00 | 0.00 | 0.00 | 0.00 | 0.00 |
