## Supplementary material 5: DEG_Comparison_HB_vs_SSB for "Bovine in vitro blastocysts with distinct morphokinetic patterns show transcriptomic differences at genome activation"

| DEG_comparison_HB_vs_SSB | | | | | | | | | | | |
|---|---|---|---|---|---|---|---|---|---|---|---|
| **Geneid** | **Gene name** | **log2FC** | **padj** | **HB-1** | **HB-2** | **HB-3** | **HB-4** | **SSB-1** | **SSB-2** | **SSB-3** | **SSB-4** |
| **ENSBTAG00000000707** | WISP1 | -32.77 | 9.74E-11 | 0.00 | 0.00 | 0.00 | 0.00 | 0.00 | 134.58 | 0.00 | 0.00 |
| **ENSBTAG00000022759** | PAG11 | -31.61 | 9.74E-11 | 0.00 | 0.00 | 0.00 | 0.00 | 0.00 | 74.32 | 0.00 | 0.00 |
| **ENSBTAG00000017294** | ORM1 | -25.66 | 7.41E-06 | 0.00 | 0.00 | 0.00 | 0.00 | 4.25 | 0.00 | 0.00 | 0.00 |
| **ENSBTAG00000007927** | SLAMF1 | -24.41 | 1.99E-05 | 0.00 | 0.00 | 0.00 | 0.00 | 68.79 | 0.00 | 0.00 | 0.00 |
| **ENSBTAG00000006432** | KCNE4 | -21.84 | 4.83E-04 | 0.00 | 0.00 | 0.00 | 0.00 | 0.00 | 111.48 | 0.00 | 0.00 |
| **ENSBTAG00000016444** | RETREG1 | -20.93 | 9.24E-04 | 0.00 | 0.00 | 0.00 | 0.00 | 25.48 | 32.14 | 0.00 | 0.00 |
| **ENSBTAG00000007596** | GEM | -20.87 | 2.82E-06 | 0.00 | 0.00 | 0.00 | 0.00 | 0.00 | 63.27 | 0.00 | 1.14 |
| **ENSBTAG00000019428** | CCR1 | -20.66 | 1.99E-05 | 0.00 | 0.00 | 0.00 | 0.00 | 24.63 | 27.12 | 0.00 | 0.00 |
| **ENSBTAG00000053807** | | -18.84 | 7.06E-03 | 0.00 | 0.00 | 0.00 | 0.00 | 0.00 | 13.06 | 0.00 | 0.00 |
| **ENSBTAG00000013339** | NEBL | -6.92 | 2.95E-04 | 0.00 | 0.00 | 0.00 | 0.00 | 39.07 | 5.02 | 28.87 | 11.41 |
| **ENSBTAG00000019859** | B3GALT2 | -6.91 | 3.54E-02 | 0.00 | 0.00 | 0.00 | 0.00 | 17.84 | 3.01 | 33.68 | 29.66 |
| **ENSBTAG00000003871** | CYP2B6 | -6.65 | 1.02E-02 | 8.94 | 1.62 | 1.62 | 0.00 | 9.34 | 1203.21 | 10.83 | 3.42 |
| **ENSBTAG00000015296** | PTPRB | -6.60 | 4.98E-04 | 0.00 | 0.00 | 1.62 | 0.00 | 18.68 | 73.32 | 60.14 | 28.52 |
| **ENSBTAG00000046155** | RGN | -6.23 | 4.43E-02 | 0.00 | 0.81 | 0.00 | 0.00 | 62.00 | 11.05 | 13.23 | 0.00 |
| **ENSBTAG00000025401** | DOK3 | -5.09 | 4.86E-02 | 0.00 | 0.00 | 0.00 | 2.50 | 11.04 | 26.11 | 10.83 | 18.25 |
| **ENSBTAG00000021696** | CCIN | -4.44 | 4.13E-02 | 4.97 | 0.00 | 0.00 | 0.00 | 44.16 | 26.11 | 19.25 | 15.97 |
| **ENSBTAG00000043512** | RF00263 | -2.07 | 4.13E-02 | 169.91 | 222.14 | 283.98 | 122.37 | 213.18 | 255.11 | 856.43 | 2035.29 |
| **ENSBTAG00000025441** | HSPA1A | -1.32 | 6.41E-04 | 2042.95 | 1438.25 | 3072.68 | 2195.21 | 5752.41 | 6104.44 | 6079.23 | 3909.72 |
| **ENSBTAG00000009284** | GLS2 | -1.11 | 4.07E-02 | 437.21 | 328.35 | 245.85 | 329.66 | 717.67 | 533.31 | 673.60 | 975.43 |
| **ENSBTAG00000040206** | ZNF770 | 1.26 | 7.06E-03 | 597.18 | 360.78 | 362.69 | 299.69 | 205.53 | 130.57 | 141.94 | 197.37 |
| **ENSBTAG00000032299** | CCDC172 | 1.58 | 4.98E-04 | 1120.84 | 734.53 | 777.30 | 1937.98 | 405.12 | 348.51 | 309.13 | 465.47 |
| **ENSBTAG00000011738** | TFR2 | 1.91 | 5.48E-03 | 57.63 | 54.32 | 59.23 | 63.68 | 22.08 | 15.07 | 7.22 | 17.11 |
| **ENSBTAG00000055051** | RNF125 | 3.51 | 4.43E-02 | 22.85 | 22.70 | 13.79 | 29.97 | 0.00 | 0.00 | 8.42 | 0.00 |
| **ENSBTAG00000007196** | TAGLN | 18.96 | 3.43E-03 | 0.00 | 15.40 | 0.00 | 3.75 | 0.00 | 0.00 | 0.00 | 0.00 |
| **ENSBTAG00000018563** | SFRP2 | 20.30 | 1.27E-07 | 24.84 | 8.92 | 17.04 | 0.00 | 0.00 | 0.00 | 0.00 | 0.00 |
