## Supplementary material 6: DEG_Comparison_HB_vs_ADB for "Bovine in vitro blastocysts with distinct morphokinetic patterns show transcriptomic differences at genome activation"

| DEG_comparison_HB_vs_ADB | | | | | | | | | | | |
|---|---|---|---|---|---|---|---|---|---|---|---|
| **Geneid** | **Gene name** | **log2FC** | **padj** | **HB-1** | **HB-2** | **HB-3** | **HB-4** | **ADB-1** | **ADB-2** | **ADB-3** | **ADB-4** |
| **ENSBTAG00000017294** | ORM1 | -32.00 | 6.42E-11 | 0.00 | 0.00 | 0.00 | 0.00 | 0.69 | 0.00 | 0.00 | 346.81 |
| **ENSBTAG00000000707** | WISP1 | -31.86 | 7.71E-11 | 0.00 | 0.00 | 0.00 | 0.00 | 0.00 | 0.00 | 0.00 | 71.91 |
| **ENSBTAG00000022759** | PAG11 | -28.05 | 1.05E-08 | 0.00 | 0.00 | 0.00 | 0.00 | 6.25 | 0.00 | 0.00 | 0.00 |
| **ENSBTAG00000004547** | OLR1 | -26.25 | 2.10E-07 | 0.00 | 0.00 | 0.00 | 0.00 | 0.00 | 0.00 | 32.24 | 2978.42 |
| **ENSBTAG00000053807** | | -22.44 | 2.03E-05 | 0.00 | 0.00 | 0.00 | 0.00 | 0.00 | 0.00 | 0.00 | 158.39 |
| **ENSBTAG00000007596** | GEM | -22.06 | 4.30E-08 | 0.00 | 0.00 | 0.00 | 0.00 | 9.72 | 0.00 | 7.84 | 129.26 |
| **ENSBTAG00000016444** | RETREG1 | -21.94 | 3.50E-05 | 0.00 | 0.00 | 0.00 | 0.00 | 0.00 | 12.72 | 0.00 | 103.77 |
| **ENSBTAG00000019428** | CCR1 | -20.98 | 1.42E-06 | 0.00 | 0.00 | 0.00 | 0.00 | 0.00 | 3.47 | 21.78 | 39.14 |
| **ENSBTAG00000007881** | IFIT1 | -20.76 | 1.08E-04 | 0.00 | 0.00 | 0.00 | 0.00 | 7.64 | 0.00 | 0.00 | 46.42 |
| **ENSBTAG00000013339** | NEBL | -8.11 | 2.07E-07 | 0.00 | 0.00 | 0.00 | 0.00 | 90.26 | 21.96 | 31.37 | 49.15 |
| **ENSBTAG00000033352** | DMRT1 | -7.94 | 7.02E-04 | 0.00 | 0.00 | 0.00 | 0.00 | 76.38 | 12.72 | 15.68 | 67.36 |
| **ENSBTAG00000017348** | FGF5 | -7.92 | 1.39E-03 | 0.00 | 0.00 | 0.00 | 0.00 | 27.08 | 91.33 | 11.33 | 40.05 |
| **ENSBTAG00000016645** | GABRA4 | -7.64 | 2.09E-02 | 0.00 | 0.00 | 0.00 | 0.00 | 54.85 | 0.00 | 46.18 | 38.23 |
| **ENSBTAG00000006039** | ARHGDIB | -7.57 | 4.40E-05 | 0.00 | 0.00 | 0.00 | 0.00 | 20.14 | 50.87 | 18.30 | 43.69 |
| **ENSBTAG00000037844** | CDH18 | -7.53 | 1.12E-03 | 0.00 | 0.00 | 0.00 | 0.00 | 4.86 | 41.62 | 23.52 | 59.17 |
| **ENSBTAG00000004813** | SUSD1 | -7.43 | 2.89E-03 | 0.00 | 0.00 | 0.00 | 0.00 | 1.39 | 24.28 | 48.79 | 46.42 |
| **ENSBTAG00000046771** | FNDC7 | -7.38 | 4.26E-03 | 0.00 | 0.00 | 0.00 | 0.00 | 4.17 | 3.47 | 11.33 | 97.40 |
| **ENSBTAG00000033748** | IL18RAP | -7.36 | 5.32E-03 | 0.00 | 0.00 | 0.00 | 0.00 | 6.94 | 24.28 | 57.50 | 26.40 |
| **ENSBTAG00000005015** | SFXN3 | -7.28 | 1.27E-02 | 0.00 | 0.00 | 0.00 | 0.00 | 23.61 | 1.16 | 0.00 | 83.75 |
| **ENSBTAG00000006141** | WDR93 | -7.27 | 3.38E-02 | 0.00 | 0.00 | 0.00 | 0.00 | 11.11 | 0.00 | 55.76 | 40.96 |
| **ENSBTAG00000046155** | RGN | -7.26 | 4.23E-04 | 0.00 | 0.81 | 0.00 | 0.00 | 0.69 | 25.43 | 19.17 | 131.08 |
| **ENSBTAG00000050057** | | -7.18 | 5.81E-03 | 0.00 | 0.00 | 0.00 | 0.00 | 2.78 | 27.74 | 23.52 | 47.33 |
| **ENSBTAG00000032858** | DYDC1 | -7.08 | 4.91E-03 | 0.00 | 0.00 | 0.00 | 0.00 | 4.17 | 17.34 | 57.50 | 15.47 |
| **ENSBTAG00000011458** | CPXM1 | -6.98 | 4.43E-02 | 0.00 | 0.00 | 0.00 | 0.00 | 1.39 | 1.16 | 0.00 | 85.57 |
| **ENSBTAG00000021730** | GJD2 | -6.96 | 1.07E-03 | 0.00 | 0.00 | 0.00 | 0.00 | 23.61 | 24.28 | 13.07 | 26.40 |
| **ENSBTAG00000047595** | FSTL5 | -6.92 | 8.48E-03 | 0.00 | 0.00 | 0.00 | 0.00 | 8.33 | 16.18 | 17.43 | 42.78 |
| **ENSBTAG00000054469** | SLC25A53 | -6.90 | 4.14E-02 | 0.00 | 0.00 | 0.00 | 0.00 | 45.83 | 0.00 | 12.20 | 25.49 |
| **ENSBTAG00000018707** | LDB3 | -6.83 | 3.49E-02 | 0.00 | 0.00 | 0.00 | 0.00 | 1.39 | 8.09 | 38.34 | 31.86 |
| **ENSBTAG00000003959** | ARHGAP24 | -6.82 | 1.86E-02 | 0.00 | 0.00 | 0.00 | 0.00 | 47.21 | 21.96 | 1.74 | 8.19 |
| **ENSBTAG00000012652** | TINAG | -6.68 | 4.73E-02 | 0.00 | 0.00 | 0.00 | 0.00 | 4.86 | 55.49 | 11.33 | 0.00 |
| **ENSBTAG00000026139** | ARMCX1 | -6.63 | 3.05E-02 | 0.00 | 0.00 | 0.00 | 0.00 | 11.80 | 21.96 | 0.00 | 35.50 |

| | | | | | | | | | | | |
|---|---|---|---|---|---|---|---|---|---|---|---|
| **ENSBTAG00000012706** | RTP1 | -6.62 | 6.61E-03 | 0.00 | 0.00 | 0.00 | 0.00 | 11.11 | 10.40 | 24.40 | 22.76 |
| **ENSBTAG00000019060** | CLDN11 | -6.47 | 4.46E-03 | 0.00 | 0.00 | 0.00 | 0.00 | 0.69 | 26.59 | 32.24 | 2.73 |
| **ENSBTAG00000019504** | ADRA1D | -6.45 | 3.01E-02 | 0.00 | 0.00 | 0.00 | 0.00 | 20.14 | 3.47 | 20.04 | 17.30 |
| **ENSBTAG00000018397** | HHLA2 | -6.42 | 4.17E-02 | 0.00 | 0.00 | 0.00 | 0.00 | 4.17 | 27.74 | 0.00 | 28.22 |
| **ENSBTAG00000012628** | SLC1A2 | -6.26 | 2.12E-02 | 0.00 | 0.00 | 0.00 | 0.00 | 9.72 | 23.12 | 20.04 | 0.91 |
| **ENSBTAG00000047423** | ERICH4 | -6.25 | 3.55E-02 | 0.00 | 0.00 | 0.00 | 0.00 | 10.42 | 21.96 | 20.04 | 0.91 |
| **ENSBTAG00000013898** | NPB | -6.23 | 3.62E-02 | 0.99 | 0.00 | 0.00 | 0.00 | 2.08 | 0.00 | 0.00 | 84.66 |
| **ENSBTAG00000020755** | SELP | -6.21 | 1.04E-02 | 0.00 | 0.81 | 0.81 | 0.00 | 26.38 | 16.18 | 47.05 | 46.42 |
| **ENSBTAG00000004680** | SLC13A5 | -6.20 | 3.92E-02 | 0.00 | 0.00 | 0.00 | 0.00 | 13.89 | 18.50 | 19.17 | 0.00 |
| **ENSBTAG00000014315** | PBLD | -6.19 | 4.07E-02 | 0.00 | 0.00 | 0.00 | 0.00 | 38.19 | 0.00 | 0.00 | 12.74 |
| **ENSBTAG00000016026** | PCOLCE2 | -6.15 | 3.22E-02 | 0.00 | 0.00 | 0.00 | 1.25 | 22.91 | 0.00 | 0.00 | 59.17 |
| **ENSBTAG00000010082** | COL15A1 | -6.11 | 4.93E-02 | 0.00 | 0.00 | 0.81 | 1.25 | 26.38 | 0.00 | 0.00 | 105.59 |
| **ENSBTAG00000049205** | | -5.96 | 1.68E-02 | 0.00 | 0.00 | 0.00 | 0.00 | 13.19 | 17.34 | 13.07 | 0.00 |
| **ENSBTAG00000050721** | | -5.94 | 3.06E-02 | 0.00 | 0.81 | 0.81 | 0.00 | 3.47 | 41.62 | 56.63 | 10.92 |
| **ENSBTAG00000027126** | MUC15 | -5.91 | 1.94E-04 | 0.00 | 0.00 | 3.25 | 0.00 | 42.35 | 75.14 | 67.96 | 26.40 |
| **ENSBTAG00000032880** | PRM2 | -5.85 | 2.37E-02 | 0.00 | 0.00 | 0.00 | 0.00 | 3.47 | 30.06 | 1.74 | 5.46 |
| **ENSBTAG00000054931** | CACNG4 | -5.82 | 2.86E-02 | 0.00 | 7.30 | 1.62 | 0.00 | 61.10 | 0.00 | 66.22 | 385.05 |
| **ENSBTAG00000010145** | SLC12A1 | -5.80 | 3.31E-02 | 0.99 | 0.00 | 0.00 | 0.00 | 21.52 | 21.96 | 5.23 | 15.47 |
| **ENSBTAG00000049009** | | -5.73 | 4.13E-02 | 0.00 | 0.81 | 0.00 | 0.00 | 20.14 | 1.16 | 27.88 | 11.83 |
| **ENSBTAG00000020836** | FAM92B | -5.62 | 4.67E-02 | 0.00 | 0.00 | 0.00 | 0.00 | 0.00 | 18.50 | 6.10 | 10.01 |
| **ENSBTAG00000021696** | CCIN | -5.51 | 1.29E-04 | 4.97 | 0.00 | 0.00 | 0.00 | 34.72 | 58.96 | 85.39 | 41.87 |
| **ENSBTAG00000021132** | SYNPO2L | -5.50 | 4.36E-02 | 0.00 | 0.00 | 0.00 | 0.00 | 7.64 | 6.94 | 3.49 | 13.65 |
| **ENSBTAG00000048460** | | -5.48 | 3.25E-03 | 0.00 | 0.00 | 0.00 | 3.75 | 13.19 | 33.52 | 34.85 | 64.63 |
| **ENSBTAG00000046630** | | -5.43 | 1.96E-02 | 0.00 | 4.86 | 4.06 | 64.93 | 261.07 | 4.62 | 518.41 | 2394.93 |
| **ENSBTAG00000008380** | ITGA11 | -5.40 | 4.41E-02 | 0.99 | 0.00 | 0.00 | 0.00 | 29.86 | 1.16 | 13.07 | 4.55 |
| **ENSBTAG00000027442** | NFIB | -5.26 | 1.38E-02 | 0.00 | 1.62 | 2.43 | 0.00 | 40.97 | 85.55 | 25.27 | 11.83 |
| **ENSBTAG00000034693** | SYT1 | -5.16 | 3.59E-02 | 2.98 | 0.00 | 0.00 | 0.00 | 0.00 | 16.18 | 26.14 | 62.81 |
| **ENSBTAG00000002199** | CORIN | -5.15 | 1.42E-03 | 2.98 | 0.81 | 0.00 | 0.00 | 13.19 | 11.56 | 26.14 | 83.75 |
| **ENSBTAG00000000246** | ME3 | -5.14 | 1.71E-02 | 0.00 | 0.00 | 0.00 | 1.25 | 9.72 | 3.47 | 17.43 | 10.01 |
| **ENSBTAG00000049646** | | -5.12 | 2.36E-02 | 0.99 | 8.11 | 0.00 | 18.73 | 256.21 | 17.34 | 292.75 | 394.15 |
| **ENSBTAG00000019859** | B3GALT2 | -5.11 | 3.64E-02 | 0.00 | 0.00 | 0.00 | 0.00 | 14.58 | 1.16 | 0.00 | 8.19 |
| **ENSBTAG00000051262** | | -5.07 | 2.96E-02 | 0.00 | 3.24 | 0.00 | 0.00 | 14.58 | 46.24 | 27.88 | 26.40 |
| **ENSBTAG00000003472** | ABCA12 | -4.92 | 3.93E-02 | 0.00 | 1.62 | 1.62 | 0.00 | 19.44 | 1.16 | 81.03 | 1.82 |

| | | | | | | | | | | | |
|---|---|---|---|---|---|---|---|---|---|---|---|
| **ENSBTAG00000020984** | RAPGEF4 | -4.88 | 1.25E-03 | 2.98 | 0.00 | 8.11 | 0.00 | 78.46 | 137.57 | 35.72 | 82.83 |
| **ENSBTAG00000025401** | DOK3 | -4.86 | 5.97E-03 | 0.00 | 0.00 | 0.00 | 2.50 | 14.58 | 3.47 | 5.23 | 32.77 |
| **ENSBTAG00000053102** | | -4.80 | 2.27E-04 | 6.96 | 12.97 | 3.25 | 9.99 | 180.53 | 21.96 | 252.67 | 469.70 |
| **ENSBTAG00000019146** | OSBP2 | -4.73 | 4.42E-02 | 0.00 | 0.81 | 0.81 | 1.25 | 2.78 | 5.78 | 13.07 | 53.71 |
| **ENSBTAG00000047810** | CCDC96 | -4.69 | 1.38E-02 | 0.00 | 0.00 | 0.81 | 3.75 | 35.41 | 47.40 | 16.55 | 9.10 |
| **ENSBTAG00000053475** | | -4.67 | 9.78E-03 | 32.79 | 326.73 | 48.68 | 38.71 | 3594.56 | 277.45 | 1800.94 | 5672.83 |
| **ENSBTAG00000009833** | DTHD1 | -4.65 | 4.41E-03 | 0.00 | 6.49 | 0.00 | 0.00 | 9.72 | 73.99 | 45.31 | 42.78 |
| **ENSBTAG00000039012** | MFSD6L | -4.56 | 2.43E-02 | 0.00 | 5.68 | 0.00 | 0.00 | 13.89 | 50.87 | 28.75 | 46.42 |
| **ENSBTAG00000054147** | | -4.50 | 4.57E-05 | 5.96 | 14.59 | 36.51 | 26.22 | 47.21 | 45.09 | 247.44 | 1541.09 |
| **ENSBTAG00000015296** | PTPRB | -4.49 | 1.01E-02 | 0.00 | 0.00 | 1.62 | 0.00 | 17.36 | 8.09 | 1.74 | 14.56 |
| **ENSBTAG00000033656** | | -4.44 | 8.99E-03 | 0.00 | 140.26 | 52.74 | 99.90 | 1746.94 | 249.70 | 1404.51 | 2978.42 |
| **ENSBTAG00000004608** | | -4.40 | 4.13E-02 | 0.00 | 0.00 | 1.62 | 0.00 | 5.55 | 23.12 | 6.10 | 4.55 |
| **ENSBTAG00000052032** | | -4.40 | 3.06E-02 | 0.00 | 0.81 | 3.25 | 0.00 | 6.94 | 36.99 | 17.43 | 29.13 |
| **ENSBTAG00000009197** | | -4.26 | 1.12E-03 | 86.45 | 919.38 | 238.54 | 258.48 | 9017.31 | 393.05 | 7050.42 | 12285.06 |
| **ENSBTAG00000015032** | CD14 | -4.17 | 1.71E-02 | 0.00 | 12.16 | 0.00 | 0.00 | 62.49 | 23.12 | 39.21 | 100.13 |
| **ENSBTAG00000013632** | GRM4 | -4.09 | 1.06E-03 | 2.98 | 5.68 | 0.00 | 2.50 | 45.13 | 39.31 | 67.09 | 38.23 |
| **ENSBTAG00000051910** | FGF10 | -4.06 | 4.90E-04 | 0.00 | 8.11 | 0.00 | 6.24 | 53.46 | 26.59 | 33.11 | 123.80 |
| **ENSBTAG00000046547** | CIDEA | -4.06 | 2.67E-02 | 0.00 | 0.00 | 15.42 | 4.99 | 40.27 | 110.98 | 64.47 | 125.62 |
| **ENSBTAG00000015415** | UNC80 | -4.04 | 1.12E-02 | 0.00 | 7.30 | 4.06 | 0.00 | 25.00 | 60.11 | 69.70 | 39.14 |
| **ENSBTAG00000047956** | NAGLU | -4.03 | 2.21E-02 | 0.00 | 4.86 | 0.00 | 11.24 | 62.49 | 61.27 | 80.16 | 53.71 |
| **ENSBTAG00000020516** | | -4.03 | 4.29E-02 | 0.00 | 0.00 | 0.00 | 4.99 | 5.55 | 16.18 | 27.01 | 24.58 |
| **ENSBTAG00000048635** | | -4.01 | 8.16E-04 | 0.99 | 1.62 | 0.81 | 4.99 | 13.19 | 26.59 | 42.69 | 45.51 |
| **ENSBTAG00000051436** | | -3.93 | 3.50E-02 | 0.00 | 3.24 | 0.00 | 11.24 | 20.83 | 16.18 | 128.95 | 48.24 |
| **ENSBTAG00000002261** | LBX1 | -3.93 | 2.37E-03 | 0.99 | 0.81 | 0.00 | 21.23 | 113.18 | 65.89 | 105.43 | 50.07 |
| **ENSBTAG00000024595** | HECA | -3.91 | 1.02E-02 | 6.96 | 0.81 | 0.81 | 3.75 | 36.11 | 2.31 | 36.59 | 106.50 |
| **ENSBTAG00000018549** | SPAG16 | -3.81 | 4.17E-02 | 0.00 | 2.43 | 0.00 | 3.75 | 11.11 | 9.25 | 46.18 | 17.30 |
| **ENSBTAG00000019421** | DACT1 | -3.71 | 3.65E-02 | 0.00 | 14.59 | 4.06 | 0.00 | 88.87 | 56.65 | 46.18 | 57.35 |
| **ENSBTAG00000049108** | TRAT1 | -3.70 | 1.37E-02 | 10.93 | 20.27 | 31.64 | 3.75 | 95.12 | 298.26 | 209.98 | 268.53 |
| **ENSBTAG00000049550** | | -3.68 | 2.94E-02 | 0.99 | 3.24 | 1.62 | 0.00 | 20.14 | 13.87 | 19.17 | 25.49 |
| **ENSBTAG00000053317** | YPEL4 | -3.68 | 4.13E-04 | 11.92 | 4.05 | 2.43 | 12.49 | 29.86 | 157.22 | 97.58 | 103.77 |
| **ENSBTAG00000012721** | HOGA1 | -3.65 | 3.30E-02 | 0.00 | 12.97 | 5.68 | 0.00 | 20.14 | 43.93 | 61.86 | 113.78 |
| **ENSBTAG00000048789** | NKX1-1 | -3.54 | 2.33E-03 | 8.94 | 2.43 | 4.06 | 1.25 | 27.77 | 69.36 | 47.92 | 50.07 |
| **ENSBTAG00000001004** | ESAM | -3.53 | 1.12E-02 | 1.99 | 5.68 | 0.00 | 3.75 | 27.08 | 15.03 | 74.06 | 14.56 |

| | | | | | | | | | | | |
|---|---|---|---|---|---|---|---|---|---|---|---|
| **ENSBTAG00000004769** | NEIL2 | -3.50 | 1.06E-04 | 7.95 | 47.83 | 20.28 | 16.23 | 61.80 | 197.68 | 362.45 | 426.92 |
| **ENSBTAG00000055266** | | -3.48 | 4.55E-02 | 10.93 | 4.86 | 4.06 | 0.00 | 7.64 | 9.25 | 48.79 | 156.57 |
| **ENSBTAG00000001021** | CYP1A1 | -3.46 | 3.66E-04 | 238.48 | 225.39 | 311.57 | 179.81 | 3716.07 | 2273.91 | 1401.89 | 3139.54 |
| **ENSBTAG00000021177** | ADAMTS14 | -3.45 | 1.20E-02 | 0.00 | 12.97 | 5.68 | 6.24 | 93.04 | 35.84 | 116.75 | 28.22 |
| **ENSBTAG00000031462** | FAM209A | -3.41 | 2.06E-02 | 1.99 | 0.00 | 6.49 | 2.50 | 32.63 | 10.40 | 52.28 | 21.85 |
| **ENSBTAG00000011733** | GIPC2 | -3.39 | 1.68E-10 | 22.85 | 83.51 | 43.00 | 104.89 | 510.34 | 1175.68 | 649.98 | 328.61 |
| **ENSBTAG00000009620** | WDR27 | -3.38 | 1.34E-02 | 0.00 | 9.73 | 0.00 | 4.99 | 8.33 | 36.99 | 54.02 | 54.62 |
| **ENSBTAG00000019734** | CD276 | -3.36 | 1.51E-02 | 0.99 | 17.84 | 5.68 | 0.00 | 45.83 | 95.95 | 42.69 | 73.73 |
| **ENSBTAG00000017420** | ZMAT4 | -3.31 | 2.00E-02 | 0.00 | 9.73 | 25.96 | 24.97 | 100.68 | 211.55 | 168.16 | 121.98 |
| **ENSBTAG00000005104** | MGAT5B | -3.29 | 5.60E-07 | 35.77 | 26.75 | 31.64 | 13.74 | 181.92 | 343.34 | 260.51 | 275.81 |
| **ENSBTAG00000018133** | SEMA3A | -3.29 | 3.16E-04 | 1.99 | 17.84 | 5.68 | 9.99 | 42.35 | 102.89 | 54.89 | 149.28 |
| **ENSBTAG00000016377** | IPP | -3.28 | 3.84E-04 | 14.90 | 64.86 | 65.72 | 12.49 | 213.85 | 534.09 | 461.78 | 334.98 |
| **ENSBTAG00000051889** | TDRD15 | -3.22 | 3.09E-02 | 10.93 | 17.84 | 2.43 | 9.99 | 69.43 | 141.04 | 44.44 | 130.17 |
| **ENSBTAG00000017024** | PPARGC1A | -3.21 | 7.60E-04 | 15.90 | 66.48 | 68.97 | 18.73 | 299.26 | 440.45 | 488.79 | 352.28 |
| **ENSBTAG00000022381** | TET3 | -3.15 | 3.90E-02 | 0.00 | 107.83 | 25.96 | 42.46 | 654.76 | 245.08 | 366.81 | 298.57 |
| **ENSBTAG00000010103** | TRIM9 | -3.13 | 3.72E-02 | 0.00 | 26.75 | 10.55 | 16.23 | 41.66 | 153.75 | 144.63 | 129.26 |
| **ENSBTAG00000027017** | SIX3 | -3.11 | 7.30E-04 | 2.98 | 129.72 | 40.57 | 9.99 | 349.25 | 279.76 | 426.93 | 539.79 |
| **ENSBTAG00000001110** | SLC16A5 | -3.11 | 1.25E-02 | 7.95 | 0.00 | 2.43 | 11.24 | 29.86 | 60.11 | 39.21 | 51.89 |
| **ENSBTAG00000011266** | ZBTB16 | -3.09 | 8.88E-04 | 0.99 | 153.23 | 12.17 | 71.18 | 401.32 | 537.55 | 534.97 | 546.16 |
| **ENSBTAG00000003103** | EIF4E1B | -3.08 | 1.20E-07 | 170.91 | 825.33 | 451.12 | 413.32 | 2664.85 | 4269.22 | 3690.75 | 5072.96 |
| **ENSBTAG00000012139** | SIX1 | -3.06 | 3.76E-03 | 7.95 | 99.72 | 56.80 | 32.47 | 331.89 | 284.38 | 473.98 | 558.00 |
| **ENSBTAG00000005301** | KLHL32 | -3.04 | 2.05E-03 | 13.91 | 16.21 | 24.34 | 7.49 | 95.82 | 137.57 | 129.82 | 151.11 |
| **ENSBTAG00000017251** | SLC26A8 | -3.02 | 9.90E-03 | 5.96 | 4.86 | 4.87 | 0.00 | 18.75 | 52.02 | 24.40 | 35.50 |
| **ENSBTAG00000033268** | ZFHX4 | -3.01 | 2.23E-02 | 0.99 | 4.05 | 21.91 | 2.50 | 64.57 | 8.09 | 54.89 | 113.78 |
| **ENSBTAG00000000016** | LTA | -3.01 | 1.91E-02 | 7.95 | 17.84 | 0.00 | 0.00 | 108.32 | 21.96 | 72.32 | 8.19 |
| **ENSBTAG00000000189** | LHFPL3 | -3.00 | 9.11E-03 | 23.85 | 228.63 | 84.38 | 127.37 | 691.56 | 1088.98 | 1043.80 | 895.71 |
| **ENSBTAG00000005259** | UCP3 | -3.00 | 4.72E-02 | 11.92 | 7.30 | 9.74 | 0.00 | 27.77 | 17.34 | 46.18 | 142.00 |
| **ENSBTAG00000011664** | ZIM2 | -2.96 | 2.88E-02 | 0.99 | 2.43 | 0.00 | 49.95 | 79.15 | 203.46 | 59.25 | 66.45 |
| **ENSBTAG00000004955** | SYTL5 | -2.95 | 2.09E-02 | 9.94 | 9.73 | 24.34 | 0.00 | 75.68 | 169.94 | 48.79 | 50.07 |
| **ENSBTAG00000005702** | CNGB1 | -2.95 | 3.32E-02 | 6.96 | 8.92 | 1.62 | 0.00 | 15.28 | 21.96 | 43.56 | 56.44 |
| **ENSBTAG00000024061** | NOTO | -2.95 | 3.26E-02 | 8.94 | 34.86 | 21.91 | 21.23 | 241.63 | 167.62 | 159.44 | 101.95 |
| **ENSBTAG00000014225** | SLC5A9 | -2.94 | 2.12E-02 | 0.99 | 87.56 | 17.85 | 9.99 | 100.68 | 216.18 | 302.34 | 281.27 |
| **ENSBTAG00000026836** | | -2.94 | 3.70E-02 | 16.89 | 23.51 | 21.91 | 23.73 | 103.46 | 260.11 | 189.94 | 106.50 |

| | | | | | | | | | | | |
|---|---|---|---|---|---|---|---|---|---|---|---|
| **ENSBTAG00000002526** | BDH2 | -2.91 | 1.19E-02 | 14.90 | 5.68 | 30.02 | 3.75 | 43.05 | 85.55 | 24.40 | 257.61 |
| **ENSBTAG00000053893** | | -2.87 | 1.06E-04 | 182.83 | 1543.65 | 487.64 | 690.53 | 5685.20 | 4277.31 | 6288.92 | 5036.55 |
| **ENSBTAG00000011274** | ENO2 | -2.87 | 4.99E-02 | 3.97 | 0.81 | 6.49 | 2.50 | 23.61 | 30.06 | 18.30 | 29.13 |
| **ENSBTAG00000026384** | ZAR1L | -2.85 | 1.54E-04 | 270.27 | 2194.67 | 1098.60 | 1595.83 | 4869.36 | 11211.18 | 10930.24 | 10103.13 |
| **ENSBTAG00000001652** | SLCO3A1 | -2.85 | 3.81E-02 | 6.96 | 24.32 | 34.08 | 18.73 | 157.61 | 213.87 | 76.67 | 157.48 |
| **ENSBTAG00000045526** | INSM1 | -2.83 | 3.44E-06 | 27.82 | 18.65 | 12.98 | 24.97 | 104.84 | 120.23 | 199.52 | 173.86 |
| **ENSBTAG00000001509** | ELK3 | -2.83 | 1.78E-04 | 20.87 | 50.27 | 8.93 | 22.48 | 128.45 | 250.86 | 204.75 | 149.28 |
| **ENSBTAG00000044273** | RF00026 | -2.82 | 1.37E-02 | 0.00 | 2.43 | 9.74 | 11.24 | 24.30 | 47.40 | 51.41 | 40.96 |
| **ENSBTAG00000018784** | CTSZ | -2.82 | 2.62E-03 | 48.69 | 205.93 | 160.65 | 58.69 | 543.66 | 440.45 | 622.10 | 1745.00 |
| **ENSBTAG00000019979** | CMPK2 | -2.82 | 4.25E-03 | 0.99 | 12.97 | 30.83 | 11.24 | 187.47 | 93.64 | 39.21 | 77.37 |
| **ENSBTAG00000053831** | | -2.78 | 3.22E-04 | 23.85 | 114.31 | 62.48 | 44.95 | 120.12 | 426.58 | 676.12 | 473.34 |
| **ENSBTAG00000006722** | FGF16 | -2.77 | 1.07E-03 | 3.97 | 136.20 | 63.29 | 58.69 | 351.33 | 508.65 | 399.05 | 532.51 |
| **ENSBTAG00000006727** | | -2.77 | 2.01E-02 | 2.98 | 10.54 | 8.93 | 4.99 | 21.52 | 76.30 | 27.88 | 63.72 |
| **ENSBTAG00000000623** | CPEB1 | -2.75 | 2.07E-08 | 1424.90 | 5010.37 | 2430.07 | 2541.10 | 11675.22 | 22879.00 | 22287.42 | 20046.96 |
| **ENSBTAG00000019872** | PIK3AP1 | -2.74 | 4.97E-02 | 0.00 | 1.62 | 15.42 | 14.98 | 63.88 | 36.99 | 80.16 | 31.86 |
| **ENSBTAG00000007746** | AHR | -2.74 | 9.02E-06 | 55.64 | 42.16 | 38.95 | 77.42 | 347.17 | 401.14 | 345.03 | 334.07 |
| **ENSBTAG00000015952** | ADAP1 | -2.73 | 3.26E-12 | 90.42 | 135.39 | 38.95 | 57.44 | 563.80 | 463.57 | 568.08 | 544.34 |
| **ENSBTAG00000001322** | SAXO1 | -2.72 | 3.58E-05 | 294.12 | 2109.55 | 551.74 | 949.01 | 4121.56 | 8569.65 | 5602.35 | 7514.31 |
| **ENSBTAG00000040061** | ZBTB38 | -2.71 | 1.39E-02 | 0.00 | 68.91 | 63.29 | 9.99 | 152.75 | 105.20 | 215.21 | 464.24 |
| **ENSBTAG00000048673** | | -2.71 | 1.38E-02 | 2.98 | 13.78 | 13.79 | 6.24 | 35.41 | 77.45 | 37.47 | 93.76 |
| **ENSBTAG00000017679** | PAK5 | -2.71 | 9.10E-05 | 33.78 | 211.60 | 42.19 | 54.94 | 585.32 | 502.87 | 385.11 | 774.64 |
| **ENSBTAG00000004613** | CSGALNACT1 | -2.71 | 2.24E-03 | 19.87 | 5.68 | 4.87 | 34.96 | 65.27 | 141.04 | 99.33 | 113.78 |
| **ENSBTAG00000045782** | BMP15 | -2.70 | 1.57E-03 | 386.53 | 2820.56 | 1389.08 | 891.57 | 8818.04 | 9617.02 | 9364.55 | 7907.55 |
| **ENSBTAG00000000502** | DAZL | -2.66 | 9.53E-03 | 7.95 | 119.99 | 55.17 | 26.22 | 227.05 | 347.97 | 293.62 | 462.42 |
| **ENSBTAG00000032517** | BCL7A | -2.66 | 2.30E-02 | 3.97 | 13.78 | 12.17 | 18.73 | 68.04 | 72.83 | 78.42 | 87.39 |
| **ENSBTAG00000008091** | SELENBP1 | -2.65 | 5.94E-08 | 36.77 | 120.80 | 107.10 | 99.90 | 412.43 | 582.64 | 633.42 | 656.31 |
| **ENSBTAG00000018438** | RRAGD | -2.64 | 2.93E-03 | 2.98 | 205.12 | 125.76 | 23.73 | 343.00 | 499.41 | 758.02 | 634.46 |
| **ENSBTAG00000026307** | ZNF629 | -2.61 | 1.67E-02 | 22.85 | 4.86 | 2.43 | 9.99 | 65.27 | 30.06 | 73.19 | 74.64 |
| **ENSBTAG00000019580** | TCL1A | -2.61 | 1.27E-03 | 9.94 | 89.99 | 23.53 | 21.23 | 121.51 | 210.40 | 158.57 | 395.97 |
| **ENSBTAG00000005567** | BEND4 | -2.60 | 3.40E-03 | 8.94 | 35.67 | 10.55 | 4.99 | 193.02 | 53.18 | 67.96 | 55.53 |
| **ENSBTAG00000054056** | | -2.59 | 3.53E-03 | 14.90 | 23.51 | 4.06 | 19.98 | 49.30 | 90.17 | 134.18 | 101.95 |
| **ENSBTAG00000024648** | PGR | -2.59 | 2.11E-02 | 4.97 | 19.46 | 18.66 | 38.71 | 79.15 | 26.59 | 166.42 | 218.47 |
| **ENSBTAG00000017969** | CA4 | -2.59 | 2.23E-02 | 39.75 | 7.30 | 3.25 | 3.75 | 69.43 | 50.87 | 182.97 | 21.85 |

| | | | | | | | | | | | |
|---|---|---|---|---|---|---|---|---|---|---|---|
| **ENSBTAG00000026972** | MYF5 | -2.59 | 6.84E-08 | 395.47 | 1306.91 | 477.90 | 349.64 | 1991.35 | 4783.65 | 4497.56 | 3931.47 |
| **ENSBTAG00000047658** | PHF7 | -2.56 | 3.02E-04 | 35.77 | 46.21 | 16.23 | 26.22 | 190.94 | 124.85 | 197.78 | 222.11 |
| **ENSBTAG00000000534** | PNOC | -2.56 | 4.11E-02 | 10.93 | 22.70 | 4.06 | 0.00 | 52.77 | 55.49 | 50.53 | 66.45 |
| **ENSBTAG00000005644** | GALNT9 | -2.56 | 7.69E-03 | 9.94 | 4.05 | 11.36 | 2.50 | 28.47 | 32.37 | 74.06 | 30.95 |
| **ENSBTAG00000009478** | GDF9 | -2.56 | 3.52E-04 | 112.28 | 949.38 | 375.67 | 211.03 | 1496.98 | 3499.30 | 1931.63 | 2772.69 |
| **ENSBTAG00000020620** | RGS2 | -2.55 | 1.04E-02 | 23.85 | 304.84 | 156.60 | 89.91 | 484.64 | 1343.31 | 776.31 | 779.19 |
| **ENSBTAG00000027348** | OOSP2 | -2.54 | 5.92E-04 | 571.35 | 769.39 | 2000.04 | 1673.25 | 4768.68 | 11807.69 | 6643.53 | 5980.50 |
| **ENSBTAG00000034366** | RGS2 | -2.54 | 3.44E-03 | 559.43 | 4469.61 | 1626.00 | 2037.87 | 7097.48 | 17061.85 | 14700.28 | 11699.75 |
| **ENSBTAG00000018123** | FBLN5 | -2.53 | 2.26E-02 | 2.98 | 68.91 | 14.60 | 18.73 | 162.47 | 285.54 | 105.43 | 59.17 |
| **ENSBTAG00000013991** | NR2E1 | -2.53 | 2.32E-02 | 5.96 | 2.43 | 12.17 | 2.50 | 20.14 | 11.56 | 52.28 | 50.98 |
| **ENSBTAG00000046152** | MGAM | -2.53 | 4.41E-02 | 0.99 | 55.13 | 8.93 | 7.49 | 76.38 | 166.47 | 113.27 | 65.54 |
| **ENSBTAG00000012629** | ZNF362 | -2.53 | 1.29E-02 | 4.97 | 51.89 | 54.36 | 13.74 | 132.62 | 105.20 | 253.54 | 233.03 |
| **ENSBTAG00000033319** | CD200 | -2.52 | 5.69E-06 | 58.63 | 89.18 | 67.34 | 134.86 | 232.60 | 631.19 | 528.87 | 616.26 |
| **ENSBTAG00000013274** | CENPV | -2.52 | 2.70E-06 | 1079.11 | 2599.23 | 2207.75 | 1559.62 | 6068.47 | 14260.79 | 10654.92 | 11768.93 |
| **ENSBTAG00000052058** | | -2.52 | 2.60E-02 | 1.99 | 11.35 | 0.00 | 1.25 | 28.47 | 15.03 | 27.01 | 15.47 |
| **ENSBTAG00000023752** | | -2.51 | 6.52E-03 | 18.88 | 96.48 | 60.04 | 48.70 | 137.48 | 463.57 | 261.38 | 416.00 |
| **ENSBTAG00000006409** | LRRC34 | -2.51 | 2.37E-02 | 0.00 | 3.24 | 17.85 | 7.49 | 56.94 | 17.34 | 40.95 | 48.24 |
| **ENSBTAG00000030425** | ID3 | -2.50 | 1.85E-03 | 348.77 | 2855.43 | 1163.51 | 1336.11 | 3149.50 | 11844.69 | 7628.95 | 9749.03 |
| **ENSBTAG00000006689** | TECTB | -2.50 | 3.32E-02 | 58.63 | 282.95 | 130.63 | 133.61 | 592.96 | 1062.39 | 921.82 | 853.84 |
| **ENSBTAG00000046122** | NRARP | -2.50 | 3.26E-02 | 21.86 | 16.21 | 16.23 | 9.99 | 11.11 | 70.52 | 114.14 | 168.40 |
| **ENSBTAG00000015737** | KPNA7 | -2.49 | 4.39E-03 | 190.78 | 923.43 | 538.75 | 914.05 | 3187.69 | 4038.01 | 4175.19 | 3060.34 |
| **ENSBTAG00000018833** | SVOP | -2.49 | 4.65E-02 | 0.00 | 1.62 | 24.34 | 9.99 | 106.23 | 34.68 | 25.27 | 36.41 |
| **ENSBTAG00000047879** | KCNA3 | -2.49 | 3.18E-03 | 31.80 | 4.86 | 31.64 | 17.48 | 148.59 | 93.64 | 146.38 | 91.94 |
| **ENSBTAG00000002997** | ADAMTSL1 | -2.48 | 2.20E-02 | 0.00 | 125.66 | 27.59 | 21.23 | 272.18 | 156.06 | 278.81 | 271.26 |
| **ENSBTAG00000019675** | ATXN1 | -2.48 | 1.18E-04 | 16.89 | 91.61 | 35.70 | 58.69 | 222.88 | 379.18 | 224.79 | 307.67 |
| **ENSBTAG00000046425** | | -2.48 | 4.08E-02 | 89.43 | 391.59 | 317.25 | 129.86 | 1751.80 | 249.70 | 1523.00 | 1649.42 |
| **ENSBTAG00000005111** | BMP2 | -2.48 | 3.98E-02 | 76.51 | 179.98 | 43.00 | 29.97 | 218.72 | 279.76 | 159.44 | 1179.72 |
| **ENSBTAG00000037508** | EBF1 | -2.48 | 3.94E-04 | 66.57 | 121.61 | 69.78 | 41.21 | 300.65 | 396.52 | 476.59 | 496.10 |
| **ENSBTAG00000023377** | SH3BP5 | -2.47 | 2.26E-04 | 14.90 | 129.72 | 65.72 | 89.91 | 358.97 | 306.35 | 518.41 | 484.27 |
| **ENSBTAG00000018838** | IQCA1L | -2.47 | 2.00E-02 | 5.96 | 0.81 | 4.06 | 9.99 | 39.58 | 16.18 | 36.59 | 19.12 |
| **ENSBTAG00000001188** | ROM1 | -2.47 | 2.26E-02 | 6.96 | 8.11 | 2.43 | 6.24 | 15.28 | 24.28 | 23.52 | 68.27 |
| **ENSBTAG00000015981** | ETV1 | -2.47 | 4.71E-04 | 9.94 | 70.53 | 12.17 | 39.96 | 106.93 | 176.87 | 230.89 | 219.38 |
| **ENSBTAG00000008916** | CYFIP2 | -2.46 | 7.66E-04 | 29.81 | 248.90 | 42.19 | 111.13 | 585.32 | 575.70 | 491.40 | 727.31 |

| | | | | | | | | | | | |
|---|---|---|---|---|---|---|---|---|---|---|---|
| **ENSBTAG00000006005** | MPND | -2.44 | 9.05E-06 | 49.68 | 92.42 | 159.84 | 101.14 | 332.59 | 665.87 | 594.21 | 598.96 |
| **ENSBTAG00000052435** | | -2.44 | 2.44E-02 | 4.97 | 2.43 | 8.93 | 4.99 | 22.22 | 55.49 | 27.01 | 11.83 |
| **ENSBTAG00000053626** | | -2.44 | 4.81E-02 | 0.00 | 94.05 | 10.55 | 11.24 | 156.92 | 149.13 | 187.33 | 138.36 |
| **ENSBTAG00000040086** | SLC38A8 | -2.43 | 2.05E-02 | 5.96 | 20.27 | 17.04 | 4.99 | 63.88 | 77.45 | 59.25 | 62.81 |
| **ENSBTAG00000019940** | RASGRF1 | -2.42 | 4.95E-02 | 0.00 | 34.05 | 2.43 | 14.98 | 79.15 | 67.05 | 65.35 | 65.54 |
| **ENSBTAG00000016711** | PPIF | -2.42 | 1.36E-02 | 16.89 | 622.65 | 335.91 | 202.29 | 944.29 | 1569.89 | 1779.16 | 2018.08 |
| **ENSBTAG00000001812** | H1FOO | -2.42 | 4.10E-03 | 189.79 | 971.27 | 532.26 | 846.62 | 2125.35 | 4488.87 | 4000.06 | 2979.33 |
| **ENSBTAG00000007403** | SLC7A3 | -2.41 | 4.07E-04 | 64.59 | 304.84 | 97.37 | 132.36 | 583.24 | 1164.12 | 681.34 | 756.44 |
| **ENSBTAG00000019946** | USP44 | -2.39 | 1.15E-03 | 59.62 | 282.14 | 136.31 | 71.18 | 340.22 | 1273.95 | 557.62 | 710.01 |
| **ENSBTAG00000018600** | VGLL4 | -2.38 | 1.43E-10 | 136.13 | 414.29 | 241.79 | 112.38 | 1000.53 | 1308.63 | 1413.22 | 1008.58 |
| **ENSBTAG00000034346** | BTG4 | -2.38 | 1.36E-02 | 1330.50 | 16821.23 | 6067.47 | 4191.88 | 16773.01 | 58927.49 | 30122.86 | 42014.61 |
| **ENSBTAG00000050101** | NRXN3 | -2.38 | 1.75E-02 | 20.87 | 70.53 | 48.68 | 0.00 | 120.81 | 161.84 | 244.83 | 205.72 |
| **ENSBTAG00000012053** | ACCSL | -2.38 | 1.25E-03 | 301.08 | 1143.14 | 742.41 | 701.77 | 4680.50 | 3036.89 | 4655.26 | 2632.51 |
| **ENSBTAG00000020214** | LRRC8B | -2.38 | 2.78E-02 | 19.87 | 219.71 | 38.95 | 41.21 | 389.52 | 439.29 | 298.85 | 534.33 |
| **ENSBTAG00000025329** | IRF2BPL | -2.37 | 4.48E-02 | 8.94 | 4.05 | 12.17 | 7.49 | 19.44 | 32.37 | 38.34 | 79.19 |
| **ENSBTAG00000018753** | TMEM163 | -2.37 | 9.47E-05 | 72.54 | 424.02 | 185.81 | 197.29 | 700.58 | 1327.12 | 1201.50 | 1319.90 |
| **ENSBTAG00000006064** | C7H19orf57 | -2.37 | 3.11E-02 | 19.87 | 9.73 | 10.55 | 0.00 | 88.18 | 38.15 | 52.28 | 30.04 |
| **ENSBTAG00000019674** | CLIP3 | -2.37 | 1.84E-02 | 8.94 | 49.46 | 107.10 | 11.24 | 329.81 | 75.14 | 235.25 | 274.90 |
| **ENSBTAG00000007216** | DPF1 | -2.36 | 6.90E-03 | 58.63 | 47.02 | 17.85 | 44.95 | 222.88 | 194.21 | 263.13 | 182.05 |
| **ENSBTAG00000053612** | | -2.34 | 1.58E-03 | 53.66 | 85.94 | 103.04 | 62.43 | 335.36 | 84.39 | 449.58 | 679.06 |
| **ENSBTAG00000012413** | FGF12 | -2.34 | 4.28E-02 | 5.96 | 38.10 | 15.42 | 6.24 | 54.16 | 101.73 | 56.63 | 122.89 |
| **ENSBTAG00000004305** | RGS16 | -2.34 | 8.48E-03 | 293.13 | 1902.81 | 705.09 | 477.00 | 4198.63 | 5306.18 | 4454.87 | 3098.57 |
| **ENSBTAG00000006188** | USH2A | -2.33 | 4.28E-05 | 60.61 | 187.28 | 98.18 | 143.60 | 379.80 | 664.72 | 598.57 | 821.98 |
| **ENSBTAG00000054800** | HES6 | -2.32 | 8.41E-03 | 15.90 | 102.96 | 41.38 | 49.95 | 149.98 | 48.55 | 400.79 | 455.14 |
| **ENSBTAG00000031052** | CDK5R2 | -2.32 | 4.18E-03 | 16.89 | 102.96 | 40.57 | 56.19 | 228.44 | 216.18 | 311.92 | 328.61 |
| **ENSBTAG00000033015** | FAHD1 | -2.32 | 5.06E-04 | 25.83 | 28.38 | 38.13 | 52.45 | 58.32 | 201.15 | 136.79 | 323.15 |
| **ENSBTAG00000048279** | BCAR4 | -2.32 | 8.59E-06 | 69.56 | 445.10 | 165.52 | 189.80 | 601.99 | 1396.48 | 1165.78 | 1174.25 |
| **ENSBTAG00000006703** | PTGDR | -2.31 | 3.34E-03 | 40.74 | 72.97 | 39.76 | 77.42 | 155.53 | 104.04 | 557.62 | 329.52 |
| **ENSBTAG00000022580** | INKA2 | -2.31 | 3.47E-05 | 23.85 | 119.18 | 80.33 | 56.19 | 358.28 | 476.28 | 250.06 | 309.49 |
| **ENSBTAG00000014821** | SLC7A7 | -2.31 | 1.62E-02 | 62.60 | 18.65 | 2.43 | 73.67 | 154.14 | 278.60 | 236.12 | 104.68 |
| **ENSBTAG00000046199** | NRXN1 | -2.30 | 2.81E-03 | 68.56 | 242.41 | 50.31 | 31.22 | 506.17 | 483.22 | 482.69 | 466.97 |
| **ENSBTAG00000016462** | TCF4 | -2.30 | 1.35E-02 | 16.89 | 235.93 | 103.04 | 89.91 | 343.00 | 499.41 | 862.57 | 493.37 |
| **ENSBTAG00000051066** | ZNF71 | -2.30 | 2.83E-02 | 20.87 | 19.46 | 3.25 | 22.48 | 46.52 | 104.04 | 108.04 | 64.63 |

| | | | | | | | | | | | |
|---|---|---|---|---|---|---|---|---|---|---|---|
| **ENSBTAG00000002490** | CHPT1 | -2.29 | 1.09E-02 | 17.89 | 55.13 | 55.98 | 49.95 | 157.61 | 166.47 | 245.70 | 305.85 |
| **ENSBTAG00000004115** | MYLIP | -2.29 | 1.76E-12 | 404.42 | 787.23 | 351.33 | 555.67 | 1658.07 | 2705.11 | 3499.07 | 2381.28 |
| **ENSBTAG00000003102** | GCC1 | -2.29 | 1.25E-02 | 1.99 | 128.10 | 51.12 | 43.70 | 307.59 | 278.60 | 276.20 | 236.67 |
| **ENSBTAG00000053278** | | -2.28 | 1.53E-03 | 310.02 | 1777.14 | 703.46 | 1078.87 | 3188.38 | 5628.71 | 5698.19 | 4287.39 |
| **ENSBTAG00000002545** | | -2.28 | 2.88E-04 | 98.37 | 73.78 | 208.52 | 122.37 | 538.11 | 199.99 | 418.22 | 1286.22 |
| **ENSBTAG00000047294** | | -2.28 | 3.34E-02 | 53.66 | 150.80 | 0.00 | 66.18 | 335.36 | 365.31 | 281.42 | 328.61 |
| **ENSBTAG00000008921** | NEXN | -2.27 | 6.47E-04 | 51.67 | 271.60 | 100.61 | 46.20 | 340.22 | 566.45 | 764.99 | 598.05 |
| **ENSBTAG00000048053** | SLBP2 | -2.27 | 1.85E-03 | 316.97 | 1730.93 | 613.40 | 468.26 | 2447.53 | 4520.08 | 4087.19 | 3996.10 |
| **ENSBTAG00000037710** | | -2.25 | 2.09E-09 | 801.88 | 2029.28 | 933.08 | 1190.01 | 3646.64 | 6054.13 | 5509.12 | 8335.38 |
| **ENSBTAG00000009830** | PLEKHB1 | -2.24 | 1.56E-02 | 18.88 | 34.05 | 23.53 | 28.72 | 91.65 | 188.43 | 103.68 | 114.69 |
| **ENSBTAG00000016879** | KIAA1147 | -2.24 | 1.50E-02 | 30.80 | 41.35 | 25.96 | 42.46 | 177.06 | 143.35 | 209.98 | 133.81 |
| **ENSBTAG00000014543** | PLXNA4 | -2.24 | 5.78E-05 | 42.73 | 81.07 | 72.21 | 18.73 | 443.68 | 249.70 | 208.24 | 118.34 |
| **ENSBTAG00000047998** | COL5A1 | -2.24 | 4.68E-02 | 33.78 | 12.16 | 17.04 | 7.49 | 16.66 | 41.62 | 109.78 | 164.76 |
| **ENSBTAG00000003039** | PSMB8 | -2.23 | 4.44E-02 | 0.00 | 17.84 | 18.66 | 8.74 | 39.58 | 68.21 | 63.60 | 43.69 |
| **ENSBTAG00000018403** | ARHGAP9 | -2.23 | 2.63E-02 | 40.74 | 3.24 | 14.60 | 19.98 | 114.57 | 143.35 | 65.35 | 44.60 |
| **ENSBTAG00000021779** | MGST2 | -2.23 | 2.36E-03 | 35.77 | 542.39 | 81.95 | 53.69 | 658.23 | 1063.55 | 911.36 | 724.58 |
| **ENSBTAG00000048664** | CDK14 | -2.23 | 8.01E-03 | 6.96 | 52.70 | 27.59 | 33.71 | 68.74 | 182.65 | 140.28 | 177.50 |
| **ENSBTAG00000000357** | PRAG1 | -2.22 | 1.79E-03 | 69.56 | 68.10 | 33.27 | 38.71 | 302.73 | 234.67 | 224.79 | 215.73 |
| **ENSBTAG00000025760** | | -2.22 | 5.54E-03 | 65.58 | 248.09 | 114.40 | 71.18 | 399.94 | 623.10 | 616.87 | 685.44 |
| **ENSBTAG00000010875** | MSX1 | -2.21 | 2.09E-10 | 576.32 | 1566.35 | 902.25 | 1001.46 | 4063.24 | 5272.65 | 5053.44 | 4325.62 |
| **ENSBTAG00000040202** | | -2.21 | 1.14E-02 | 47.70 | 132.96 | 47.06 | 43.70 | 205.52 | 223.11 | 249.19 | 576.20 |
| **ENSBTAG00000037389** | TRIM44 | -2.20 | 1.80E-03 | 8.94 | 34.86 | 47.06 | 12.49 | 96.51 | 99.42 | 111.52 | 171.13 |
| **ENSBTAG00000053420** | | -2.19 | 4.93E-02 | 15.90 | 10.54 | 8.93 | 37.46 | 116.65 | 57.80 | 98.45 | 55.53 |
| **ENSBTAG00000017767** | GPR173 | -2.19 | 2.54E-03 | 83.47 | 196.20 | 17.85 | 37.46 | 377.02 | 320.22 | 436.51 | 395.06 |
| **ENSBTAG00000000245** | NHSL1 | -2.18 | 3.39E-03 | 158.98 | 1058.83 | 494.94 | 248.49 | 1671.96 | 2256.57 | 2907.47 | 2079.07 |
| **ENSBTAG00000008078** | TRIM59 | -2.18 | 3.74E-04 | 268.29 | 457.26 | 64.10 | 128.62 | 384.66 | 1780.29 | 750.17 | 1260.73 |
| **ENSBTAG00000018577** | SLC7A1 | -2.18 | 4.70E-03 | 24.84 | 37.29 | 25.15 | 3.75 | 127.76 | 113.29 | 99.33 | 76.46 |
| **ENSBTAG00000047600** | ROS1 | -2.18 | 4.37E-02 | 5.96 | 89.18 | 66.53 | 58.69 | 119.43 | 439.29 | 176.00 | 263.98 |
| **ENSBTAG00000048560** | TMEM233 | -2.18 | 1.15E-02 | 9.94 | 122.42 | 29.21 | 14.98 | 141.64 | 278.60 | 156.83 | 225.75 |
| **ENSBTAG00000024021** | NRXN1 | -2.17 | 1.28E-02 | 2.98 | 141.07 | 55.98 | 86.16 | 301.34 | 412.70 | 281.42 | 297.66 |
| **ENSBTAG00000005230** | SHOX2 | -2.17 | 7.63E-03 | 42.73 | 85.13 | 34.89 | 59.94 | 229.13 | 208.09 | 277.07 | 284.92 |
| **ENSBTAG00000036087** | ARMC2 | -2.17 | 4.41E-04 | 126.19 | 686.70 | 184.99 | 171.07 | 841.53 | 1205.74 | 1526.49 | 1676.72 |
| **ENSBTAG00000007778** | TRPM3 | -2.16 | 6.97E-03 | 20.87 | 178.36 | 83.57 | 18.73 | 169.42 | 419.64 | 345.90 | 420.55 |

| | | | | | | | | | | | |
|---|---|---|---|---|---|---|---|---|---|---|---|
| **ENSBTAG00000020548** | AZIN2 | -2.16 | 1.11E-02 | 46.70 | 60.81 | 32.46 | 39.96 | 193.02 | 268.20 | 210.85 | 131.08 |
| **ENSBTAG00000019009** | | -2.16 | 1.04E-02 | 53.66 | 51.89 | 162.28 | 112.38 | 401.32 | 362.99 | 549.78 | 381.40 |
| **ENSBTAG00000014903** | ABCA3 | -2.15 | 3.34E-02 | 13.91 | 20.27 | 25.15 | 4.99 | 77.77 | 75.14 | 89.74 | 46.42 |
| **ENSBTAG00000039015** | TMEM145 | -2.15 | 3.16E-02 | 17.89 | 90.80 | 12.98 | 63.68 | 180.53 | 235.83 | 314.53 | 89.21 |
| **ENSBTAG00000015525** | SLC39A12 | -2.14 | 3.03E-02 | 29.81 | 71.35 | 42.19 | 58.69 | 115.26 | 369.93 | 268.36 | 140.18 |
| **ENSBTAG00000016691** | LACC1 | -2.14 | 1.23E-03 | 27.82 | 82.70 | 81.14 | 12.49 | 154.14 | 215.02 | 349.38 | 190.25 |
| **ENSBTAG00000010351** | SNX25 | -2.12 | 5.81E-04 | 50.68 | 277.27 | 143.61 | 129.86 | 581.85 | 649.69 | 778.06 | 609.88 |
| **ENSBTAG00000007306** | RAB3C | -2.11 | 1.40E-04 | 33.78 | 95.67 | 114.40 | 141.10 | 231.91 | 544.49 | 321.50 | 568.01 |
| **ENSBTAG00000049459** | | -2.11 | 1.88E-02 | 5.96 | 34.86 | 12.17 | 7.49 | 40.27 | 86.70 | 81.03 | 57.35 |
| **ENSBTAG00000014699** | FAM81A | -2.11 | 5.62E-06 | 262.32 | 989.91 | 503.05 | 514.46 | 1845.54 | 2858.86 | 2628.66 | 2472.30 |
| **ENSBTAG00000027375** | DUSP22 | -2.11 | 4.10E-03 | 4.97 | 16.21 | 11.36 | 8.74 | 31.25 | 46.24 | 57.50 | 45.51 |
| **ENSBTAG00000010241** | UNC5D | -2.11 | 1.54E-04 | 21.86 | 17.84 | 17.85 | 32.47 | 80.54 | 105.20 | 72.32 | 126.53 |
| **ENSBTAG00000044125** | TLCD4 | -2.11 | 5.82E-03 | 42.73 | 379.43 | 162.28 | 78.67 | 373.55 | 1132.91 | 573.30 | 783.75 |
| **ENSBTAG00000002733** | DBX1 | -2.11 | 1.60E-04 | 146.07 | 389.16 | 90.87 | 203.54 | 798.48 | 765.29 | 1004.59 | 1007.67 |
| **ENSBTAG00000001704** | | -2.10 | 3.98E-02 | 6.96 | 5.68 | 7.30 | 19.98 | 36.11 | 25.43 | 53.15 | 53.71 |
| **ENSBTAG00000031246** | CCNI2 | -2.10 | 1.49E-04 | 80.49 | 265.92 | 63.29 | 101.14 | 377.72 | 649.69 | 639.52 | 529.78 |
| **ENSBTAG00000000418** | TOX3 | -2.08 | 2.71E-02 | 48.69 | 150.80 | 45.44 | 27.47 | 204.13 | 381.49 | 312.79 | 256.70 |
| **ENSBTAG00000007439** | TENM4 | -2.08 | 2.91E-06 | 44.71 | 40.54 | 33.27 | 56.19 | 193.72 | 153.75 | 212.59 | 173.86 |
| **ENSBTAG00000043996** | SAMD12 | -2.07 | 6.55E-03 | 24.84 | 293.49 | 89.25 | 112.38 | 229.13 | 633.50 | 598.57 | 728.22 |
| **ENSBTAG00000003212** | NNAT | -2.07 | 6.90E-03 | 37.76 | 28.38 | 21.91 | 21.23 | 135.40 | 69.36 | 113.27 | 141.09 |
| **ENSBTAG00000016381** | REM2 | -2.07 | 1.83E-02 | 10.93 | 70.53 | 34.89 | 24.97 | 95.12 | 129.48 | 116.75 | 254.88 |
| **ENSBTAG00000000830** | GPR137B | -2.07 | 3.44E-03 | 40.74 | 616.97 | 162.28 | 186.06 | 775.57 | 1102.85 | 1150.09 | 1195.19 |
| **ENSBTAG00000003827** | RIMS2 | -2.07 | 2.13E-03 | 48.69 | 534.28 | 165.52 | 320.92 | 731.83 | 1396.48 | 1247.68 | 1104.16 |
| **ENSBTAG00000046509** | TENT5C | -2.07 | 3.26E-12 | 179.85 | 378.62 | 344.83 | 274.71 | 1184.53 | 1019.62 | 1157.06 | 1577.50 |
| **ENSBTAG00000012653** | CAMK2B | -2.06 | 1.64E-03 | 7.95 | 11.35 | 14.60 | 12.49 | 52.77 | 33.52 | 54.02 | 52.80 |
| **ENSBTAG00000020458** | STX6 | -2.06 | 1.49E-03 | 23.85 | 140.26 | 47.87 | 94.90 | 337.45 | 304.04 | 337.19 | 297.66 |
| **ENSBTAG00000006194** | FOSL1 | -2.05 | 5.86E-03 | 71.54 | 14.59 | 47.06 | 36.21 | 147.20 | 55.49 | 287.52 | 212.09 |
| **ENSBTAG00000038600** | LRRC15 | -2.05 | 2.55E-02 | 32.79 | 57.56 | 7.30 | 24.97 | 149.98 | 89.01 | 90.61 | 179.32 |
| **ENSBTAG00000003669** | BNC2 | -2.05 | 4.87E-08 | 112.28 | 194.58 | 211.77 | 124.87 | 717.25 | 455.48 | 877.38 | 620.81 |
| **ENSBTAG00000034925** | KHDC3L | -2.04 | 2.83E-03 | 979.74 | 2901.64 | 3018.32 | 3019.35 | 4633.98 | 10497.91 | 10792.58 | 15008.60 |
| **ENSBTAG00000021497** | CDH23 | -2.04 | 4.09E-02 | 5.96 | 36.48 | 28.40 | 19.98 | 72.21 | 73.99 | 96.71 | 132.90 |
| **ENSBTAG00000002683** | PFKP | -2.04 | 4.99E-02 | 16.89 | 95.67 | 13.79 | 7.49 | 184.00 | 80.92 | 67.96 | 221.20 |
| **ENSBTAG00000001729** | DUSP10 | -2.04 | 1.12E-03 | 63.59 | 347.00 | 411.37 | 509.47 | 1251.88 | 1419.61 | 1614.49 | 1180.63 |

| | | | | | | | | | | | |
|---|---|---|---|---|---|---|---|---|---|---|---|
| **ENSBTAG00000052473** | CNTNAP2 | -2.03 | 9.00E-04 | 329.89 | 1692.83 | 870.61 | 674.30 | 2681.52 | 5034.51 | 2980.66 | 3898.70 |
| **ENSBTAG00000006734** | BICD1 | -2.02 | 2.21E-02 | 35.77 | 21.08 | 12.98 | 2.50 | 94.43 | 50.87 | 81.90 | 68.27 |
| **ENSBTAG00000017502** | RIMKLA | -2.02 | 4.21E-05 | 36.77 | 71.35 | 22.72 | 59.94 | 107.62 | 270.51 | 202.14 | 192.98 |
| **ENSBTAG00000005738** | ATP8B1 | -2.02 | 2.81E-02 | 1.99 | 79.45 | 7.30 | 41.21 | 148.59 | 123.70 | 152.47 | 101.04 |
| **ENSBTAG00000017370** | ARFGEF3 | -2.01 | 5.76E-03 | 17.89 | 98.91 | 43.81 | 34.96 | 194.41 | 227.74 | 201.27 | 167.49 |
| **ENSBTAG00000020348** | ASTL | -2.00 | 6.61E-03 | 17.89 | 68.91 | 34.08 | 13.74 | 162.47 | 80.92 | 146.38 | 153.84 |
| **ENSBTAG00000002417** | RAB30 | -2.00 | 8.10E-06 | 212.64 | 344.56 | 305.08 | 97.40 | 783.21 | 1295.91 | 882.61 | 876.59 |
| **ENSBTAG00000005413** | NLRC5 | -2.00 | 1.02E-03 | 28.82 | 109.45 | 61.66 | 58.69 | 395.08 | 166.47 | 250.93 | 220.29 |
| **ENSBTAG00000002908** | GNG4 | -1.99 | 2.00E-03 | 59.62 | 181.61 | 92.50 | 38.71 | 182.61 | 361.84 | 400.79 | 544.34 |
| **ENSBTAG00000009587** | NRG3 | -1.99 | 1.15E-02 | 11.92 | 19.46 | 30.02 | 17.48 | 31.25 | 78.61 | 92.36 | 112.87 |
| **ENSBTAG00000022006** | TRIM65 | -1.99 | 9.47E-03 | 142.09 | 220.52 | 54.36 | 168.57 | 381.19 | 691.31 | 940.11 | 312.22 |
| **ENSBTAG00000019145** | MOS | -1.99 | 4.44E-02 | 162.96 | 805.06 | 724.56 | 1364.83 | 1948.30 | 4537.42 | 2320.23 | 3308.85 |
| **ENSBTAG00000054829** | | -1.99 | 1.62E-02 | 15.90 | 21.89 | 17.04 | 14.98 | 35.41 | 115.60 | 87.13 | 40.05 |
| **ENSBTAG00000013219** | PEX5L | -1.98 | 1.26E-03 | 146.07 | 274.84 | 47.87 | 176.07 | 527.00 | 616.16 | 860.83 | 544.34 |
| **ENSBTAG00000013706** | MEGF9 | -1.98 | 4.16E-02 | 5.96 | 34.05 | 7.30 | 47.45 | 144.42 | 62.43 | 57.50 | 105.59 |
| **ENSBTAG00000004862** | TUB | -1.98 | 1.37E-04 | 41.73 | 52.70 | 61.66 | 39.96 | 156.23 | 190.75 | 311.05 | 115.60 |
| **ENSBTAG00000001455** | HOXA7 | -1.98 | 1.29E-02 | 6.96 | 74.59 | 17.04 | 42.46 | 80.54 | 226.58 | 114.14 | 134.72 |
| **ENSBTAG00000002929** | IRF4 | -1.98 | 9.43E-04 | 21.86 | 13.78 | 42.19 | 31.22 | 123.59 | 65.89 | 152.47 | 85.57 |
| **ENSBTAG00000037547** | OLIG1 | -1.97 | 1.36E-02 | 7.95 | 68.10 | 44.63 | 119.87 | 222.88 | 146.82 | 352.87 | 217.56 |
| **ENSBTAG00000024015** | PTPRM | -1.97 | 4.68E-04 | 36.77 | 43.78 | 8.93 | 61.19 | 112.48 | 174.56 | 156.83 | 141.09 |
| **ENSBTAG00000017824** | IRF8 | -1.96 | 5.62E-06 | 220.59 | 539.14 | 455.99 | 530.70 | 1421.99 | 1935.19 | 1938.60 | 1494.67 |
| **ENSBTAG00000011350** | CCNQ | -1.95 | 8.17E-06 | 1014.52 | 2164.68 | 1321.73 | 1503.43 | 2039.95 | 7767.37 | 6128.60 | 7340.45 |
| **ENSBTAG00000006844** | LEF1 | -1.95 | 1.53E-03 | 129.17 | 792.90 | 193.92 | 385.85 | 1042.19 | 910.95 | 2053.61 | 1816.00 |
| **ENSBTAG00000039720** | TMEM225B | -1.95 | 5.62E-06 | 248.41 | 764.53 | 366.74 | 297.19 | 1037.33 | 2206.86 | 1697.26 | 1561.12 |
| **ENSBTAG00000017063** | EPB41L4B | -1.94 | 1.95E-06 | 190.78 | 274.03 | 118.46 | 270.97 | 763.77 | 1120.19 | 777.18 | 624.45 |
| **ENSBTAG00000017561** | HHIPL2 | -1.94 | 4.72E-02 | 11.92 | 4.86 | 9.74 | 6.24 | 9.72 | 36.99 | 44.44 | 35.50 |
| **ENSBTAG00000004886** | ZAR1 | -1.94 | 1.37E-03 | 99.37 | 489.69 | 88.44 | 186.06 | 540.89 | 833.50 | 843.40 | 1099.61 |
| **ENSBTAG00000006209** | EXT1 | -1.93 | 1.46E-05 | 88.43 | 291.06 | 146.86 | 181.06 | 508.95 | 721.36 | 568.95 | 901.17 |
| **ENSBTAG00000020445** | ZNF398 | -1.91 | 1.32E-04 | 162.96 | 321.86 | 93.31 | 223.52 | 683.22 | 705.18 | 961.03 | 661.77 |
| **ENSBTAG00000012436** | HES7 | -1.90 | 3.42E-02 | 17.89 | 47.02 | 37.32 | 34.96 | 73.60 | 110.98 | 127.21 | 202.08 |
| **ENSBTAG00000047582** | CSNK1E | -1.90 | 3.37E-11 | 321.94 | 468.61 | 401.63 | 388.34 | 1631.68 | 1399.95 | 1589.22 | 1268.01 |
| **ENSBTAG00000034140** | MEGF11 | -1.89 | 2.71E-02 | 24.84 | 30.00 | 18.66 | 6.24 | 35.41 | 91.33 | 55.76 | 116.52 |
| **ENSBTAG00000016277** | MARC2 | -1.89 | 6.23E-03 | 27.82 | 27.57 | 53.55 | 11.24 | 59.02 | 77.45 | 129.82 | 183.88 |

| | | | | | | | | | | | |
|---|---|---|---|---|---|---|---|---|---|---|---|
| **ENSBTAG00000030259** | RASGRF2 | -1.89 | 4.10E-02 | 17.89 | 15.40 | 28.40 | 2.50 | 59.71 | 40.46 | 93.23 | 47.33 |
| **ENSBTAG00000008150** | PKIA | -1.89 | 3.11E-02 | 20.87 | 64.86 | 79.51 | 42.46 | 168.72 | 127.16 | 148.12 | 325.88 |
| **ENSBTAG00000012010** | PANX1 | -1.89 | 2.62E-03 | 68.56 | 63.24 | 21.91 | 149.84 | 356.89 | 298.26 | 250.93 | 209.36 |
| **ENSBTAG00000004593** | TOP2B | -1.88 | 5.94E-08 | 655.81 | 1455.28 | 947.69 | 907.80 | 2676.66 | 4657.65 | 3088.70 | 4232.77 |
| **ENSBTAG00000054813** | | -1.88 | 1.55E-04 | 88.43 | 279.71 | 120.90 | 164.83 | 411.05 | 698.24 | 700.51 | 601.69 |
| **ENSBTAG00000012389** | | -1.88 | 5.33E-04 | 178.86 | 510.77 | 423.54 | 206.04 | 1852.48 | 543.33 | 1132.67 | 1330.82 |
| **ENSBTAG00000015979** | SEZ6 | -1.88 | 4.23E-04 | 128.18 | 93.24 | 156.60 | 48.70 | 361.05 | 409.23 | 415.60 | 388.69 |
| **ENSBTAG00000008111** | ESYT3 | -1.88 | 4.18E-05 | 61.61 | 55.13 | 125.76 | 58.69 | 418.68 | 257.79 | 244.83 | 187.52 |
| **ENSBTAG00000023172** | ITSN2 | -1.87 | 1.23E-03 | 113.28 | 462.93 | 182.56 | 238.50 | 488.81 | 1238.11 | 993.26 | 917.56 |
| **ENSBTAG00000013491** | EML1 | -1.86 | 1.52E-02 | 8.94 | 30.81 | 67.34 | 18.73 | 101.37 | 129.48 | 104.55 | 126.53 |
| **ENSBTAG00000033967** | | -1.86 | 1.68E-02 | 13.91 | 47.83 | 41.38 | 39.96 | 66.66 | 135.26 | 90.61 | 226.66 |
| **ENSBTAG00000052895** | H2AC10 | -1.86 | 4.15E-02 | 56.64 | 68.91 | 34.08 | 34.96 | 172.89 | 18.50 | 183.84 | 329.52 |
| **ENSBTAG00000019024** | JAZF1 | -1.86 | 3.69E-03 | 39.75 | 175.93 | 97.37 | 73.67 | 337.45 | 375.71 | 385.98 | 304.03 |
| **ENSBTAG00000010597** | GGCT | -1.85 | 1.49E-03 | 155.01 | 1171.52 | 457.62 | 440.79 | 958.87 | 2762.91 | 2027.48 | 2301.17 |
| **ENSBTAG00000018773** | RND1 | -1.85 | 9.99E-04 | 390.50 | 466.18 | 577.70 | 560.66 | 1384.50 | 1371.05 | 2088.47 | 2369.44 |
| **ENSBTAG00000005429** | BCL2L10 | -1.85 | 8.85E-05 | 311.01 | 646.16 | 413.80 | 467.01 | 888.05 | 2001.09 | 2087.59 | 1668.53 |
| **ENSBTAG00000054352** | | -1.85 | 2.06E-02 | 36.77 | 9.73 | 29.21 | 52.45 | 100.68 | 89.01 | 128.08 | 142.00 |
| **ENSBTAG00000014915** | ETV5 | -1.85 | 2.81E-02 | 137.12 | 175.12 | 75.46 | 93.65 | 389.52 | 230.05 | 457.42 | 662.68 |
| **ENSBTAG00000050187** | | -1.85 | 3.32E-03 | 28.82 | 121.61 | 99.80 | 82.41 | 142.34 | 428.89 | 210.85 | 422.37 |
| **ENSBTAG00000010658** | PFKL | -1.85 | 2.69E-03 | 130.17 | 99.72 | 226.37 | 94.90 | 787.37 | 478.60 | 258.77 | 465.15 |
| **ENSBTAG00000016455** | CACHD1 | -1.84 | 7.63E-03 | 50.68 | 542.39 | 133.07 | 98.65 | 672.11 | 775.70 | 787.64 | 730.95 |
| **ENSBTAG00000023666** | | -1.84 | 7.86E-05 | 110.30 | 407.80 | 172.82 | 234.76 | 665.17 | 950.26 | 653.46 | 1043.17 |
| **ENSBTAG00000011234** | FOXO3 | -1.84 | 1.55E-04 | 37.76 | 231.87 | 103.86 | 142.35 | 449.23 | 361.84 | 538.45 | 495.19 |
| **ENSBTAG00000002289** | NLRP14 | -1.83 | 9.57E-04 | 139.11 | 850.47 | 391.08 | 278.46 | 1383.81 | 2100.51 | 1307.80 | 1131.47 |
| **ENSBTAG00000032686** | HOMEZ | -1.83 | 2.02E-02 | 20.87 | 153.23 | 45.44 | 31.22 | 135.40 | 208.09 | 261.38 | 293.11 |
| **ENSBTAG00000015177** | PRSS23 | -1.83 | 9.24E-08 | 515.70 | 587.79 | 473.03 | 349.64 | 1540.03 | 1873.93 | 2196.50 | 1258.00 |
| **ENSBTAG00000037804** | IKZF2 | -1.83 | 1.68E-02 | 11.92 | 72.97 | 27.59 | 27.47 | 73.60 | 99.42 | 203.88 | 121.98 |
| **ENSBTAG00000039766** | FBRSL1 | -1.83 | 9.37E-07 | 349.77 | 566.71 | 312.38 | 390.84 | 2422.53 | 876.27 | 1229.38 | 1212.48 |
| **ENSBTAG00000006095** | PRDM13 | -1.82 | 1.87E-03 | 62.60 | 140.26 | 47.87 | 89.91 | 180.53 | 387.27 | 372.91 | 262.16 |
| **ENSBTAG00000038532** | | -1.81 | 2.41E-03 | 77.50 | 34.05 | 47.06 | 56.19 | 151.36 | 228.89 | 210.85 | 162.94 |
| **ENSBTAG00000006054** | ACBD4 | -1.81 | 6.76E-03 | 14.90 | 29.19 | 46.25 | 37.46 | 128.45 | 99.42 | 144.63 | 76.46 |
| **ENSBTAG00000003880** | EMILIN2 | -1.81 | 1.78E-02 | 35.77 | 113.50 | 30.83 | 28.72 | 79.85 | 260.11 | 276.20 | 119.25 |
| **ENSBTAG00000002361** | TMCC2 | -1.80 | 1.07E-03 | 48.69 | 111.88 | 68.97 | 23.73 | 237.46 | 121.38 | 228.28 | 303.12 |

| | | | | | | | | | | | |
|---|---|---|---|---|---|---|---|---|---|---|---|
| **ENSBTAG00000018446** | GCA | -1.80 | 8.50E-04 | 86.45 | 582.11 | 311.57 | 259.73 | 892.91 | 1218.46 | 1137.90 | 1058.65 |
| **ENSBTAG00000009822** | PPP4R1 | -1.79 | 3.33E-03 | 40.74 | 382.67 | 126.57 | 152.34 | 922.77 | 557.21 | 532.35 | 426.92 |
| **ENSBTAG00000000815** | EPHA2 | -1.79 | 3.98E-02 | 93.40 | 78.64 | 32.46 | 38.71 | 289.54 | 223.11 | 209.11 | 121.98 |
| **ENSBTAG00000009443** | GCNT3 | -1.79 | 1.69E-04 | 189.79 | 244.84 | 59.23 | 124.87 | 556.16 | 452.01 | 575.92 | 560.73 |
| **ENSBTAG00000002253** | FKBP6 | -1.79 | 7.34E-03 | 39.75 | 248.09 | 160.65 | 106.14 | 363.83 | 442.76 | 574.18 | 542.52 |
| **ENSBTAG00000017268** | PROCA1 | -1.79 | 5.81E-06 | 266.30 | 581.30 | 349.70 | 275.96 | 1027.61 | 1227.70 | 1586.61 | 1255.27 |
| **ENSBTAG00000013726** | RNPEP | -1.79 | 2.17E-04 | 96.38 | 214.85 | 158.22 | 254.73 | 347.86 | 569.92 | 888.71 | 692.72 |
| **ENSBTAG00000004710** | HNF1B | -1.78 | 1.88E-02 | 21.86 | 135.39 | 85.19 | 32.47 | 148.59 | 317.91 | 152.47 | 331.34 |
| **ENSBTAG00000015232** | RAPGEFL1 | -1.78 | 6.66E-03 | 39.75 | 81.88 | 40.57 | 22.48 | 112.48 | 168.78 | 176.00 | 180.23 |
| **ENSBTAG00000050871** | | -1.78 | 1.60E-02 | 33.78 | 14.59 | 55.17 | 47.45 | 54.16 | 198.84 | 182.10 | 81.92 |
| **ENSBTAG00000027625** | SFRP1 | -1.78 | 4.93E-02 | 40.74 | 25.94 | 14.60 | 2.50 | 51.38 | 70.52 | 73.19 | 93.76 |
| **ENSBTAG00000046348** | ELAVL2 | -1.77 | 2.57E-04 | 158.98 | 878.03 | 404.07 | 345.89 | 879.03 | 1891.27 | 1844.51 | 1468.27 |
| **ENSBTAG00000009950** | PAX3 | -1.77 | 2.94E-05 | 197.74 | 749.93 | 285.60 | 289.70 | 984.56 | 1145.63 | 1334.81 | 1717.69 |
| **ENSBTAG00000009749** | USP2 | -1.77 | 3.26E-12 | 632.96 | 1307.72 | 660.46 | 912.80 | 3114.78 | 2897.01 | 3127.91 | 2805.46 |
| **ENSBTAG00000050602** | HAS3 | -1.76 | 2.80E-02 | 14.90 | 104.59 | 45.44 | 22.48 | 132.62 | 153.75 | 107.17 | 246.68 |
| **ENSBTAG00000005729** | FBXL4 | -1.76 | 1.92E-04 | 120.23 | 406.99 | 216.64 | 379.60 | 1010.95 | 890.14 | 892.19 | 1017.69 |
| **ENSBTAG00000020491** | SOX30 | -1.76 | 3.06E-02 | 30.80 | 47.83 | 25.96 | 7.49 | 87.49 | 113.29 | 72.32 | 110.14 |
| **ENSBTAG00000011857** | PPM1H | -1.76 | 6.49E-04 | 75.52 | 144.31 | 43.81 | 34.96 | 335.36 | 322.53 | 225.66 | 131.99 |
| **ENSBTAG00000015904** | RORA | -1.76 | 1.02E-03 | 61.61 | 54.32 | 85.19 | 34.96 | 185.39 | 253.17 | 220.43 | 142.00 |
| **ENSBTAG00000008661** | C15H11orf52 | -1.76 | 9.11E-03 | 42.73 | 37.29 | 77.08 | 27.47 | 119.43 | 141.04 | 217.82 | 147.46 |
| **ENSBTAG00000025898** | TBC1D8 | -1.74 | 1.64E-03 | 84.46 | 291.87 | 258.02 | 181.06 | 547.83 | 598.82 | 1049.90 | 535.24 |
| **ENSBTAG00000048246** | DPP4 | -1.74 | 4.04E-03 | 62.60 | 30.00 | 66.53 | 67.43 | 168.72 | 287.85 | 179.48 | 121.07 |
| **ENSBTAG00000017116** | RASD2 | -1.74 | 1.96E-02 | 37.76 | 137.83 | 42.19 | 63.68 | 268.71 | 150.28 | 234.38 | 288.56 |
| **ENSBTAG00000014636** | ZFHX3 | -1.74 | 1.28E-02 | 12.92 | 21.08 | 31.64 | 73.67 | 142.34 | 109.82 | 120.24 | 88.30 |
| **ENSBTAG00000033089** | HACD1 | -1.74 | 1.80E-02 | 12.92 | 58.37 | 94.12 | 27.47 | 110.40 | 206.93 | 105.43 | 224.84 |
| **ENSBTAG00000020796** | UBE2D1 | -1.74 | 1.35E-02 | 31.80 | 418.34 | 124.14 | 139.85 | 209.69 | 879.74 | 566.33 | 730.04 |
| **ENSBTAG00000006686** | NPNT | -1.74 | 2.47E-02 | 86.45 | 119.99 | 103.04 | 6.24 | 153.45 | 323.69 | 305.82 | 273.08 |
| **ENSBTAG00000019497** | ILDR2 | -1.73 | 3.33E-03 | 30.80 | 100.53 | 77.08 | 38.71 | 112.48 | 223.11 | 156.83 | 334.07 |
| **ENSBTAG00000021487** | CIART | -1.73 | 1.90E-04 | 435.22 | 1317.45 | 1642.22 | 554.42 | 2614.86 | 3714.33 | 3723.86 | 3053.97 |
| **ENSBTAG00000008940** | NPTX1 | -1.73 | 1.29E-04 | 194.76 | 243.22 | 219.88 | 255.98 | 1040.11 | 297.10 | 900.91 | 788.30 |
| **ENSBTAG00000004791** | VPS26C | -1.73 | 1.09E-03 | 112.28 | 318.62 | 79.51 | 234.76 | 492.98 | 567.61 | 816.39 | 588.04 |
| **ENSBTAG00000016269** | ME2 | -1.72 | 1.07E-03 | 141.10 | 193.77 | 240.98 | 389.59 | 624.21 | 601.14 | 716.19 | 1241.61 |
| **ENSBTAG00000030721** | UNCX | -1.72 | 2.97E-05 | 236.49 | 558.60 | 288.04 | 298.44 | 1047.75 | 1055.46 | 1096.07 | 1350.85 |

| | | | | | | | | | | | |
|---|---|---|---|---|---|---|---|---|---|---|---|
| **ENSBTAG00000052570** | | -1.72 | 8.92E-03 | 49.68 | 140.26 | 77.08 | 121.12 | 354.80 | 373.40 | 304.95 | 243.95 |
| **ENSBTAG00000001491** | NWD2 | -1.71 | 3.87E-02 | 18.88 | 59.99 | 94.93 | 53.69 | 56.94 | 150.28 | 292.75 | 248.50 |
| **ENSBTAG00000017983** | PRELID3A | -1.71 | 2.81E-02 | 5.96 | 76.21 | 47.06 | 39.96 | 158.31 | 104.04 | 175.13 | 119.25 |
| **ENSBTAG00000012012** | CYB5A | -1.71 | 1.43E-04 | 179.85 | 575.63 | 504.68 | 212.28 | 638.79 | 1369.90 | 1270.33 | 1556.57 |
| **ENSBTAG00000007139** | WSB2 | -1.71 | 1.61E-05 | 280.21 | 791.28 | 584.19 | 228.51 | 1478.24 | 1907.45 | 1258.13 | 1530.17 |
| **ENSBTAG00000018394** | SDR42E1 | -1.70 | 5.31E-06 | 183.83 | 184.85 | 103.04 | 255.98 | 422.85 | 751.42 | 488.79 | 692.72 |
| **ENSBTAG00000009493** | BCL3 | -1.69 | 4.28E-02 | 14.90 | 218.09 | 62.48 | 83.66 | 265.93 | 396.52 | 406.89 | 160.21 |
| **ENSBTAG00000001080** | SPAG17 | -1.69 | 1.37E-03 | 111.29 | 580.49 | 283.98 | 395.84 | 769.32 | 1595.32 | 792.87 | 1275.29 |
| **ENSBTAG00000002224** | UHRF1 | -1.69 | 1.51E-07 | 968.81 | 1937.67 | 1081.56 | 1437.25 | 4277.79 | 3968.65 | 5680.76 | 3545.52 |
| **ENSBTAG00000017354** | MPP7 | -1.68 | 6.16E-03 | 245.43 | 1008.56 | 312.38 | 389.59 | 1105.38 | 2036.93 | 1690.29 | 1448.25 |
| **ENSBTAG00000006618** | HLF | -1.68 | 2.32E-04 | 169.91 | 505.09 | 288.85 | 409.57 | 747.10 | 1166.43 | 1326.09 | 1154.23 |
| **ENSBTAG00000011044** | TACC3 | -1.68 | 1.92E-19 | 4706.93 | 5835.71 | 5586.32 | 5138.39 | 18095.02 | 14713.95 | 19635.23 | 15537.47 |
| **ENSBTAG00000030369** | IFFO1 | -1.67 | 1.46E-02 | 33.78 | 110.26 | 117.65 | 22.48 | 261.76 | 172.25 | 220.43 | 255.79 |
| **ENSBTAG00000050426** | MVB12B | -1.67 | 1.04E-02 | 34.78 | 54.32 | 16.23 | 44.95 | 93.74 | 142.19 | 98.45 | 142.91 |
| **ENSBTAG00000013825** | BCAT1 | -1.66 | 3.79E-02 | 118.24 | 232.68 | 159.03 | 139.85 | 453.40 | 587.26 | 400.79 | 614.43 |
| **ENSBTAG00000008575** | CGNL1 | -1.66 | 1.57E-02 | 72.54 | 111.88 | 64.91 | 126.12 | 508.25 | 234.67 | 283.17 | 157.48 |
| **ENSBTAG00000046918** | NPM2 | -1.66 | 1.32E-03 | 204.69 | 928.30 | 579.32 | 399.58 | 1276.88 | 2177.96 | 1346.13 | 1862.42 |
| **ENSBTAG00000008353** | CDKN1A | -1.66 | 4.71E-03 | 127.19 | 236.74 | 77.08 | 103.64 | 254.13 | 247.39 | 690.06 | 526.14 |
| **ENSBTAG00000020126** | MYO10 | -1.65 | 1.42E-04 | 143.09 | 262.68 | 184.99 | 108.64 | 699.19 | 542.18 | 399.92 | 565.28 |
| **ENSBTAG00000007841** | WTIP | -1.65 | 4.91E-03 | 127.19 | 227.82 | 104.67 | 108.64 | 203.44 | 358.37 | 617.74 | 608.06 |
| **ENSBTAG00000017053** | ABCB10 | -1.65 | 7.15E-04 | 173.89 | 613.73 | 249.09 | 203.54 | 702.67 | 1265.85 | 837.30 | 1095.97 |
| **ENSBTAG00000023283** | JAML | -1.65 | 1.60E-04 | 39.75 | 109.45 | 133.88 | 126.12 | 257.60 | 292.48 | 325.86 | 405.07 |
| **ENSBTAG00000007071** | RAI14 | -1.65 | 8.65E-04 | 221.58 | 449.96 | 234.49 | 295.94 | 1181.76 | 863.55 | 762.37 | 953.06 |
| **ENSBTAG00000038195** | CD302 | -1.64 | 4.05E-02 | 64.59 | 19.46 | 22.72 | 38.71 | 66.66 | 78.61 | 147.25 | 158.39 |
| **ENSBTAG00000033621** | SMCO4 | -1.64 | 1.09E-19 | 604.14 | 525.36 | 514.41 | 585.64 | 1456.71 | 1706.30 | 2174.72 | 1584.79 |
| **ENSBTAG00000023730** | TUBB3 | -1.63 | 5.62E-06 | 425.28 | 625.89 | 395.14 | 359.62 | 1412.27 | 891.30 | 1976.94 | 1321.72 |
| **ENSBTAG00000001514** | ASB11 | -1.63 | 5.62E-06 | 272.26 | 461.31 | 302.64 | 241.00 | 599.90 | 1394.17 | 1043.80 | 929.39 |
| **ENSBTAG00000014605** | ETV6 | -1.63 | 1.01E-02 | 91.42 | 306.46 | 128.20 | 139.85 | 175.67 | 624.26 | 580.27 | 685.44 |
| **ENSBTAG00000048098** | PKP3 | -1.63 | 6.93E-03 | 73.53 | 31.62 | 69.78 | 39.96 | 245.79 | 78.61 | 207.37 | 131.99 |
| **ENSBTAG00000015313** | CEACAM19 | -1.63 | 1.12E-02 | 33.78 | 36.48 | 11.36 | 21.23 | 72.91 | 38.15 | 108.04 | 98.31 |
| **ENSBTAG00000018845** | DND1 | -1.62 | 4.13E-04 | 56.64 | 125.66 | 111.97 | 174.82 | 333.28 | 387.27 | 441.74 | 273.99 |
| **ENSBTAG00000019456** | SPRED2 | -1.61 | 8.81E-04 | 177.86 | 452.39 | 301.83 | 227.26 | 735.30 | 756.04 | 1086.49 | 969.44 |
| **ENSBTAG00000053613** | | -1.61 | 1.72E-02 | 150.04 | 131.34 | 127.39 | 259.73 | 535.33 | 191.90 | 356.35 | 947.60 |

| | | | | | | | | | | | |
|---|---|---|---|---|---|---|---|---|---|---|---|
| **ENSBTAG00000004888** | SLC10A4 | -1.61 | 1.23E-02 | 43.72 | 38.92 | 75.46 | 89.91 | 308.98 | 250.86 | 67.09 | 125.62 |
| **ENSBTAG00000009704** | CECR2 | -1.60 | 7.15E-03 | 211.65 | 1130.17 | 377.29 | 477.00 | 1669.87 | 1784.91 | 1706.84 | 1520.16 |
| **ENSBTAG00000002385** | SLC35E4 | -1.60 | 2.37E-03 | 51.67 | 135.39 | 111.16 | 139.85 | 536.72 | 225.43 | 338.06 | 227.57 |
| **ENSBTAG00000026637** | | -1.60 | 1.09E-02 | 17.89 | 26.75 | 37.32 | 21.23 | 48.60 | 94.79 | 79.29 | 92.85 |
| **ENSBTAG00000006965** | NLRP8 | -1.60 | 2.84E-04 | 907.20 | 2152.51 | 1781.78 | 1273.67 | 6637.83 | 3525.89 | 4045.37 | 4334.72 |
| **ENSBTAG00000011639** | STK11 | -1.60 | 2.07E-04 | 210.65 | 382.67 | 279.92 | 337.15 | 1133.85 | 583.80 | 1013.30 | 933.94 |
| **ENSBTAG00000004283** | PPFIBP1 | -1.59 | 4.13E-02 | 135.14 | 108.64 | 133.88 | 126.12 | 246.49 | 332.94 | 254.41 | 685.44 |
| **ENSBTAG00000004037** | JUN | -1.59 | 4.70E-02 | 33.78 | 178.36 | 86.01 | 48.70 | 177.75 | 302.88 | 271.84 | 295.84 |
| **ENSBTAG00000020701** | MEF2C | -1.59 | 2.99E-02 | 68.56 | 72.16 | 17.85 | 38.71 | 185.39 | 198.84 | 113.27 | 96.49 |
| **ENSBTAG00000048876** | FAM117B | -1.59 | 4.34E-05 | 415.35 | 769.39 | 399.20 | 342.14 | 1337.29 | 1397.64 | 1541.30 | 1517.43 |
| **ENSBTAG00000018249** | KCNN3 | -1.58 | 9.10E-03 | 77.50 | 167.01 | 78.70 | 34.96 | 310.37 | 203.46 | 325.86 | 238.49 |
| **ENSBTAG00000044007** | KLF12 | -1.58 | 1.38E-02 | 44.71 | 12.97 | 16.23 | 27.47 | 31.94 | 99.42 | 78.42 | 92.85 |
| **ENSBTAG00000016293** | TRIM77 | -1.58 | 3.24E-07 | 620.04 | 662.37 | 674.25 | 666.80 | 1162.31 | 2166.40 | 2436.98 | 2100.00 |
| **ENSBTAG00000012509** | DYRK1B | -1.57 | 3.09E-04 | 99.37 | 319.43 | 254.77 | 163.58 | 490.89 | 675.12 | 701.38 | 628.09 |
| **ENSBTAG00000013666** | SLC25A29 | -1.57 | 1.64E-03 | 44.71 | 89.99 | 81.95 | 69.93 | 261.07 | 139.88 | 126.34 | 324.97 |
| **ENSBTAG00000007578** | SHTN1 | -1.57 | 3.21E-02 | 47.70 | 319.43 | 45.44 | 197.29 | 261.07 | 610.38 | 472.24 | 463.33 |
| **ENSBTAG00000030599** | SMOC1 | -1.56 | 8.67E-08 | 220.59 | 120.80 | 121.71 | 169.82 | 420.07 | 443.92 | 514.93 | 484.27 |
| **ENSBTAG00000010704** | LRRC4 | -1.56 | 2.26E-02 | 27.82 | 89.99 | 49.49 | 33.71 | 131.23 | 163.00 | 113.27 | 187.52 |
| **ENSBTAG00000002594** | ZNF436 | -1.56 | 6.03E-04 | 114.27 | 242.41 | 131.44 | 166.08 | 328.42 | 458.94 | 372.91 | 767.36 |
| **ENSBTAG00000048850** | | -1.56 | 7.54E-03 | 36.77 | 81.07 | 55.98 | 43.70 | 100.68 | 204.62 | 196.04 | 142.00 |
| **ENSBTAG00000010360** | LRIG1 | -1.56 | 5.41E-04 | 56.64 | 158.09 | 97.37 | 133.61 | 428.40 | 151.44 | 382.49 | 347.72 |
| **ENSBTAG00000008100** | GOLGA7B | -1.56 | 3.04E-02 | 64.59 | 417.53 | 183.37 | 128.62 | 343.70 | 635.82 | 500.12 | 858.39 |
| **ENSBTAG00000013881** | GJA4 | -1.55 | 1.74E-02 | 19.87 | 133.77 | 105.48 | 99.90 | 331.89 | 317.91 | 209.11 | 194.80 |
| **ENSBTAG00000002214** | TAT | -1.55 | 7.84E-03 | 50.68 | 73.78 | 87.63 | 33.71 | 131.23 | 154.91 | 253.54 | 183.88 |
| **ENSBTAG00000014267** | ZCCHC14 | -1.55 | 9.41E-07 | 276.24 | 708.59 | 438.14 | 402.08 | 1037.33 | 1195.34 | 1836.66 | 1280.76 |
| **ENSBTAG00000007584** | INPP1 | -1.55 | 2.21E-03 | 96.38 | 245.65 | 177.69 | 86.16 | 351.33 | 576.86 | 312.79 | 534.33 |
| **ENSBTAG00000014191** | QSOX1 | -1.54 | 7.24E-03 | 97.38 | 93.24 | 64.10 | 23.73 | 180.53 | 149.13 | 175.13 | 310.40 |
| **ENSBTAG00000031184** | CDKN1C | -1.54 | 3.15E-02 | 53.66 | 175.93 | 89.25 | 171.07 | 143.73 | 342.18 | 276.20 | 658.13 |
| **ENSBTAG00000011910** | STK33 | -1.53 | 1.56E-02 | 78.50 | 33.24 | 39.76 | 74.92 | 115.26 | 191.90 | 189.94 | 156.57 |
| **ENSBTAG00000012278** | MTMR12 | -1.53 | 1.39E-04 | 245.43 | 299.97 | 138.75 | 290.95 | 860.97 | 360.68 | 858.21 | 732.77 |
| **ENSBTAG00000016108** | BTBD2 | -1.53 | 7.33E-03 | 55.64 | 269.17 | 111.16 | 164.83 | 450.62 | 507.50 | 483.56 | 291.29 |
| **ENSBTAG00000018399** | MYH15 | -1.53 | 4.04E-03 | 134.14 | 237.55 | 200.41 | 91.15 | 200.66 | 704.02 | 574.18 | 436.02 |
| **ENSBTAG00000003889** | PER1 | -1.52 | 1.71E-06 | 639.91 | 686.70 | 544.43 | 342.14 | 1752.50 | 1346.78 | 1804.43 | 1443.69 |

| | | | | | | | | | | | |
|---|---|---|---|---|---|---|---|---|---|---|---|
| **ENSBTAG00000015119** | SIK3 | -1.52 | 1.11E-02 | 74.52 | 105.40 | 128.20 | 123.62 | 517.28 | 265.89 | 189.94 | 261.25 |
| **ENSBTAG00000006948** | GARNL3 | -1.52 | 2.06E-02 | 93.40 | 58.37 | 31.64 | 108.64 | 326.34 | 147.97 | 223.92 | 132.90 |
| **ENSBTAG00000013750** | B3GALNT2 | -1.51 | 8.48E-03 | 92.41 | 108.64 | 111.16 | 122.37 | 274.26 | 543.33 | 161.19 | 260.34 |
| **ENSBTAG00000006961** | NLRP13 | -1.51 | 4.28E-02 | 41.73 | 137.02 | 21.10 | 47.45 | 257.60 | 184.96 | 185.58 | 75.55 |
| **ENSBTAG00000038777** | MSMP | -1.50 | 3.84E-02 | 13.91 | 25.13 | 19.47 | 12.49 | 83.32 | 24.28 | 55.76 | 38.23 |
| **ENSBTAG00000049619** | GFOD1 | -1.50 | 4.23E-02 | 39.75 | 98.10 | 38.13 | 19.98 | 119.43 | 89.01 | 253.54 | 94.67 |
| **ENSBTAG00000044105** | FOXO1 | -1.49 | 4.51E-06 | 469.00 | 1090.45 | 478.71 | 688.03 | 2005.24 | 1871.61 | 1802.68 | 1979.85 |
| **ENSBTAG00000002037** | ATXN7L1 | -1.49 | 2.45E-02 | 40.74 | 186.47 | 68.97 | 53.69 | 243.02 | 277.45 | 230.02 | 234.85 |
| **ENSBTAG00000033890** | PRAME | -1.49 | 1.26E-03 | 1385.15 | 1274.48 | 1252.76 | 1017.69 | 4813.81 | 1284.35 | 2991.99 | 4742.53 |
| **ENSBTAG00000004011** | GALK2 | -1.49 | 3.30E-02 | 17.89 | 38.92 | 60.04 | 26.22 | 90.96 | 104.04 | 56.63 | 152.02 |
| **ENSBTAG00000038945** | PADI6 | -1.48 | 3.40E-03 | 86.45 | 264.30 | 153.35 | 179.81 | 607.54 | 505.19 | 460.91 | 341.35 |
| **ENSBTAG00000011851** | FYN | -1.48 | 2.28E-02 | 91.42 | 461.31 | 215.01 | 176.07 | 345.08 | 608.07 | 642.14 | 1046.82 |
| **ENSBTAG00000011540** | SPG21 | -1.48 | 8.42E-06 | 983.71 | 1706.61 | 1166.76 | 987.72 | 1907.33 | 5277.28 | 3396.26 | 2948.38 |
| **ENSBTAG00000012253** | AGO1 | -1.48 | 6.12E-03 | 65.58 | 73.78 | 45.44 | 164.83 | 223.58 | 194.21 | 214.34 | 334.98 |
| **ENSBTAG00000018598** | HSPB6 | -1.47 | 7.60E-04 | 154.02 | 63.24 | 164.71 | 114.88 | 406.88 | 290.16 | 360.71 | 322.24 |
| **ENSBTAG00000017442** | CDO1 | -1.47 | 7.86E-05 | 505.77 | 740.21 | 731.86 | 508.22 | 892.91 | 1990.68 | 2003.95 | 2018.08 |
| **ENSBTAG00000021844** | COCH | -1.47 | 3.99E-02 | 33.78 | 75.40 | 55.98 | 19.98 | 171.50 | 60.11 | 117.62 | 166.58 |
| **ENSBTAG00000047594** | CCNO | -1.47 | 8.28E-03 | 40.74 | 100.53 | 66.53 | 41.21 | 203.44 | 131.79 | 189.94 | 167.49 |
| **ENSBTAG00000007796** | POLM | -1.47 | 2.80E-02 | 61.61 | 92.42 | 101.42 | 54.94 | 202.75 | 171.09 | 272.71 | 213.91 |
| **ENSBTAG00000002363** | SESN2 | -1.47 | 2.90E-02 | 157.00 | 353.48 | 106.29 | 108.64 | 901.24 | 253.17 | 447.84 | 402.34 |
| **ENSBTAG00000021420** | EPHA7 | -1.45 | 4.95E-03 | 65.58 | 154.04 | 117.65 | 54.94 | 233.30 | 228.89 | 281.42 | 335.89 |
| **ENSBTAG00000003051** | FER | -1.45 | 3.59E-05 | 976.76 | 2124.14 | 1475.89 | 1493.44 | 2389.90 | 5514.27 | 4755.46 | 3911.45 |
| **ENSBTAG00000004442** | AGPAT1 | -1.44 | 6.08E-03 | 85.45 | 139.45 | 194.73 | 129.86 | 290.23 | 240.45 | 582.02 | 384.14 |
| **ENSBTAG00000013848** | ADGRD1 | -1.44 | 2.75E-02 | 165.94 | 411.86 | 297.77 | 127.37 | 726.97 | 736.39 | 315.40 | 949.42 |
| **ENSBTAG00000006870** | RASGEF1A | -1.44 | 9.96E-04 | 193.76 | 231.87 | 215.01 | 179.81 | 357.58 | 410.39 | 656.95 | 801.04 |
| **ENSBTAG00000026085** | ZP4 | -1.44 | 1.08E-03 | 243.44 | 507.52 | 303.45 | 703.02 | 1199.81 | 1174.53 | 865.18 | 1512.88 |
| **ENSBTAG00000009800** | MSI1 | -1.43 | 3.38E-02 | 23.85 | 169.44 | 95.74 | 59.94 | 211.08 | 224.27 | 299.72 | 209.36 |
| **ENSBTAG00000018297** | PLEKHH1 | -1.43 | 2.43E-02 | 69.56 | 82.70 | 51.93 | 56.19 | 286.07 | 124.85 | 137.66 | 152.93 |
| **ENSBTAG00000027182** | NR3C2 | -1.42 | 1.63E-06 | 337.84 | 839.93 | 668.57 | 518.21 | 1919.83 | 1152.56 | 1541.30 | 1737.71 |
| **ENSBTAG00000016091** | KLHL25 | -1.42 | 1.91E-03 | 141.10 | 320.24 | 144.42 | 133.61 | 504.09 | 286.70 | 697.03 | 496.10 |
| **ENSBTAG00000010452** | PODXL | -1.42 | 3.55E-03 | 60.61 | 129.72 | 88.44 | 124.87 | 360.36 | 228.89 | 232.63 | 256.70 |
| **ENSBTAG00000008005** | DRC7 | -1.42 | 2.61E-02 | 40.74 | 189.71 | 77.08 | 56.19 | 236.07 | 199.99 | 221.31 | 318.60 |
| **ENSBTAG00000001002** | TCF7 | -1.42 | 1.78E-02 | 81.48 | 165.39 | 44.63 | 51.20 | 169.42 | 216.18 | 286.65 | 246.68 |

| | | | | | | | | | | | |
|---|---|---|---|---|---|---|---|---|---|---|---|
| **ENSBTAG00000054030** | | -1.42 | 3.67E-03 | 62.60 | 124.85 | 103.86 | 111.13 | 111.79 | 406.92 | 192.55 | 366.84 |
| **ENSBTAG00000055205** | | -1.41 | 3.88E-04 | 88.43 | 117.56 | 77.08 | 66.18 | 174.97 | 189.59 | 301.46 | 266.71 |
| **ENSBTAG00000010947** | PHYHIPL | -1.41 | 2.86E-03 | 821.75 | 1036.94 | 589.87 | 847.87 | 1471.99 | 3028.80 | 2221.77 | 2049.94 |
| **ENSBTAG00000000473** | ATP10D | -1.41 | 6.52E-03 | 97.38 | 257.82 | 279.92 | 272.22 | 302.73 | 667.03 | 585.50 | 859.30 |
| **ENSBTAG00000002728** | ARID1B | -1.41 | 1.06E-04 | 200.72 | 250.52 | 125.76 | 143.60 | 514.50 | 425.42 | 429.54 | 546.16 |
| **ENSBTAG00000014791** | CTH | -1.41 | 5.64E-03 | 102.35 | 144.31 | 41.38 | 89.91 | 164.56 | 245.08 | 359.84 | 232.12 |
| **ENSBTAG00000004498** | ESR2 | -1.40 | 7.17E-03 | 90.42 | 32.43 | 68.16 | 47.45 | 138.17 | 175.72 | 165.54 | 152.02 |
| **ENSBTAG00000002144** | ADRB2 | -1.40 | 3.88E-04 | 657.80 | 964.78 | 765.94 | 1058.90 | 1355.34 | 2070.45 | 3971.31 | 1699.48 |
| **ENSBTAG00000019707** | GATA2 | -1.40 | 8.12E-08 | 303.06 | 511.58 | 388.65 | 349.64 | 838.75 | 969.91 | 1177.10 | 1115.09 |
| **ENSBTAG00000000949** | DNAJC9 | -1.39 | 2.05E-05 | 345.79 | 870.73 | 426.78 | 548.18 | 1250.49 | 1462.38 | 1474.21 | 1579.33 |
| **ENSBTAG00000007494** | SMARCA2 | -1.39 | 9.75E-04 | 215.62 | 449.96 | 447.07 | 398.33 | 740.16 | 941.01 | 1092.59 | 1197.92 |
| **ENSBTAG00000007490** | SULF2 | -1.39 | 1.84E-02 | 46.70 | 45.40 | 47.06 | 51.20 | 61.10 | 154.91 | 153.35 | 131.08 |
| **ENSBTAG00000016658** | ABHD4 | -1.39 | 3.13E-03 | 251.39 | 444.29 | 112.78 | 243.50 | 739.47 | 517.90 | 664.79 | 838.36 |
| **ENSBTAG00000002069** | BOLA | -1.39 | 1.11E-03 | 132.16 | 239.98 | 132.25 | 124.87 | 575.60 | 194.21 | 369.42 | 511.57 |
| **ENSBTAG00000007898** | CYBRD1 | -1.39 | 1.52E-02 | 106.32 | 167.01 | 163.90 | 77.42 | 342.31 | 375.71 | 302.34 | 332.25 |
| **ENSBTAG00000033008** | MYOZ1 | -1.39 | 2.98E-03 | 48.69 | 30.00 | 31.64 | 43.70 | 118.73 | 105.20 | 103.68 | 73.73 |
| **ENSBTAG00000016164** | LMBRD1 | -1.39 | 9.01E-04 | 1068.18 | 2280.61 | 816.24 | 1070.13 | 2519.04 | 4920.07 | 2863.04 | 3390.77 |
| **ENSBTAG00000017155** | TRIM32 | -1.38 | 3.08E-05 | 1150.65 | 1709.04 | 1582.99 | 1291.15 | 5453.99 | 3206.83 | 3685.53 | 2592.46 |
| **ENSBTAG00000023765** | C18H16orf46 | -1.38 | 2.21E-02 | 45.71 | 69.72 | 162.28 | 84.91 | 303.42 | 154.91 | 215.21 | 271.26 |
| **ENSBTAG00000016407** | IRX6 | -1.38 | 3.58E-02 | 56.64 | 77.02 | 32.46 | 66.18 | 202.05 | 77.45 | 199.52 | 123.80 |
| **ENSBTAG00000008105** | RBM38 | -1.38 | 1.42E-04 | 584.27 | 1068.56 | 583.38 | 691.78 | 1521.98 | 1349.09 | 2788.11 | 1943.44 |
| **ENSBTAG00000020346** | CCDC92 | -1.38 | 7.98E-03 | 64.59 | 90.80 | 68.16 | 58.69 | 197.19 | 169.94 | 196.04 | 170.22 |
| **ENSBTAG00000001116** | P4HA2 | -1.37 | 3.32E-02 | 157.00 | 141.88 | 99.80 | 61.19 | 265.24 | 252.01 | 284.04 | 390.51 |
| **ENSBTAG00000003512** | MYH7B | -1.37 | 4.97E-02 | 70.55 | 83.51 | 38.13 | 71.18 | 186.78 | 161.84 | 234.38 | 94.67 |
| **ENSBTAG00000052516** | CDA | -1.36 | 5.39E-04 | 125.20 | 176.74 | 99.80 | 73.67 | 377.02 | 285.54 | 280.55 | 281.27 |
| **ENSBTAG00000019839** | LTBP1 | -1.36 | 1.00E-02 | 138.12 | 144.31 | 105.48 | 79.92 | 206.91 | 273.98 | 277.07 | 442.39 |
| **ENSBTAG00000021505** | USP36 | -1.35 | 9.43E-06 | 1196.36 | 1853.35 | 1159.46 | 1111.34 | 4377.08 | 2335.18 | 4000.06 | 2869.18 |
| **ENSBTAG00000048544** | | -1.35 | 2.90E-02 | 59.62 | 306.46 | 159.03 | 113.63 | 293.70 | 397.67 | 446.10 | 496.10 |
| **ENSBTAG00000000260** | ZNRF2 | -1.35 | 7.66E-06 | 614.08 | 1185.30 | 713.20 | 1076.38 | 1375.47 | 2891.23 | 2855.19 | 2024.45 |
| **ENSBTAG00000049111** | | -1.35 | 8.50E-04 | 107.31 | 107.02 | 115.22 | 114.88 | 174.97 | 331.78 | 320.63 | 304.94 |
| **ENSBTAG00000015982** | KCTD13 | -1.35 | 1.30E-08 | 621.03 | 724.80 | 645.86 | 706.76 | 1471.99 | 1702.83 | 1732.98 | 1960.73 |
| **ENSBTAG00000013901** | PTDSS1 | -1.35 | 1.67E-03 | 455.09 | 766.15 | 628.00 | 616.86 | 2421.14 | 1268.17 | 1281.66 | 1297.14 |
| **ENSBTAG00000043957** | TDRP | -1.35 | 1.97E-02 | 28.82 | 128.10 | 51.12 | 88.66 | 152.06 | 213.87 | 203.01 | 185.70 |

| | | | | | | | | | | | |
|---|---|---|---|---|---|---|---|---|---|---|---|
| **ENSBTAG00000004694** | RPGRIP1 | -1.34 | 1.05E-02 | 253.38 | 633.19 | 207.71 | 279.71 | 626.98 | 1221.92 | 670.89 | 961.25 |
| **ENSBTAG00000009290** | FAM161A | -1.34 | 5.31E-06 | 632.96 | 1216.11 | 999.62 | 1002.70 | 1601.13 | 2656.56 | 2522.36 | 2964.76 |
| **ENSBTAG00000020797** | CABLES2 | -1.34 | 3.79E-03 | 114.27 | 115.94 | 144.42 | 219.77 | 422.85 | 368.77 | 455.68 | 249.42 |
| **ENSBTAG00000004361** | ZBTB37 | -1.34 | 2.26E-02 | 159.98 | 136.20 | 215.01 | 98.65 | 403.41 | 321.38 | 487.92 | 327.70 |
| **ENSBTAG00000021183** | MAP3K21 | -1.33 | 4.37E-02 | 104.33 | 451.58 | 150.10 | 112.38 | 434.65 | 359.53 | 670.89 | 600.78 |
| **ENSBTAG00000016836** | PDK1 | -1.33 | 8.83E-05 | 528.62 | 657.51 | 471.41 | 370.86 | 973.46 | 1443.88 | 1296.47 | 1400.00 |
| **ENSBTAG00000024204** | H2BK1 | -1.33 | 1.49E-03 | 67.57 | 113.50 | 84.38 | 141.10 | 236.77 | 198.84 | 319.76 | 262.16 |
| **ENSBTAG00000023179** | TRIB1 | -1.33 | 6.26E-04 | 517.69 | 1441.50 | 773.24 | 759.21 | 1997.60 | 2038.08 | 2470.96 | 2249.29 |
| **ENSBTAG00000050550** | FAM110B | -1.32 | 9.46E-03 | 134.14 | 301.60 | 146.86 | 76.17 | 311.06 | 431.20 | 413.86 | 495.19 |
| **ENSBTAG00000004780** | CDKL1 | -1.32 | 2.14E-02 | 97.38 | 149.18 | 81.95 | 152.34 | 149.28 | 383.80 | 319.76 | 347.72 |
| **ENSBTAG00000044155** | BSPRY | -1.32 | 3.84E-02 | 57.63 | 51.08 | 48.68 | 53.69 | 112.48 | 121.38 | 148.99 | 143.82 |
| **ENSBTAG00000050737** | | -1.32 | 9.90E-03 | 49.68 | 87.56 | 61.66 | 106.14 | 204.13 | 75.14 | 250.93 | 226.66 |
| **ENSBTAG00000018936** | LSS | -1.31 | 6.64E-03 | 318.96 | 201.87 | 167.95 | 123.62 | 563.80 | 430.04 | 423.44 | 603.51 |
| **ENSBTAG00000051258** | | -1.31 | 3.61E-02 | 29.81 | 19.46 | 27.59 | 18.73 | 71.52 | 47.40 | 58.38 | 60.08 |
| **ENSBTAG00000012387** | PAM | -1.31 | 4.43E-02 | 191.77 | 301.60 | 236.92 | 232.26 | 399.24 | 336.40 | 345.90 | 1306.24 |
| **ENSBTAG00000021176** | CRISPLD2 | -1.31 | 2.39E-02 | 170.91 | 278.89 | 82.76 | 164.83 | 294.40 | 252.01 | 690.06 | 491.55 |
| **ENSBTAG00000002630** | MRTFA | -1.31 | 1.20E-02 | 49.68 | 53.51 | 66.53 | 68.68 | 168.72 | 141.04 | 174.26 | 105.59 |
| **ENSBTAG00000021752** | DNAJB4 | -1.31 | 1.16E-03 | 109.30 | 149.18 | 248.28 | 177.31 | 485.34 | 336.40 | 327.60 | 544.34 |
| **ENSBTAG00000014619** | WASF3 | -1.31 | 1.64E-03 | 238.48 | 435.37 | 315.63 | 242.25 | 654.76 | 752.58 | 730.14 | 914.83 |
| **ENSBTAG00000012382** | KCTD9 | -1.31 | 5.14E-06 | 478.94 | 848.84 | 805.70 | 639.33 | 1131.07 | 2053.11 | 1712.94 | 1968.01 |
| **ENSBTAG00000016259** | KIAA1958 | -1.31 | 8.01E-03 | 85.45 | 170.26 | 59.23 | 94.90 | 218.02 | 334.09 | 194.30 | 268.53 |
| **ENSBTAG00000006667** | EPB41 | -1.31 | 5.08E-04 | 346.78 | 701.29 | 399.20 | 574.40 | 1347.01 | 1297.07 | 1381.85 | 969.44 |
| **ENSBTAG00000000279** | RAB26 | -1.30 | 2.96E-02 | 74.52 | 212.41 | 118.46 | 67.43 | 175.67 | 304.04 | 347.64 | 342.26 |
| **ENSBTAG00000016915** | FLT1 | -1.30 | 3.46E-02 | 94.40 | 391.59 | 159.03 | 207.28 | 530.47 | 853.15 | 324.12 | 394.15 |
| **ENSBTAG00000006573** | KLHL18 | -1.30 | 5.60E-03 | 321.94 | 625.08 | 356.19 | 575.65 | 1757.36 | 1077.42 | 1014.17 | 771.00 |
| **ENSBTAG00000005824** | SPOP | -1.30 | 7.72E-05 | 478.94 | 618.59 | 623.95 | 611.86 | 1010.26 | 1908.61 | 1281.66 | 1531.99 |
| **ENSBTAG00000018084** | CASTOR1 | -1.30 | 8.41E-05 | 492.85 | 787.23 | 597.98 | 638.08 | 1112.32 | 1491.28 | 2000.47 | 1572.95 |
| **ENSBTAG00000014861** | SLC20A2 | -1.29 | 2.22E-02 | 101.35 | 92.42 | 72.21 | 49.95 | 136.09 | 145.66 | 269.23 | 224.84 |
| **ENSBTAG00000010023** | FAM107B | -1.29 | 1.91E-02 | 74.52 | 282.14 | 201.22 | 74.92 | 345.78 | 347.97 | 388.59 | 469.70 |
| **ENSBTAG00000002888** | TMTC2 | -1.28 | 1.42E-02 | 169.91 | 273.22 | 469.79 | 239.75 | 649.90 | 802.29 | 731.88 | 621.72 |
| **ENSBTAG00000001693** | RPAP1 | -1.28 | 8.71E-06 | 467.02 | 474.28 | 305.89 | 433.30 | 1293.54 | 714.43 | 1025.50 | 1047.73 |
| **ENSBTAG00000030416** | | -1.28 | 3.65E-02 | 26.83 | 11.35 | 21.91 | 22.48 | 54.16 | 50.87 | 48.79 | 45.51 |
| **ENSBTAG00000004383** | FNBP1L | -1.27 | 2.12E-03 | 398.45 | 859.38 | 525.77 | 427.05 | 638.09 | 1781.44 | 1443.72 | 1484.66 |

| | | | | | | | | | | | |
|---|---|---|---|---|---|---|---|---|---|---|---|
| **ENSBTAG00000004036** | GJC1 | -1.27 | 5.09E-05 | 403.42 | 773.45 | 568.77 | 694.28 | 1326.18 | 1262.39 | 1633.66 | 1665.80 |
| **ENSBTAG00000030898** | MS4A13 | -1.27 | 3.16E-02 | 218.60 | 246.46 | 271.81 | 94.90 | 249.27 | 324.84 | 733.62 | 702.73 |
| **ENSBTAG00000005498** | SQLE | -1.27 | 7.08E-04 | 1252.00 | 1275.29 | 1041.00 | 990.22 | 3611.92 | 1209.21 | 2810.76 | 3356.18 |
| **ENSBTAG00000010626** | SH3GLB2 | -1.27 | 4.73E-03 | 163.95 | 211.60 | 263.70 | 223.52 | 649.20 | 393.05 | 528.87 | 505.20 |
| **ENSBTAG00000005788** | VANGL1 | -1.27 | 1.12E-02 | 86.45 | 141.07 | 83.57 | 61.19 | 231.21 | 297.10 | 252.67 | 118.34 |
| **ENSBTAG00000032021** | RALB | -1.27 | 1.17E-06 | 2806.07 | 3576.99 | 2593.16 | 2477.41 | 5516.48 | 9249.40 | 6122.50 | 6678.68 |
| **ENSBTAG00000021535** | CROT | -1.26 | 1.02E-03 | 644.88 | 959.92 | 477.90 | 711.76 | 868.61 | 2246.17 | 1448.94 | 2148.25 |
| **ENSBTAG00000043981** | SBF2 | -1.26 | 8.47E-03 | 125.20 | 300.78 | 90.06 | 157.34 | 273.57 | 499.41 | 419.09 | 426.92 |
| **ENSBTAG00000017263** | MXI1 | -1.26 | 3.50E-04 | 364.67 | 950.19 | 608.53 | 539.44 | 946.38 | 1486.66 | 1643.24 | 1821.46 |
| **ENSBTAG00000020780** | SBNO2 | -1.26 | 3.15E-02 | 83.47 | 119.18 | 71.40 | 103.64 | 345.78 | 123.70 | 233.50 | 198.44 |
| **ENSBTAG00000003836** | ADAM19 | -1.26 | 3.16E-04 | 699.53 | 1096.93 | 577.70 | 754.21 | 1243.55 | 1930.57 | 2171.24 | 2130.95 |
| **ENSBTAG00000011431** | GBA2 | -1.25 | 5.90E-04 | 179.85 | 291.87 | 234.49 | 216.02 | 814.45 | 599.98 | 400.79 | 379.58 |
| **ENSBTAG00000019581** | PLEKHF2 | -1.25 | 1.84E-04 | 652.83 | 1563.11 | 1148.10 | 1037.67 | 2061.48 | 3165.21 | 2854.32 | 2372.17 |
| **ENSBTAG00000024957** | SNCA | -1.25 | 4.09E-03 | 167.93 | 401.32 | 435.71 | 232.26 | 495.75 | 808.07 | 677.86 | 957.61 |
| **ENSBTAG00000017976** | ABL1 | -1.25 | 2.10E-03 | 330.89 | 661.56 | 466.54 | 226.01 | 1009.56 | 941.01 | 1228.51 | 821.07 |
| **ENSBTAG00000040116** | H1FX | -1.25 | 7.54E-03 | 835.66 | 963.16 | 831.66 | 1193.75 | 1506.70 | 1360.65 | 4412.18 | 1783.23 |
| **ENSBTAG00000018313** | MBNL2 | -1.24 | 4.08E-02 | 172.90 | 323.49 | 107.91 | 283.45 | 342.31 | 399.99 | 329.34 | 1031.34 |
| **ENSBTAG00000005412** | NEDD4L | -1.24 | 7.34E-03 | 213.64 | 440.23 | 281.55 | 177.31 | 751.27 | 482.06 | 845.14 | 558.00 |
| **ENSBTAG00000012343** | TSPAN5 | -1.24 | 1.68E-03 | 306.04 | 516.44 | 829.23 | 706.76 | 886.66 | 1663.53 | 1320.86 | 1703.12 |
| **ENSBTAG00000012500** | RARA | -1.24 | 1.20E-02 | 58.63 | 68.91 | 61.66 | 69.93 | 182.61 | 176.87 | 135.92 | 115.60 |
| **ENSBTAG00000021339** | SCN8A | -1.24 | 1.78E-02 | 177.86 | 579.68 | 334.29 | 218.52 | 785.99 | 1063.55 | 659.56 | 583.49 |
| **ENSBTAG00000001879** | PER2 | -1.23 | 2.30E-02 | 50.68 | 143.50 | 52.74 | 68.68 | 186.08 | 232.36 | 167.29 | 159.30 |
| **ENSBTAG00000032455** | HIST2H2BF | -1.23 | 6.75E-08 | 402.43 | 534.28 | 483.58 | 443.29 | 809.59 | 1151.41 | 1008.07 | 1420.03 |
| **ENSBTAG00000025046** | ALKBH5 | -1.23 | 2.76E-11 | 3771.90 | 5310.35 | 4342.48 | 4010.81 | 11280.84 | 9139.58 | 10416.19 | 10089.48 |
| **ENSBTAG00000003994** | IGFBP3 | -1.23 | 1.37E-05 | 332.87 | 406.99 | 477.90 | 383.35 | 1124.13 | 1119.04 | 587.24 | 925.75 |
| **ENSBTAG00000019213** | CNNM4 | -1.23 | 3.38E-02 | 423.30 | 128.10 | 581.76 | 549.43 | 1409.50 | 624.26 | 1081.26 | 825.62 |
| **ENSBTAG00000008842** | JPH1 | -1.23 | 2.78E-02 | 164.95 | 269.17 | 229.62 | 126.12 | 724.88 | 399.99 | 356.35 | 371.39 |
| **ENSBTAG00000019406** | IGF2BP3 | -1.23 | 1.26E-04 | 1473.58 | 2604.91 | 1927.02 | 1580.85 | 3150.89 | 4940.87 | 5848.05 | 3825.88 |
| **ENSBTAG00000020119** | RNF43 | -1.22 | 4.94E-02 | 56.64 | 50.27 | 63.29 | 28.72 | 74.29 | 84.39 | 194.30 | 113.78 |
| **ENSBTAG00000008895** | BPGM | -1.22 | 4.18E-05 | 835.66 | 1125.31 | 1033.69 | 752.96 | 1441.44 | 2626.50 | 2582.48 | 2103.64 |
| **ENSBTAG00000005650** | SKAP2 | -1.22 | 2.19E-02 | 207.67 | 94.05 | 211.77 | 57.44 | 347.86 | 338.72 | 216.95 | 431.47 |
| **ENSBTAG00000009565** | RASA1 | -1.22 | 3.46E-04 | 591.22 | 1088.01 | 797.58 | 708.01 | 1159.54 | 2072.76 | 1829.69 | 2363.98 |
| **ENSBTAG00000025441** | HSPA1A | -1.22 | 2.72E-04 | 2042.95 | 1438.25 | 3072.68 | 2195.21 | 5365.11 | 2967.53 | 4510.63 | 7526.15 |

| | | | | | | | | | | | |
|---|---|---|---|---|---|---|---|---|---|---|---|
| **ENSBTAG00000010192** | LYSMD4 | -1.22 | 1.57E-02 | 158.98 | 470.23 | 135.50 | 260.98 | 508.25 | 759.51 | 636.04 | 483.36 |
| **ENSBTAG00000014501** | FERMT2 | -1.22 | 5.12E-03 | 656.80 | 2720.84 | 1333.90 | 1421.02 | 2478.08 | 3678.49 | 4663.98 | 3439.93 |
| **ENSBTAG00000027868** | DLG2 | -1.22 | 2.09E-02 | 110.30 | 104.59 | 50.31 | 148.59 | 181.92 | 211.55 | 264.00 | 301.30 |
| **ENSBTAG00000011905** | RRAS2 | -1.21 | 8.12E-08 | 3626.83 | 6055.42 | 3673.10 | 4582.72 | 9926.19 | 12739.46 | 9263.48 | 9663.47 |
| **ENSBTAG00000000561** | OCLN | -1.21 | 2.07E-08 | 3675.52 | 3176.48 | 4369.26 | 3736.10 | 6852.38 | 11264.36 | 8380.87 | 8127.84 |
| **ENSBTAG00000007519** | ADAR | -1.20 | 4.13E-04 | 329.89 | 458.07 | 452.75 | 317.17 | 1115.10 | 667.03 | 880.87 | 923.93 |
| **ENSBTAG00000021747** | | -1.20 | 1.08E-04 | 573.34 | 454.83 | 432.46 | 630.59 | 793.62 | 1310.94 | 1427.16 | 1276.20 |
| **ENSBTAG00000021527** | IGF1R | -1.20 | 4.17E-03 | 147.06 | 181.61 | 249.90 | 129.86 | 498.53 | 393.05 | 362.45 | 371.39 |
| **ENSBTAG00000016297** | CCNYL1 | -1.19 | 3.50E-05 | 435.22 | 445.10 | 425.16 | 440.79 | 729.05 | 1069.33 | 1307.80 | 885.70 |
| **ENSBTAG00000021933** | ALOX12 | -1.19 | 2.94E-02 | 47.70 | 70.53 | 45.44 | 29.97 | 111.79 | 62.43 | 143.76 | 125.62 |
| **ENSBTAG00000020854** | BCL6B | -1.19 | 1.98E-02 | 109.30 | 94.86 | 183.37 | 134.86 | 199.97 | 418.48 | 376.39 | 195.71 |
| **ENSBTAG00000008638** | FAM110A | -1.18 | 6.12E-03 | 328.90 | 713.45 | 426.78 | 288.45 | 574.91 | 1558.33 | 957.54 | 899.35 |
| **ENSBTAG00000006760** | OTX1 | -1.18 | 9.72E-04 | 411.37 | 429.69 | 397.57 | 530.70 | 1336.59 | 701.71 | 970.61 | 993.11 |
| **ENSBTAG00000019044** | BAIAP2 | -1.18 | 3.23E-03 | 751.20 | 1058.83 | 558.23 | 766.70 | 1874.01 | 1285.51 | 2174.72 | 1755.01 |
| **ENSBTAG00000004899** | ABLIM1 | -1.18 | 4.07E-05 | 793.93 | 807.50 | 645.86 | 718.00 | 1166.48 | 2102.82 | 1665.02 | 1770.48 |
| **ENSBTAG00000004976** | CDCA7L | -1.17 | 1.40E-02 | 145.07 | 428.07 | 285.60 | 324.66 | 458.26 | 760.67 | 729.26 | 722.76 |
| **ENSBTAG00000004386** | SOCS1 | -1.17 | 7.50E-05 | 420.31 | 447.53 | 335.10 | 337.15 | 1029.00 | 983.78 | 724.91 | 728.22 |
| **ENSBTAG00000051328** | ZNF621 | -1.17 | 4.85E-02 | 67.57 | 36.48 | 30.02 | 106.14 | 174.28 | 105.20 | 135.05 | 120.16 |
| **ENSBTAG00000004561** | PAX6 | -1.17 | 1.08E-02 | 58.63 | 58.37 | 61.66 | 51.20 | 113.18 | 76.30 | 155.09 | 172.04 |
| **ENSBTAG00000001830** | LRRC46 | -1.17 | 4.05E-02 | 97.38 | 26.75 | 115.22 | 87.41 | 241.63 | 115.60 | 165.54 | 208.45 |
| **ENSBTAG00000020533** | FBXO34 | -1.17 | 2.20E-04 | 348.77 | 774.26 | 475.47 | 586.89 | 967.21 | 1567.58 | 1069.94 | 1301.69 |
| **ENSBTAG00000033248** | CDH3 | -1.16 | 5.67E-03 | 186.81 | 329.97 | 362.69 | 515.71 | 858.89 | 687.84 | 891.32 | 682.71 |
| **ENSBTAG00000006985** | TM7SF3 | -1.16 | 5.41E-04 | 256.36 | 473.47 | 701.84 | 477.00 | 1340.76 | 1238.11 | 715.32 | 979.45 |
| **ENSBTAG00000019251** | EPB41L3 | -1.16 | 3.86E-02 | 115.26 | 211.60 | 116.84 | 158.58 | 578.38 | 248.55 | 279.68 | 234.85 |
| **ENSBTAG00000001294** | PPP1R15A | -1.16 | 2.62E-03 | 522.66 | 673.72 | 481.96 | 362.12 | 1562.94 | 758.36 | 1301.70 | 923.93 |
| **ENSBTAG00000007123** | ENSA | -1.15 | 4.41E-09 | 1740.88 | 2416.82 | 2018.70 | 1992.92 | 3675.11 | 5555.88 | 4156.89 | 4769.84 |
| **ENSBTAG00000001185** | KMT5B | -1.15 | 5.94E-08 | 1591.83 | 1876.05 | 1775.29 | 2462.43 | 4237.52 | 4500.43 | 4160.38 | 4193.63 |
| **ENSBTAG00000051440** | | -1.15 | 1.04E-02 | 57.63 | 66.48 | 70.59 | 131.11 | 151.36 | 164.16 | 224.79 | 178.41 |
| **ENSBTAG00000010835** | CRELD1 | -1.14 | 8.76E-04 | 221.58 | 257.00 | 284.79 | 274.71 | 629.76 | 393.05 | 777.18 | 493.37 |
| **ENSBTAG00000002315** | RNF34 | -1.14 | 1.22E-07 | 2170.13 | 3022.44 | 3143.27 | 3398.95 | 4221.55 | 6864.51 | 7549.67 | 7294.03 |
| **ENSBTAG00000009907** | MAPK4 | -1.14 | 5.98E-03 | 65.58 | 68.10 | 86.01 | 61.19 | 133.31 | 180.34 | 176.87 | 131.99 |
| **ENSBTAG00000046218** | KLF11 | -1.14 | 3.43E-03 | 389.51 | 594.27 | 324.55 | 380.85 | 843.62 | 1057.77 | 809.42 | 1003.12 |
| **ENSBTAG00000004266** | DDX25 | -1.14 | 1.38E-02 | 637.92 | 476.72 | 505.49 | 344.64 | 946.38 | 562.99 | 1922.05 | 885.70 |

| | | | | | | | | | | | |
|---|---|---|---|---|---|---|---|---|---|---|---|
| **ENSBTAG00000045904** | PIM3 | -1.13 | 1.64E-03 | 706.49 | 899.11 | 764.32 | 570.65 | 2105.91 | 973.38 | 1828.82 | 1540.18 |
| **ENSBTAG00000011646** | ZNF512B | -1.13 | 3.53E-04 | 198.73 | 185.66 | 143.61 | 282.21 | 482.56 | 393.05 | 471.36 | 419.64 |
| **ENSBTAG00000006022** | ATP6V0E2 | -1.13 | 4.58E-02 | 145.07 | 39.73 | 51.12 | 43.70 | 134.70 | 168.78 | 148.99 | 158.39 |
| **ENSBTAG00000007147** | CUEDC1 | -1.13 | 1.98E-02 | 204.69 | 367.27 | 165.52 | 216.02 | 583.93 | 394.21 | 569.82 | 532.51 |
| **ENSBTAG00000001060** | CXCR4 | -1.12 | 1.54E-04 | 1429.86 | 1752.01 | 2651.58 | 1816.85 | 3539.02 | 3693.52 | 4298.04 | 5104.82 |
| **ENSBTAG00000003504** | GSS | -1.12 | 5.39E-03 | 129.17 | 256.19 | 133.07 | 76.17 | 263.85 | 331.78 | 326.73 | 373.21 |
| **ENSBTAG00000011504** | ZP2 | -1.12 | 1.62E-03 | 368.64 | 741.02 | 524.15 | 370.86 | 978.32 | 1109.79 | 828.59 | 1435.50 |
| **ENSBTAG00000009760** | | -1.11 | 2.23E-03 | 546.51 | 532.66 | 619.89 | 480.75 | 1353.95 | 1218.46 | 1251.16 | 893.89 |
| **ENSBTAG00000048999** | TRIM6 | -1.11 | 1.09E-02 | 731.33 | 1910.10 | 917.67 | 849.11 | 2804.41 | 2358.30 | 2285.37 | 2087.26 |
| **ENSBTAG00000002689** | NME7 | -1.11 | 1.42E-02 | 152.03 | 120.80 | 227.19 | 137.36 | 231.91 | 412.70 | 331.09 | 404.16 |
| **ENSBTAG00000021458** | DLX6 | -1.11 | 3.15E-02 | 147.06 | 393.21 | 214.20 | 179.81 | 426.32 | 497.09 | 572.43 | 525.23 |
| **ENSBTAG00000003474** | RAB15 | -1.11 | 2.77E-02 | 207.67 | 518.06 | 345.65 | 419.56 | 1176.90 | 431.20 | 887.84 | 722.76 |
| **ENSBTAG00000015743** | GMPR | -1.11 | 4.47E-03 | 459.07 | 435.37 | 415.42 | 345.89 | 1610.85 | 809.22 | 582.02 | 570.74 |
| **ENSBTAG00000009194** | OSBPL10 | -1.11 | 1.81E-02 | 209.66 | 215.66 | 69.78 | 137.36 | 298.56 | 261.26 | 456.55 | 347.72 |
| **ENSBTAG00000011150** | PFN2 | -1.10 | 3.76E-03 | 443.17 | 840.74 | 544.43 | 583.14 | 1188.00 | 1530.58 | 1143.12 | 1324.45 |
| **ENSBTAG00000054533** | GNG12 | -1.10 | 1.06E-03 | 364.67 | 747.50 | 678.31 | 478.25 | 906.11 | 1152.56 | 1351.36 | 1465.54 |
| **ENSBTAG00000021021** | C21H15orf39 | -1.10 | 4.37E-03 | 969.80 | 877.22 | 753.77 | 689.28 | 3040.49 | 1246.20 | 1429.78 | 1344.47 |
| **ENSBTAG00000018272** | RERE | -1.10 | 1.18E-04 | 234.50 | 366.45 | 264.51 | 280.96 | 679.75 | 530.62 | 740.59 | 509.75 |
| **ENSBTAG00000001592** | INSIG1 | -1.10 | 1.52E-02 | 827.71 | 615.35 | 724.56 | 556.92 | 1352.56 | 865.87 | 1428.03 | 2193.76 |
| **ENSBTAG00000014011** | TMOD2 | -1.10 | 6.92E-04 | 248.41 | 351.05 | 237.73 | 272.22 | 603.38 | 579.17 | 647.36 | 547.07 |
| **ENSBTAG00000018910** | WDR6 | -1.10 | 5.14E-03 | 1965.44 | 1879.30 | 1445.06 | 1964.20 | 6380.23 | 2034.61 | 4023.58 | 3080.37 |
| **ENSBTAG00000012555** | RBM15 | -1.10 | 5.75E-03 | 322.94 | 496.98 | 487.64 | 393.34 | 658.23 | 564.14 | 1116.11 | 1298.05 |
| **ENSBTAG00000047502** | FKBP5 | -1.09 | 3.30E-03 | 376.59 | 538.33 | 758.64 | 691.78 | 1176.90 | 1676.24 | 1016.79 | 1183.36 |
| **ENSBTAG00000018421** | WWP2 | -1.09 | 5.77E-03 | 329.89 | 443.47 | 606.10 | 295.94 | 1035.25 | 649.69 | 776.31 | 1105.98 |
| **ENSBTAG00000010582** | BRD3 | -1.08 | 1.42E-04 | 938.01 | 1283.40 | 910.36 | 930.28 | 2280.89 | 1910.92 | 2216.54 | 2192.85 |
| **ENSBTAG00000052673** | | -1.08 | 1.67E-02 | 178.86 | 243.22 | 201.22 | 162.33 | 340.22 | 490.16 | 383.36 | 449.68 |
| **ENSBTAG00000047717** | FAM222B | -1.08 | 1.26E-04 | 2297.32 | 2808.40 | 2216.68 | 2400.00 | 6689.21 | 3900.45 | 5428.09 | 4494.02 |
| **ENSBTAG00000007634** | HOOK3 | -1.08 | 7.29E-04 | 766.11 | 771.82 | 851.94 | 1376.06 | 2045.51 | 2199.93 | 1706.84 | 1984.40 |
| **ENSBTAG00000015527** | MYO1D | -1.08 | 8.41E-05 | 362.68 | 328.35 | 461.67 | 337.15 | 597.82 | 1018.46 | 838.17 | 689.99 |
| **ENSBTAG00000019168** | PHOX2A | -1.07 | 4.63E-03 | 155.01 | 218.09 | 202.03 | 149.84 | 236.77 | 483.22 | 508.83 | 300.39 |
| **ENSBTAG00000020861** | CHAMP1 | -1.07 | 3.50E-05 | 1104.94 | 1352.31 | 751.33 | 1105.10 | 1935.11 | 2308.59 | 2756.74 | 2066.32 |
| **ENSBTAG00000002796** | KLHL3 | -1.07 | 3.13E-02 | 99.37 | 137.83 | 113.59 | 81.17 | 164.56 | 260.11 | 223.05 | 262.16 |
| **ENSBTAG00000000854** | SEC16B | -1.07 | 9.68E-03 | 210.65 | 469.42 | 243.41 | 383.35 | 915.83 | 567.61 | 460.91 | 794.67 |

| | | | | | | | | | | | |
|---|---|---|---|---|---|---|---|---|---|---|---|
| **ENSBTAG00000002846** | TRAF3IP3 | -1.07 | 2.87E-02 | 85.45 | 146.74 | 97.37 | 91.15 | 295.09 | 300.57 | 147.25 | 141.09 |
| **ENSBTAG00000033727** | RBPMS | -1.07 | 2.20E-03 | 1526.25 | 2733.82 | 2297.82 | 1577.10 | 3100.89 | 6068.00 | 4102.00 | 3781.28 |
| **ENSBTAG00000003865** | CASR | -1.06 | 1.25E-02 | 301.08 | 406.18 | 259.64 | 171.07 | 765.16 | 373.40 | 692.67 | 546.16 |
| **ENSBTAG00000000919** | HEY2 | -1.06 | 1.37E-02 | 139.11 | 284.57 | 241.79 | 153.59 | 436.74 | 301.72 | 466.14 | 506.11 |
| **ENSBTAG00000016730** | ANP32E | -1.06 | 4.68E-06 | 2093.62 | 2573.29 | 2559.08 | 2061.60 | 3634.84 | 5566.29 | 4760.69 | 5387.00 |
| **ENSBTAG00000013210** | ADAMTS4 | -1.06 | 2.27E-02 | 182.83 | 201.06 | 163.90 | 79.92 | 331.20 | 202.31 | 384.24 | 390.51 |
| **ENSBTAG00000025762** | CNP | -1.06 | 3.01E-02 | 241.46 | 184.04 | 158.22 | 275.96 | 383.27 | 357.21 | 434.77 | 609.88 |
| **ENSBTAG00000015817** | ELK1 | -1.05 | 3.24E-02 | 369.64 | 484.82 | 363.50 | 305.93 | 1487.26 | 326.00 | 706.61 | 642.65 |
| **ENSBTAG00000040093** | FOXI3 | -1.05 | 1.47E-02 | 787.97 | 693.18 | 877.10 | 807.91 | 2966.89 | 1032.34 | 1378.37 | 1191.55 |
| **ENSBTAG00000050347** | MAML1 | -1.05 | 4.09E-03 | 290.15 | 388.34 | 289.66 | 327.16 | 883.89 | 477.44 | 794.61 | 523.41 |
| **ENSBTAG00000001828** | SCRN2 | -1.05 | 2.67E-02 | 190.78 | 207.55 | 164.71 | 101.14 | 424.24 | 389.58 | 378.14 | 183.88 |
| **ENSBTAG00000055062** | SMAD5 | -1.05 | 2.96E-03 | 347.78 | 380.24 | 277.49 | 374.61 | 420.07 | 869.33 | 981.06 | 580.75 |
| **ENSBTAG00000014333** | SLC10A3 | -1.04 | 2.70E-03 | 457.08 | 379.43 | 400.82 | 463.27 | 1264.38 | 752.58 | 751.92 | 735.50 |
| **ENSBTAG00000012025** | LMX1A | -1.04 | 1.33E-02 | 43.72 | 75.40 | 77.08 | 63.68 | 122.90 | 129.48 | 189.94 | 94.67 |
| **ENSBTAG00000018902** | ZC2HC1A | -1.04 | 1.78E-05 | 3785.81 | 6183.51 | 4768.46 | 5599.16 | 7380.07 | 13468.91 | 12121.29 | 8897.93 |
| **ENSBTAG00000019810** | AARS | -1.04 | 3.62E-03 | 2454.32 | 3437.54 | 2369.22 | 2360.04 | 8543.77 | 4887.70 | 4075.86 | 4355.66 |
| **ENSBTAG00000026585** | ELOA | -1.04 | 2.96E-02 | 261.33 | 616.97 | 401.63 | 427.05 | 976.93 | 775.70 | 732.75 | 1025.88 |
| **ENSBTAG00000010542** | SPIRE1 | -1.04 | 5.81E-03 | 339.83 | 638.05 | 653.16 | 498.23 | 1294.93 | 897.08 | 1120.47 | 1063.20 |
| **ENSBTAG00000010324** | GTPBP1 | -1.04 | 5.50E-04 | 617.06 | 542.39 | 485.20 | 339.65 | 1349.09 | 938.70 | 940.99 | 847.47 |
| **ENSBTAG00000020853** | SLC16A13 | -1.04 | 2.33E-02 | 96.38 | 100.53 | 132.25 | 172.32 | 219.41 | 210.40 | 243.96 | 353.19 |
| **ENSBTAG00000009126** | YBX2 | -1.04 | 9.52E-04 | 500.80 | 678.59 | 438.95 | 445.78 | 1039.42 | 845.06 | 1322.61 | 1027.70 |
| **ENSBTAG00000016828** | TAPBP | -1.04 | 3.96E-02 | 365.66 | 402.94 | 398.39 | 279.71 | 745.02 | 435.82 | 563.72 | 1222.50 |
| **ENSBTAG00000021151** | MYH10 | -1.03 | 4.30E-02 | 289.15 | 449.96 | 501.43 | 345.89 | 793.62 | 1034.65 | 546.29 | 874.77 |
| **ENSBTAG00000017245** | COPG2 | -1.03 | 1.42E-02 | 319.96 | 521.31 | 491.69 | 494.48 | 636.70 | 1267.01 | 837.30 | 999.48 |
| **ENSBTAG00000018050** | CDR2 | -1.03 | 2.93E-02 | 604.14 | 616.97 | 557.42 | 599.37 | 2171.18 | 598.82 | 1202.37 | 887.52 |
| **ENSBTAG00000014471** | AKAP2 | -1.03 | 7.64E-03 | 245.43 | 431.31 | 279.11 | 196.05 | 622.82 | 500.56 | 609.90 | 618.99 |
| **ENSBTAG00000008991** | EIF4ENIF1 | -1.02 | 2.88E-04 | 835.66 | 1136.66 | 635.31 | 734.23 | 2087.86 | 1521.34 | 1579.64 | 1609.36 |
| **ENSBTAG00000009861** | FRS3 | -1.02 | 2.09E-02 | 156.00 | 119.99 | 142.80 | 87.41 | 290.93 | 260.11 | 264.00 | 213.91 |
| **ENSBTAG00000038541** | ZNF879 | -1.02 | 2.20E-02 | 121.23 | 235.93 | 266.94 | 178.56 | 322.17 | 362.99 | 405.15 | 537.97 |
| **ENSBTAG00000002121** | CDK2AP2 | -1.01 | 1.52E-07 | 3650.67 | 3840.48 | 3656.06 | 2842.03 | 4769.38 | 8337.29 | 8461.03 | 6694.16 |
| **ENSBTAG00000019517** | ELN | -1.01 | 4.05E-02 | 158.98 | 349.43 | 206.90 | 148.59 | 305.51 | 450.85 | 382.49 | 608.97 |
| **ENSBTAG00000017850** | ARHGEF6 | -1.01 | 3.55E-02 | 132.16 | 182.42 | 128.20 | 197.29 | 323.56 | 278.60 | 346.77 | 338.62 |
| **ENSBTAG00000007101** | F3 | -1.01 | 2.58E-03 | 871.43 | 1733.36 | 1068.58 | 880.33 | 1461.57 | 2410.32 | 3069.53 | 2232.90 |

| | | | | | | | | | | | |
|---|---|---|---|---|---|---|---|---|---|---|---|
| **ENSBTAG00000002378** | NUCB1 | -1.01 | 1.16E-02 | 610.10 | 479.96 | 672.63 | 483.25 | 1351.17 | 998.81 | 626.45 | 1545.65 |
| **ENSBTAG00000019290** | PACSIN2 | -1.01 | 2.55E-02 | 685.62 | 1806.33 | 1148.10 | 698.02 | 2413.50 | 2625.35 | 1836.66 | 1851.50 |
| **ENSBTAG00000045896** | NPTX2 | -1.01 | 5.39E-03 | 321.94 | 535.90 | 455.18 | 343.39 | 1017.89 | 522.53 | 965.38 | 824.71 |
| **ENSBTAG00000010309** | XPO6 | -1.00 | 1.42E-02 | 216.62 | 565.09 | 285.60 | 333.40 | 845.00 | 706.33 | 512.31 | 749.16 |
| **ENSBTAG00000004964** | PCGF5 | -1.00 | 1.04E-02 | 351.75 | 314.57 | 322.93 | 194.80 | 433.26 | 766.45 | 645.62 | 533.42 |
| **ENSBTAG00000018022** | SPINDOC | -1.00 | 2.25E-03 | 157.00 | 268.35 | 139.56 | 146.10 | 333.97 | 374.55 | 352.87 | 365.93 |
| **ENSBTAG00000020283** | DUSP5 | -1.00 | 1.29E-04 | 1161.58 | 985.05 | 782.98 | 724.24 | 1649.74 | 2300.50 | 1677.22 | 1685.83 |
| **ENSBTAG00000021570** | PLEKHG4 | -1.00 | 2.81E-02 | 1105.93 | 1357.99 | 946.06 | 1063.89 | 4315.98 | 1125.97 | 1512.55 | 1991.68 |
| **ENSBTAG00000038783** | | 1.00 | 4.79E-02 | 86.45 | 157.28 | 94.93 | 73.67 | 44.44 | 55.49 | 68.83 | 38.23 |
| **ENSBTAG00000052352** | | 1.00 | 8.41E-05 | 216.62 | 192.96 | 228.81 | 182.31 | 111.09 | 86.70 | 121.11 | 90.12 |
| **ENSBTAG00000052977** | | 1.02 | 1.39E-03 | 200.72 | 374.56 | 307.51 | 270.97 | 125.67 | 153.75 | 116.75 | 175.68 |
| **ENSBTAG00000021118** | CYP26A1 | 1.02 | 1.16E-02 | 842.62 | 477.53 | 1522.95 | 952.76 | 366.61 | 409.23 | 481.82 | 610.79 |
| **ENSBTAG00000013478** | MARVELD1 | 1.04 | 1.09E-02 | 4647.31 | 1939.29 | 3759.10 | 3791.04 | 2619.03 | 1786.07 | 1279.91 | 1192.46 |
| **ENSBTAG00000023659** | MT2A | 1.05 | 5.08E-03 | 32265.84 | 22089.42 | 38083.55 | 32840.73 | 8367.41 | 16043.39 | 23205.75 | 12932.26 |
| **ENSBTAG00000052047** | | 1.06 | 2.42E-03 | 395.47 | 461.31 | 288.04 | 332.15 | 146.50 | 198.84 | 153.35 | 213.00 |
| **ENSBTAG00000046022** | PYY | 1.07 | 1.04E-03 | 4153.46 | 3322.41 | 3319.34 | 3234.13 | 1692.09 | 1513.24 | 1814.01 | 1670.35 |
| **ENSBTAG00000048743** | | 1.09 | 4.06E-02 | 249.41 | 176.74 | 140.37 | 382.10 | 144.42 | 113.29 | 90.61 | 94.67 |
| **ENSBTAG00000016438** | TERB1 | 1.10 | 5.69E-06 | 11375.32 | 7366.38 | 8938.12 | 11681.56 | 3746.62 | 5573.22 | 4692.73 | 4342.92 |
| **ENSBTAG00000038706** | MT1E | 1.10 | 3.67E-02 | 734.31 | 344.56 | 546.87 | 518.21 | 136.09 | 255.48 | 358.10 | 247.59 |
| **ENSBTAG00000047865** | RF01164 | 1.12 | 1.09E-04 | 359.70 | 271.60 | 365.12 | 359.62 | 123.59 | 186.12 | 129.82 | 186.61 |
| **ENSBTAG00000003027** | EMX2 | 1.13 | 5.09E-05 | 529.62 | 372.94 | 490.88 | 536.94 | 191.64 | 208.09 | 184.71 | 295.84 |
| **ENSBTAG00000009252** | KLRA1 | 1.14 | 1.21E-02 | 575.32 | 266.73 | 360.25 | 458.27 | 193.02 | 99.42 | 196.91 | 260.34 |
| **ENSBTAG00000050728** | | 1.16 | 3.44E-02 | 88.43 | 98.10 | 135.50 | 118.63 | 68.74 | 53.18 | 37.47 | 37.32 |
| **ENSBTAG00000052759** | | 1.16 | 3.71E-03 | 514.71 | 262.68 | 309.95 | 453.28 | 205.52 | 161.84 | 127.21 | 192.98 |
| **ENSBTAG00000001043** | LUZP2 | 1.16 | 1.04E-02 | 410.38 | 428.07 | 399.20 | 332.15 | 103.46 | 159.53 | 179.48 | 259.43 |
| **ENSBTAG00000051842** | | 1.19 | 3.84E-02 | 93.40 | 117.56 | 77.08 | 56.19 | 33.33 | 31.21 | 33.98 | 52.80 |
| **ENSBTAG00000054655** | | 1.19 | 3.82E-02 | 46.70 | 59.18 | 30.02 | 61.19 | 16.66 | 24.28 | 17.43 | 28.22 |
| **ENSBTAG00000002956** | ZNF674 | 1.22 | 8.50E-04 | 840.63 | 569.14 | 391.08 | 771.69 | 229.82 | 322.53 | 363.32 | 192.07 |
| **ENSBTAG00000007268** | F13A1 | 1.22 | 2.03E-02 | 509.74 | 266.73 | 406.50 | 358.38 | 97.90 | 277.45 | 157.70 | 131.99 |
| **ENSBTAG00000038900** | | 1.24 | 2.64E-02 | 46.70 | 55.13 | 46.25 | 84.91 | 26.38 | 16.18 | 25.27 | 30.04 |
| **ENSBTAG00000040206** | ZNF770 | 1.26 | 5.26E-04 | 597.18 | 360.78 | 362.69 | 299.69 | 184.69 | 152.60 | 224.79 | 115.60 |
| **ENSBTAG00000006806** | KRT17 | 1.26 | 1.05E-02 | 802.87 | 1425.28 | 392.71 | 422.06 | 204.13 | 413.86 | 363.32 | 286.74 |
| **ENSBTAG00000005749** | | 1.28 | 4.56E-02 | 161.97 | 179.17 | 133.88 | 126.12 | 28.47 | 57.80 | 96.71 | 65.54 |

| | | | | | | | | | | | |
|---|---|---|---|---|---|---|---|---|---|---|---|
| **ENSBTAG00000001877** | HMGN5 | 1.28 | 2.66E-02 | 206.68 | 522.12 | 377.29 | 467.01 | 61.10 | 282.07 | 164.67 | 141.09 |
| **ENSBTAG00000054034** | | 1.32 | 1.10E-02 | 1878.00 | 3023.25 | 2249.95 | 1071.38 | 470.76 | 541.02 | 1240.71 | 1045.90 |
| **ENSBTAG00000005330** | KRTDAP | 1.35 | 2.19E-02 | 514.71 | 736.15 | 391.08 | 934.03 | 306.20 | 397.67 | 107.17 | 198.44 |
| **ENSBTAG00000043258** | RF00425 | 1.37 | 1.50E-02 | 173.89 | 173.50 | 155.78 | 192.30 | 49.99 | 49.71 | 64.47 | 105.59 |
| **ENSBTAG00000035643** | | 1.37 | 7.20E-03 | 620.04 | 489.69 | 278.30 | 511.97 | 115.26 | 276.29 | 170.77 | 172.04 |
| **ENSBTAG00000021098** | OR2T29 | 1.38 | 1.62E-03 | 133.15 | 209.98 | 172.82 | 133.61 | 65.27 | 48.55 | 55.76 | 80.10 |
| **ENSBTAG00000027696** | TTC30B | 1.38 | 2.96E-02 | 151.03 | 156.47 | 335.10 | 288.45 | 77.77 | 121.38 | 47.92 | 111.05 |
| **ENSBTAG00000032299** | CCDC172 | 1.38 | 4.83E-04 | 1120.84 | 734.53 | 777.30 | 1937.98 | 254.13 | 462.41 | 419.96 | 617.17 |
| **ENSBTAG00000049228** | | 1.46 | 2.83E-02 | 65.58 | 138.64 | 54.36 | 84.91 | 43.74 | 39.31 | 18.30 | 23.67 |
| **ENSBTAG00000009848** | GPR171 | 1.47 | 9.75E-03 | 505.77 | 630.76 | 346.46 | 146.10 | 153.45 | 176.87 | 141.15 | 117.43 |
| **ENSBTAG00000044441** | RF00553 | 1.47 | 1.05E-02 | 141.10 | 62.43 | 141.18 | 128.62 | 54.16 | 26.59 | 30.49 | 58.26 |
| **ENSBTAG00000043222** | RF00586 | 1.48 | 7.01E-05 | 238.48 | 141.88 | 158.22 | 218.52 | 79.85 | 40.46 | 67.96 | 81.01 |
| **ENSBTAG00000004817** | | 1.56 | 1.36E-02 | 259.34 | 222.14 | 74.65 | 103.64 | 56.24 | 67.05 | 48.79 | 51.89 |
| **ENSBTAG00000009664** | SP140L | 1.64 | 2.82E-02 | 205.69 | 423.21 | 103.04 | 78.67 | 43.05 | 79.77 | 74.06 | 64.63 |
| **ENSBTAG00000008338** | PLCB1 | 1.95 | 3.63E-02 | 99.37 | 91.61 | 32.46 | 42.46 | 31.25 | 13.87 | 9.58 | 13.65 |
| **ENSBTAG00000047719** | DRD1 | 2.40 | 2.86E-02 | 32.79 | 105.40 | 127.39 | 71.18 | 20.14 | 16.18 | 17.43 | 10.01 |
| **ENSBTAG00000047412** | TNFSF18 | 2.59 | 3.86E-02 | 41.73 | 178.36 | 182.56 | 2.50 | 4.17 | 18.50 | 29.62 | 15.47 |
| **ENSBTAG00000007135** | REG3G | 2.98 | 1.22E-02 | 34.78 | 225.39 | 146.05 | 56.19 | 20.83 | 10.40 | 27.01 | 0.00 |
| **ENSBTAG00000050086** | | 2.99 | 7.44E-03 | 20.87 | 17.03 | 60.04 | 8.74 | 2.08 | 10.40 | 1.74 | 0.00 |
| **ENSBTAG00000042363** | RF00425 | 3.28 | 4.93E-02 | 20.87 | 31.62 | 4.06 | 18.73 | 2.08 | 1.16 | 2.61 | 1.82 |
| **ENSBTAG00000000620** | MCMDC2 | 3.49 | 1.58E-02 | 36.77 | 35.67 | 8.11 | 46.20 | 9.72 | 0.00 | 0.00 | 0.91 |
| **ENSBTAG00000054257** | bta-mir-2887-2 | 3.51 | 2.96E-03 | 89.43 | 855.33 | 263.70 | 47.45 | 9.03 | 86.70 | 7.84 | 7.28 |
| **ENSBTAG00000000575** | TNC | 4.06 | 2.14E-02 | 10.93 | 450.77 | 47.06 | 19.98 | 5.55 | 8.09 | 0.00 | 18.21 |
| **ENSBTAG00000029994** | MIRLET7I | 4.61 | 1.79E-02 | 22.85 | 5.68 | 17.85 | 13.74 | 1.39 | 0.00 | 0.87 | 0.00 |
| **ENSBTAG00000038496** | CR2 | 6.17 | 4.70E-02 | 22.85 | 0.81 | 23.53 | 0.00 | 0.00 | 0.00 | 0.00 | 0.00 |
| **ENSBTAG00000025952** | OBP | 6.21 | 1.63E-02 | 52.66 | 13.78 | 0.00 | 13.74 | 0.00 | 1.16 | 0.00 | 0.00 |
