## Supplementary material 7: DEG_Comparison_SSB_vs_ADB for "Bovine in vitro blastocysts with distinct morphokinetic patterns show transcriptomic differences at genome activation"

| DEG_comparison_SSB_vs_ADB | | | | | | | | | | | |
|---|---|---|---|---|---|---|---|---|---|---|---|
| **Geneid** | **Gene name** | **log2FC** | **padj** | **SSB-1** | **SSB-2** | **SSB-3** | **SSB-4** | **ADB-1** | **ADB-2** | **ADB-3** | **ADB-4** |
| **ENSBTAG00000004547** | OLR1 | -26.40 | 1.20E-06 | 0.00 | 0.00 | 0.00 | 0.00 | 0.00 | 0.00 | 32.24 | 2978.42 |
| **ENSBTAG00000007196** | TAGLN | -23.98 | 4.38E-06 | 0.00 | 0.00 | 0.00 | 0.00 | 6.25 | 0.00 | 0.00 | 616.26 |
| **ENSBTAG00000018563** | SFRP2 | -23.06 | 2.17E-10 | 0.00 | 0.00 | 0.00 | 0.00 | 2.08 | 0.00 | 13.07 | 328.61 |
| **ENSBTAG00000007881** | IFIT1 | -20.62 | 3.78E-04 | 0.00 | 0.00 | 0.00 | 0.00 | 7.64 | 0.00 | 0.00 | 46.42 |
| **ENSBTAG00000004476** | ADHFE1 | -7.87 | 4.37E-03 | 0.00 | 0.00 | 0.00 | 0.00 | 0.00 | 50.87 | 128.08 | 0.00 |
| **ENSBTAG00000039556** | WIPI1 | -7.00 | 4.25E-02 | 0.00 | 0.00 | 0.00 | 0.00 | 29.86 | 0.00 | 0.00 | 68.27 |
| **ENSBTAG00000047749** | | -6.72 | 1.71E-02 | 0.85 | 0.00 | 0.00 | 0.00 | 10.42 | 50.87 | 59.25 | 12.74 |
| **ENSBTAG00000006451** | GAP43 | -6.63 | 2.28E-02 | 0.00 | 0.00 | 0.00 | 0.00 | 9.72 | 17.34 | 12.20 | 36.41 |
| **ENSBTAG00000000951** | JAKMIP3 | -6.34 | 4.24E-02 | 0.00 | 1.00 | 0.00 | 0.00 | 6.94 | 35.84 | 14.81 | 44.60 |
| **ENSBTAG00000015459** | KCNA2 | -5.80 | 2.31E-03 | 2.55 | 0.00 | 2.41 | 0.00 | 113.18 | 104.04 | 45.31 | 16.38 |
| **ENSBTAG00000013196** | STK17B | -5.62 | 3.39E-02 | 0.00 | 3.01 | 0.00 | 0.00 | 36.80 | 16.18 | 54.89 | 42.78 |
| **ENSBTAG00000009337** | PNMA8A | -4.95 | 1.05E-02 | 3.40 | 3.01 | 0.00 | 0.00 | 35.41 | 13.87 | 94.97 | 61.90 |
| **ENSBTAG00000024791** | | -4.74 | 2.72E-02 | 5.95 | 1.00 | 2.41 | 2.28 | 68.04 | 4.62 | 22.65 | 219.38 |
| **ENSBTAG00000055051** | RNF125 | -4.61 | 1.55E-04 | 0.00 | 0.00 | 8.42 | 0.00 | 22.22 | 91.33 | 50.53 | 27.31 |
| **ENSBTAG00000037921** | OVCH2 | -4.41 | 3.48E-02 | 0.00 | 0.00 | 0.00 | 6.85 | 18.05 | 26.59 | 36.59 | 60.08 |
| **ENSBTAG00000054147** | | -4.08 | 8.09E-04 | 16.14 | 12.05 | 63.75 | 19.39 | 47.21 | 45.09 | 247.44 | 1541.09 |
| **ENSBTAG00000013632** | GRM4 | -3.67 | 1.04E-02 | 5.10 | 6.03 | 3.61 | 0.00 | 45.13 | 39.31 | 67.09 | 38.23 |
| **ENSBTAG00000001010** | ADAMTS18 | -3.60 | 2.85E-02 | 1.70 | 9.04 | 20.45 | 0.00 | 111.79 | 71.67 | 130.69 | 59.17 |
| **ENSBTAG00000014589** | CLDN18 | -3.58 | 4.25E-02 | 3.40 | 2.01 | 3.61 | 0.00 | 6.25 | 45.09 | 27.01 | 30.95 |
| **ENSBTAG00000040202** | | -3.57 | 4.78E-05 | 55.21 | 14.06 | 30.07 | 5.70 | 205.52 | 223.11 | 249.19 | 576.20 |
| **ENSBTAG00000021447** | SIM2 | -3.31 | 1.15E-02 | 1.70 | 3.01 | 6.01 | 11.41 | 106.23 | 26.59 | 33.98 | 48.24 |
| **ENSBTAG00000018133** | SEMA3A | -3.10 | 2.31E-03 | 12.74 | 17.07 | 3.61 | 6.85 | 42.35 | 102.89 | 54.89 | 149.28 |
| **ENSBTAG00000032517** | BCL7A | -3.04 | 2.72E-02 | 22.93 | 7.03 | 0.00 | 6.85 | 68.04 | 72.83 | 78.42 | 87.39 |
| **ENSBTAG00000033330** | HOXD11 | -2.99 | 1.60E-03 | 4.25 | 34.15 | 8.42 | 4.56 | 128.45 | 129.48 | 44.44 | 106.50 |
| **ENSBTAG00000048635** | | -2.87 | 4.07E-02 | 4.25 | 5.02 | 4.81 | 3.42 | 13.19 | 26.59 | 42.69 | 45.51 |
| **ENSBTAG00000012629** | ZNF362 | -2.84 | 1.40E-02 | 41.62 | 48.21 | 2.41 | 7.99 | 132.62 | 105.20 | 253.54 | 233.03 |
| **ENSBTAG00000020520** | RASD1 | -2.63 | 1.24E-03 | 8.49 | 81.35 | 33.68 | 23.96 | 163.17 | 158.38 | 417.34 | 173.86 |
| **ENSBTAG00000013920** | | -2.59 | 3.42E-02 | 58.60 | 66.29 | 20.45 | 17.11 | 127.76 | 375.71 | 238.73 | 239.40 |
| **ENSBTAG00000048560** | TMEM233 | -2.58 | 7.12E-03 | 28.03 | 48.21 | 32.48 | 25.10 | 141.64 | 278.60 | 156.83 | 225.75 |
| **ENSBTAG00000034366** | RGS2 | -2.52 | 1.20E-02 | 1915.20 | 3482.08 | 1015.21 | 2380.97 | 7097.48 | 17061.85 | 14700.28 | 11699.75 |
| **ENSBTAG00000015402** | GREB1 | -2.45 | 4.06E-03 | 13.59 | 5.02 | 8.42 | 21.68 | 85.40 | 70.52 | 54.02 | 55.53 |

| | | | | | | | | | | | |
|---|---|---|---|---|---|---|---|---|---|---|---|
| **ENSBTAG00000034346** | BTG4 | -2.43 | 3.55E-02 | 5447.50 | 8049.87 | 5850.69 | 8088.68 | 16773.01 | 58927.49 | 30122.86 | 42014.61 |
| **ENSBTAG00000006188** | USH2A | -2.41 | 7.40E-05 | 95.12 | 168.73 | 105.85 | 93.55 | 379.80 | 664.72 | 598.57 | 821.98 |
| **ENSBTAG00000019580** | TCL1A | -2.40 | 1.08E-02 | 39.92 | 70.30 | 20.45 | 36.51 | 121.51 | 210.40 | 158.57 | 395.97 |
| **ENSBTAG00000037964** | HIST3H2A | -2.35 | 4.25E-02 | 10.19 | 116.50 | 98.63 | 118.65 | 52.08 | 139.88 | 1004.59 | 548.90 |
| **ENSBTAG00000002908** | GNG4 | -2.32 | 8.09E-04 | 66.25 | 72.31 | 37.29 | 123.21 | 182.61 | 361.84 | 400.79 | 544.34 |
| **ENSBTAG00000016958** | PHF19 | -2.30 | 2.49E-02 | 9.34 | 9.04 | 4.81 | 12.55 | 57.63 | 23.12 | 35.72 | 60.08 |
| **ENSBTAG00000048053** | SLBP2 | -2.28 | 5.74E-03 | 792.41 | 1068.63 | 501.59 | 734.71 | 2447.53 | 4520.08 | 4087.19 | 3996.10 |
| **ENSBTAG00000003827** | RIMS2 | -2.27 | 1.95E-03 | 169.01 | 306.33 | 289.89 | 162.00 | 731.83 | 1396.48 | 1247.68 | 1104.16 |
| **ENSBTAG00000026972** | MYF5 | -2.27 | 1.77E-05 | 536.77 | 1062.60 | 761.41 | 788.33 | 1991.35 | 4783.65 | 4497.56 | 3931.47 |
| **ENSBTAG00000005230** | SHOX2 | -2.24 | 1.76E-02 | 77.29 | 67.29 | 33.68 | 33.08 | 229.13 | 208.09 | 277.07 | 284.92 |
| **ENSBTAG00000030425** | ID3 | -2.23 | 2.10E-02 | 849.31 | 2762.97 | 956.27 | 2346.75 | 3149.50 | 11844.69 | 7628.95 | 9749.03 |
| **ENSBTAG00000033319** | CD200 | -2.20 | 3.78E-04 | 116.36 | 101.44 | 103.45 | 115.23 | 232.60 | 631.19 | 528.87 | 616.26 |
| **ENSBTAG00000006095** | PRDM13 | -2.20 | 3.78E-04 | 50.11 | 112.49 | 54.13 | 44.49 | 180.53 | 387.27 | 372.91 | 262.16 |
| **ENSBTAG00000001812** | H1FOO | -2.18 | 3.47E-02 | 524.88 | 930.03 | 425.81 | 1116.90 | 2125.35 | 4488.87 | 4000.06 | 2979.33 |
| **ENSBTAG00000009478** | GDF9 | -2.15 | 1.07E-02 | 338.03 | 751.25 | 336.80 | 754.11 | 1496.98 | 3499.30 | 1931.63 | 2772.69 |
| **ENSBTAG00000053831** | | -2.15 | 2.33E-02 | 45.86 | 112.49 | 91.42 | 132.34 | 120.12 | 426.58 | 676.12 | 473.34 |
| **ENSBTAG00000019024** | JAZF1 | -2.15 | 1.95E-03 | 74.74 | 112.49 | 60.14 | 68.45 | 337.45 | 375.71 | 385.98 | 304.03 |
| **ENSBTAG00000033089** | HACD1 | -2.15 | 8.38E-03 | 32.27 | 56.24 | 22.85 | 34.23 | 110.40 | 206.93 | 105.43 | 224.84 |
| **ENSBTAG00000038195** | CD302 | -2.14 | 1.59E-02 | 61.15 | 14.06 | 10.83 | 14.83 | 66.66 | 78.61 | 147.25 | 158.39 |
| **ENSBTAG00000017205** | WISP3 | -2.14 | 3.46E-02 | 79.84 | 42.18 | 52.93 | 13.69 | 203.44 | 204.62 | 189.07 | 237.58 |
| **ENSBTAG00000007767** | TBX15 | -2.13 | 3.17E-02 | 21.23 | 26.11 | 6.01 | 50.20 | 67.35 | 84.39 | 174.26 | 127.44 |
| **ENSBTAG00000044125** | TLCD4 | -2.13 | 1.65E-02 | 140.99 | 171.74 | 42.10 | 301.19 | 373.55 | 1132.91 | 573.30 | 783.75 |
| **ENSBTAG00000031246** | CCNI2 | -2.12 | 3.99E-04 | 200.44 | 95.41 | 96.23 | 110.66 | 377.72 | 649.69 | 639.52 | 529.78 |
| **ENSBTAG00000015952** | ADAP1 | -2.11 | 1.20E-06 | 176.66 | 114.50 | 87.81 | 114.09 | 563.80 | 463.57 | 568.08 | 544.34 |
| **ENSBTAG00000008916** | CYFIP2 | -2.11 | 1.49E-02 | 264.99 | 173.75 | 66.16 | 44.49 | 585.32 | 575.70 | 491.40 | 727.31 |
| **ENSBTAG00000000830** | GPR137B | -2.10 | 9.71E-03 | 284.52 | 233.01 | 184.04 | 285.21 | 775.57 | 1102.85 | 1150.09 | 1195.19 |
| **ENSBTAG00000046744** | PALM3 | -2.09 | 4.25E-02 | 27.18 | 44.19 | 31.27 | 33.08 | 120.12 | 172.25 | 94.10 | 191.16 |
| **ENSBTAG00000048664** | CDK14 | -2.08 | 4.46E-02 | 35.67 | 43.19 | 3.61 | 51.34 | 68.74 | 182.65 | 140.28 | 177.50 |
| **ENSBTAG00000033015** | FAHD1 | -2.08 | 7.12E-03 | 46.71 | 53.23 | 32.48 | 37.65 | 58.32 | 201.15 | 136.79 | 323.15 |
| **ENSBTAG00000007403** | SLC7A3 | -2.07 | 9.63E-03 | 196.19 | 248.07 | 87.81 | 224.75 | 583.24 | 1164.12 | 681.34 | 756.44 |
| **ENSBTAG00000026384** | ZAR1L | -2.07 | 2.81E-02 | 1925.40 | 1931.37 | 1853.60 | 3128.23 | 4869.36 | 11211.18 | 10930.24 | 10103.13 |
| **ENSBTAG00000019164** | RHOBTB1 | -2.06 | 1.48E-02 | 30.58 | 43.19 | 10.83 | 12.55 | 47.91 | 95.95 | 144.63 | 121.07 |
| **ENSBTAG00000019964** | GAS6 | -2.02 | 1.28E-02 | 49.26 | 19.08 | 24.06 | 26.24 | 86.79 | 139.88 | 208.24 | 50.07 |

| | | | | | | | | | | | |
|---|---|---|---|---|---|---|---|---|---|---|---|
| **ENSBTAG00000036087** | ARMC2 | -2.02 | 3.75E-03 | 280.27 | 415.80 | 209.30 | 386.75 | 841.53 | 1205.74 | 1526.49 | 1676.72 |
| **ENSBTAG00000008091** | SELENBP1 | -2.01 | 3.22E-04 | 218.27 | 136.59 | 72.17 | 139.18 | 412.43 | 582.64 | 633.42 | 656.31 |
| **ENSBTAG00000013274** | CENPV | -1.99 | 1.24E-03 | 2210.77 | 4062.60 | 1530.03 | 2992.47 | 6068.47 | 14260.79 | 10654.92 | 11768.93 |
| **ENSBTAG00000004710** | HNF1B | -1.98 | 2.49E-02 | 69.64 | 78.34 | 42.10 | 50.20 | 148.59 | 317.91 | 152.47 | 331.34 |
| **ENSBTAG00000002115** | C13H20orf96 | -1.97 | 1.10E-03 | 84.93 | 168.73 | 56.53 | 69.59 | 446.46 | 270.51 | 480.95 | 286.74 |
| **ENSBTAG00000001729** | DUSP10 | -1.96 | 6.37E-03 | 291.31 | 605.62 | 256.21 | 252.13 | 1251.88 | 1419.61 | 1614.49 | 1180.63 |
| **ENSBTAG00000002929** | IRF4 | -1.96 | 3.60E-03 | 32.27 | 30.13 | 16.84 | 30.80 | 123.59 | 65.89 | 152.47 | 85.57 |
| **ENSBTAG00000022580** | INKA2 | -1.95 | 1.97E-03 | 100.22 | 93.40 | 110.66 | 55.90 | 358.28 | 476.28 | 250.06 | 309.49 |
| **ENSBTAG00000053612** | | -1.93 | 3.61E-02 | 175.81 | 78.34 | 49.32 | 101.54 | 335.36 | 84.39 | 449.58 | 679.06 |
| **ENSBTAG00000053893** | | -1.93 | 4.46E-02 | 1811.59 | 1854.03 | 732.54 | 1192.20 | 5685.20 | 4277.31 | 6288.92 | 5036.55 |
| **ENSBTAG00000048279** | BCAR4 | -1.91 | 1.19E-03 | 221.67 | 307.33 | 262.22 | 359.37 | 601.99 | 1396.48 | 1165.78 | 1174.25 |
| **ENSBTAG00000008078** | TRIM59 | -1.91 | 7.23E-03 | 202.99 | 414.80 | 226.14 | 263.54 | 384.66 | 1780.29 | 750.17 | 1260.73 |
| **ENSBTAG00000020796** | UBE2D1 | -1.91 | 1.75E-02 | 168.16 | 170.74 | 105.85 | 188.24 | 209.69 | 879.74 | 566.33 | 730.04 |
| **ENSBTAG00000038815** | | -1.91 | 8.02E-03 | 46.71 | 56.24 | 19.25 | 85.56 | 133.31 | 246.23 | 240.47 | 161.12 |
| **ENSBTAG00000020346** | CCDC92 | -1.89 | 4.92E-04 | 58.60 | 77.33 | 14.43 | 46.78 | 197.19 | 169.94 | 196.04 | 170.22 |
| **ENSBTAG00000001322** | SAXO1 | -1.88 | 2.22E-02 | 2046.00 | 1916.30 | 971.91 | 2068.38 | 4121.56 | 8569.65 | 5602.35 | 7514.31 |
| **ENSBTAG00000037508** | EBF1 | -1.86 | 3.38E-02 | 67.95 | 198.86 | 134.72 | 58.18 | 300.65 | 396.52 | 476.59 | 496.10 |
| **ENSBTAG00000001080** | SPAG17 | -1.86 | 1.14E-03 | 293.01 | 299.30 | 215.31 | 412.99 | 769.32 | 1595.32 | 792.87 | 1275.29 |
| **ENSBTAG00000019402** | CFAP100 | -1.86 | 5.09E-03 | 20.38 | 42.18 | 33.68 | 29.66 | 163.17 | 64.74 | 148.12 | 77.37 |
| **ENSBTAG00000012389** | | -1.84 | 2.31E-03 | 366.90 | 497.15 | 173.21 | 318.30 | 1852.48 | 543.33 | 1132.67 | 1330.82 |
| **ENSBTAG00000003103** | EIF4E1B | -1.84 | 1.33E-02 | 1204.33 | 1484.43 | 716.90 | 990.26 | 2664.85 | 4269.22 | 3690.75 | 5072.96 |
| **ENSBTAG00000050426** | MVB12B | -1.84 | 1.38E-02 | 41.62 | 51.22 | 25.26 | 14.83 | 93.74 | 142.19 | 98.45 | 142.91 |
| **ENSBTAG00000004886** | ZAR1 | -1.82 | 9.57E-03 | 397.48 | 234.01 | 115.47 | 188.24 | 540.89 | 833.50 | 843.40 | 1099.61 |
| **ENSBTAG00000018394** | SDR42E1 | -1.82 | 3.95E-06 | 195.34 | 211.92 | 161.18 | 99.25 | 422.85 | 751.42 | 488.79 | 692.72 |
| **ENSBTAG00000006961** | NLRP13 | -1.82 | 3.59E-02 | 48.41 | 51.22 | 68.56 | 31.94 | 257.60 | 184.96 | 185.58 | 75.55 |
| **ENSBTAG00000006844** | LEF1 | -1.81 | 1.19E-02 | 472.22 | 514.23 | 344.02 | 325.14 | 1042.19 | 910.95 | 2053.61 | 1816.00 |
| **ENSBTAG00000053278** | | -1.81 | 4.84E-02 | 1746.19 | 1493.47 | 905.75 | 1213.87 | 3188.38 | 5628.71 | 5698.19 | 4287.39 |
| **ENSBTAG00000013249** | SALL2 | -1.81 | 1.33E-02 | 50.11 | 73.32 | 44.51 | 67.31 | 366.61 | 213.87 | 124.59 | 117.43 |
| **ENSBTAG00000054813** | | -1.79 | 1.10E-03 | 225.92 | 176.77 | 116.68 | 175.69 | 411.05 | 698.24 | 700.51 | 601.69 |
| **ENSBTAG00000010875** | MSX1 | -1.79 | 3.95E-06 | 1643.42 | 1532.64 | 935.82 | 1302.86 | 4063.24 | 5272.65 | 5053.44 | 4325.62 |
| **ENSBTAG00000010597** | GGCT | -1.79 | 7.70E-03 | 378.79 | 509.21 | 488.36 | 958.32 | 958.87 | 2762.91 | 2027.48 | 2301.17 |
| **ENSBTAG00000045526** | INSM1 | -1.77 | 1.82E-02 | 18.68 | 73.32 | 49.32 | 34.23 | 104.84 | 120.23 | 199.52 | 173.86 |
| **ENSBTAG00000054434** | | -1.76 | 3.07E-02 | 153.73 | 186.81 | 97.43 | 102.68 | 261.07 | 647.38 | 451.32 | 478.80 |

| | | | | | | | | | | | |
|---|---|---|---|---|---|---|---|---|---|---|---|
| **ENSBTAG00000052516** | CDA | -1.75 | 1.77E-05 | 89.18 | 111.48 | 90.21 | 74.16 | 377.02 | 285.54 | 280.55 | 281.27 |
| **ENSBTAG00000023666** | | -1.74 | 6.72E-04 | 258.19 | 287.24 | 269.44 | 179.11 | 665.17 | 950.26 | 653.46 | 1043.17 |
| **ENSBTAG00000004791** | VPS26C | -1.73 | 3.49E-03 | 191.95 | 187.81 | 135.92 | 227.03 | 492.98 | 567.61 | 816.39 | 588.04 |
| **ENSBTAG00000018845** | DND1 | -1.73 | 4.37E-04 | 63.70 | 127.55 | 151.56 | 91.27 | 333.28 | 387.27 | 441.74 | 273.99 |
| **ENSBTAG00000002289** | NLRP14 | -1.73 | 7.00E-03 | 642.08 | 415.80 | 445.06 | 286.36 | 1383.81 | 2100.51 | 1307.80 | 1131.47 |
| **ENSBTAG00000012012** | CYB5A | -1.72 | 4.16E-04 | 404.27 | 326.41 | 329.58 | 403.86 | 638.79 | 1369.90 | 1270.33 | 1556.57 |
| **ENSBTAG00000011733** | GIPC2 | -1.72 | 1.28E-02 | 225.92 | 215.94 | 187.65 | 181.40 | 510.34 | 1175.68 | 649.98 | 328.61 |
| **ENSBTAG00000017824** | IRF8 | -1.71 | 3.78E-04 | 680.30 | 734.18 | 223.73 | 433.53 | 1421.99 | 1935.19 | 1938.60 | 1494.67 |
| **ENSBTAG00000001414** | KCTD12 | -1.70 | 4.27E-02 | 28.03 | 81.35 | 18.04 | 85.56 | 247.18 | 105.20 | 179.48 | 158.39 |
| **ENSBTAG00000009341** | CCDC146 | -1.70 | 1.75E-02 | 21.23 | 35.15 | 38.49 | 35.37 | 80.54 | 122.54 | 120.24 | 97.40 |
| **ENSBTAG00000018600** | VGLL4 | -1.70 | 7.37E-05 | 375.40 | 472.04 | 310.34 | 302.33 | 1000.53 | 1308.63 | 1413.22 | 1008.58 |
| **ENSBTAG00000016407** | IRX6 | -1.68 | 2.57E-02 | 30.58 | 84.37 | 50.52 | 22.82 | 202.05 | 77.45 | 199.52 | 123.80 |
| **ENSBTAG00000011738** | TFR2 | -1.68 | 7.30E-03 | 22.08 | 15.07 | 7.22 | 17.11 | 39.58 | 41.62 | 52.28 | 65.54 |
| **ENSBTAG00000019675** | ATXN1 | -1.68 | 4.27E-02 | 118.05 | 111.48 | 55.33 | 69.59 | 222.88 | 379.18 | 224.79 | 307.67 |
| **ENSBTAG00000030721** | UNCX | -1.66 | 1.96E-04 | 318.49 | 473.05 | 282.67 | 365.07 | 1047.75 | 1055.46 | 1096.07 | 1350.85 |
| **ENSBTAG00000002733** | DBX1 | -1.65 | 1.42E-02 | 202.14 | 392.70 | 182.83 | 363.93 | 798.48 | 765.29 | 1004.59 | 1007.67 |
| **ENSBTAG00000000623** | CPEB1 | -1.65 | 7.40E-03 | 6756.30 | 6351.51 | 4809.02 | 6629.53 | 11675.22 | 22879.00 | 22287.42 | 20046.96 |
| **ENSBTAG00000039720** | TMEM225B | -1.65 | 6.72E-04 | 574.14 | 676.93 | 239.37 | 585.26 | 1037.33 | 2206.86 | 1697.26 | 1561.12 |
| **ENSBTAG00000007439** | TENM4 | -1.62 | 1.32E-03 | 77.29 | 68.30 | 38.49 | 53.62 | 193.72 | 153.75 | 212.59 | 173.86 |
| **ENSBTAG00000052473** | CNTNAP2 | -1.60 | 3.70E-02 | 1063.34 | 1150.99 | 1321.94 | 1273.20 | 2681.52 | 5034.51 | 2980.66 | 3898.70 |
| **ENSBTAG00000012010** | PANX1 | -1.60 | 4.09E-02 | 135.04 | 39.17 | 104.65 | 88.99 | 356.89 | 298.26 | 250.93 | 209.36 |
| **ENSBTAG00000017354** | MPP7 | -1.59 | 3.17E-02 | 802.60 | 673.92 | 345.22 | 256.69 | 1105.38 | 2036.93 | 1690.29 | 1448.25 |
| **ENSBTAG00000004561** | PAX6 | -1.59 | 1.09E-03 | 36.52 | 43.19 | 38.49 | 53.62 | 113.18 | 76.30 | 155.09 | 172.04 |
| **ENSBTAG00000021487** | CIART | -1.58 | 2.40E-03 | 765.23 | 1410.11 | 925.00 | 1270.92 | 2614.86 | 3714.33 | 3723.86 | 3053.97 |
| **ENSBTAG00000009861** | FRS3 | -1.58 | 4.45E-04 | 64.55 | 134.58 | 72.17 | 73.01 | 290.93 | 260.11 | 264.00 | 213.91 |
| **ENSBTAG00000018446** | GCA | -1.58 | 1.29E-02 | 347.37 | 317.37 | 340.41 | 436.95 | 892.91 | 1218.46 | 1137.90 | 1058.65 |
| **ENSBTAG00000011851** | FYN | -1.57 | 4.56E-02 | 369.45 | 316.37 | 86.61 | 115.23 | 345.08 | 608.07 | 642.14 | 1046.82 |
| **ENSBTAG00000004115** | MYLIP | -1.56 | 2.26E-05 | 931.70 | 1056.58 | 588.20 | 893.29 | 1658.07 | 2705.11 | 3499.07 | 2381.28 |
| **ENSBTAG00000046348** | ELAVL2 | -1.56 | 5.09E-03 | 517.23 | 713.09 | 344.02 | 490.57 | 879.03 | 1891.27 | 1844.51 | 1468.27 |
| **ENSBTAG00000007306** | RAB3C | -1.55 | 2.40E-02 | 179.21 | 83.36 | 193.66 | 111.80 | 231.91 | 544.49 | 321.50 | 568.01 |
| **ENSBTAG00000046218** | KLF11 | -1.55 | 1.17E-04 | 492.60 | 359.56 | 248.99 | 168.85 | 843.62 | 1057.77 | 809.42 | 1003.12 |
| **ENSBTAG00000037710** | | -1.54 | 4.23E-04 | 1997.59 | 2604.28 | 1477.11 | 2001.07 | 3646.64 | 6054.13 | 5509.12 | 8335.38 |
| **ENSBTAG00000008842** | JPH1 | -1.54 | 1.29E-02 | 98.52 | 169.74 | 157.57 | 211.06 | 724.88 | 399.99 | 356.35 | 371.39 |

| | | | | | | | | | | | |
|---|---|---|---|---|---|---|---|---|---|---|---|
| **ENSBTAG00000007841** | WTIP | -1.54 | 2.98E-02 | 166.47 | 200.87 | 168.40 | 78.72 | 203.44 | 358.37 | 617.74 | 608.06 |
| **ENSBTAG00000006209** | EXT1 | -1.54 | 2.86E-03 | 286.22 | 178.77 | 253.80 | 211.06 | 508.95 | 721.36 | 568.95 | 901.17 |
| **ENSBTAG00000003669** | BNC2 | -1.53 | 3.78E-04 | 269.23 | 278.21 | 169.60 | 207.64 | 717.25 | 455.48 | 877.38 | 620.81 |
| **ENSBTAG00000012509** | DYRK1B | -1.51 | 1.76E-03 | 285.37 | 239.04 | 150.36 | 198.51 | 490.89 | 675.12 | 701.38 | 628.09 |
| **ENSBTAG00000014699** | FAM81A | -1.51 | 7.28E-03 | 1157.62 | 930.03 | 740.96 | 624.05 | 1845.54 | 2858.86 | 2628.66 | 2472.30 |
| **ENSBTAG00000002594** | ZNF436 | -1.50 | 3.13E-03 | 150.33 | 246.07 | 165.99 | 116.37 | 328.42 | 458.94 | 372.91 | 767.36 |
| **ENSBTAG00000007898** | CYBRD1 | -1.50 | 2.49E-02 | 78.14 | 102.44 | 147.95 | 149.45 | 342.31 | 375.71 | 302.34 | 332.25 |
| **ENSBTAG00000048544** | | -1.50 | 4.17E-02 | 105.32 | 166.72 | 146.75 | 158.58 | 293.70 | 397.67 | 446.10 | 496.10 |
| **ENSBTAG00000021458** | DLX6 | -1.50 | 7.17E-03 | 78.14 | 209.91 | 259.82 | 167.71 | 426.32 | 497.09 | 572.43 | 525.23 |
| **ENSBTAG00000046918** | NPM2 | -1.49 | 1.39E-02 | 620.85 | 616.67 | 317.55 | 810.01 | 1276.88 | 2177.96 | 1346.13 | 1862.42 |
| **ENSBTAG00000014791** | CTH | -1.49 | 9.71E-03 | 87.48 | 100.44 | 64.95 | 102.68 | 164.56 | 245.08 | 359.84 | 232.12 |
| **ENSBTAG00000033672** | JMJD8 | -1.49 | 2.94E-02 | 86.63 | 44.19 | 52.93 | 17.11 | 136.09 | 112.13 | 149.86 | 169.31 |
| **ENSBTAG00000009950** | PAX3 | -1.49 | 1.94E-03 | 475.62 | 447.94 | 430.62 | 496.27 | 984.56 | 1145.63 | 1334.81 | 1717.69 |
| **ENSBTAG00000020445** | ZNF398 | -1.46 | 1.50E-02 | 332.08 | 378.64 | 230.95 | 148.31 | 683.22 | 705.18 | 961.03 | 661.77 |
| **ENSBTAG00000044208** | DUSP4 | -1.46 | 3.24E-02 | 93.42 | 135.59 | 61.35 | 42.21 | 196.50 | 174.56 | 225.66 | 321.33 |
| **ENSBTAG00000004694** | RPGRIP1 | -1.45 | 1.52E-02 | 391.53 | 436.89 | 134.72 | 305.75 | 626.98 | 1221.92 | 670.89 | 961.25 |
| **ENSBTAG00000017976** | ABL1 | -1.45 | 8.09E-04 | 407.67 | 500.17 | 268.24 | 288.64 | 1009.56 | 941.01 | 1228.51 | 821.07 |
| **ENSBTAG00000006618** | HLF | -1.44 | 6.64E-03 | 558.85 | 438.90 | 241.77 | 376.48 | 747.10 | 1166.43 | 1326.09 | 1154.23 |
| **ENSBTAG00000043957** | TDRP | -1.44 | 3.64E-02 | 92.58 | 58.25 | 40.90 | 85.56 | 152.06 | 213.87 | 203.01 | 185.70 |
| **ENSBTAG00000007071** | RAI14 | -1.44 | 1.35E-02 | 565.64 | 505.19 | 184.04 | 130.06 | 1181.76 | 863.55 | 762.37 | 953.06 |
| **ENSBTAG00000019839** | LTBP1 | -1.43 | 1.94E-02 | 106.16 | 147.64 | 98.63 | 92.41 | 206.91 | 273.98 | 277.07 | 442.39 |
| **ENSBTAG00000002361** | TMCC2 | -1.42 | 4.07E-02 | 95.12 | 107.47 | 63.75 | 65.03 | 237.46 | 121.38 | 228.28 | 303.12 |
| **ENSBTAG00000004593** | TOP2B | -1.42 | 3.78E-04 | 2043.45 | 1599.93 | 864.85 | 949.19 | 2676.66 | 4657.65 | 3088.70 | 4232.77 |
| **ENSBTAG00000008535** | SOCS7 | -1.42 | 4.46E-02 | 170.71 | 200.87 | 61.35 | 168.85 | 305.51 | 387.27 | 528.87 | 390.51 |
| **ENSBTAG00000054366** | | -1.42 | 4.90E-02 | 110.41 | 101.44 | 92.62 | 76.44 | 271.48 | 89.01 | 317.15 | 339.53 |
| **ENSBTAG00000004383** | FNBP1L | -1.42 | 1.61E-03 | 633.59 | 464.01 | 549.70 | 353.67 | 638.09 | 1781.44 | 1443.72 | 1484.66 |
| **ENSBTAG00000032455** | HIST2H2BF | -1.41 | 1.07E-09 | 407.67 | 440.91 | 352.44 | 446.08 | 809.59 | 1151.41 | 1008.07 | 1420.03 |
| **ENSBTAG00000011044** | TACC3 | -1.41 | 1.03E-12 | 8431.15 | 6169.72 | 5570.42 | 5432.77 | 18095.02 | 14713.95 | 19635.23 | 15537.47 |
| **ENSBTAG00000011350** | CCNQ | -1.41 | 7.86E-03 | 1547.45 | 2859.39 | 1302.69 | 3071.19 | 2039.95 | 7767.37 | 6128.60 | 7340.45 |
| **ENSBTAG00000010192** | LYSMD4 | -1.40 | 1.37E-02 | 268.38 | 250.08 | 139.53 | 243.00 | 508.25 | 759.51 | 636.04 | 483.36 |
| **ENSBTAG00000025898** | TBC1D8 | -1.40 | 4.60E-02 | 442.49 | 213.93 | 223.73 | 154.02 | 547.83 | 598.82 | 1049.90 | 535.24 |
| **ENSBTAG00000033803** | FABP7 | -1.39 | 1.75E-02 | 106.16 | 79.34 | 54.13 | 75.30 | 213.16 | 342.18 | 116.75 | 156.57 |
| **ENSBTAG00000017268** | PROCA1 | -1.39 | 2.47E-03 | 693.89 | 578.51 | 338.00 | 335.41 | 1027.61 | 1227.70 | 1586.61 | 1255.27 |

| | | | | | | | | | | | |
|---|---|---|---|---|---|---|---|---|---|---|---|
| **ENSBTAG00000011234** | FOXO3 | -1.39 | 1.95E-02 | 216.58 | 188.82 | 170.81 | 128.92 | 449.23 | 361.84 | 538.45 | 495.19 |
| **ENSBTAG00000011540** | SPG21 | -1.37 | 1.69E-04 | 1376.74 | 1514.56 | 1107.83 | 1235.55 | 1907.33 | 5277.28 | 3396.26 | 2948.38 |
| **ENSBTAG00000008111** | ESYT3 | -1.37 | 1.33E-02 | 129.10 | 85.37 | 127.50 | 87.85 | 418.68 | 257.79 | 244.83 | 187.52 |
| **ENSBTAG00000004862** | TUB | -1.35 | 4.14E-02 | 99.37 | 113.49 | 45.71 | 43.35 | 156.23 | 190.75 | 311.05 | 115.60 |
| **ENSBTAG00000007139** | WSB2 | -1.35 | 3.37E-03 | 633.59 | 776.36 | 507.60 | 501.98 | 1478.24 | 1907.45 | 1258.13 | 1530.17 |
| **ENSBTAG00000038945** | PADI6 | -1.34 | 2.85E-02 | 313.40 | 214.93 | 120.29 | 103.82 | 607.54 | 505.19 | 460.91 | 341.35 |
| **ENSBTAG00000019456** | SPRED2 | -1.34 | 2.31E-02 | 455.23 | 405.76 | 223.73 | 317.16 | 735.30 | 756.04 | 1086.49 | 969.44 |
| **ENSBTAG00000054533** | GNG12 | -1.34 | 1.55E-04 | 502.79 | 553.40 | 387.32 | 487.15 | 906.11 | 1152.56 | 1351.36 | 1465.54 |
| **ENSBTAG00000010947** | PHYHIPL | -1.33 | 1.65E-02 | 837.42 | 1196.18 | 603.83 | 843.09 | 1471.99 | 3028.80 | 2221.77 | 2049.94 |
| **ENSBTAG00000005429** | BCL2L10 | -1.33 | 2.41E-02 | 874.79 | 750.25 | 481.14 | 544.19 | 888.05 | 2001.09 | 2087.59 | 1668.53 |
| **ENSBTAG00000000260** | ZNRF2 | -1.32 | 4.80E-05 | 921.51 | 1077.67 | 766.22 | 891.01 | 1375.47 | 2891.23 | 2855.19 | 2024.45 |
| **ENSBTAG00000003865** | CASR | -1.32 | 4.24E-03 | 232.71 | 218.95 | 221.33 | 280.65 | 765.16 | 373.40 | 692.67 | 546.16 |
| **ENSBTAG00000050550** | FAM110B | -1.32 | 3.07E-02 | 158.82 | 243.05 | 123.89 | 136.90 | 311.06 | 431.20 | 413.86 | 495.19 |
| **ENSBTAG00000010693** | LMO7 | -1.30 | 7.23E-04 | 231.01 | 311.35 | 185.24 | 394.74 | 633.93 | 898.24 | 709.22 | 529.78 |
| **ENSBTAG00000051378** | BEX4 | -1.30 | 9.63E-03 | 84.93 | 176.77 | 122.69 | 132.34 | 256.21 | 506.34 | 280.55 | 230.30 |
| **ENSBTAG00000002417** | RAB30 | -1.30 | 2.16E-02 | 542.71 | 508.20 | 191.25 | 316.02 | 783.21 | 1295.91 | 882.61 | 876.59 |
| **ENSBTAG00000047502** | FKBP5 | -1.29 | 1.28E-03 | 544.41 | 755.27 | 277.86 | 492.85 | 1176.90 | 1676.24 | 1016.79 | 1183.36 |
| **ENSBTAG00000003051** | FER | -1.28 | 1.09E-03 | 1938.99 | 1497.49 | 1910.13 | 1476.27 | 2389.90 | 5514.27 | 4755.46 | 3911.45 |
| **ENSBTAG00000015177** | PRSS23 | -1.28 | 1.73E-03 | 1047.20 | 803.48 | 383.71 | 595.53 | 1540.03 | 1873.93 | 2196.50 | 1258.00 |
| **ENSBTAG00000017063** | EPB41L4B | -1.27 | 1.19E-02 | 326.99 | 444.93 | 261.02 | 325.14 | 763.77 | 1120.19 | 777.18 | 624.45 |
| **ENSBTAG00000021535** | CROT | -1.27 | 2.95E-03 | 695.59 | 746.23 | 590.60 | 742.70 | 868.61 | 2246.17 | 1448.94 | 2148.25 |
| **ENSBTAG00000013996** | SH3BP2 | -1.27 | 3.71E-02 | 103.62 | 124.54 | 116.68 | 78.72 | 392.99 | 291.32 | 170.77 | 165.67 |
| **ENSBTAG00000009749** | USP2 | -1.27 | 9.78E-06 | 1392.03 | 1516.57 | 1157.15 | 897.86 | 3114.78 | 2897.01 | 3127.91 | 2805.46 |
| **ENSBTAG00000047582** | CSNK1E | -1.26 | 1.55E-04 | 710.88 | 771.34 | 470.32 | 497.41 | 1631.68 | 1399.95 | 1589.22 | 1268.01 |
| **ENSBTAG00000004939** | ZNF569 | -1.26 | 5.00E-02 | 138.44 | 136.59 | 102.24 | 92.41 | 229.13 | 299.41 | 187.33 | 410.53 |
| **ENSBTAG00000004976** | CDCA7L | -1.26 | 2.45E-02 | 378.79 | 305.32 | 135.92 | 296.62 | 458.26 | 760.67 | 729.26 | 722.76 |
| **ENSBTAG00000019168** | PHOX2A | -1.25 | 2.31E-03 | 189.40 | 133.58 | 132.31 | 184.82 | 236.77 | 483.22 | 508.83 | 300.39 |
| **ENSBTAG00000024204** | H2BK1 | -1.25 | 1.01E-02 | 94.27 | 147.64 | 86.61 | 99.25 | 236.77 | 198.84 | 319.76 | 262.16 |
| **ENSBTAG00000007256** | DYRK2 | -1.25 | 4.11E-02 | 73.89 | 90.39 | 64.95 | 158.58 | 274.26 | 166.47 | 162.06 | 314.95 |
| **ENSBTAG00000024957** | SNCA | -1.25 | 1.33E-02 | 202.99 | 442.92 | 322.37 | 271.52 | 495.75 | 808.07 | 677.86 | 957.61 |
| **ENSBTAG00000006965** | NLRP8 | -1.25 | 2.08E-02 | 2517.37 | 2691.66 | 1620.25 | 993.69 | 6637.83 | 3525.89 | 4045.37 | 4334.72 |
| **ENSBTAG00000018084** | CASTOR1 | -1.24 | 5.15E-04 | 629.34 | 666.89 | 706.08 | 605.80 | 1112.32 | 1491.28 | 2000.47 | 1572.95 |
| **ENSBTAG00000005252** | NUDT11 | -1.24 | 3.17E-02 | 191.10 | 270.17 | 119.08 | 268.10 | 419.38 | 489.00 | 651.72 | 437.84 |

| | | | | | | | | | | | |
|---|---|---|---|---|---|---|---|---|---|---|---|
| **ENSBTAG00000017747** | PLSCR3 | -1.23 | 3.66E-02 | 106.16 | 108.47 | 66.16 | 50.20 | 224.96 | 283.23 | 128.08 | 145.64 |
| **ENSBTAG00000031793** | NUDT2 | -1.23 | 1.69E-04 | 270.93 | 274.19 | 187.65 | 264.68 | 479.09 | 628.88 | 555.01 | 681.80 |
| **ENSBTAG00000005729** | FBXL4 | -1.23 | 4.46E-02 | 485.81 | 553.40 | 340.41 | 246.43 | 1010.95 | 890.14 | 892.19 | 1017.69 |
| **ENSBTAG00000016836** | PDK1 | -1.23 | 1.10E-03 | 617.45 | 464.01 | 441.45 | 659.42 | 973.46 | 1443.88 | 1296.47 | 1400.00 |
| **ENSBTAG00000044105** | FOXO1 | -1.22 | 8.53E-04 | 798.36 | 1230.33 | 621.88 | 629.75 | 2005.24 | 1871.61 | 1802.68 | 1979.85 |
| **ENSBTAG00000039766** | FBRSL1 | -1.22 | 7.50E-03 | 749.94 | 808.50 | 425.81 | 483.72 | 2422.53 | 876.27 | 1229.38 | 1212.48 |
| **ENSBTAG00000003889** | PER1 | -1.21 | 8.09E-04 | 980.96 | 715.10 | 611.05 | 430.10 | 1752.50 | 1346.78 | 1804.43 | 1443.69 |
| **ENSBTAG00000055062** | SMAD5 | -1.21 | 1.42E-03 | 301.51 | 355.54 | 292.29 | 281.79 | 420.07 | 869.33 | 981.06 | 580.75 |
| **ENSBTAG00000005999** | FSD1 | -1.19 | 6.61E-03 | 152.03 | 290.26 | 153.97 | 96.97 | 312.45 | 398.83 | 408.63 | 468.79 |
| **ENSBTAG00000012382** | KCTD9 | -1.19 | 1.55E-04 | 697.29 | 685.97 | 999.57 | 626.33 | 1131.07 | 2053.11 | 1712.94 | 1968.01 |
| **ENSBTAG00000016658** | ABHD4 | -1.19 | 4.46E-02 | 326.99 | 453.97 | 169.60 | 262.40 | 739.47 | 517.90 | 664.79 | 838.36 |
| **ENSBTAG00000013175** | KIAA0355 | -1.17 | 2.58E-03 | 443.34 | 505.19 | 283.87 | 433.53 | 806.82 | 879.74 | 969.74 | 1102.34 |
| **ENSBTAG00000019155** | FRS2 | -1.16 | 1.53E-02 | 295.56 | 89.39 | 179.23 | 140.33 | 395.77 | 443.92 | 327.60 | 415.08 |
| **ENSBTAG00000055205** | | -1.16 | 1.39E-02 | 106.16 | 132.57 | 90.21 | 86.71 | 174.97 | 189.59 | 301.46 | 266.71 |
| **ENSBTAG00000016805** | SGMS2 | -1.16 | 2.64E-03 | 411.07 | 1046.53 | 761.41 | 659.42 | 965.12 | 1840.40 | 1843.63 | 1770.48 |
| **ENSBTAG00000011905** | RRAS2 | -1.15 | 2.77E-06 | 5879.80 | 4308.66 | 3458.21 | 5050.58 | 9926.19 | 12739.46 | 9263.48 | 9663.47 |
| **ENSBTAG00000001514** | ASB11 | -1.15 | 7.97E-03 | 360.96 | 400.74 | 371.68 | 650.29 | 599.90 | 1394.17 | 1043.80 | 929.39 |
| **ENSBTAG00000009290** | FAM161A | -1.15 | 4.23E-04 | 942.74 | 1189.15 | 1366.44 | 884.17 | 1601.13 | 2656.56 | 2522.36 | 2964.76 |
| **ENSBTAG00000007519** | ADAR | -1.15 | 2.68E-03 | 601.31 | 443.92 | 335.60 | 238.44 | 1115.10 | 667.03 | 880.87 | 923.93 |
| **ENSBTAG00000048876** | FAM117B | -1.14 | 1.59E-02 | 889.23 | 877.80 | 469.11 | 385.61 | 1337.29 | 1397.64 | 1541.30 | 1517.43 |
| **ENSBTAG00000003836** | ADAM19 | -1.14 | 4.04E-03 | 834.88 | 1203.21 | 417.39 | 933.22 | 1243.55 | 1930.57 | 2171.24 | 2130.95 |
| **ENSBTAG00000003994** | IGFBP3 | -1.14 | 2.38E-04 | 427.21 | 475.06 | 334.39 | 466.61 | 1124.13 | 1119.04 | 587.24 | 925.75 |
| **ENSBTAG00000049111** | | -1.14 | 1.89E-02 | 90.03 | 145.63 | 179.23 | 100.40 | 174.97 | 331.78 | 320.63 | 304.94 |
| **ENSBTAG00000019250** | BTK | -1.13 | 2.66E-02 | 610.66 | 347.51 | 279.06 | 426.68 | 1056.08 | 1101.70 | 602.93 | 884.79 |
| **ENSBTAG00000018272** | RERE | -1.13 | 2.38E-04 | 324.44 | 286.24 | 256.21 | 257.83 | 679.75 | 530.62 | 740.59 | 509.75 |
| **ENSBTAG00000005824** | SPOP | -1.13 | 2.31E-03 | 711.73 | 920.99 | 609.85 | 377.62 | 1010.26 | 1908.61 | 1281.66 | 1531.99 |
| **ENSBTAG00000020264** | CCDC181 | -1.12 | 3.90E-03 | 490.05 | 513.22 | 530.46 | 506.54 | 557.55 | 1502.84 | 1086.49 | 1305.33 |
| **ENSBTAG00000014501** | FERMT2 | -1.12 | 3.47E-02 | 2197.18 | 1548.71 | 1024.83 | 1787.72 | 2478.08 | 3678.49 | 4663.98 | 3439.93 |
| **ENSBTAG00000052673** | | -1.12 | 4.11E-02 | 136.74 | 197.86 | 174.41 | 258.97 | 340.22 | 490.16 | 383.36 | 449.68 |
| **ENSBTAG00000027182** | NR3C2 | -1.11 | 1.10E-03 | 814.49 | 871.78 | 648.34 | 604.65 | 1919.83 | 1152.56 | 1541.30 | 1737.71 |
| **ENSBTAG00000011779** | MAL2 | -1.10 | 1.59E-02 | 686.25 | 794.44 | 488.36 | 812.29 | 1185.92 | 2064.67 | 1363.56 | 1364.50 |
| **ENSBTAG00000021505** | USP36 | -1.10 | 1.61E-03 | 1720.71 | 2118.18 | 1281.04 | 1203.61 | 4377.08 | 2335.18 | 4000.06 | 2869.18 |
| **ENSBTAG00000052180** | | -1.09 | 1.12E-02 | 519.78 | 523.27 | 511.21 | 466.61 | 1065.80 | 504.03 | 1286.88 | 1454.62 |

| | | | | | | | | | | | |
|---|---|---|---|---|---|---|---|---|---|---|---|
| **ENSBTAG00000019707** | GATA2 | -1.09 | 2.45E-04 | 607.26 | 359.56 | 436.64 | 520.23 | 838.75 | 969.91 | 1177.10 | 1115.09 |
| **ENSBTAG00000014267** | ZCCHC14 | -1.09 | 3.88E-03 | 517.23 | 763.31 | 697.66 | 533.92 | 1037.33 | 1195.34 | 1836.66 | 1280.76 |
| **ENSBTAG00000023179** | TRIB1 | -1.09 | 2.01E-02 | 1603.51 | 856.71 | 733.74 | 917.25 | 1997.60 | 2038.08 | 2470.96 | 2249.29 |
| **ENSBTAG00000018562** | TMEM159 | -1.08 | 1.93E-02 | 604.71 | 799.46 | 256.21 | 562.44 | 766.54 | 1623.07 | 1097.82 | 1226.14 |
| **ENSBTAG00000020533** | FBXO34 | -1.08 | 2.31E-03 | 697.29 | 640.78 | 525.65 | 452.92 | 967.21 | 1567.58 | 1069.94 | 1301.69 |
| **ENSBTAG00000032021** | RALB | -1.08 | 2.04E-04 | 3645.26 | 4417.13 | 2577.72 | 2395.80 | 5516.48 | 9249.40 | 6122.50 | 6678.68 |
| **ENSBTAG00000033727** | RBPMS | -1.08 | 6.44E-03 | 2790.85 | 1997.65 | 1167.97 | 2109.45 | 3100.89 | 6068.00 | 4102.00 | 3781.28 |
| **ENSBTAG00000002224** | UHRF1 | -1.08 | 7.17E-03 | 2838.41 | 2358.21 | 1641.90 | 1452.31 | 4277.79 | 3968.65 | 5680.76 | 3545.52 |
| **ENSBTAG00000023730** | TUBB3 | -1.07 | 1.75E-02 | 918.96 | 796.45 | 394.54 | 557.88 | 1412.27 | 891.30 | 1976.94 | 1321.72 |
| **ENSBTAG00000046509** | TENT5C | -1.07 | 4.22E-03 | 788.16 | 602.61 | 433.03 | 528.22 | 1184.53 | 1019.62 | 1157.06 | 1577.50 |
| **ENSBTAG00000014619** | WASF3 | -1.07 | 4.11E-02 | 363.51 | 342.48 | 306.73 | 443.79 | 654.76 | 752.58 | 730.14 | 914.83 |
| **ENSBTAG00000007123** | ENSA | -1.06 | 6.91E-07 | 1988.25 | 2424.50 | 1775.41 | 2492.78 | 3675.11 | 5555.88 | 4156.89 | 4769.84 |
| **ENSBTAG00000017245** | COPG2 | -1.06 | 3.60E-02 | 567.34 | 609.64 | 261.02 | 354.81 | 636.70 | 1267.01 | 837.30 | 999.48 |
| **ENSBTAG00000021527** | IGF1R | -1.06 | 4.11E-02 | 144.38 | 235.02 | 232.15 | 169.99 | 498.53 | 393.05 | 362.45 | 371.39 |
| **ENSBTAG00000008105** | RBM38 | -1.05 | 1.75E-02 | 741.45 | 1297.62 | 732.54 | 909.26 | 1521.98 | 1349.09 | 2788.11 | 1943.44 |
| **ENSBTAG00000016094** | SLC36A1 | -1.04 | 1.48E-02 | 181.75 | 177.77 | 182.83 | 90.13 | 467.29 | 312.13 | 297.98 | 224.84 |
| **ENSBTAG00000010363** | LSM11 | -1.03 | 4.71E-02 | 160.52 | 125.54 | 79.39 | 120.93 | 232.60 | 306.35 | 196.04 | 261.25 |
| **ENSBTAG00000010582** | BRD3 | -1.03 | 1.09E-03 | 1087.12 | 1334.78 | 860.04 | 934.36 | 2280.89 | 1910.92 | 2216.54 | 2192.85 |
| **ENSBTAG00000002728** | ARID1B | -1.02 | 2.20E-02 | 335.48 | 273.18 | 145.55 | 185.96 | 514.50 | 425.42 | 429.54 | 546.16 |
| **ENSBTAG00000025462** | GADD45B | -1.02 | 4.37E-04 | 2054.49 | 1744.56 | 1751.36 | 1958.85 | 2372.54 | 3105.10 | 5876.80 | 3855.92 |
| **ENSBTAG00000013278** | MAGIX | -1.02 | 2.81E-02 | 152.88 | 144.63 | 209.30 | 241.86 | 322.17 | 410.39 | 496.63 | 284.92 |
| **ENSBTAG00000022777** | CDC42BPA | -1.01 | 4.11E-02 | 185.15 | 232.00 | 116.68 | 191.66 | 285.37 | 568.77 | 203.01 | 412.35 |
| **ENSBTAG00000008895** | BPGM | -1.01 | 3.07E-03 | 1201.78 | 1253.43 | 790.28 | 1091.80 | 1441.44 | 2626.50 | 2582.48 | 2103.64 |
| **ENSBTAG00000045832** | PHLPP1 | -1.01 | 3.88E-02 | 210.63 | 184.80 | 222.53 | 200.79 | 374.94 | 501.72 | 443.48 | 329.52 |
| **ENSBTAG00000015887** | FOXJ3 | -1.01 | 5.75E-03 | 873.94 | 662.87 | 381.31 | 797.46 | 1185.92 | 1454.29 | 1623.20 | 1205.20 |
| **ENSBTAG00000030483** | KLK7 | -1.01 | 4.69E-02 | 497.70 | 360.56 | 261.02 | 493.99 | 1152.59 | 908.64 | 585.50 | 599.87 |
| **ENSBTAG00000017442** | CDO1 | -1.01 | 3.38E-02 | 911.31 | 1136.92 | 496.78 | 887.59 | 892.91 | 1990.68 | 2003.95 | 2018.08 |
| **ENSBTAG00000021747** | | -1.01 | 5.09E-03 | 636.99 | 434.88 | 564.14 | 756.39 | 793.62 | 1310.94 | 1427.16 | 1276.20 |
| **ENSBTAG00000026243** | PPP1R35 | -1.01 | 3.62E-05 | 2016.27 | 2514.89 | 1705.65 | 1963.42 | 2764.14 | 5183.64 | 4467.07 | 4065.28 |
| **ENSBTAG00000020824** | KRT10 | 1.00 | 1.53E-02 | 778.82 | 938.06 | 1538.45 | 2580.62 | 775.57 | 625.41 | 731.01 | 781.93 |
| **ENSBTAG00000020105** | AQP7 | 1.01 | 5.60E-03 | 1078.63 | 1090.72 | 1465.08 | 1114.62 | 475.62 | 596.51 | 514.93 | 763.72 |
| **ENSBTAG00000030490** | | 1.02 | 4.62E-02 | 88.33 | 217.94 | 133.52 | 146.03 | 52.77 | 76.30 | 78.42 | 81.92 |
| **ENSBTAG00000049134** | | 1.05 | 2.58E-02 | 114.66 | 139.60 | 155.17 | 255.55 | 77.77 | 73.99 | 70.57 | 97.40 |

| | | | | | | | | | | | |
|---|---|---|---|---|---|---|---|---|---|---|---|
| **ENSBTAG00000008280** | HNF4G | 1.06 | 4.69E-02 | 285.37 | 346.50 | 624.28 | 313.74 | 176.36 | 239.30 | 176.87 | 162.94 |
| **ENSBTAG00000025161** | AGPAT2 | 1.07 | 1.71E-02 | 3067.72 | 2876.46 | 3056.46 | 2052.40 | 695.03 | 1183.78 | 1726.88 | 1672.17 |
| **ENSBTAG00000002166** | CRISP2 | 1.12 | 2.63E-02 | 696.44 | 881.82 | 1723.69 | 2030.73 | 663.78 | 778.01 | 710.10 | 309.49 |
| **ENSBTAG00000009252** | KLRA1 | 1.12 | 4.46E-02 | 297.26 | 336.46 | 328.38 | 671.97 | 193.02 | 99.42 | 196.91 | 260.34 |
| **ENSBTAG00000052352** | | 1.13 | 1.78E-05 | 198.74 | 209.91 | 240.57 | 252.13 | 111.09 | 86.70 | 121.11 | 90.12 |
| **ENSBTAG00000042354** | RF00334 | 1.14 | 1.69E-02 | 357.56 | 443.92 | 713.29 | 1092.94 | 218.02 | 254.33 | 331.96 | 377.76 |
| **ENSBTAG00000021292** | ANKFN1 | 1.26 | 2.81E-02 | 779.67 | 459.99 | 704.87 | 940.07 | 178.44 | 485.53 | 227.40 | 312.22 |
| **ENSBTAG00000022564** | AKR1C4 | 1.28 | 2.94E-02 | 484.11 | 520.25 | 573.76 | 204.21 | 150.67 | 270.51 | 103.68 | 212.09 |
| **ENSBTAG00000052382** | H2B | 1.28 | 1.30E-02 | 3927.23 | 3329.42 | 2480.29 | 5044.87 | 612.40 | 1624.22 | 2756.74 | 1099.61 |
| **ENSBTAG00000043641** | RF00413 | 1.41 | 2.95E-02 | 388.99 | 564.44 | 631.50 | 1897.25 | 236.07 | 275.14 | 250.93 | 550.72 |
| **ENSBTAG00000021098** | OR2T29 | 1.43 | 3.16E-03 | 106.16 | 140.61 | 147.95 | 281.79 | 65.27 | 48.55 | 55.76 | 80.10 |
| **ENSBTAG00000043258** | RF00425 | 1.54 | 1.53E-02 | 112.96 | 120.52 | 137.13 | 416.41 | 49.99 | 49.71 | 64.47 | 105.59 |
| **ENSBTAG00000042408** | RF00443 | 1.56 | 1.51E-02 | 149.48 | 188.82 | 280.27 | 636.60 | 118.04 | 83.23 | 84.51 | 140.18 |
| **ENSBTAG00000007268** | F13A1 | 1.59 | 4.95E-03 | 247.15 | 693.00 | 455.88 | 596.67 | 97.90 | 277.45 | 157.70 | 131.99 |
| **ENSBTAG00000043359** | RF00412 | 1.64 | 1.34E-02 | 290.47 | 538.33 | 991.15 | 2120.86 | 229.82 | 284.38 | 255.29 | 491.55 |
| **ENSBTAG00000004347** | ADGRF5 | 1.72 | 2.10E-02 | 180.90 | 376.63 | 114.27 | 305.75 | 109.70 | 31.21 | 128.08 | 27.31 |
| **ENSBTAG00000043222** | RF00586 | 1.75 | 3.95E-06 | 145.23 | 251.09 | 261.02 | 256.69 | 79.85 | 40.46 | 67.96 | 81.01 |
| **ENSBTAG00000000888** | SPTA1 | 1.84 | 4.87E-02 | 92.58 | 65.28 | 186.44 | 308.03 | 61.10 | 71.67 | 26.14 | 23.67 |
| **ENSBTAG00000042280** | RF00561 | 1.87 | 4.46E-02 | 45.86 | 70.30 | 132.31 | 467.75 | 50.69 | 49.71 | 28.75 | 66.45 |
| **ENSBTAG00000044441** | RF00553 | 1.89 | 1.93E-03 | 103.62 | 99.43 | 143.14 | 286.36 | 54.16 | 26.59 | 30.49 | 58.26 |
| **ENSBTAG00000042559** | RF00406 | 1.93 | 4.44E-02 | 45.86 | 83.36 | 92.62 | 402.72 | 35.41 | 20.81 | 30.49 | 76.46 |
| **ENSBTAG00000046066** | | 1.99 | 1.02E-02 | 170.71 | 169.74 | 1533.64 | 263.54 | 106.93 | 126.01 | 160.32 | 144.73 |
| **ENSBTAG00000055165** | RF00045 | 2.04 | 3.73E-04 | 536.77 | 671.91 | 1390.50 | 2299.97 | 174.97 | 316.75 | 183.84 | 514.30 |
| **ENSBTAG00000042702** | RF00394 | 2.09 | 1.34E-02 | 334.63 | 205.89 | 525.65 | 2307.96 | 175.67 | 153.75 | 100.20 | 362.29 |
| **ENSBTAG00000042224** | RF00413 | 2.19 | 2.49E-02 | 42.47 | 79.34 | 167.20 | 521.37 | 24.30 | 41.62 | 47.05 | 64.63 |
| **ENSBTAG00000043128** | RF00265 | 2.20 | 7.17E-03 | 180.90 | 167.73 | 276.66 | 1136.29 | 76.38 | 55.49 | 84.51 | 167.49 |
| **ENSBTAG00000043243** | RF00322 | 2.25 | 1.53E-02 | 76.44 | 109.47 | 277.86 | 879.60 | 67.35 | 30.06 | 34.85 | 149.28 |
| **ENSBTAG00000043394** | RF00421 | 2.32 | 2.81E-02 | 85.78 | 71.31 | 190.05 | 630.89 | 16.66 | 72.83 | 20.04 | 86.48 |
| **ENSBTAG00000021358** | BLNK | 2.35 | 3.64E-02 | 107.86 | 24.10 | 91.42 | 110.66 | 3.47 | 4.62 | 47.05 | 10.01 |
| **ENSBTAG00000043512** | RF00263 | 2.38 | 1.54E-03 | 213.18 | 255.11 | 856.43 | 2035.29 | 106.93 | 135.26 | 148.99 | 255.79 |
| **ENSBTAG00000042475** | RF00092 | 2.39 | 1.65E-02 | 593.67 | 550.38 | 1539.65 | 4865.76 | 230.52 | 428.89 | 186.45 | 596.23 |
| **ENSBTAG00000012759** | FBXO41 | 3.12 | 7.17E-03 | 31.42 | 73.32 | 18.04 | 41.07 | 4.17 | 1.16 | 10.46 | 2.73 |
| **ENSBTAG00000000620** | MCMDC2 | 3.87 | 1.95E-02 | 33.97 | 16.07 | 39.69 | 75.30 | 9.72 | 0.00 | 0.00 | 0.91 |

| | | | | | | | | | | | |
|---|---|---|---|---|---|---|---|---|---|---|---|
| **ENSBTAG00000025952** | OBP | 6.93 | 1.94E-02 | 24.63 | 1.00 | 22.85 | 83.28 | 0.00 | 1.16 | 0.00 | 0.00 |
| **ENSBTAG00000006432** | KCNE4 | 22.00 | 1.17E-04 | 0.00 | 111.48 | 0.00 | 0.00 | 0.00 | 0.00 | 0.00 | 0.00 |
| **ENSBTAG00000007927** | SLAMF1 | 24.53 | 8.94E-06 | 68.79 | 0.00 | 0.00 | 0.00 | 0.00 | 0.00 | 0.00 | 0.00 |
