## Supplementary material 8: DEG_Comparison_ADB_vs_ALL for "Bovine in vitro blastocysts with distinct morphokinetic patterns show transcriptomic differences at genome activation"

| DEGin ADB common to all | | | | | | | | | | | | | | | | | |
|---|---|---|---|---|---|---|---|---|---|---|---|---|---|---|---|---|---|
| GeneID | Gene name | EHB-1 | EHB-2 | EHB-3 | EHB-4 | HB-1 | HB-2 | HB-3 | HB-4 | SSB-1 | SSB-2 | SSB-3 | SSB-4 | ADB-1 | ADB-2 | ADB-3 | ADB-4 |
| ENSBTAG00000016658 | ABHD4 | 217.63 | 173.61 | 313.83 | 216.35 | 251.39 | 444.29 | 112.78 | 243.50 | 326.99 | 453.97 | 169.60 | 262.40 | 739.47 | 517.90 | 664.79 | 838.36 |
| ENSBTAG00000017976 | ABL1 | 417.13 | 234.53 | 332.29 | 222.81 | 330.89 | 661.56 | 466.54 | 226.01 | 407.67 | 500.17 | 268.24 | 288.64 | 1009.56 | 941.01 | 1228.51 | 821.07 |
| ENSBTAG00000003836 | ADAM19 | 670.03 | 607.13 | 659.66 | 578.00 | 699.53 | 1096.93 | 577.70 | 754.21 | 834.88 | 1203.21 | 417.39 | 933.22 | 1243.55 | 1930.57 | 2171.24 | 2130.95 |
| ENSBTAG00000015952 | ADAP1 | 86.65 | 55.84 | 72.61 | 87.18 | 90.42 | 135.39 | 38.95 | 57.44 | 176.66 | 114.50 | 87.81 | 114.09 | 563.80 | 463.57 | 568.08 | 544.34 |
| ENSBTAG00000007519 | ADAR | 418.14 | 278.18 | 342.14 | 302.46 | 329.89 | 458.07 | 452.75 | 317.17 | 601.31 | 443.92 | 335.60 | 238.44 | 1115.10 | 667.03 | 880.87 | 923.93 |
| ENSBTAG00000036087 | ARMC2 | 152.14 | 54.82 | 152.61 | 38.75 | 126.19 | 686.70 | 184.99 | 171.07 | 280.27 | 415.80 | 209.30 | 386.75 | 841.53 | 1205.74 | 1526.49 | 1676.72 |
| ENSBTAG00000001514 | ASB11 | 221.67 | 288.34 | 376.60 | 367.04 | 272.26 | 461.31 | 302.64 | 241.00 | 360.96 | 400.74 | 371.68 | 650.29 | 599.90 | 1394.17 | 1043.80 | 929.39 |
| ENSBTAG00000019675 | ATXN1 | 48.36 | 18.27 | 78.77 | 23.68 | 16.89 | 91.61 | 35.70 | 58.69 | 118.05 | 111.48 | 55.33 | 69.59 | 222.88 | 379.18 | 224.79 | 307.67 |
| ENSBTAG00000048279 | BCAR4 | 141.06 | 89.34 | 198.14 | 82.88 | 69.56 | 445.10 | 165.52 | 189.80 | 221.67 | 307.33 | 262.22 | 359.37 | 601.99 | 1396.48 | 1165.78 | 1174.25 |
| ENSBTAG00000032517 | BCL7A | 4.03 | 11.17 | 3.69 | 0.00 | 3.97 | 13.78 | 12.17 | 18.73 | 22.93 | 7.03 | 0.00 | 6.85 | 68.04 | 72.83 | 78.42 | 87.39 |
| ENSBTAG00000003669 | BNC2 | 282.12 | 143.15 | 172.30 | 116.25 | 112.28 | 194.58 | 211.77 | 124.87 | 269.23 | 278.21 | 169.60 | 207.64 | 717.25 | 455.48 | 877.38 | 620.81 |
| ENSBTAG00000008895 | BPGM | 1087.17 | 793.94 | 1053.48 | 575.85 | 835.66 | 1125.31 | 1033.69 | 752.96 | 1201.78 | 1253.43 | 790.28 | 1091.80 | 1441.44 | 2626.50 | 2582.48 | 2103.64 |
| ENSBTAG00000010582 | BRD3 | 957.19 | 438.60 | 996.87 | 753.45 | 938.01 | 1283.40 | 910.36 | 930.28 | 1087.12 | 1334.78 | 860.04 | 934.36 | 2280.89 | 1910.92 | 2216.54 | 2192.85 |
| ENSBTAG00000034346 | BTG4 | 3545.64 | 1071.11 | 7651.30 | 1505.82 | 1330.50 | 16821.23 | 6067.47 | 4191.88 | 5447.50 | 8049.87 | 5850.69 | 8088.68 | 16773.01 | 58927.49 | 30122.86 | 42014.61 |
| ENSBTAG00000003865 | CASR | 215.62 | 181.73 | 285.52 | 107.64 | 301.08 | 406.18 | 259.64 | 171.07 | 232.71 | 218.95 | 221.33 | 280.65 | 765.16 | 373.40 | 692.67 | 546.16 |
| ENSBTAG00000018084 | CASTOR1 | 697.24 | 342.15 | 575.97 | 278.78 | 492.85 | 787.23 | 597.98 | 638.08 | 629.34 | 666.89 | 706.08 | 605.80 | 1112.32 | 1491.28 | 2000.47 | 1572.95 |
| ENSBTAG00000020346 | CCDC92 | 97.73 | 60.92 | 94.76 | 62.43 | 64.59 | 90.80 | 68.16 | 58.69 | 58.60 | 77.33 | 14.43 | 46.78 | 197.19 | 169.94 | 196.04 | 170.22 |
| ENSBTAG00000031246 | CCNI2 | 113.86 | 42.64 | 68.92 | 47.36 | 80.49 | 265.92 | 63.29 | 101.14 | 200.44 | 95.41 | 96.23 | 110.66 | 377.72 | 649.69 | 639.52 | 529.78 |
| ENSBTAG00000011350 | CCNQ | 1091.20 | 673.12 | 1281.16 | 733.00 | 1014.52 | 2164.68 | 1321.73 | 1503.43 | 1547.45 | 2859.39 | 1302.69 | 3071.19 | 2039.95 | 7767.37 | 6128.60 | 7340.45 |
| ENSBTAG00000033319 | CD200 | 111.84 | 22.34 | 135.38 | 48.44 | 58.63 | 89.18 | 67.34 | 134.86 | 116.36 | 101.44 | 103.45 | 115.23 | 232.60 | 631.19 | 528.87 | 616.26 |
| ENSBTAG00000052516 | CDA | 79.60 | 159.40 | 153.84 | 122.70 | 125.20 | 176.74 | 99.80 | 73.67 | 89.18 | 111.48 | 90.21 | 74.16 | 377.02 | 285.54 | 280.55 | 281.27 |
| ENSBTAG00000004976 | CDCA7L | 258.95 | 327.93 | 182.14 | 433.77 | 145.07 | 428.07 | 285.60 | 324.66 | 378.79 | 305.32 | 135.92 | 296.62 | 458.26 | 760.67 | 729.26 | 722.76 |
| ENSBTAG00000017442 | CDO1 | 682.12 | 1182.79 | 601.82 | 729.77 | 505.77 | 740.21 | 731.86 | 508.22 | 911.31 | 1136.92 | 496.78 | 887.59 | 892.91 | 1990.68 | 2003.95 | 2018.08 |
| ENSBTAG00000021487 | CIART | 990.44 | 339.10 | 452.90 | 613.52 | 435.22 | 1317.45 | 1642.22 | 554.42 | 765.23 | 1410.11 | 925.00 | 1270.92 | 2614.86 | 3714.33 | 3723.86 | 3053.97 |
| ENSBTAG00000017245 | COPG2 | 355.67 | 527.94 | 633.81 | 276.62 | 319.96 | 521.31 | 491.69 | 494.48 | 567.34 | 609.64 | 261.02 | 354.81 | 636.70 | 1267.01 | 837.30 | 999.48 |
| ENSBTAG00000021535 | CROT | 1029.74 | 433.52 | 884.88 | 725.46 | 644.88 | 959.92 | 477.90 | 711.76 | 695.59 | 746.23 | 590.60 | 742.70 | 868.61 | 2246.17 | 1448.94 | 2148.25 |
| ENSBTAG00000047582 | CSNK1E | 557.19 | 258.89 | 364.29 | 259.40 | 321.94 | 468.61 | 401.63 | 388.34 | 710.88 | 771.34 | 470.32 | 497.41 | 1631.68 | 1399.95 | 1589.22 | 1268.01 |
| ENSBTAG00000014791 | CTH | 69.52 | 113.71 | 86.15 | 146.38 | 102.35 | 144.31 | 41.38 | 89.91 | 87.48 | 100.44 | 64.95 | 102.68 | 164.56 | 245.08 | 359.84 | 232.12 |
| ENSBTAG00000012012 | CYB5A | 361.72 | 309.66 | 377.83 | 180.83 | 179.85 | 575.63 | 504.68 | 212.28 | 404.27 | 326.41 | 329.58 | 403.86 | 638.79 | 1369.90 | 1270.33 | 1556.57 |

| | | | | | | | | | | | | | | | | | |
|---|---|---|---|---|---|---|---|---|---|---|---|---|---|---|---|---|---|
| ENSBTAG00000007898 | CYBRD1 | 86.65 | 70.05 | 52.92 | 11.84 | 106.32 | 167.01 | 163.90 | 77.42 | 78.14 | 102.44 | 147.95 | 149.45 | 342.31 | 375.71 | 302.34 | 332.25 |
| ENSBTAG00000008916 | CYFIP2 | 143.07 | 58.89 | 124.30 | 97.95 | 29.81 | 248.90 | 42.19 | 111.13 | 264.99 | 173.75 | 66.16 | 44.49 | 585.32 | 575.70 | 491.40 | 727.31 |
| ENSBTAG00000002733 | DBX1 | 86.65 | 33.50 | 114.46 | 181.90 | 146.07 | 389.16 | 90.87 | 203.54 | 202.14 | 392.70 | 182.83 | 363.93 | 798.48 | 765.29 | 1004.59 | 1007.67 |
| ENSBTAG00000021458 | DLX6 | 291.19 | 114.73 | 204.30 | 176.52 | 147.06 | 393.21 | 214.20 | 179.81 | 78.14 | 209.91 | 259.82 | 167.71 | 426.32 | 497.09 | 572.43 | 525.23 |
| ENSBTAG00000018845 | DND1 | 80.61 | 55.84 | 89.84 | 107.64 | 56.64 | 125.66 | 111.97 | 174.82 | 63.70 | 127.55 | 151.56 | 91.27 | 333.28 | 387.27 | 441.74 | 273.99 |
| ENSBTAG00000001729 | DUSP10 | 336.53 | 67.01 | 219.07 | 117.32 | 63.59 | 347.00 | 411.37 | 509.47 | 291.31 | 605.62 | 256.21 | 252.13 | 1251.88 | 1419.61 | 1614.49 | 1180.63 |
| ENSBTAG00000012509 | DYRK1B | 200.51 | 106.60 | 214.14 | 116.25 | 99.37 | 319.43 | 254.77 | 163.58 | 285.37 | 239.04 | 150.36 | 198.51 | 490.89 | 675.12 | 701.38 | 628.09 |
| ENSBTAG00000037508 | EBF1 | 65.49 | 19.29 | 7.38 | 77.50 | 66.57 | 121.61 | 69.78 | 41.21 | 67.95 | 198.86 | 134.72 | 58.18 | 300.65 | 396.52 | 476.59 | 496.10 |
| ENSBTAG00000003103 | EIF4E1B | 443.33 | 133.00 | 380.29 | 105.48 | 170.91 | 825.33 | 451.12 | 413.32 | 1204.33 | 1484.43 | 716.90 | 990.26 | 2664.85 | 4269.22 | 3690.75 | 5072.96 |
| ENSBTAG00000046348 | ELAVL2 | 483.63 | 296.46 | 454.13 | 470.37 | 158.98 | 878.03 | 404.07 | 345.89 | 517.23 | 713.09 | 344.02 | 490.57 | 879.03 | 1891.27 | 1844.51 | 1468.27 |
| ENSBTAG00000007123 | ENSA | 2397.01 | 2253.90 | 2449.10 | 1792.13 | 1740.88 | 2416.82 | 2018.70 | 1992.92 | 1988.25 | 2424.50 | 1775.41 | 2492.78 | 3675.11 | 5555.88 | 4156.89 | 4769.84 |
| ENSBTAG00000017063 | EPB41L4B | 228.72 | 177.67 | 150.15 | 69.96 | 190.78 | 274.03 | 118.46 | 270.97 | 326.99 | 444.93 | 261.02 | 325.14 | 763.77 | 1120.19 | 777.18 | 624.45 |
| ENSBTAG00000008111 | ESYT3 | 87.66 | 144.17 | 148.92 | 82.88 | 61.61 | 55.13 | 125.76 | 58.69 | 129.10 | 85.37 | 127.50 | 87.85 | 418.68 | 257.79 | 244.83 | 187.52 |
| ENSBTAG00000006209 | EXT1 | 170.28 | 101.53 | 287.99 | 136.70 | 88.43 | 291.06 | 146.86 | 181.06 | 286.22 | 178.77 | 253.80 | 211.06 | 508.95 | 721.36 | 568.95 | 901.17 |
| ENSBTAG00000033015 | FAHD1 | 30.23 | 32.49 | 83.69 | 26.91 | 25.83 | 28.38 | 38.13 | 52.45 | 46.71 | 53.23 | 32.48 | 37.65 | 58.32 | 201.15 | 136.79 | 323.15 |
| ENSBTAG00000050550 | FAM110B | 127.96 | 110.66 | 212.91 | 94.72 | 134.14 | 301.60 | 146.86 | 76.17 | 158.82 | 243.05 | 123.89 | 136.90 | 311.06 | 431.20 | 413.86 | 495.19 |
| ENSBTAG00000048876 | FAM117B | 297.23 | 241.63 | 439.36 | 207.74 | 415.35 | 769.39 | 399.20 | 342.14 | 889.23 | 877.80 | 469.11 | 385.61 | 1337.29 | 1397.64 | 1541.30 | 1517.43 |
| ENSBTAG00000009290 | FAM161A | 983.39 | 661.96 | 1142.10 | 1061.28 | 632.96 | 1216.11 | 999.62 | 1002.70 | 942.74 | 1189.15 | 1366.44 | 884.17 | 1601.13 | 2656.56 | 2522.36 | 2964.76 |
| ENSBTAG00000014699 | FAM81A | 485.65 | 409.15 | 820.88 | 176.52 | 262.32 | 989.91 | 503.05 | 514.46 | 1157.62 | 930.03 | 740.96 | 624.05 | 1845.54 | 2858.86 | 2628.66 | 2472.30 |
| ENSBTAG00000039766 | FBRSL1 | 358.69 | 268.03 | 380.29 | 456.37 | 349.77 | 566.71 | 312.38 | 390.84 | 749.94 | 808.50 | 425.81 | 483.72 | 2422.53 | 876.27 | 1229.38 | 1212.48 |
| ENSBTAG00000005729 | FBXL4 | 153.15 | 60.92 | 203.07 | 121.63 | 120.23 | 406.99 | 216.64 | 379.60 | 485.81 | 553.40 | 340.41 | 246.43 | 1010.95 | 890.14 | 892.19 | 1017.69 |
| ENSBTAG00000020533 | FBXO34 | 560.21 | 437.58 | 440.59 | 342.28 | 348.77 | 774.26 | 475.47 | 586.89 | 697.29 | 640.78 | 525.65 | 452.92 | 967.21 | 1567.58 | 1069.94 | 1301.69 |
| ENSBTAG00000003051 | FER | 1438.81 | 964.51 | 1321.78 | 629.67 | 976.76 | 2124.14 | 1475.89 | 1493.44 | 1938.99 | 1497.49 | 1910.13 | 1476.27 | 2389.90 | 5514.27 | 4755.46 | 3911.45 |
| ENSBTAG00000014501 | FERMT2 | 1631.25 | 1183.80 | 1590.07 | 911.67 | 656.80 | 2720.84 | 1333.90 | 1421.02 | 2197.18 | 1548.71 | 1024.83 | 1787.72 | 2478.08 | 3678.49 | 4663.98 | 3439.93 |
| ENSBTAG00000047502 | FKBP5 | 561.22 | 446.72 | 555.05 | 745.91 | 376.59 | 538.33 | 758.64 | 691.78 | 544.41 | 755.27 | 277.86 | 492.85 | 1176.90 | 1676.24 | 1016.79 | 1183.36 |
| ENSBTAG00000004383 | FNBP1L | 464.49 | 603.07 | 292.91 | 473.60 | 398.45 | 859.38 | 525.77 | 427.05 | 633.59 | 464.01 | 549.70 | 353.67 | 638.09 | 1781.44 | 1443.72 | 1484.66 |
| ENSBTAG00000044105 | FOXO1 | 636.78 | 510.68 | 695.35 | 540.33 | 469.00 | 1090.45 | 478.71 | 688.03 | 798.36 | 1230.33 | 621.88 | 629.75 | 2005.24 | 1871.61 | 1802.68 | 1979.85 |
| ENSBTAG00000011234 | FOXO3 | 136.02 | 150.26 | 107.07 | 104.41 | 37.76 | 231.87 | 103.86 | 142.35 | 216.58 | 188.82 | 170.81 | 128.92 | 449.23 | 361.84 | 538.45 | 495.19 |
| ENSBTAG00000019707 | GATA2 | 321.41 | 283.26 | 436.90 | 368.11 | 303.06 | 511.58 | 388.65 | 349.64 | 607.26 | 359.56 | 436.64 | 520.23 | 838.75 | 969.91 | 1177.10 | 1115.09 |
| ENSBTAG00000018446 | GCA | 325.44 | 135.03 | 211.68 | 89.34 | 86.45 | 582.11 | 311.57 | 259.73 | 347.37 | 317.37 | 340.41 | 436.95 | 892.91 | 1218.46 | 1137.90 | 1058.65 |
| ENSBTAG00000009478 | GDF9 | 237.79 | 142.14 | 204.30 | 184.06 | 112.28 | 949.38 | 375.67 | 211.03 | 338.03 | 751.25 | 336.80 | 754.11 | 1496.98 | 3499.30 | 1931.63 | 2772.69 |

| | | | | | | | | | | | | | | | | | |
|---|---|---|---|---|---|---|---|---|---|---|---|---|---|---|---|---|---|
| ENSBTAG00000010597 | GGCT | 471.54 | 377.68 | 531.66 | 206.66 | 155.01 | 1171.52 | 457.62 | 440.79 | 378.79 | 509.21 | 488.36 | 958.32 | 958.87 | 2762.91 | 2027.48 | 2301.17 |
| ENSBTAG00000011733 | GIPC2 | 51.39 | 15.23 | 32.00 | 19.37 | 22.85 | 83.51 | 43.00 | 104.89 | 225.92 | 215.94 | 187.65 | 181.40 | 510.34 | 1175.68 | 649.98 | 328.61 |
| ENSBTAG00000054533 | GNG12 | 641.82 | 337.07 | 414.75 | 635.05 | 364.67 | 747.50 | 678.31 | 478.25 | 502.79 | 553.40 | 387.32 | 487.15 | 906.11 | 1152.56 | 1351.36 | 1465.54 |
| ENSBTAG00000002908 | GNG4 | 32.24 | 40.61 | 83.69 | 34.44 | 59.62 | 181.61 | 92.50 | 38.71 | 66.25 | 72.31 | 37.29 | 123.21 | 182.61 | 361.84 | 400.79 | 544.34 |
| ENSBTAG00000000830 | GPR137B | 124.94 | 36.55 | 179.68 | 58.12 | 40.74 | 616.97 | 162.28 | 186.06 | 284.52 | 233.01 | 184.04 | 285.21 | 775.57 | 1102.85 | 1150.09 | 1195.19 |
| ENSBTAG00000001812 | H1FOO | 541.06 | 210.16 | 1361.16 | 149.61 | 189.79 | 971.27 | 532.26 | 846.62 | 524.88 | 930.03 | 425.81 | 1116.90 | 2125.35 | 4488.87 | 4000.06 | 2979.33 |
| ENSBTAG00000033089 | HACD1 | 52.39 | 82.24 | 46.77 | 32.29 | 12.92 | 58.37 | 94.12 | 27.47 | 32.27 | 56.24 | 22.85 | 34.23 | 110.40 | 206.93 | 105.43 | 224.84 |
| ENSBTAG00000032455 | HIST2H2BF | 308.32 | 255.85 | 235.06 | 326.14 | 402.43 | 534.28 | 483.58 | 443.29 | 407.67 | 440.91 | 352.44 | 446.08 | 809.59 | 1151.41 | 1008.07 | 1420.03 |
| ENSBTAG00000006618 | HLF | 216.63 | 78.18 | 231.37 | 206.66 | 169.91 | 505.09 | 288.85 | 409.57 | 558.85 | 438.90 | 241.77 | 376.48 | 747.10 | 1166.43 | 1326.09 | 1154.23 |
| ENSBTAG00000004710 | HNF1B | 64.48 | 41.63 | 45.54 | 11.84 | 21.86 | 135.39 | 85.19 | 32.47 | 69.64 | 78.34 | 42.10 | 50.20 | 148.59 | 317.91 | 152.47 | 331.34 |
| ENSBTAG00000030425 | ID3 | 1133.52 | 619.31 | 1676.22 | 339.05 | 348.77 | 2855.43 | 1163.51 | 1336.11 | 849.31 | 2762.97 | 956.27 | 2346.75 | 3149.50 | 11844.69 | 7628.95 | 9749.03 |
| ENSBTAG00000021527 | IGF1R | 204.54 | 146.20 | 171.07 | 68.89 | 147.06 | 181.61 | 249.90 | 129.86 | 144.38 | 235.02 | 232.15 | 169.99 | 498.53 | 393.05 | 362.45 | 371.39 |
| ENSBTAG00000003994 | IGFBP3 | 317.38 | 396.97 | 508.28 | 362.73 | 332.87 | 406.99 | 477.90 | 383.35 | 427.21 | 475.06 | 334.39 | 466.61 | 1124.13 | 1119.04 | 587.24 | 925.75 |
| ENSBTAG00000022580 | INKA2 | 57.43 | 58.89 | 100.92 | 29.06 | 23.85 | 119.18 | 80.33 | 56.19 | 100.22 | 93.40 | 110.66 | 55.90 | 358.28 | 476.28 | 250.06 | 309.49 |
| ENSBTAG00000045526 | INSM1 | 60.45 | 18.27 | 34.46 | 33.37 | 27.82 | 18.65 | 12.98 | 24.97 | 18.68 | 73.32 | 49.32 | 34.23 | 104.84 | 120.23 | 199.52 | 173.86 |
| ENSBTAG00000002929 | IRF4 | 12.09 | 12.18 | 16.00 | 11.84 | 21.86 | 13.78 | 42.19 | 31.22 | 32.27 | 30.13 | 16.84 | 30.80 | 123.59 | 65.89 | 152.47 | 85.57 |
| ENSBTAG00000017824 | IRF8 | 349.63 | 334.02 | 748.27 | 377.80 | 220.59 | 539.14 | 455.99 | 530.70 | 680.30 | 734.18 | 223.73 | 433.53 | 1421.99 | 1935.19 | 1938.60 | 1494.67 |
| ENSBTAG00000019024 | JAZF1 | 111.84 | 56.86 | 52.92 | 13.99 | 39.75 | 175.93 | 97.37 | 73.67 | 74.74 | 112.49 | 60.14 | 68.45 | 337.45 | 375.71 | 385.98 | 304.03 |
| ENSBTAG00000008842 | JPH1 | 325.44 | 110.66 | 83.69 | 130.24 | 164.95 | 269.17 | 229.62 | 126.12 | 98.52 | 169.74 | 157.57 | 211.06 | 724.88 | 399.99 | 356.35 | 371.39 |
| ENSBTAG00000012382 | KCTD9 | 449.38 | 679.22 | 559.97 | 597.38 | 478.94 | 848.84 | 805.70 | 639.33 | 697.29 | 685.97 | 999.57 | 626.33 | 1131.07 | 2053.11 | 1712.94 | 1968.01 |
| ENSBTAG00000046218 | KLF11 | 577.34 | 456.87 | 302.75 | 490.82 | 389.51 | 594.27 | 324.55 | 380.85 | 492.60 | 359.56 | 248.99 | 168.85 | 843.62 | 1057.77 | 809.42 | 1003.12 |
| ENSBTAG00000006844 | LEF1 | 178.34 | 67.01 | 338.44 | 137.77 | 129.17 | 792.90 | 193.92 | 385.85 | 472.22 | 514.23 | 344.02 | 325.14 | 1042.19 | 910.95 | 2053.61 | 1816.00 |
| ENSBTAG00000010192 | LYSMD4 | 116.88 | 105.59 | 241.22 | 149.61 | 158.98 | 470.23 | 135.50 | 260.98 | 268.38 | 250.08 | 139.53 | 243.00 | 508.25 | 759.51 | 636.04 | 483.36 |
| ENSBTAG00000017354 | MPP7 | 294.21 | 96.45 | 424.59 | 137.77 | 245.43 | 1008.56 | 312.38 | 389.59 | 802.60 | 673.92 | 345.22 | 256.69 | 1105.38 | 2036.93 | 1690.29 | 1448.25 |
| ENSBTAG00000010875 | MSX1 | 785.90 | 466.01 | 1309.47 | 588.77 | 576.32 | 1566.35 | 902.25 | 1001.46 | 1643.42 | 1532.64 | 935.82 | 1302.86 | 4063.24 | 5272.65 | 5053.44 | 4325.62 |
| ENSBTAG00000026972 | MYF5 | 324.44 | 256.86 | 358.14 | 99.02 | 395.47 | 1306.91 | 477.90 | 349.64 | 536.77 | 1062.60 | 761.41 | 788.33 | 1991.35 | 4783.65 | 4497.56 | 3931.47 |
| ENSBTAG00000004115 | MYLIP | 590.44 | 519.82 | 849.19 | 480.05 | 404.42 | 787.23 | 351.33 | 555.67 | 931.70 | 1056.58 | 588.20 | 893.29 | 1658.07 | 2705.11 | 3499.07 | 2381.28 |
| ENSBTAG00000002289 | NLRP14 | 412.10 | 196.96 | 198.14 | 124.86 | 139.11 | 850.47 | 391.08 | 278.46 | 642.08 | 415.80 | 445.06 | 286.36 | 1383.81 | 2100.51 | 1307.80 | 1131.47 |
| ENSBTAG00000006965 | NLRP8 | 1438.81 | 579.72 | 1745.14 | 891.22 | 907.20 | 2152.51 | 1781.78 | 1273.67 | 2517.37 | 2691.66 | 1620.25 | 993.69 | 6637.83 | 3525.89 | 4045.37 | 4334.72 |
| ENSBTAG00000046918 | NPM2 | 397.99 | 255.85 | 318.75 | 116.25 | 204.69 | 928.30 | 579.32 | 399.58 | 620.85 | 616.67 | 317.55 | 810.01 | 1276.88 | 2177.96 | 1346.13 | 1862.42 |
| ENSBTAG00000027182 | NR3C2 | 479.60 | 469.05 | 414.75 | 386.41 | 337.84 | 839.93 | 668.57 | 518.21 | 814.49 | 871.78 | 648.34 | 604.65 | 1919.83 | 1152.56 | 1541.30 | 1737.71 |

| | | | | | | | | | | | | | | | | | |
|---|---|---|---|---|---|---|---|---|---|---|---|---|---|---|---|---|---|
| ENSBTAG00000038945 | PADI6 | 112.85 | 74.11 | 153.84 | 96.87 | 86.45 | 264.30 | 153.35 | 179.81 | 313.40 | 214.93 | 120.29 | 103.82 | 607.54 | 505.19 | 460.91 | 341.35 |
| ENSBTAG00000012010 | PANX1 | 128.97 | 108.63 | 61.54 | 79.65 | 68.56 | 63.24 | 21.91 | 149.84 | 135.04 | 39.17 | 104.65 | 88.99 | 356.89 | 298.26 | 250.93 | 209.36 |
| ENSBTAG00000009950 | PAX3 | 322.42 | 424.38 | 327.37 | 190.51 | 197.74 | 749.93 | 285.60 | 289.70 | 475.62 | 447.94 | 430.62 | 496.27 | 984.56 | 1145.63 | 1334.81 | 1717.69 |
| ENSBTAG00000004561 | PAX6 | 60.45 | 36.55 | 38.15 | 52.74 | 58.63 | 58.37 | 61.66 | 51.20 | 36.52 | 43.19 | 38.49 | 53.62 | 113.18 | 76.30 | 155.09 | 172.04 |
| ENSBTAG00000016836 | PDK1 | 233.76 | 282.25 | 568.59 | 283.08 | 528.62 | 657.51 | 471.41 | 370.86 | 617.45 | 464.01 | 441.45 | 659.42 | 973.46 | 1443.88 | 1296.47 | 1400.00 |
| ENSBTAG00000003889 | PER1 | 382.88 | 231.48 | 374.13 | 354.12 | 639.91 | 686.70 | 544.43 | 342.14 | 980.96 | 715.10 | 611.05 | 430.10 | 1752.50 | 1346.78 | 1804.43 | 1443.69 |
| ENSBTAG00000019168 | PHOX2A | 164.23 | 169.55 | 204.30 | 121.63 | 155.01 | 218.09 | 202.03 | 149.84 | 189.40 | 133.58 | 132.31 | 184.82 | 236.77 | 483.22 | 508.83 | 300.39 |
| ENSBTAG00000010947 | PHYHIPL | 923.94 | 190.87 | 552.59 | 294.92 | 821.75 | 1036.94 | 589.87 | 847.87 | 837.42 | 1196.18 | 603.83 | 843.09 | 1471.99 | 3028.80 | 2221.77 | 2049.94 |
| ENSBTAG00000006095 | PRDM13 | 94.71 | 30.46 | 39.38 | 54.89 | 62.60 | 140.26 | 47.87 | 89.91 | 50.11 | 112.49 | 54.13 | 44.49 | 180.53 | 387.27 | 372.91 | 262.16 |
| ENSBTAG00000017268 | PROCA1 | 422.17 | 320.83 | 500.90 | 196.97 | 266.30 | 581.30 | 349.70 | 275.96 | 693.89 | 578.51 | 338.00 | 335.41 | 1027.61 | 1227.70 | 1586.61 | 1255.27 |
| ENSBTAG00000015177 | PRSS23 | 422.17 | 793.94 | 804.88 | 597.38 | 515.70 | 587.79 | 473.03 | 349.64 | 1047.20 | 803.48 | 383.71 | 595.53 | 1540.03 | 1873.93 | 2196.50 | 1258.00 |
| ENSBTAG00000002417 | RAB30 | 249.88 | 414.23 | 236.30 | 289.54 | 212.64 | 344.56 | 305.08 | 97.40 | 542.71 | 508.20 | 191.25 | 316.02 | 783.21 | 1295.91 | 882.61 | 876.59 |
| ENSBTAG00000007306 | RAB3C | 63.48 | 24.37 | 48.00 | 52.74 | 33.78 | 95.67 | 114.40 | 141.10 | 179.21 | 83.36 | 193.66 | 111.80 | 231.91 | 544.49 | 321.50 | 568.01 |
| ENSBTAG00000007071 | RAI14 | 418.14 | 328.95 | 339.67 | 226.03 | 221.58 | 449.96 | 234.49 | 295.94 | 565.64 | 505.19 | 184.04 | 130.06 | 1181.76 | 863.55 | 762.37 | 953.06 |
| ENSBTAG00000032021 | RALB | 2200.53 | 1914.80 | 2590.64 | 1734.01 | 2806.07 | 3576.99 | 2593.16 | 2477.41 | 3645.26 | 4417.13 | 2577.72 | 2395.80 | 5516.48 | 9249.40 | 6122.50 | 6678.68 |
| ENSBTAG00000008105 | RBM38 | 829.23 | 413.21 | 508.28 | 483.28 | 584.27 | 1068.56 | 583.38 | 691.78 | 741.45 | 1297.62 | 732.54 | 909.26 | 1521.98 | 1349.09 | 2788.11 | 1943.44 |
| ENSBTAG00000033727 | RBPMS | 1504.30 | 2415.33 | 2284.19 | 1766.30 | 1526.25 | 2733.82 | 2297.82 | 1577.10 | 2790.85 | 1997.65 | 1167.97 | 2109.45 | 3100.89 | 6068.00 | 4102.00 | 3781.28 |
| ENSBTAG00000018272 | RERE | 286.15 | 208.13 | 173.53 | 163.61 | 234.50 | 366.45 | 264.51 | 280.96 | 324.44 | 286.24 | 256.21 | 257.83 | 679.75 | 530.62 | 740.59 | 509.75 |
| ENSBTAG00000034366 | RGS2 | 1367.27 | 395.96 | 2313.73 | 287.39 | 559.43 | 4469.61 | 1626.00 | 2037.87 | 1915.20 | 3482.08 | 1015.21 | 2380.97 | 7097.48 | 17061.85 | 14700.28 | 11699.75 |
| ENSBTAG00000003827 | RIMS2 | 125.95 | 39.60 | 183.38 | 207.74 | 48.69 | 534.28 | 165.52 | 320.92 | 169.01 | 306.33 | 289.89 | 162.00 | 731.83 | 1396.48 | 1247.68 | 1104.16 |
| ENSBTAG00000004694 | RPGRIP1 | 468.52 | 195.95 | 297.83 | 315.37 | 253.38 | 633.19 | 207.71 | 279.71 | 391.53 | 436.89 | 134.72 | 305.75 | 626.98 | 1221.92 | 670.89 | 961.25 |
| ENSBTAG00000011905 | RRAS2 | 4417.18 | 4517.95 | 5420.03 | 4242.99 | 3626.83 | 6055.42 | 3673.10 | 4582.72 | 5879.80 | 4308.66 | 3458.21 | 5050.58 | 9926.19 | 12739.46 | 9263.48 | 9663.47 |
| ENSBTAG00000001322 | SAXO1 | 793.96 | 375.65 | 776.58 | 328.29 | 294.12 | 2109.55 | 551.74 | 949.01 | 2046.00 | 1916.30 | 971.91 | 2068.38 | 4121.56 | 8569.65 | 5602.35 | 7514.31 |
| ENSBTAG00000018394 | SDR42E1 | 172.29 | 198.99 | 162.45 | 150.69 | 183.83 | 184.85 | 103.04 | 255.98 | 195.34 | 211.92 | 161.18 | 99.25 | 422.85 | 751.42 | 488.79 | 692.72 |
| ENSBTAG00000008091 | SELENBP1 | 99.75 | 64.98 | 65.23 | 27.99 | 36.77 | 120.80 | 107.10 | 99.90 | 218.27 | 136.59 | 72.17 | 139.18 | 412.43 | 582.64 | 633.42 | 656.31 |
| ENSBTAG00000018133 | SEMA3A | 3.02 | 7.11 | 3.69 | 5.38 | 1.99 | 17.84 | 5.68 | 9.99 | 12.74 | 17.07 | 3.61 | 6.85 | 42.35 | 102.89 | 54.89 | 149.28 |
| ENSBTAG00000005230 | SHOX2 | 21.16 | 38.58 | 48.00 | 0.00 | 42.73 | 85.13 | 34.89 | 59.94 | 77.29 | 67.29 | 33.68 | 33.08 | 229.13 | 208.09 | 277.07 | 284.92 |
| ENSBTAG00000048053 | SLBP2 | 577.34 | 149.24 | 694.12 | 266.94 | 316.97 | 1730.93 | 613.40 | 468.26 | 792.41 | 1068.63 | 501.59 | 734.71 | 2447.53 | 4520.08 | 4087.19 | 3996.10 |
| ENSBTAG00000007403 | SLC7A3 | 156.17 | 19.29 | 150.15 | 49.51 | 64.59 | 304.84 | 97.37 | 132.36 | 196.19 | 248.07 | 87.81 | 224.75 | 583.24 | 1164.12 | 681.34 | 756.44 |
| ENSBTAG00000055062 | SMAD5 | 466.50 | 237.57 | 319.98 | 292.77 | 347.78 | 380.24 | 277.49 | 374.61 | 301.51 | 355.54 | 292.29 | 281.79 | 420.07 | 869.33 | 981.06 | 580.75 |
| ENSBTAG00000024957 | SNCA | 282.12 | 178.69 | 322.44 | 204.51 | 167.93 | 401.32 | 435.71 | 232.26 | 202.99 | 442.92 | 322.37 | 271.52 | 495.75 | 808.07 | 677.86 | 957.61 |

| | | | | | | | | | | | | | | | | | |
|---|---|---|---|---|---|---|---|---|---|---|---|---|---|---|---|---|---|
| ENSBTAG00000001080 | SPAG17 | 199.50 | 186.81 | 339.67 | 111.94 | 111.29 | 580.49 | 283.98 | 395.84 | 293.01 | 299.30 | 215.31 | 412.99 | 769.32 | 1595.32 | 792.87 | 1275.29 |
| ENSBTAG00000011540 | SPG21 | 1458.96 | 944.20 | 1158.09 | 795.43 | 983.71 | 1706.61 | 1166.76 | 987.72 | 1376.74 | 1514.56 | 1107.83 | 1235.55 | 1907.33 | 5277.28 | 3396.26 | 2948.38 |
| ENSBTAG00000005824 | SPOP | 677.09 | 434.54 | 751.96 | 597.38 | 478.94 | 618.59 | 623.95 | 611.86 | 711.73 | 920.99 | 609.85 | 377.62 | 1010.26 | 1908.61 | 1281.66 | 1531.99 |
| ENSBTAG00000019456 | SPRED2 | 237.79 | 164.47 | 379.06 | 85.03 | 177.86 | 452.39 | 301.83 | 227.26 | 455.23 | 405.76 | 223.73 | 317.16 | 735.30 | 756.04 | 1086.49 | 969.44 |
| ENSBTAG00000011044 | TACC3 | 5569.84 | 4202.20 | 5416.34 | 3752.17 | 4706.93 | 5835.71 | 5586.32 | 5138.39 | 8431.15 | 6169.72 | 5570.42 | 5432.77 | 18095.02 | 14713.95 | 19635.23 | 15537.47 |
| ENSBTAG00000025898 | TBC1D8 | 211.59 | 106.60 | 61.54 | 93.64 | 84.46 | 291.87 | 258.02 | 181.06 | 442.49 | 213.93 | 223.73 | 154.02 | 547.83 | 598.82 | 1049.90 | 535.24 |
| ENSBTAG00000019580 | TCL1A | 25.19 | 12.18 | 35.69 | 11.84 | 9.94 | 89.99 | 23.53 | 21.23 | 39.92 | 70.30 | 20.45 | 36.51 | 121.51 | 210.40 | 158.57 | 395.97 |
| ENSBTAG00000043957 | TDRP | 66.50 | 70.05 | 98.46 | 51.66 | 28.82 | 128.10 | 51.12 | 88.66 | 92.58 | 58.25 | 40.90 | 85.56 | 152.06 | 213.87 | 203.01 | 185.70 |
| ENSBTAG00000007439 | TENM4 | 72.54 | 44.67 | 43.07 | 105.48 | 44.71 | 40.54 | 33.27 | 56.19 | 77.29 | 68.30 | 38.49 | 53.62 | 193.72 | 153.75 | 212.59 | 173.86 |
| ENSBTAG00000046509 | TENT5C | 476.58 | 277.17 | 438.13 | 355.20 | 179.85 | 378.62 | 344.83 | 274.71 | 788.16 | 602.61 | 433.03 | 528.22 | 1184.53 | 1019.62 | 1157.06 | 1577.50 |
| ENSBTAG00000002361 | TMCC2 | 78.59 | 64.98 | 68.92 | 45.21 | 48.69 | 111.88 | 68.97 | 23.73 | 95.12 | 107.47 | 63.75 | 65.03 | 237.46 | 121.38 | 228.28 | 303.12 |
| ENSBTAG00000039720 | TMEM225B | 259.95 | 190.87 | 262.14 | 298.15 | 248.41 | 764.53 | 366.74 | 297.19 | 574.14 | 676.93 | 239.37 | 585.26 | 1037.33 | 2206.86 | 1697.26 | 1561.12 |
| ENSBTAG00000048560 | TMEM233 | 9.07 | 5.08 | 19.69 | 16.15 | 9.94 | 122.42 | 29.21 | 14.98 | 28.03 | 48.21 | 32.48 | 25.10 | 141.64 | 278.60 | 156.83 | 225.75 |
| ENSBTAG00000004593 | TOP2B | 1205.05 | 691.40 | 1175.32 | 961.18 | 655.81 | 1455.28 | 947.69 | 907.80 | 2043.45 | 1599.93 | 864.85 | 949.19 | 2676.66 | 4657.65 | 3088.70 | 4232.77 |
| ENSBTAG00000023179 | TRIB1 | 947.12 | 1058.93 | 958.72 | 494.05 | 517.69 | 1441.50 | 773.24 | 759.21 | 1603.51 | 856.71 | 733.74 | 917.25 | 1997.60 | 2038.08 | 2470.96 | 2249.29 |
| ENSBTAG00000008078 | TRIM59 | 211.59 | 141.12 | 161.22 | 212.04 | 268.29 | 457.26 | 64.10 | 128.62 | 202.99 | 414.80 | 226.14 | 263.54 | 384.66 | 1780.29 | 750.17 | 1260.73 |
| ENSBTAG00000004862 | TUB | 72.54 | 62.95 | 48.00 | 54.89 | 41.73 | 52.70 | 61.66 | 39.96 | 99.37 | 113.49 | 45.71 | 43.35 | 156.23 | 190.75 | 311.05 | 115.60 |
| ENSBTAG00000023730 | TUBB3 | 432.25 | 296.46 | 503.36 | 529.57 | 425.28 | 625.89 | 395.14 | 359.62 | 918.96 | 796.45 | 394.54 | 557.88 | 1412.27 | 891.30 | 1976.94 | 1321.72 |
| ENSBTAG00000020796 | UBE2D1 | 174.31 | 253.82 | 183.38 | 166.83 | 31.80 | 418.34 | 124.14 | 139.85 | 168.16 | 170.74 | 105.85 | 188.24 | 209.69 | 879.74 | 566.33 | 730.04 |
| ENSBTAG00000002224 | UHRF1 | 1634.28 | 907.65 | 1358.70 | 837.40 | 968.81 | 1937.67 | 1081.56 | 1437.25 | 2838.41 | 2358.21 | 1641.90 | 1452.31 | 4277.79 | 3968.65 | 5680.76 | 3545.52 |
| ENSBTAG00000030721 | UNCX | 346.60 | 106.60 | 364.29 | 268.01 | 236.49 | 558.60 | 288.04 | 298.44 | 318.49 | 473.05 | 282.67 | 365.07 | 1047.75 | 1055.46 | 1096.07 | 1350.85 |
| ENSBTAG00000006188 | USH2A | 56.42 | 15.23 | 118.15 | 62.43 | 60.61 | 187.28 | 98.18 | 143.60 | 95.12 | 168.73 | 105.85 | 93.55 | 379.80 | 664.72 | 598.57 | 821.98 |
| ENSBTAG00000009749 | USP2 | 648.87 | 687.34 | 887.34 | 740.53 | 632.96 | 1307.72 | 660.46 | 912.80 | 1392.03 | 1516.57 | 1157.15 | 897.86 | 3114.78 | 2897.01 | 3127.91 | 2805.46 |
| ENSBTAG00000021505 | USP36 | 1141.58 | 699.52 | 999.33 | 1177.53 | 1196.36 | 1853.35 | 1159.46 | 1111.34 | 1720.71 | 2118.18 | 1281.04 | 1203.61 | 4377.08 | 2335.18 | 4000.06 | 2869.18 |
| ENSBTAG00000018600 | VGLL4 | 221.67 | 211.18 | 280.60 | 223.88 | 136.13 | 414.29 | 241.79 | 112.38 | 375.40 | 472.04 | 310.34 | 302.33 | 1000.53 | 1308.63 | 1413.22 | 1008.58 |
| ENSBTAG00000004791 | VPS26C | 282.12 | 272.09 | 104.61 | 115.17 | 112.28 | 318.62 | 79.51 | 234.76 | 191.95 | 187.81 | 135.92 | 227.03 | 492.98 | 567.61 | 816.39 | 588.04 |
| ENSBTAG00000014619 | WASF3 | 215.62 | 103.56 | 375.37 | 350.89 | 238.48 | 435.37 | 315.63 | 242.25 | 363.51 | 342.48 | 306.73 | 443.79 | 654.76 | 752.58 | 730.14 | 914.83 |
| ENSBTAG00000007139 | WSB2 | 602.53 | 419.31 | 696.58 | 428.39 | 280.21 | 791.28 | 584.19 | 228.51 | 633.59 | 776.36 | 507.60 | 501.98 | 1478.24 | 1907.45 | 1258.13 | 1530.17 |
| ENSBTAG00000007841 | WTIP | 178.34 | 135.03 | 103.38 | 41.98 | 127.19 | 227.82 | 104.67 | 108.64 | 166.47 | 200.87 | 168.40 | 78.72 | 203.44 | 358.37 | 617.74 | 608.06 |
| ENSBTAG00000004886 | ZAR1 | 134.01 | 125.89 | 171.07 | 77.50 | 99.37 | 489.69 | 88.44 | 186.06 | 397.48 | 234.01 | 115.47 | 188.24 | 540.89 | 833.50 | 843.40 | 1099.61 |
| ENSBTAG00000026384 | ZAR1L | 1244.35 | 510.68 | 1970.36 | 360.58 | 270.27 | 2194.67 | 1098.60 | 1595.83 | 1925.40 | 1931.37 | 1853.60 | 3128.23 | 4869.36 | 11211.18 | 10930.24 | 10103.13 |

| | | | | | | | | | | | | | | | | | |
|---|---|---|---|---|---|---|---|---|---|---|---|---|---|---|---|---|---|
| ENSBTAG00000014267 | ZCCHC14 | 467.51 | 394.94 | 520.59 | 396.10 | 276.24 | 708.59 | 438.14 | 402.08 | 517.23 | 763.31 | 697.66 | 533.92 | 1037.33 | 1195.34 | 1836.66 | 1280.76 |
| ENSBTAG00000012629 | ZNF362 | 5.04 | 7.11 | 8.61 | 6.46 | 4.97 | 51.89 | 54.36 | 13.74 | 41.62 | 48.21 | 2.41 | 7.99 | 132.62 | 105.20 | 253.54 | 233.03 |
| ENSBTAG00000020445 | ZNF398 | 178.34 | 70.05 | 239.99 | 121.63 | 162.96 | 321.86 | 93.31 | 223.52 | 332.08 | 378.64 | 230.95 | 148.31 | 683.22 | 705.18 | 961.03 | 661.77 |
| ENSBTAG00000002594 | ZNF436 | 87.66 | 139.09 | 66.46 | 75.34 | 114.27 | 242.41 | 131.44 | 166.08 | 150.33 | 246.07 | 165.99 | 116.37 | 328.42 | 458.94 | 372.91 | 767.36 |
| ENSBTAG00000000260 | ZNRF2 | 698.25 | 650.79 | 772.88 | 838.48 | 614.08 | 1185.30 | 713.20 | 1076.38 | 921.51 | 1077.67 | 766.22 | 891.01 | 1375.47 | 2891.23 | 2855.19 | 2024.45 |
| ENSBTAG00000000623 | | 2218.67 | 1317.82 | 3151.84 | 923.51 | 1424.90 | 5010.37 | 2430.07 | 2541.10 | 6756.30 | 6351.51 | 4809.02 | 6629.53 | 11675.22 | 22879.00 | 22287.42 | 20046.96 |
| ENSBTAG00000005429 | | 341.57 | 305.60 | 748.27 | 158.22 | 311.01 | 646.16 | 413.80 | 467.01 | 874.79 | 750.25 | 481.14 | 544.19 | 888.05 | 2001.09 | 2087.59 | 1668.53 |
| ENSBTAG00000006961 | | 33.25 | 40.61 | 27.08 | 19.37 | 41.73 | 137.02 | 21.10 | 47.45 | 48.41 | 51.22 | 68.56 | 31.94 | 257.60 | 184.96 | 185.58 | 75.55 |
| ENSBTAG00000012389 | | 609.58 | 268.03 | 433.21 | 200.20 | 178.86 | 510.77 | 423.54 | 206.04 | 366.90 | 497.15 | 173.21 | 318.30 | 1852.48 | 543.33 | 1132.67 | 1330.82 |
| ENSBTAG00000013274 | | 1850.90 | 1027.45 | 2814.62 | 741.61 | 1079.11 | 2599.23 | 2207.75 | 1559.62 | 2210.77 | 4062.60 | 1530.03 | 2992.47 | 6068.47 | 14260.79 | 10654.92 | 11768.93 |
| ENSBTAG00000021747 | | 409.07 | 249.76 | 409.83 | 487.59 | 573.34 | 454.83 | 432.46 | 630.59 | 636.99 | 434.88 | 564.14 | 756.39 | 793.62 | 1310.94 | 1427.16 | 1276.20 |
| ENSBTAG00000023666 | | 446.35 | 248.74 | 308.91 | 145.31 | 110.30 | 407.80 | 172.82 | 234.76 | 258.19 | 287.24 | 269.44 | 179.11 | 665.17 | 950.26 | 653.46 | 1043.17 |
| ENSBTAG00000024204 | | 94.71 | 51.78 | 108.30 | 69.96 | 67.57 | 113.50 | 84.38 | 141.10 | 94.27 | 147.64 | 86.61 | 99.25 | 236.77 | 198.84 | 319.76 | 262.16 |
| ENSBTAG00000037710 | | 1432.76 | 651.80 | 931.64 | 539.25 | 801.88 | 2029.28 | 933.08 | 1190.01 | 1997.59 | 2604.28 | 1477.11 | 2001.07 | 3646.64 | 6054.13 | 5509.12 | 8335.38 |
| ENSBTAG00000040202 | | 13.10 | 2.03 | 23.38 | 5.38 | 47.70 | 132.96 | 47.06 | 43.70 | 55.21 | 14.06 | 30.07 | 5.70 | 205.52 | 223.11 | 249.19 | 576.20 |
| ENSBTAG00000044125 | | 34.26 | 69.04 | 56.61 | 26.91 | 42.73 | 379.43 | 162.28 | 78.67 | 140.99 | 171.74 | 42.10 | 301.19 | 373.55 | 1132.91 | 573.30 | 783.75 |
| ENSBTAG00000048544 | | 52.39 | 85.28 | 72.61 | 18.30 | 59.62 | 306.46 | 159.03 | 113.63 | 105.32 | 166.72 | 146.75 | 158.58 | 293.70 | 397.67 | 446.10 | 496.10 |
| ENSBTAG00000048635 | | 2.02 | 1.02 | 6.15 | 3.23 | 0.99 | 1.62 | 0.81 | 4.99 | 4.25 | 5.02 | 4.81 | 3.42 | 13.19 | 26.59 | 42.69 | 45.51 |
| ENSBTAG00000049111 | | 116.88 | 53.81 | 91.07 | 83.96 | 107.31 | 107.02 | 115.22 | 114.88 | 90.03 | 145.63 | 179.23 | 100.40 | 174.97 | 331.78 | 320.63 | 304.94 |
| ENSBTAG00000052473 | | 429.22 | 272.09 | 979.64 | 304.61 | 329.89 | 1692.83 | 870.61 | 674.30 | 1063.34 | 1150.99 | 1321.94 | 1273.20 | 2681.52 | 5034.51 | 2980.66 | 3898.70 |
| ENSBTAG00000052673 | | 89.67 | 117.77 | 111.99 | 284.16 | 178.86 | 243.22 | 201.22 | 162.33 | 136.74 | 197.86 | 174.41 | 258.97 | 340.22 | 490.16 | 383.36 | 449.68 |
| ENSBTAG00000053278 | | 784.90 | 268.03 | 1335.32 | 240.03 | 310.02 | 1777.14 | 703.46 | 1078.87 | 1746.19 | 1493.47 | 905.75 | 1213.87 | 3188.38 | 5628.71 | 5698.19 | 4287.39 |
| ENSBTAG00000053612 | | 142.07 | 25.38 | 109.53 | 44.13 | 53.66 | 85.94 | 103.04 | 62.43 | 175.81 | 78.34 | 49.32 | 101.54 | 335.36 | 84.39 | 449.58 | 679.06 |
| ENSBTAG00000053831 | | 53.40 | 37.56 | 70.15 | 5.38 | 23.85 | 114.31 | 62.48 | 44.95 | 45.86 | 112.49 | 91.42 | 132.34 | 120.12 | 426.58 | 676.12 | 473.34 |
| ENSBTAG00000053893 | | 498.75 | 181.73 | 674.43 | 132.39 | 182.83 | 1543.65 | 487.64 | 690.53 | 1811.59 | 1854.03 | 732.54 | 1192.20 | 5685.20 | 4277.31 | 6288.92 | 5036.55 |
| ENSBTAG00000054147 | | 39.30 | 14.21 | 82.46 | 15.07 | 5.96 | 14.59 | 36.51 | 26.22 | 16.14 | 12.05 | 63.75 | 19.39 | 47.21 | 45.09 | 247.44 | 1541.09 |
| ENSBTAG00000054813 | | 152.14 | 58.89 | 180.91 | 71.04 | 88.43 | 279.71 | 120.90 | 164.83 | 225.92 | 176.77 | 116.68 | 175.69 | 411.05 | 698.24 | 700.51 | 601.69 |
| ENSBTAG00000055205 | | 96.73 | 50.76 | 83.69 | 97.95 | 88.43 | 117.56 | 77.08 | 66.18 | 106.16 | 132.57 | 90.21 | 86.71 | 174.97 | 189.59 | 301.46 | 266.71 |
