## Supplementary material 9: Gene overlap list par pathway for "Bovine in vitro blastocysts with distinct morphokinetic patterns show transcriptomic differences at genome activation"

| Ontology | Term order | Term | Overlap | Adjusted P-value | Odds Ratio | Genes |
|---|---|---|---|---|---|---|
| Go Biological Process 2025 | 1 | **Regulation of Transcription by RNA Polymerase II (GO:0006357)** | 132/2250 | 3.1E-03 | 1.63 | SPI1;HDAC10;FOXI3;UBE2D1;RORA;PRDM1;AHR;IKZF2;NR3C2;SALL2;MYB;ZXDC;DPF1;MYOZ1;SOX5;HES7;EOMES;HES6;ZNF18;BSX;LMO3;USP2;EBF1;MRTFA;HNF1B;LMO7;FOXP3;HCFC1;DMTF1;MAGEB3;NOBOX;ZNF436;ZNF398;HOMEZ;DLX6;SHOX2;TCF7;GATA2;FOXO3;FOXO1;NPAS3;SBNO2;ATXN1;ABL1;PPARGC1A;TFAP2B;ZFHX2;EGR4;ZBTB16;MICAL2;NFATC1;MAFA;SMARCA2;SMAD5;BMP6;ESR2;PHOX2A;SOX30;FOSL2;FOSL1;ID3;LHX4;BMPR1B;MYF5;BARHL2;PHF2;TENM2;ONECUT2;THRA;SIX1;HOXC13;IRF2BPL;BCL7A;ZNF529;HEY2;SIX3;DACT1;HAVCR2;BRD3;KLF11;ZHX2;ZNF362;PAX3;OLIG1;ETV1;PAX6;ETV5;ETV6;TOX3;ELF3;IRF4;IRF2;TET3;RARA;IRF8;CREB5;HLF;UHRF1;LEF1;DBX1;WIZ;HOXD13;BARX2;LRP6;GBX2;ZNF629;DNAJB4;MXI1;NOTO;MSX1;TAF9B;HES2;JAZF1;SNCA;ZNF462;FOXJ3;IRX4;DAB2IP;NR1H3;GRHL3;TBX18;POU4F3;PER2;MYO1E;PER1;TBX15;KLF7;ZNF71;NFIA;BCL6B;INSM1;HSPA1A |
| | 2 | **Positive Regulation of DNA-templated Transcription (GO:0045893)** | 83/1274 | 4.5E-03 | 1.79 | FANK1;SPI1;ONECUT2;THRA;BNC1;RASL11A;SIX1;RORA;HOXC13;AHR;IRF2BPL;SALL2;HEY2;MYB;ZXDC;KCNH2;EOMES;BRD3;HES6;LMO3;MRTFA;PAX3;DYRK1B;HNF1B;OLIG1;ETV1;PAX6;LMO7;FOXP3;ETV5;HCFC1;TOX3;ELF3;IRF4;IRF2;TET3;RARA;IRF8;CREB5;ZNF398;HLF;UHRF1;LEF1;HOXD13;GATA2;FOXO3;BARX2;FOXO1;NPAS3;LRP6;CAND2;SBNO2;ABL1;PPARGC1A;TRIM44;ZNF462;TFAP2B;EGR4;ZBTB16;FOXJ3;IRX4;DAB2IP;MICAL2;NR1H3;NFATC1;GRHL3;MAFA;SMARCA2;SMAD5;BMP6;ESR2;PHOX2A;SOX30;FOSL2;POU4F3;FOSL1;MYO1E;PER1;KLF7;NFIA;BCL3;LHX4;BMPR1B |
| | 3 | **Positive Regulation of Transcription by RNA Polymerase II (GO:0045944)** | 67/983 | 5.2E-03 | 1.87 | SPI1;ONECUT2;THRA;SIX1;RORA;AHR;IRF2BPL;SALL2;HEY2;MYB;EOMES;BRD3;HES6;LMO3;MRTFA;PAX3;HNF1B;OLIG1;ETV1;PAX6;LMO7;FOXP3;ETV5;HCFC1;TOX3;ELF3;IRF4;IRF2;TET3;RARA;IRF8;HLF;UHRF1;LEF1;HOXD13;GATA2;FOXO3;BARX2;FOXO1;LRP6;SBNO2;ABL1;PPARGC1A;ZNF462;TFAP2B;EGR4;FOXJ3;DAB2IP;MICAL2;NR1H3;NFATC1;GRHL3;MAFA;SMARCA2;SMAD5;BMP6;ESR2;PHOX2A;SOX30;FOSL2;POU4F3;MYO1E;PER1;KLF7;NFIA;LHX4;BMPR1B |
| | 4 | **Regulation of DNA-templated Transcription (GO:0006355)** | 123/2139 | 5.2E-03 | 1.59 | SPI1;HDAC10;FOXI3;RORA;PRDM1;AHR;IKZF2;NR3C2;SALL2;MYB;ZXDC;DPF1;SOX5;HES7;KCNH2;EOMES;MAGI1;HES6;ZNF18;BSX;EBF1;DYRK1B;HNF1B;FOXP3;HCFC1;DMTF1;NOBOX;ZNF436;ZNF398;HOMEZ;CALCA;DLX6;SHOX2;TCF7;FOXO3;FOXO1;NPAS3;SBNO2;ATXN1;PPARGC1A;TFAP2B;ZFHX2;EGR4;ZBTB16;NFATC1;FOXN3;MAFA;SMARCA2;SMAD5;ESR2;PHOX2A;SOX30;FOSL2;FOSL1;ZFAT;BCL3;ID3;LHX4;MYF5;FANK1;BARHL2;PHF2;ONECUT2;BNC2;THRA;BNC1;SIX1;HOXC13;ELAVL2;BCL7A;ZNF529;HEY2;SIX3;HAVCR2;BRD3;KLF11;ZHX2;ZNF362;PAX3;ETV1;PAX6;ETV5;ETV6;TOX3;ELF3;IRF4;IRF2;RARA;IRF8;VGLL4;CREB5;USP13;HLF;LEF1;DBX1;WIZ;HOXD13;BARX2;LRP6;CIART;CAND2;GBX2;ZNF629;MXI1;NOTO;PRAME;MSX1;HES2;TRIM44;FOXJ3;IRX4;DAB2IP;NR1H3;GRHL3;WTIP;TBX18;POU4F3;PER2;PER1;TBX15;KLF7;ZNF71;NFIA |
| | 5 | **Regulation of Nervous System Development (GO:0051960)** | 9/45 | 2.8E-02 | 6.19 | HES7;PER2;HES6;TENM4;GBX2;LEF1;HEY2;HES2;WNT3 |
| | 6 | **Negative Regulation of DNA-templated Transcription (GO:0045892)** | 64/1006 | 4.0E-02 | 1.72 | TENM2;SPI1;HDAC10;THRA;UBE2D1;PRDM1;AHR;IRF2BPL;BCL7A;HEY2;MYB;SIX3;MYOZ1;DACT1;EOMES;MAGI1;KLF11;ZHX2;USP2;PAX6;FOXP3;ETV5;ETV6;HCFC1;ELF3;IRF2;RARA;IRF8;MAGEB3;VGLL4;CALCA;UHRF1;SHOX2;LEF1;GATA2;FOXO3;FOXO1;CIART;SBNO2;ATXN1;DNAJB4;MXI1;PRAME;TAF9B;JAZF1;SNCA;TFAP2B;ZBTB16;DAB2IP;NR1H3;FOXN3;SMARCA2;BMP6;TBX18;ESR2;SOX30;PER2;PER1;KLF7;BCL6B;BCL3;ID3;INSM1;HSPA1A |
| | 7 | **Negative Regulation of Transcription by RNA Polymerase II (GO:0000122)** | 50/732 | 4.0E-02 | 1.85 | TENM2;SPI1;HDAC10;THRA;UBE2D1;PRDM1;IRF2BPL;MYB;HEY2;MYOZ1;DACT1;EOMES;KLF11;ZHX2;USP2;PAX6;ETV5;HCFC1;ETV6;IRF2;RARA;MAGEB3;IRF8;UHRF1;SHOX2;LEF1;GATA2;FOXO3;ATXN1;DNAJB4;MXI1;TAF9B;JAZF1;SNCA;TFAP2B;ZBTB16;DAB2IP;NR1H3;SMARCA2;BMP6;ESR2;TBX18;SOX30;PER2;KLF7;PER1;BCL6B;ID3;INSM1;HSPA1A |

| | | | | | | |
|---|---|---|---|---|---|---|
| **Go Molecular Fonction 2025** | 8 | **Sequence-Specific Double-Stranded DNA Binding (GO:1990837)** | 64/706 | 2.0E-07 | 2.57 | BARHL2;SPI1;ONECUT2;THRA;BNC1;SIX1;HOXC13;AHR;PRDM1;NR3C2;HEY2;DPF1;SIX3;SOX5;HES7;KCNH2;EOMES;HES6;KLF11;BSX;PAX3;PAX6;ETV1;FOXP3;ETV5;ELF3;IRF4;IRF2;RARA;IRF8;CREB5;HLF;DLX6;SHOX2;TCF7;LEF1;HOXD13;GATA2;FOXO3;BARX2;GBX2;NOTO;MSX1;HES2;SNCA;TFAP2B;EGR4;FOXJ3;NR1H3;UNCX;NFATC1;MAFA;SMARCA2;SMAD5;TBX18;SOX30;POU4F3;PHOX2A;FOSL1;PER2;PER1;TBX15;BCL6B;LHX4 |
| | 9 | **Sequence-Specific DNA Binding (GO:0043565)** | 64/718 | 2.0E-07 | 2.52 | BARHL2;SPI1;ONECUT2;SIX1;RORA;HOXC13;AHR;PRDM1;NR3C2;ZCWPW1;HEY2;DPF1;SIX3;HES7;EOMES;HES6;KLF11;BSX;PAX3;PAX6;ETV1;FOXP3;ETV5;ELF3;IRF4;TET3;IRF2;RARA;IRF8;NOBOX;CREB5;HLF;UHRF1;DLX6;SHOX2;TCF7;LEF1;HOXD13;GATA2;FOXO3;BARX2;FOXO1;GBX2;HLX;NOTO;MSX1;PPARGC1A;HES2;TFAP2B;EGR4;FOXJ3;UNCX;NFATC1;GRHL3;MAFA;SMAD5;TBX18;SOX30;POU4F3;PHOX2A;FOSL1;TBX15;BCL6B;LHX4 |
| | 10 | **Double-Stranded DNA Binding (GO:0003690)** | 57/651 | 2.3E-06 | 2.46 | BARHL2;ONECUT2;SIX1;HOXC13;AHR;PRDM1;NR3C2;HEY2;DPF1;SIX3;HES7;EOMES;HES6;KLF11;BSX;PAX3;PAX6;ETV1;FOXP3;ETV5;ELF3;IRF4;IRF2;RARA;IRF8;CREB5;HLF;UHRF1;DLX6;SHOX2;TCF7;LEF1;HOXD13;GATA2;FOXO3;BARX2;GBX2;NOTO;MSX1;NLRP1;HES2;TFAP2B;EGR4;FOXJ3;UNCX;NFATC1;MAFA;SMAD5;TBX18;SOX30;POU4F3;PHOX2A;FOSL1;TBX15;BCL6B;LHX4;ZCCHC3 |
| | 11 | **RNA Polymerase II Transcription Regulatory Region Sequence-Specific DNA Binding (GO:0000977)** | 85/1236 | 3.1E-05 | 1.91 | BARHL2;SPI1;ONECUT2;THRA;FOXI3;SIX1;RORA;HOXC13;PRDM1;IKZF2;SALL2;ZNF529;HEY2;MYB;SIX3;SOX5;HES7;EOMES;HES6;ZNF18;KLF11;ZNF362;BSX;EBF1;PAX3;HNF1B;ETV1;PAX6;FOXP3;ETV5;ETV6;DMTF1;IRF4;IRF2;TET3;RARA;IRF8;NOBOX;ZNF436;ZNF398;HLF;DLX6;TCF7;LEF1;WIZ;HOXD13;GATA2;FOXO3;BARX2;FOXO1;NPAS3;CIART;GBX2;MXI1;NOTO;MSX1;HES2;TFAP2B;ZFHX2;EGR4;ZBTB16;FOXJ3;IRX4;NR1H3;NFATC1;GRHL3;MAFA;SMAD5;TBX18;ESR2;PHOX2A;SOX30;FOSL2;POU4F3;FOSL1;PER1;TBX15;KLF7;ZNF71;NFIA;ZFAT;BCL6B;LHX4;INSM1;MYF5 |
| | 12 | **DNA-binding Transcription Activator Activity, RNA Polymerase II-specific (GO:0001228)** | 33/318 | 3.1E-05 | 2.92 | HLF;ONECUT2;LEF1;SIX1;HOXD13;FOXO3;BARX2;FOXO1;SALL2;MYB;SIX3;TFAP2B;EGR4;ZBTB16;FOXJ3;IRX4;NFATC1;PAX6;GRHL3;ETV1;FOXP3;ETV5;FOSL2;SOX30;POU4F3;PHOX2A;KLF7;NFIA;ZNF71;ELF3;IRF4;IRF2;LHX4 |
| | 13 | **RNA Polymerase II Cis-Regulatory Region Sequence-Specific DNA Binding (GO:0000978)** | 75/1054 | 3.1E-05 | 1.97 | SPI1;ONECUT2;THRA;FOXI3;SIX1;RORA;HOXC13;PRDM1;IKZF2;NR3C2;HEY2;MYB;SIX3;SOX5;HES7;EOMES;HES6;ZNF18;KLF11;ZNF362;BSX;EBF1;PAX3;HNF1B;OLIG1;ETV1;PAX6;FOXP3;DMTF1;IRF4;IRF2;TET3;RARA;IRF8;NOBOX;CREB5;HLF;DLX6;TCF7;LEF1;WIZ;HOXD13;GATA2;FOXO3;FOXO1;CIART;MXI1;NOTO;HES2;TFAP2B;ZFHX2;EGR4;ZBTB16;FOXJ3;IRX4;NR1H3;NFATC1;GRHL3;MAFA;SMAD5;TBX18;ESR2;SOX30;FOSL2;POU4F3;FOSL1;PER1;TBX15;KLF7;ZNF71;NFIA;ZFAT;BCL6B;INSM1;MYF5 |
| | 14 | **Cis-Regulatory Region Sequence-Specific DNA Binding (GO:0000987)** | 74/1035 | 3.1E-05 | 1.98 | SPI1;ONECUT2;THRA;FOXI3;SIX1;RORA;HOXC13;PRDM1;IKZF2;HEY2;MYB;SIX3;SOX5;HES7;EOMES;HES6;ZNF18;KLF11;ZNF362;BSX;EBF1;PAX3;HNF1B;ETV1;PAX6;FOXP3;DMTF1;IRF4;IRF2;TET3;RARA;IRF8;NOBOX;HLF;UHRF1;DLX6;TCF7;LEF1;WIZ;HOXD13;GATA2;FOXO3;FOXO1;CIART;MXI1;NOTO;HES2;TFAP2B;ZFHX2;EGR4;ZBTB16;FOXJ3;IRX4;NR1H3;NFATC1;GRHL3;FOXN3;MAFA;SMAD5;TBX18;ESR2;SOX30;FOSL2;POU4F3;FOSL1;PER1;TBX15;KLF7;ZNF71;NFIA;ZFAT;BCL6B;INSM1;MYF5 |
| | 15 | **Transcription Cis-Regulatory Region Binding (GO:0000976)** | 39/548 | 1.7E-02 | 1.92 | BARHL2;SPI1;THRA;UHRF1;TCF7;LEF1;SIX1;RORA;HOXC13;AHR;FOXO3;BARX2;NPAS3;GBX2;SALL2;ZNF529;HEY2;MXI1;MSX1;SOX5;SNCA;KCNH2;TFAP2B;KLF11;NR1H3;PAX6;FOXN3;SMARCA2;ETV5;FOSL2;PHOX2A;ETV6;PER2;PER1;IRF2;RARA;ZNF436;LHX4;ZNF398 |
| | 16 | **Actin Binding (GO:0003779)** | 18/187 | 2.5E-02 | 2.65 | MRTFA;MICAL2;CXCR4;SSH2;CORO2B;ADD1;ADD2;CNN2;ABLIM1;MICAL1;ABL1;EPS8L1;MYOZ1;MYH10;FERMT2;CDK5R2;PFN2;SNCA |
| | 17 | **Transcription Coregulator Binding (GO:0001221)** | 12/103 | 3.7E-02 | 3.27 | PER2;PER1;ZBTB16;SIX3;LEF1;RARA;WIZ;RORA;HNF1B;PAX6;GATA2;VGLL4 |

| | | | | | | |
|---|---|---|---|---|---|---|
| MGI Mammalian Phenotype Level 4 2024 | 18 | **Neonatal Lethality, Complete Penetrance MP:0011087** | 45/521 | 1.1E-03 | 2.40 | CYFIP2;TOP2B;SPAG17;SPI1;BNC2;DLX6;FOXI3;DBX1;CXCR4;LPL;JPH1;SLC7A1;NALCN;LRP6;IGF1R;HHAT;GBX2;TUBB3;ADGRA2;PODXL;HEY2;SPOP;GPC3;SFN;MSX1;MYH10;SOX5;HES7;TFAP2B;SLC12A5;ELOVL4;PAX6;TBX18;SULF2;CLIP3;TBX15;VCAN;NBEA;SMOC1;SLCO2A1;TACC3;TCF4;INSM1;MYF5;CREB5 |
| | 19 | **Enlarged Spleen MP:0000691** | 50/632 | 2.3E-03 | 2.18 | PCSK1;CDKN1A;SPI1;CDCA7L;TNFAIP3;SIX1;AHR;ITGAL;SPRED2;STK11;RGS2;MYB;CHAMP1;FBRSL1;FAM81A;PRKG1;PABPC5;EBF1;CACNA2D2;BPGM;SSTR2;FOXP3;VCAN;IRF4;IRF8;CCNO;EPB41;CXCR4;ADRB2;FOXO3;NALCN;RASGRP1;ADD2;NLRP9;MXI1;RFFL;KCNN4;EXTL3;CCL25;RBM38;TCAF2;SLC12A5;PRSS33;ESR2;GGCT;FER;ZFAT;ID3;FAS;TMCC2 |
| | 20 | **Increased Spleen Weight MP:0004952** | 24/224 | 5.1E-03 | 3.00 | CCL25;PPP1R15A;CDKN1A;SPI1;EPB42;CACNA2D2;PRDM13;AHR;FOXP3;RASGRP1;ETV6;ADD2;FER;MARVELD2;IRF4;SCN8A;BCL3;ID3;FAS;IRF8;KCNN4;DOCK2;HAVCR2;SNCA |
| | 21 | **Abnormal Cardiac Outflow Tract Development MP:0006126** | 11/56 | 5.1E-03 | 6.06 | KCNH2;ADAM19;IFT172;FGF19;ELP1;SIX1;MRTFA;PAX3;HAS2;NFATC1;MYH10 |
| | 22 | **Decreased Startle Reflex MP:0001489** | 22/196 | 5.1E-03 | 3.16 | CYFIP2;PLEKHF2;THRA;TMOD2;PALM3;NEDD4L;DYRK1B;FOXO3;FBXO30;BMP6;POU4F3;TMEM145;PEX5L;MARVELD2;SCN8A;CMYA5;TCF4;CSMD1;FAM81A;MPDZ;CUEDC1;ASIC1 |
| | 23 | **No Abnormal Phenotype Detected MP:0002169** | 112/1935 | 7.3E-03 | 1.59 | CYFIP2;ERRFI1;RAB3C;SPI1;FOXI3;MSI1;PRDM1;ADARB1;RIMS2;STK11;RGS2;FRAT2;SALL2;ADGRA2;MYB;PIM3;UBASH3A;SOX5;EOMES;MAGI1;BSX;LMO3;CACNA2D2;FBXO10;FOXP3;GTPBP1;ERN1;NEIL2;COL4A6;FKBP5;CALCA;GATA2;CACNA1E;FOXO1;KLK7;IGSF9;PODXL;ABL1;TAC3;KIF3C;MYH10;CDR2;CDT1;SMURF1;MAFA;CORO2B;SMAD5;ESR2;VANGL1;EXT1;FOSL1;PEX5L;FER;DAB1;NOXA1;CYP1A1;ID3;IL7R;BMPR1B;FERMT2;MYF5;FANK1;SPAG17;TOP2B;PRSS27;FMN1;CROT;CCR8;PRKG1;KLF11;PARP4;HUNK;PAX3;ETV1;PAX6;ETV6;VCAN;CDHR1;PITPNM2;IRF2;RARA;IRF8;PPP1R15A;RALB;HLF;NRXN1;SEZ6;DBX1;NRXN3;COCH;LRP6;CACNG7;ABLIM1;GBX2;LRIG1;HAS2;GCNT3;MSX1;TAF9B;RFFL;SNCA;PCDH8;NR1H3;GRHL3;OOSP2;POU4F3;PER2;SYT11;TACC3;ESYT3;FAS;CDK5R2 |
| | 24 | **Increased NK Cell Number MP:0008044** | 9/41 | 9.6E-03 | 6.96 | KCTD9;UHRF1;CXCR4;RRAS2;FAS;FOXO3;ADAP1;DOCK2;RASGRP1 |
| | 25 | **Abnormal Spleen Morphology MP:0000689** | 44/589 | 1.2E-02 | 2.04 | TOP2B;CDKN1A;SPI1;BNC1;CDCA7L;MGST2;CXCR4;SIX1;PRDM13;ADAR;AHR;ADRB2;NALCN;SPRED2;MYB;ABL1;CHAMP1;SFN;RFFL;KCNN4;EXTL3;FAM81A;PLXNA4;KL;TCAF2;SLC12A5;EPHA6;ZBTB16;PABPC5;EBF1;CACNA2D2;PRSS33;SSTR2;FOXP3;SELP;FER;VCAN;FGF14;IRF4;ZFAT;ID3;TMCC2;DOCK2;CCNO |
| | 26 | **Decreased Body Size MP:0001265** | 60/917 | 1.9E-02 | 1.78 | RERE;PCSK1;TOP2B;SPI1;TENM4;TNFAIP3;SIX1;RORA;ADAR;HOXC13;RIMS2;GRIP1;IBSP;CA2;HEY2;MYB;KRT25;CSGALNACT1;CASR;ABCG5;BSX;EBF1;CACNA2D2;PAX3;ETV1;PAX6;FOXP3;HCFC1;ELF3;SCN8A;VGLL4;CAMK2B;HACD1;SEMA3A;LEF1;CXCR4;SEMA3F;LTBP3;GATA2;AGPAT1;SLC7A1;ADD1;NPAS3;LRP6;ABL1;KL;SLC12A5;FOXN3;BICC1;BMP6;FOSL2;SULF2;POU4F3;SLC6A5;TBX15;DAB1;NFIA;NBEA;SMOC1;TMCC2 |
| | 27 | **Abnormal Muscle Precursor Cell Migration MP:0003090** | 4/7 | 1.9E-02 | 32.84 | SIX1;CXCR4;PAX3;MYF5 |
| | 28 | **Forelimb Oligodactyly MP:0014280** | 4/7 | 1.9E-02 | 32.84 | FMN1;PRDM1;BAIAP2;LRP6 |
| | 29 | **Fused Metatarsal Bones MP:0004642** | 4/7 | 1.9E-02 | 32.84 | ZBTB16;SMOC1;FMN1;BMPR1B |
| | 30 | **Abnormal Erythrocyte Morphology MP:0002447** | 12/84 | 2.4E-02 | 4.13 | PPP1R15A;RBM38;GGCT;ELOVL4;EPB42;EPB41;MYB;TMCC2;FOXP3;PRKG1;ADD1;ADD2 |
| | 31 | **Abnormal Retina Morphology MP:0001325** | 35/456 | 2.6E-02 | 2.09 | HACD1;TENM2;NEDD4L;SIX1;WIZ;ADRB2;FOXO3;FAM117B;RASGRP4;TMEM145;SPRED2;PODXL;FBRSL1;EPS8L1;GJA8;MPDZ;MYH10;PROM2;KL;ELOVL4;LIMCH1;TBC1D8;DYRK1B;PAX6;GRHL3;ARHGAP25;PER2;FER;FAM161A;SMOC1;CTH;MAB21L1;CDHR1;TCF4;PKIG |
| | 32 | **Abnormal Estrous Cycle MP:0001927** | 8/40 | 2.6E-02 | 6.18 | PER1;TNFAIP3;TACR3;RORA;TAC3;FOXO3;AGPAT1;ESR2 |
| | 33 | **Abnormal Spatial Learning MP:0001463** | 18/171 | 2.6E-02 | 2.93 | SLC24A2;EPHA6;TMOD2;LRRN4;SEZ6;TACR3;RORA;BAIAP2;KIF17;ESR2;SLC8A2;ADD2;ARC;ATXN1;NBEA;FAS;SHANK2;ASIC1 |

| | | | | | | |
|---|---|---|---|---|---|---|
| | 34 | **Postnatal Lethality, Incomplete Penetrance MP:0011086** | 45/646 | 2.6E-02 | 1.89 | RERE;PHF2;ERRFI1;PCSK1;SPI1;SEMA3A;SHOX2;NRXN3;TNFAIP3;RORA;HOXC13;AHR;PRDM1;GATA2;AGPAT1;ELAVL2;ELK3;NPAS3;LRP6;BRINP1;HEY2;ABL1;GPC3;PPARGC1A;MYH10;HES7;PRKCH;CACNA2D2;PAX6;LMO7;FOXN3;SNAP91;ETV5;HCFC1;SULF2;ADAM19;SHTN1;GAP43;DAB1;NFIA;DMTF1;ELF3;SCN8A;CDO1;SHANK2 |
| | 35 | **Abnormal CD4-positive, Alpha Beta T Cell Morphology MP:0002432** | 7/31 | 2.7E-02 | 7.20 | CDKN1A;IRF4;GRAP2;BCL6B;DOCK2;RASGRP1;FOXP3 |
| | 36 | **Abnormal Miniature Excitatory Postsynaptic Currents MP:0004753** | 12/89 | 2.9E-02 | 3.86 | ARC;SLC12A5;NBEA;NRXN1;KCTD13;NRXN3;PPP1R9A;SHANK2;REM2;SNAP91;PFN2;SNCA |
| | 37 | **Reduced Female Fertility MP:0001923** | 24/272 | 2.9E-02 | 2.42 | NPM2;KHDC3L;THRA;CACNA2D2;TACR3;TNFAIP3;PAX6;RORA;NRARP;AHR;FOXO3;ESR2;SULF2;BMP15;MOS;PER2;PER1;TCL1A;WEE2;ASTL;NFIA;GAS2;KPNA7;TAC3 |
| | 38 | **Interdigital Webbing MP:0000571** | 5/15 | 2.9E-02 | 12.33 | GRIP1;ZBTB16;SMOC1;RARA;HOXD13 |
| | 39 | **Abnormal Cerebellum Morphology MP:0000849** | 13/105 | 3.3E-02 | 3.50 | RERE;CACNA2D2;CXCR4;RORA;PAX6;OLIG1;LRP6;OCLN;DAB1;GBX2;CMYA5;PPARGC1A;MYH10 |
