## Supplementary figures and images for "Bovine in vitro blastocysts with distinct morphokinetic patterns show transcriptomic differences at genome activation"

### Supplementary material 1 - Figure 1S

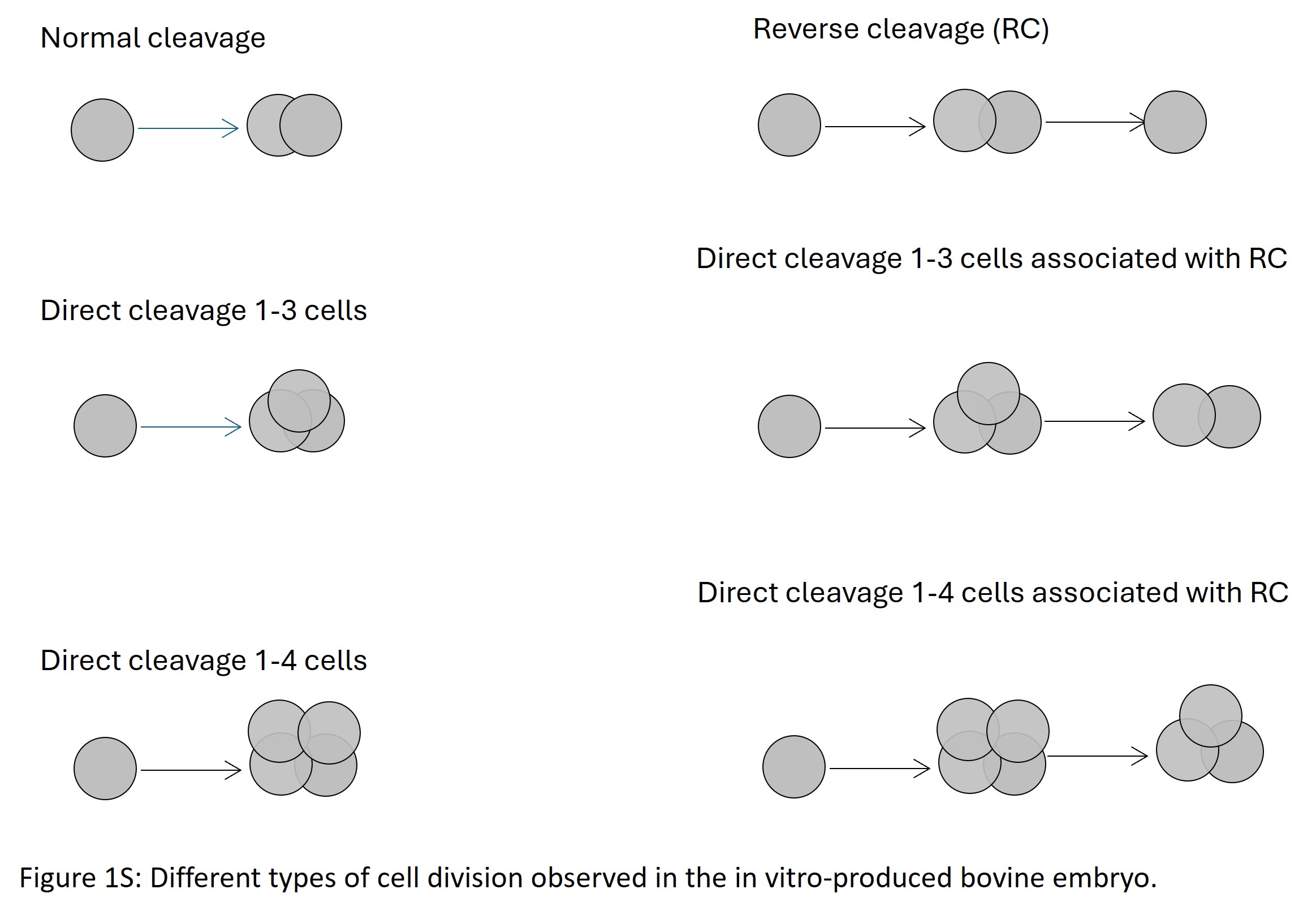

### Supplementary material 1 - Figure 2S

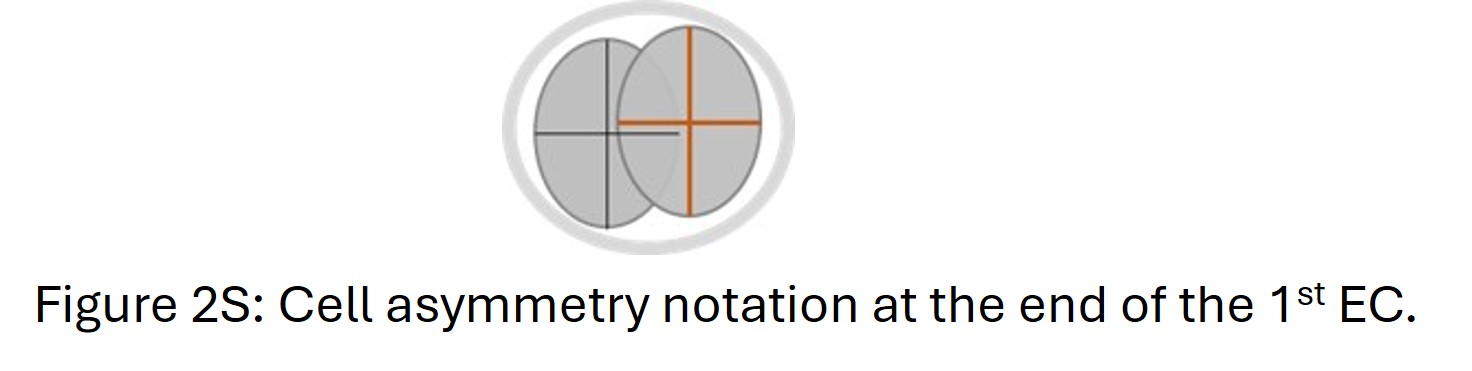

### Supplementary material 1 - Figure 3S

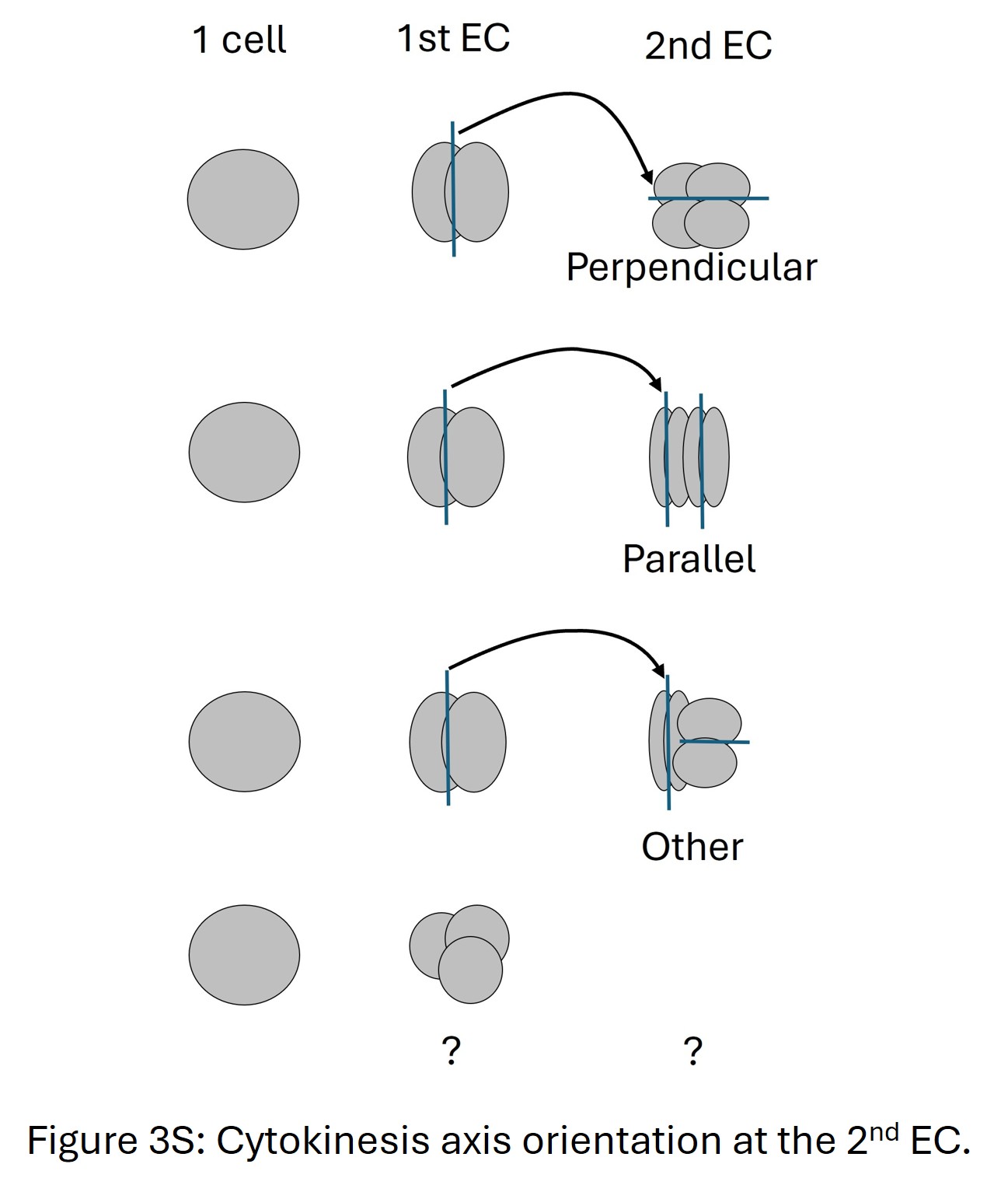

### Supplementary material 1 - Figure 4S

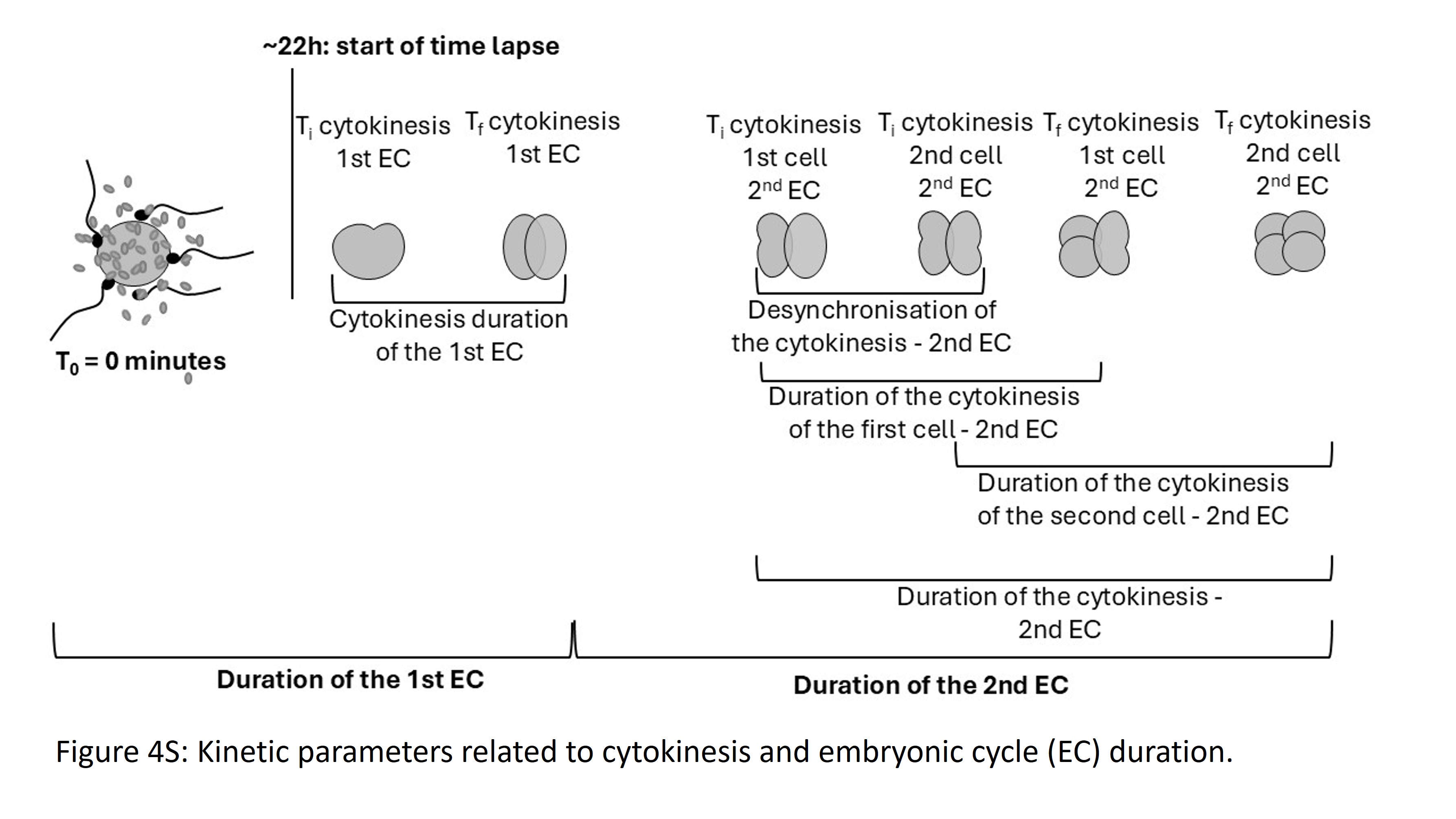

### Supplementary material 1 - Figure 5S

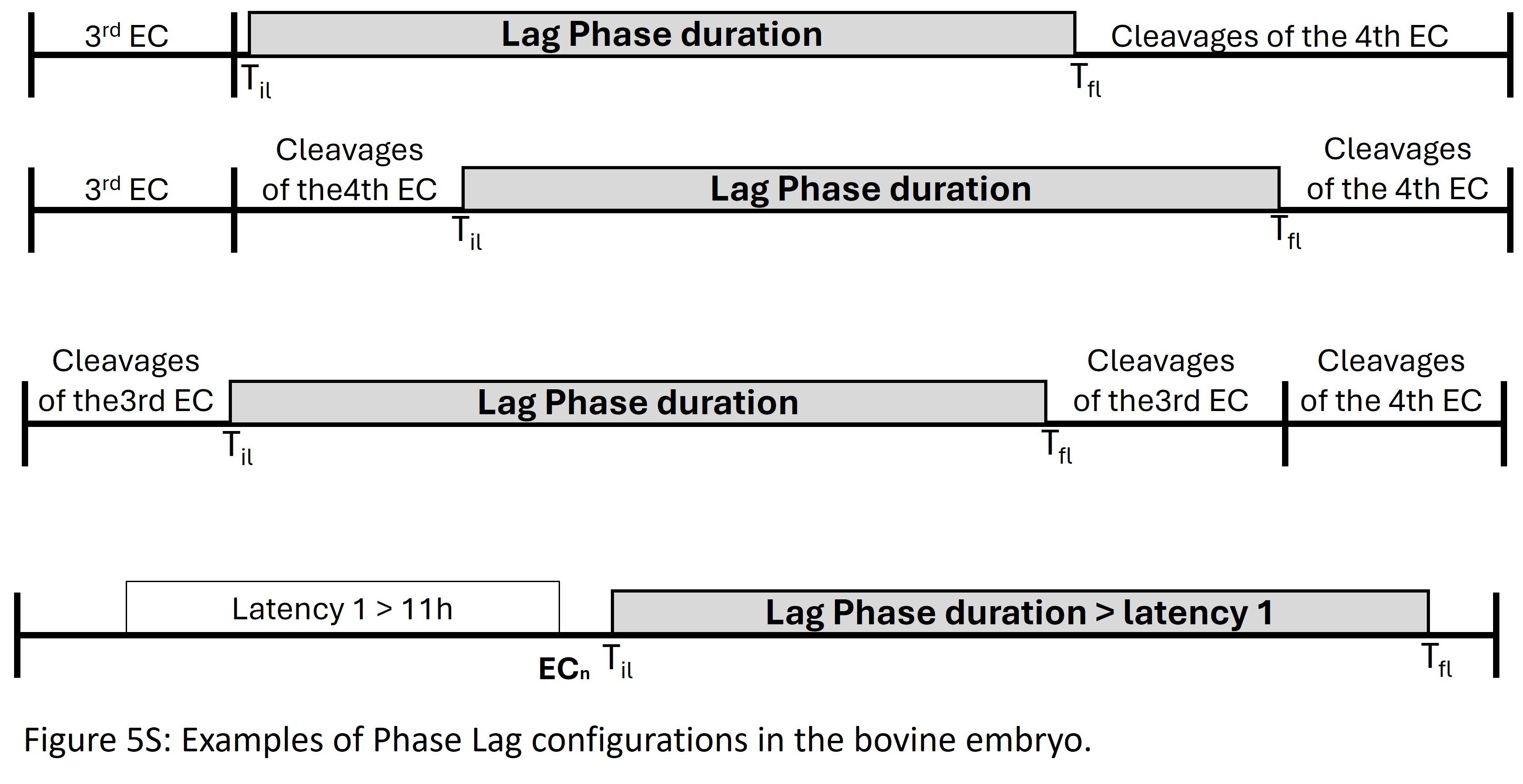
